# Un-LOK-ing a new approach for conformational selective targeting of STK10 (LOK)

**DOI:** 10.1101/2025.07.22.666149

**Authors:** Martina Dettenhöfer, Laura Nadine Tandara, Jennifer Alisa Amrhein, Christian Georg Kurz, Martin Peter Schwalm, Theresa Elisabeth Mensing, Laurenz Maximilian Wahl, Andreas Krämer, Joshua Gerninghaus, Christopher Lenz, Lewis Elson, Benedict-Tilman Berger, Martin Schröder, Krishna Saxena, Susanne Müller, Stefan Knapp, Francesco Aleksy Greco, Thomas Hanke

## Abstract

STK10 (serine/threonine kinase 10, LOK), is an important regulator of diverse cellular processes, such as cell cycle progression or lymphocyte migration. STK10 has emerged as a potential therapeutic target for diseases associated with impaired cell migration and cell division. Here we present a late-stage optimization of a macrocyclic pyrazolo[1,5-*a*]pyrimidine scaffold that led to a urea-based lead series targeting the back-pocket of STK10. Co-crystal structure analysis of **23** revealed that the optimized macrocycles adopted a unique binding mode that protrudes deep into the back pocket of STK10. Compound **23** exhibited potent on-target activity in biophysical and activity assays and displayed nanomolar activity for STK10 in cells. In addition, **23** shows good selectivity against the kinome and remarkably also against the closely related kinase SLK (STE20-like kinase). Therefore, we propose that targeting the unique and largely extended pocket in STK10 represents an opportunity to develop highly selective STK10 inhibitors.

**TOC:** 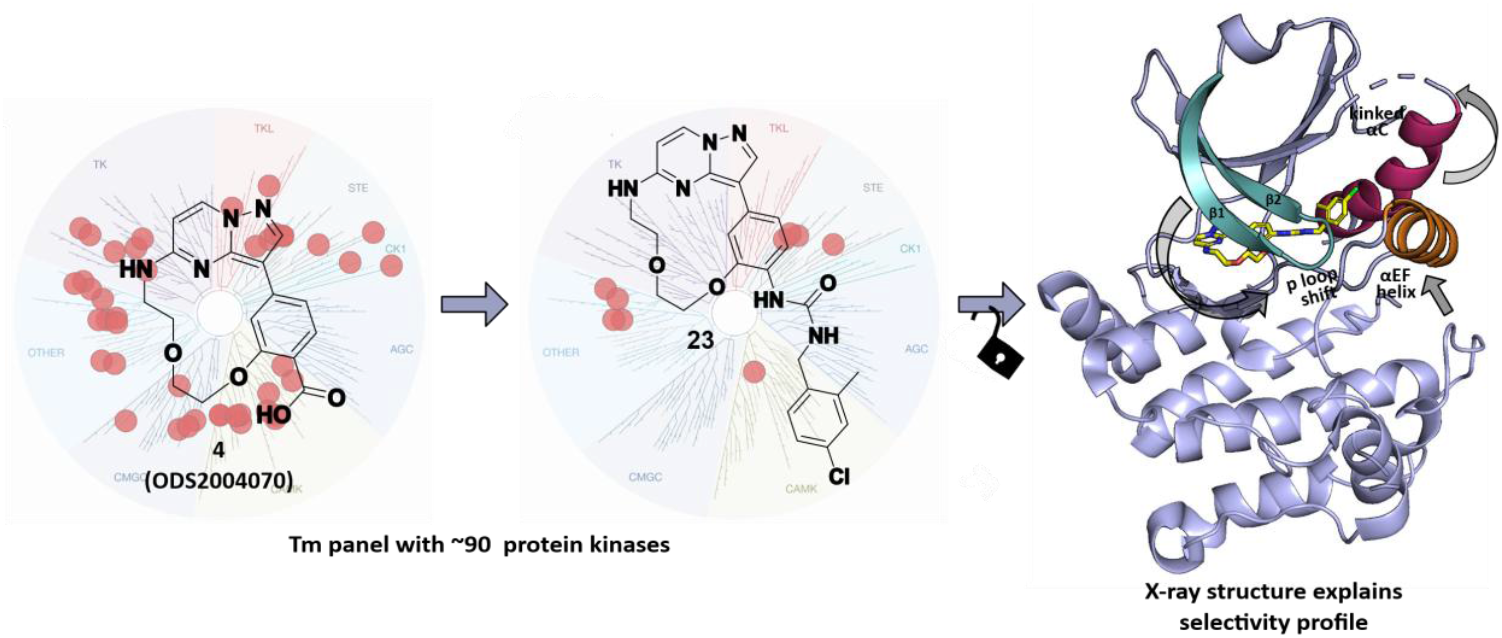

---

Serine/threonine kinase 10 (STK10) belongs to the STE20 superfamily of serine/threonine kinases, which is predominantly expressed in lymphoid tissues such as the spleen, bone marrow or thymus. The restricted expression pattern of STK10 coined its name Lymphocyte-Oriented Kinase (LOK).[1–3] STK10 shares high structural similarity with SLK (STE20-like kinase) and other STE20 family members. The common function of STK10 and SLK is the activation of ezrin/radixin/moesin (ERM) proteins by phosphorylation of a threonine residue at their N-terminal FERM domain.[4,5] ERM proteins are crucial for the structural stability and integrity of the cell by establishing links between the plasma membrane and actin filaments.[3,6] Aberrant activity of ERM proteins, or dysregulation of their phosphorylation by STK10 and SLK, leads to altered apical morphology in epithelial cells with loss of microvilli integrity.[4] These actin-rich cell protrusions are found e.g., on immune cells and serve as hubs for T-cell signalling, linking the function of the ERM protein to key components of the initial immune response.[7] However, deeper knowledge of STK10 functions in physiological and pathophysiological settings remains elusive, which can be partly attributed to the paucity of well profiled tool compounds.

It is known that a wide range of small molecule kinase inhibitors (SMKIs) can bind to STK10, including the FDA approved BCR-ABL/SRC inhibitor Bosutinib (**1**) and the c-MET/VEGFR-2 inhibitor Foretinib (**2**).[8–10] In addition, there are also several SMKIs that are not FDA-approved but known to be potent inhibitors of STK10. For example, the inhibitor SB-633825 is a commercially available, potent STK10 inhibitor that was originally developed as a TIE2 inhibitor (**Figure 1**).[11] In 2021, Serafim *et al*. developed a series of maleimide-containing compounds that are type I STK10-inhibitors, but also inhibit the closely related kinase SLK and are thus dual SLK/STK10 inhibitors with however moderate cellular activity (**Figure 1**).[12]

**Figure 1.**
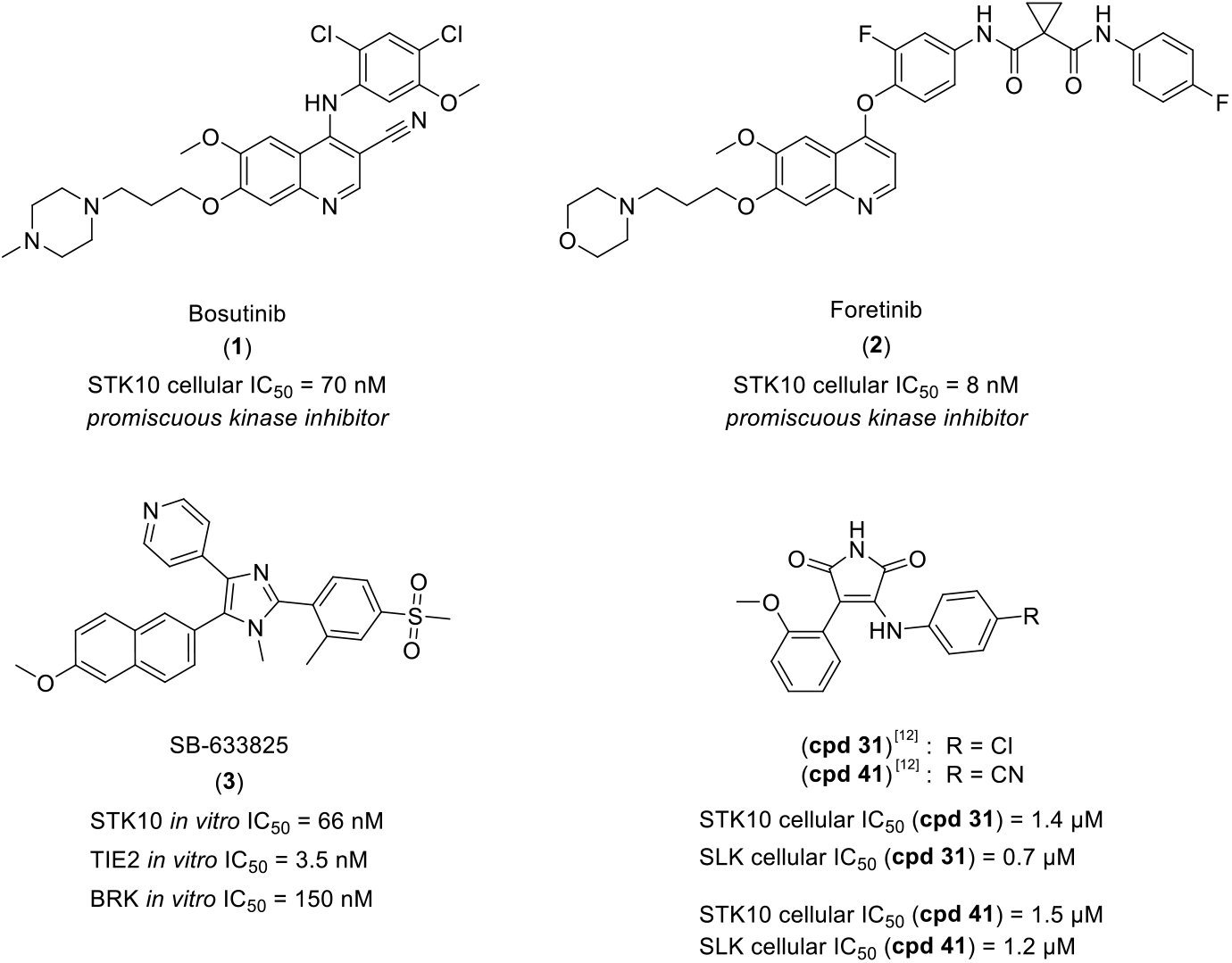
Chemical structures of highly potent and promiscuous kinase inhibitors Bosutinib (**1**) and Foretinib (**2**), SB-633825 (**3**)[11] and previous published dual STK10/SLK inhibitors **cpd 31**[12] (STK10 *in vitro* IC_50_ = 12 nM) and **cpd 41**[12] (STK10 *in vitro* IC_50_ = 23 nM) by Serafim *et al*.

This structural diversity of inhibitors indicates that STK10 is a kinase with a high degree of domain plasticity allowing it to accommodate different chemotypes that target the ATP pocket independent of the position of the DFG motif including type II (DFG-out) inhibitors.[13–15] Indeed, structural studies revealed different conformational states as well as the formation of transient dimers with domain-exchanged activation segments, a conformation that extends the ATP-binding site back pocket and plays an important role in kinase auto-activation at non-consensus sites located in the kinase activation segment (A-loop).[16,17] Thus, STK10 can accommodate highly diverse ligands and it is a frequent off-target of kinase inhibitors[11] including FDA approved and clinical inhibitors. However, potent and selective kinase inhibitors for STK10 are still lacking, in particular inhibitors that exhibit selectivity towards the closely related kinase SLK. In this study, we have modified macrocyclic pyrazolo[1,5-*a*]pyrimidines with the goal to target the back-pocket of kinases to achieve selectivity for STK10/SLK. By incorporating a urea motif, we have succeeded in addressing a pocket that has not been described in any STK10 co-crystal before and offers new avenues for selectively targeting this kinase.

The starting point for the development of our series of urea-based inhibitors was the promiscuous macrocyclic inhibitor **4** (ODS2004070), which is based on the widely used and well-established pyrazolo[1,5-*a*]pyrimidine scaffold (**Figure 2**).[18–20] The target profile of **4** (ODS2004070) includes kinases that can adopt both type I and type II binding modes which motivated us to design compounds that would allow to harness the conformational flexibility of these kinases. In our previous work we developed a series of amide-based macrocycles as part of a late-stage optimization strategy, identifying the chemical probe **CK156**, a selective inhibitor of DRAK1 (DAP kinase-related apoptosis-inducing protein kinase 1).[21] The favorable kinome wide selectivity profile of **CK156** can be attributed to its bulky tertiary amide group that points towards the back-pocket (**Figure 2**).

**Figure 2.**
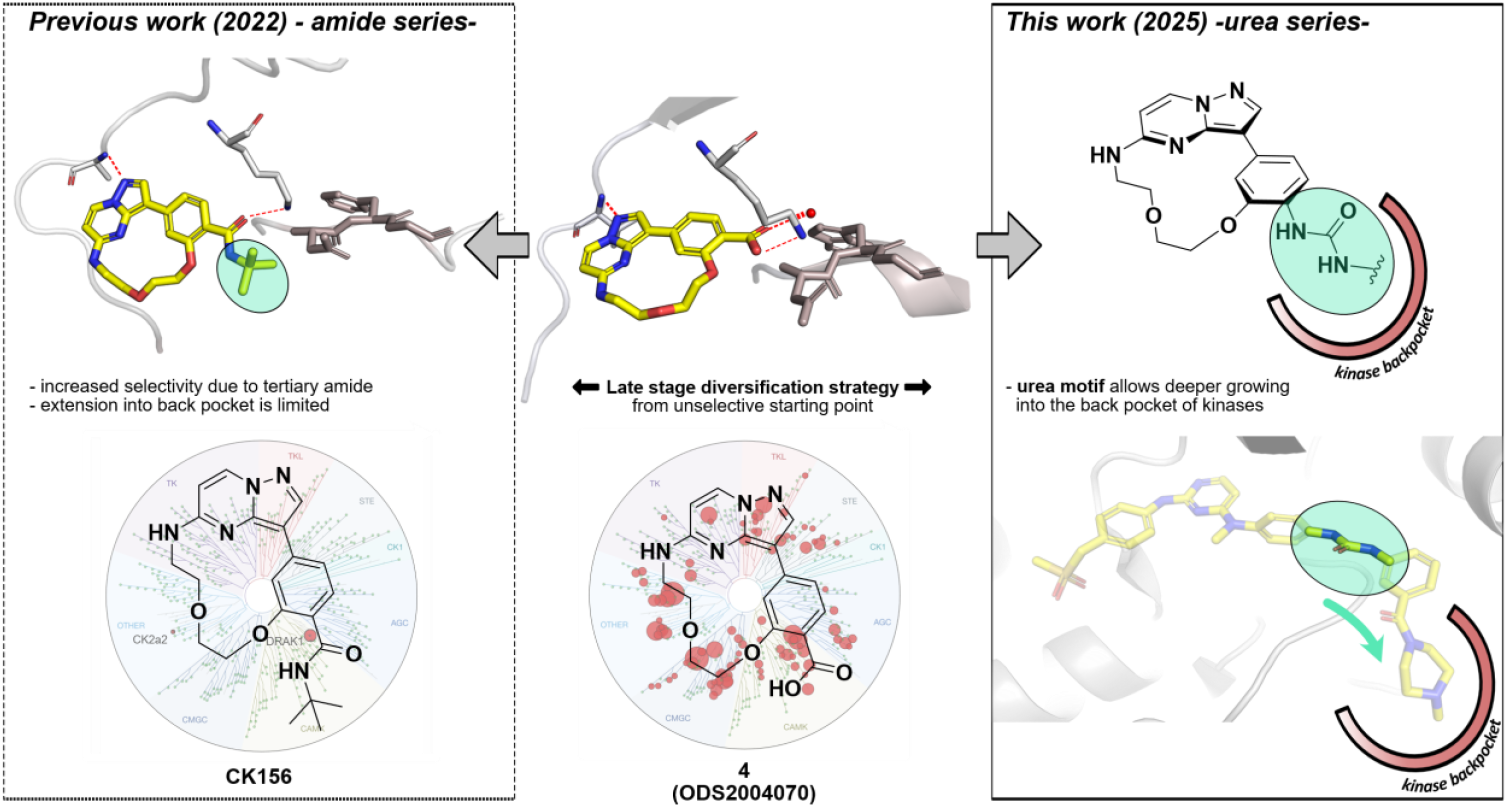
Design strategy. Previous work: Modification of the promiscuous pyrazolo[1,5-*a*]pyrimidine macrocyclic scaffold **4** (ODS2004070) (middle, PDB 6Z4Z) leads to **CK156**, a chemical probe for DRAK1 (left, PDB 7QUF).[21] This work: A “late-stage” modification strategy of the promiscuous pyrazolo[1,5-*a*]pyrimidine macrocyclic scaffold **4** (ODS2004070) by introduction of diverse back-pocket motifs, resulting in a series of 20 urea-based derivatives. The urea motif enables a deeper growing into the back-pocket of kinases (right, PDB 4AOT).

Based on our experience from the utilization of this scaffold[21,22] in conjunction with the synthesis of macrocyclic kinase inhibitors in general[23–25], the present study is oriented towards the exploration of the back-pocket interactions of compound **4** (**Figure 2**). Our optimization efforts led to the design and synthesis of a novel series of urea-based macrocycles intended to act as type II kinase inhibitors. The conformational changes in the kinase domain induced by a type II binding mode often result in altered selectivity profiles. In addition, the changes in protein confirmations upon binding of type II kinase inhibitors can alter not only catalytic but also scaffolding functions of protein kinases and can thus differentiate functionally from type I inhibitors.[26,27] The urea motif is a classical component found in a plethora of type II inhibitors (**Figure 2**, right panel).[28–31] Thus, we introduced it as linking moiety that positions the bulky benzylic moiety to protrude deep into the back pocket of the kinase, while the macrocyclic moiety occupies the ATP-binding site with shape complementary modulated by the linker used for cyclization.[32]

The macrocyclic scaffold **4** was synthesized in a 10-step synthetic route as outlined in **Scheme 1**.[22] Bromination of the commercially available 5-chloropyrazolo[1,5-*a*]pyrimidine **5** with NBS resulted in the brominated product **6** with a yield of 96%. The linker was introduced by a nucleophilic aromatic substitution of **6** with 2-(2-aminoethoxy)ethanol to obtain **7** with a yield of 80%. Subsequently, the hydroxy group of **7** was protected with TBDMS-Cl (87%) and the secondary amine with a Boc protecting group (94%), resulting in the intermediate **9**. The pinacolborane ester **11** was synthesized by Miyaura borylation from the commercially available **10**, followed by Suzuki cross-coupling between the intermediates **9** and **11** to obtain **12** with a yield of 84%. Cleavage of the silyl ether protecting group was carried out with TBAF to give **13** in 80% yield. The macrocyclic ring closure reaction was carried out by a Mitsunobu reaction under high dilution leading to macrocyclic compound **14** with a yield of 87%. Deprotection with TFA (63%) and hydrolysis with LiOH (87%) yielded the macrocyclic precursor **4**. The final urea derivatives **16**-**35** were synthesized by a Curtius rearrangement reaction using diphenylphosphoryl azide (DPPA) and triethylamine in the presence of various benzylamines, with yields ranging from 11% to 84%.

The synthesized urea-based derivatives were measured in a differential scanning fluorimetry (DSF) panel to investigate their selectivity profile. A positive ΔT_m_ in the DSF assay indicates stabilization of the protein upon ligand binding in comparison to its apo state, with higher thermal shifts generally correlating with stronger binding. For this purpose, we used our internal kinase selectivity panel (~90 kinases) along with staurosporine as a positive control. The urea series showed moderate to good binding (ΔT_m_: 4–15 °C in DSF) to various proteins of the CMGC, CAMK and STE kinase superfamilies, while most kinases of the TK and TKL class were no longer stabilized by the urea derivatives (**Figure S1**). Interestingly, the compounds showed good stabilization in the STE family especially with the closely related kinases STK10/SLK as well as with members of the Mammalian Sterile20-like (MST) family STK3/4 (also called MST2/MST1) and STK24/26 (also called MST3/MST4). When ranking the compounds according to their thermal stabilization, we noticed that STK10 showed high ΔT_m_ shifts throughout series, while ΔT_m_ shifts for SLK were less pronounced (**Table S1**). This was also reflected in the overall low selectivity score (S) throughout the series, which is determined by the ratio between the kinases that are stabilized above 5 °C and the total number of kinases tested (**Figure 3**). Motivated by these findings, the entire series was further validated by orthogonal binding and activity assays to determine the *in vitro* and cellular potency and selectivity of the developed series in comparison to the selected reference compounds.

**Scheme 1.**
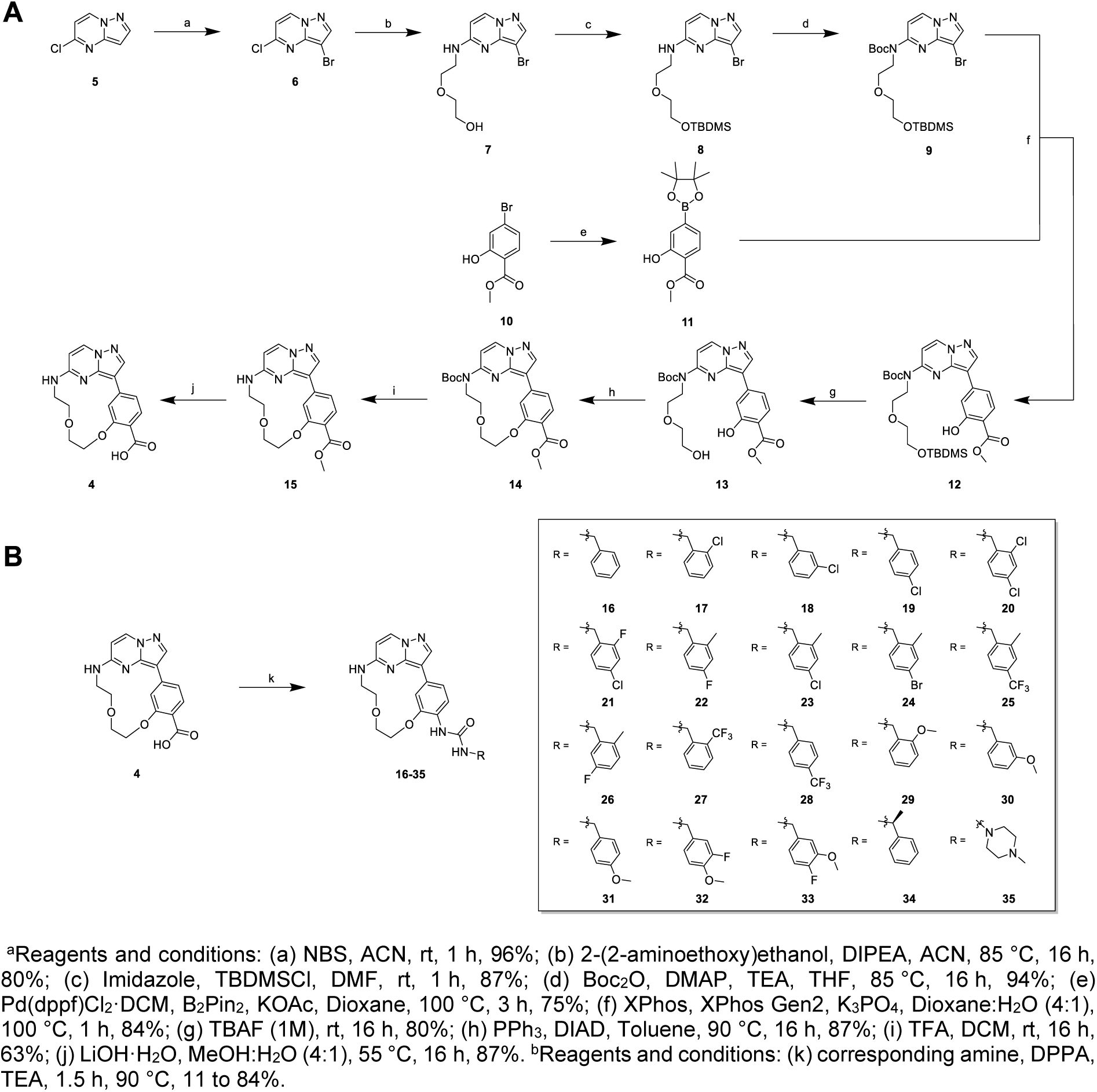
Synthesis of different urea derivatives. **A**. Synthesis of the macrocyclic precursor **4**.^a^ **B**. Synthesis of inhibitors **16**-**35**.^b^

**Figure 3.**
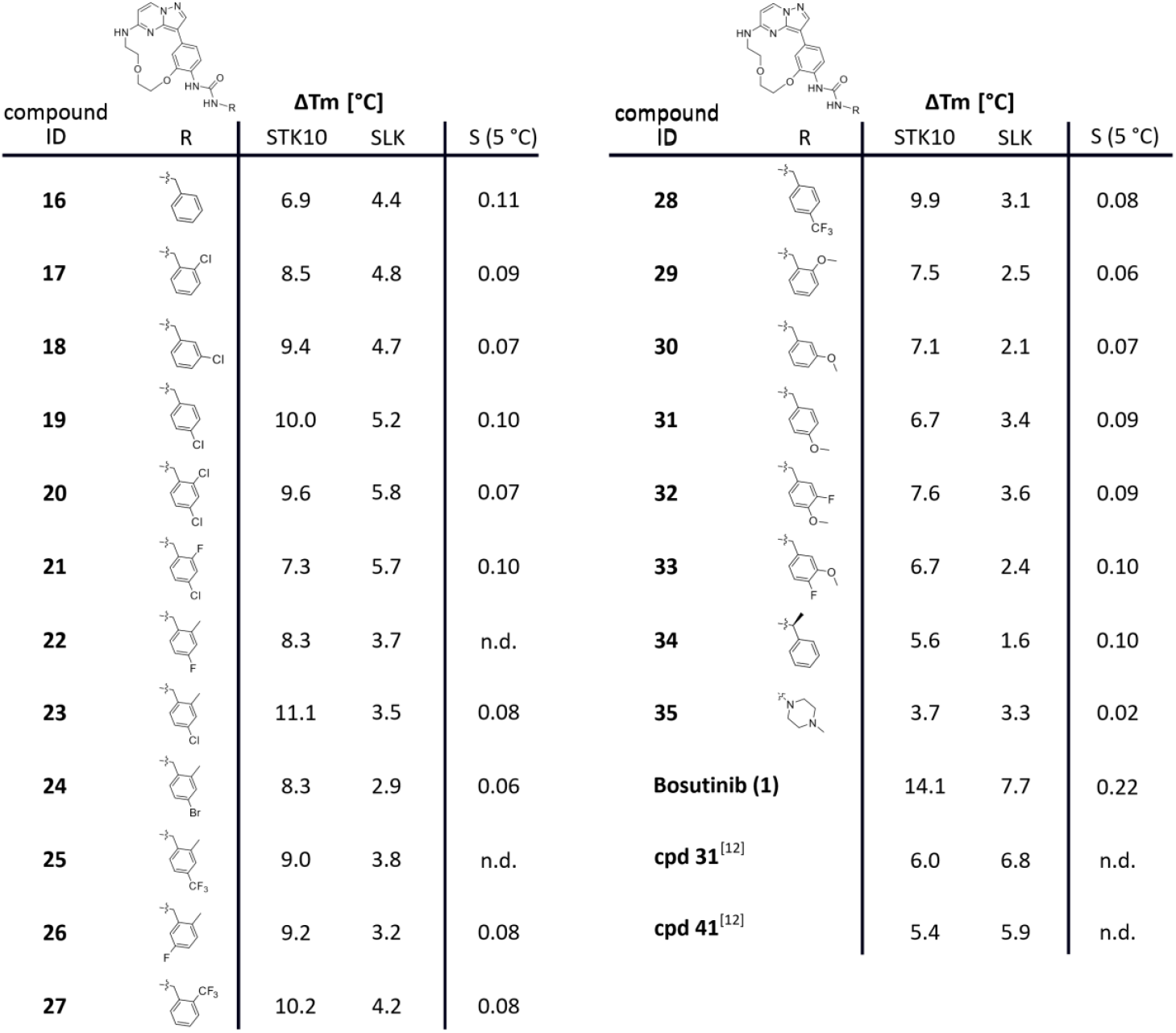
Summary of ΔT_m_ shifts of compounds **16**-**35**, Bosutinib (**1**), (**cpd 31**)[12] and (**cpd 41**)[12] for STK10 and SLK in °C. The selectivity score was calculated as the number of kinases showing a thermal shift >5 °C divided by the total number of kinases screened (see supplemental **Table S1**).

Figure 4 summarizes the *in vitro* and cellular evaluation for **16-35**, Bosutinib (**1**) and dual STK10/SLK inhibitors **cpd 31**[12] and **cpd 41**[12]. The unsubstituted benzylamine derivative **16** led to moderate stabilization of both STK10 and SLK (6.9/4.4 °C respectively) while derivatization of the methylene group of the urea moiety (**34**) and the introduction of a methyl piperazine moiety (**35**) led to lower ΔT_m_ shifts indicating that the benzylamine moiety is required for moderate stabilization of both kinases (**Figure 3**). This was further confirmed in cellular assays using the NanoBRET target engagement assay (EC_50_ = 5.7 µM **34**; EC_50_ = 15.2 µM **35** on STK10) and with a PhosphoSens *in vitro* activity assay[33] (IC_50_ = 10.2 µM **34**) (**Figure 4**). Different substitution patterns on the benzyl group were tolerated. Especially the introduction of an *ortho, meta*, and *para*-chlorine (**17**–**19**) moiety increased both stabilization (8.5–10.0 °C) of STK10 and the selectivity window towards SLK (4.7–5.2 °C). This correlated well with *in cellulo* NanoBRET data as compounds **17–19** all revealed EC_50_ values <1 µM for STK10 while no binding (>50 µM) was detected for SLK. Methoxy substituents in all three positions (*ortho, meta* and *para*) of the benzyl moiety seemed to decrease the binding affinity for STK10, since the corresponding compounds **29**–**31** showed lower ΔT_m_ shifts for STK10 (**Figure 3**). Based on this data set, we decided to explore different combinations of *ortho* and *para* substituents on the benzylamine moiety. For this purpose, we introduced a methyl group in *ortho*-position and tested different halogens at the *meta*-position (**22-25**).

Notably, the pattern of an *ortho*-methyl and a *para*-chlorine substituent (**23**) led to the highest stabilization in our DSF screen of 11.1 °C, which additionally was confirmed by an EC_50_ value of <1 µM, determined in NanoBRET (cellular) and a PhosphoSens assay (*in vitro*) (**Figures 4, 5B** and **5C**). We evaluated the selectivity profile of **23** in our in-house DSF kinase panel to better understand the influence of the back-pocket targeting moiety. Alongside STK10 other members of the sterile 20 (Ste20) superfamily such as STK3, STK4 were considerably stabilized by **23** with ΔTm values of 9.1/7.4 °C (STK3/STK4) respectively (**Figure 5B**). These off-targets in addition to the closely related kinase SLK were evaluated in cells using NanoBRET. In the NanoBRET assay, a substantial cellular effect of **23** on STK10 was observed (EC_50_ = 0.89 µM), while no effect on STK3, STK4 and SLK was detected at concentrations up to 50 µM (**Figure 5C**). Additionally, we confirmed the binding of **23** *in vitro* using surface plasmon resonance (SPR) revealing a *K*_*d*_ value of 365 ± 45 nM with fast binding kinetics (**Figure 5D**).

**Figure 4.**
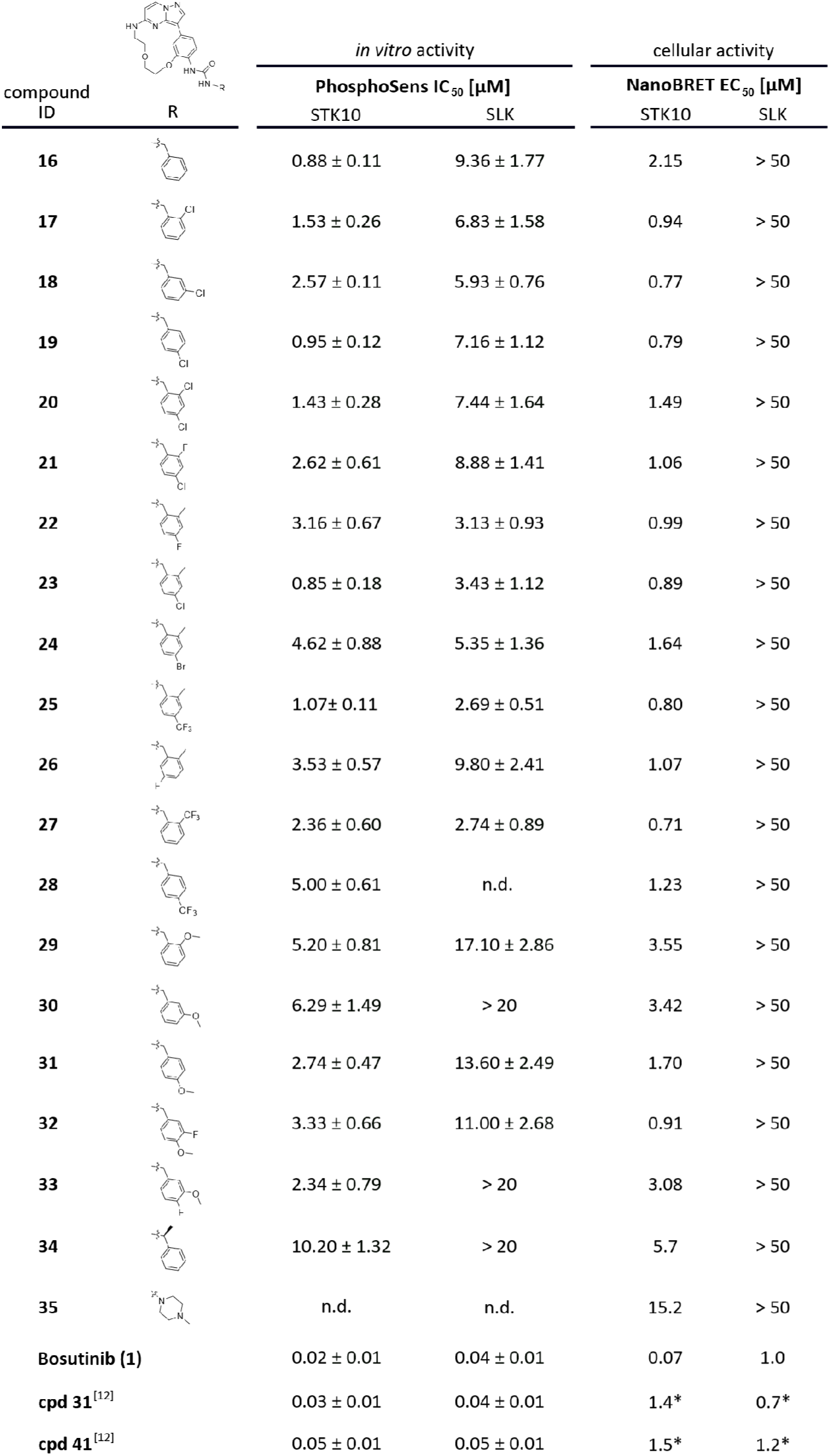
*In vitro* and cellular activity of compounds **16**-**35**, Bosutinib (**1**), **cpd 31**[12] and **cpd 41**[12] for STK10 and SLK were determined using a PhosphoSens assay (ATP concentration: 160 µM for STK10, 150 µM for SLK) and a NanoBRET assay. *reported values for **cpd 31**[12] and **cpd 41**[12] by Serafim *et al*.

**Figure 5.**
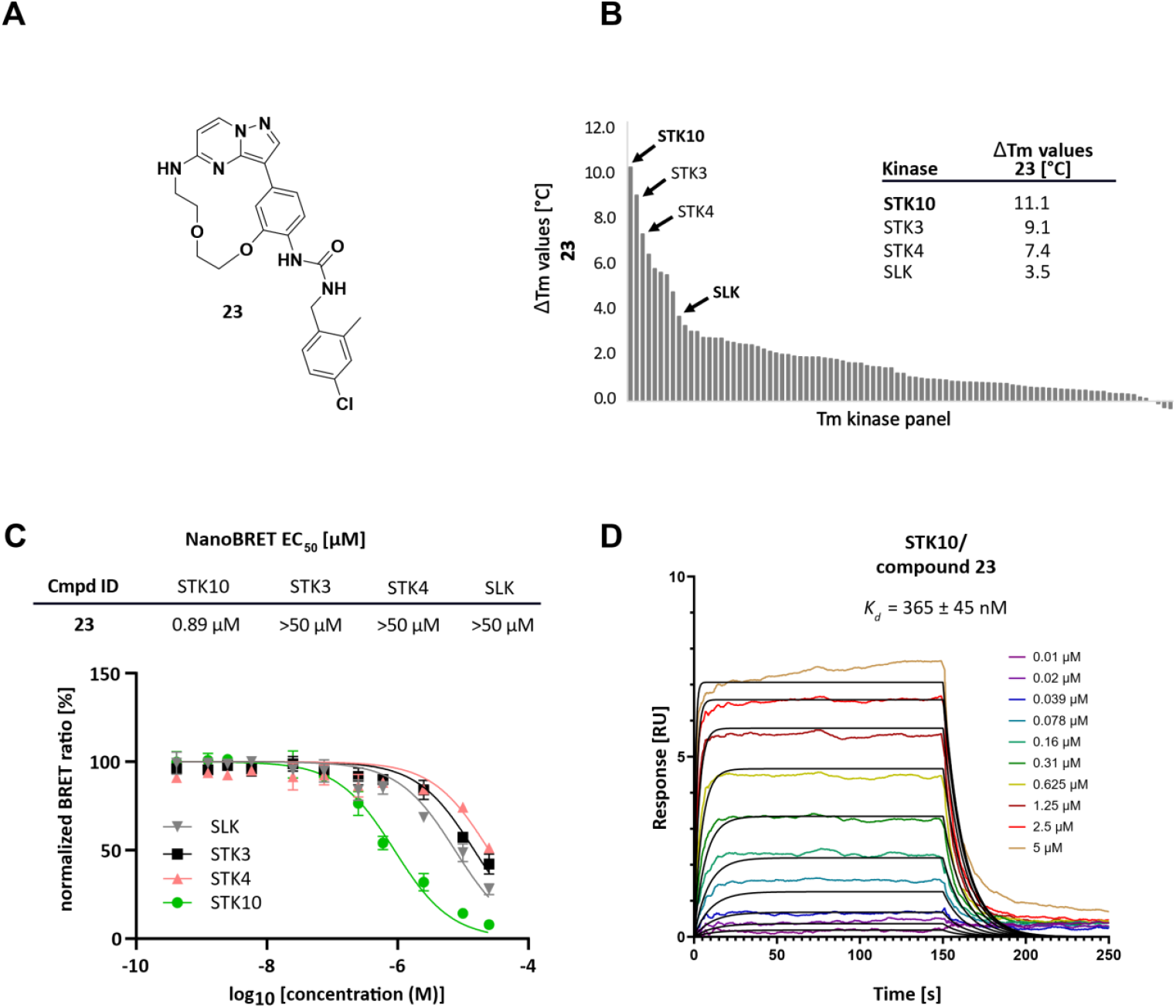
*In vitro* and cellular profiling of 23. A. Lewis structure of **23. B. 23** screened against an in-house DSF panel of 90 protein kinases. Values are shown in °C. **C**. On target in cell evaluation of **23**. EC_50_ values measured with NanoBRET against the most prominent off-targets STK3, STK4 and the closely related kinase SLK. **D**. SPR sensorgram overlay plot of **23** binding to STK10 at varying concentrations (colored) with kinetic fit (black).

To gain insides on the binding mode, we solved the crystal structure of **23** in complex with STK10. Analysis revealed a canonical hydrogen bond to the hinge region of the kinase between C113 and the N1 nitrogen of **23**. An extended water network mediated polar interactions to different amino acids in the ATP binding pocket. W1 interacted with the aniline-like nitrogen and W2 with the oxygen of the PEG linker of **23**, illustrating the contribution of the linker region of the macrocycle to the overall binding. The carbonyl functional group of the urea motif interacted with W3 and enabled polar contacts with D175 and F176 of the DFG loop of STK10 (**Figure 6A**). Non-polar contacts were likely the main driving force between the benzylic moiety of **23** and the back-pocket of the kinase. Particularly the chlorine substituent engaged by forming van der Waals contacts to hydrophobic amino acid side chains as well as a weak halogen(δ^+^)-π interaction to Y78. The bulky *N*-substituted benzylic 2-methyl-4-Chloro urea moiety protruded deep into the back-pocket of STK10 but without inducing the type II (DFG *out*) conformation expected for compounds bearing a urea motif.[28] Indeed, the compound induced an unexpected, kinked conformation of the αC helix that accommodated the bulky aromatic residue that we introduced. Additionally, the P-loop of the kinase underwent a vast conformational change that was stabilized both by hydrophobic interactions with the αEF helix of the activation segment and the benzylic residue of **23** (**Figure 6B**). To assess the influence of the conformation on the activation state of the kinase we compared the architecture of the catalytic (C) and regulatory spine (R) across different inhibitor types. The type I inhibitor Bosutinib binds to the active conformation of the kinase which is reflected in the formation of extensive hydrophobic contacts of the R-and C-spine (**Figure 6C**, left panel). In contrast a type II inhibitor binds to the inactive conformation and induces an outwards shift of F176 of the R spine (**Figure 6C**, middle panel). Interestingly, **23** stabilizes an inactive conformation of the kinase with assembled R and C-spines reminiscent of a type I binding mode, while the ATP/Mg^2+^ binding motif VAIK is repressed (**Figure 6C**, right panel).[34] At the same time the benzyl residue of **23** bound to the urea motif is perfectly positioned between the catalytic lysine (K65) of the β3-sheet and the glutamate (E81) of the αC-helix and thus stabilizes the inactive state of the kinase (**Figure S2**).

**Figure 6.**
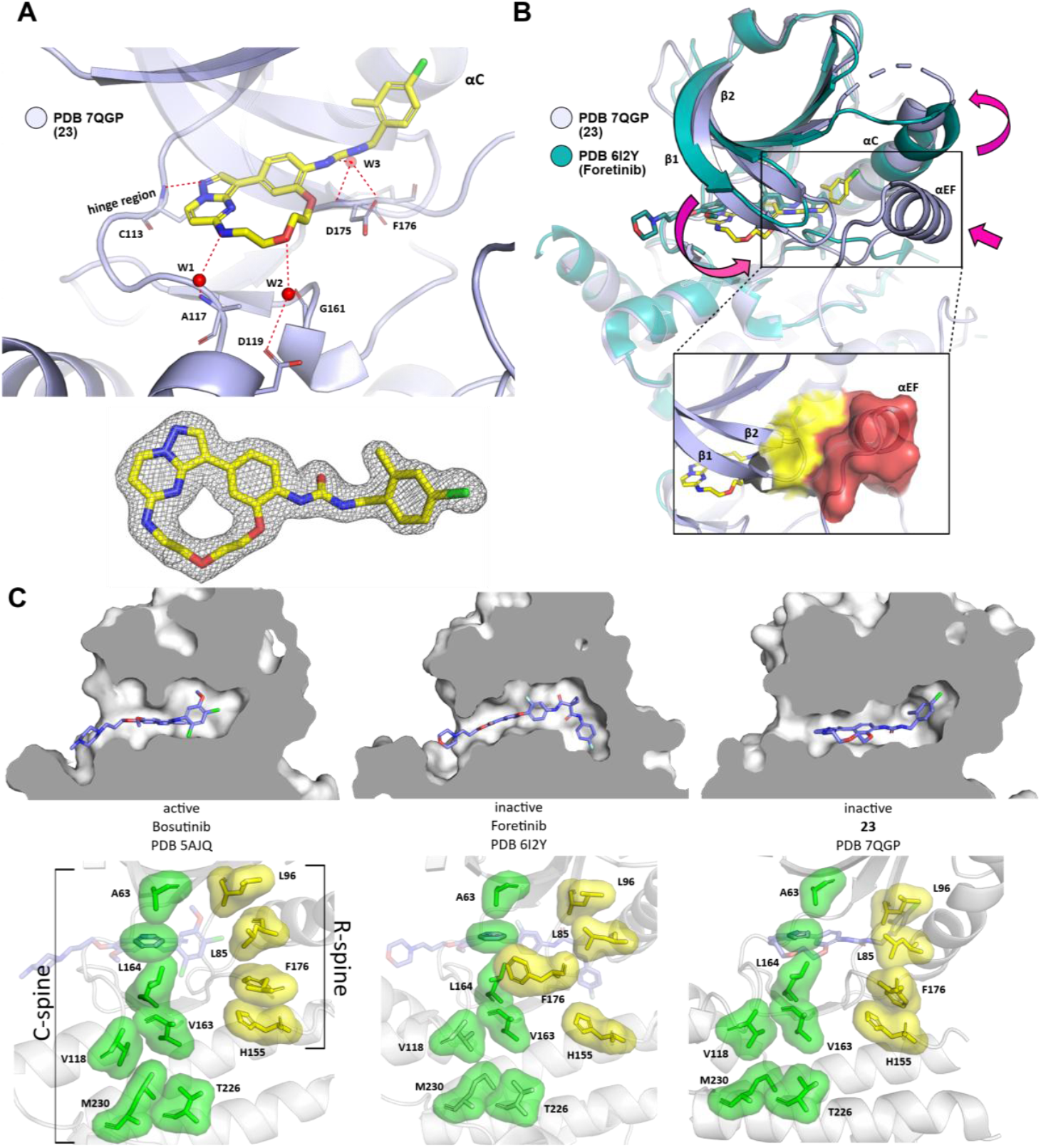
Crystal structure of 23 in complex with STK10: **A**. Top: Polar interactions of **23** (yellow sticks) with STK10 (livid, PDB 7QGP). The hydrogen bond of nitrogen N1 of **23** with C113 in the hinge region of the kinase shown as red dotted line. Water molecules W1 and W2 (red spheres) form an extended water network with amino acids A117/G161and the heteroatoms in the linker region of the macrocycle. W3 mediates the polar interaction of the urea motif with the DFG region of STK10. Bottom: electron density map of **23. B**. Overlay of PDB 62IY with Foretinib (teal) and **23** PDB 7QGP (livid). Important conformational changes are indicated by pink arrows: a substantial conformation change of the p-loop leads to an extended interaction with the well-resolved αEF helix in the activation segment (yellow and red surfaces respectively). The αC helix is found in a kinked conformation induced by the back-pocket moiety of **23. C**. Top, left to right: Cross sections of type I (Bosutinib, PDB 5AJQ), type II (Foretinib, PDB 6I2Y) and **23** (PDB 7QGP) in complex with STK10. Bottom, left to right: architecture of the C (catalytic)-spine (green) and R (regulatory)-spine (yellow). Type I inhibitors usually bind the active conformation of the kinase. Both the C-and R-spine are assembled and stabilized by extensive hydrophobic contacts between aliphatic and aromatic amino acids. The backbone of the R-spine is disrupted via an outwards movement of the aromatic residue in the DFG region (F176 in STK10) after binding of a type II inhibitor. **23** induces a bent conformation of the αC helix which allows the extension in the upper region of the back-pocket without influencing the overall assembly of both the R- and C-spines.

In this study we outline a strategy for the selective inhibition of STK10 that utilizes a late-stage modification approach of a pyrazolo[1,5-*a*]pyrimidine macrocyclic scaffold. The Curtius rearrangement was applied to the macrocyclic scaffold to introduce a variety of differently decorated urea functionalities targeting the back-pocket region of the STK10. Although many co-crystal structures of STK10 with different inhibitors have already been described in the PDB, a potent and selective tool compound for STK10 is still missing. The currently best profiled inhibitor, “**cpd 31**”, is a dual STK10/SLK inhibitor with some additional kinase off-targets. However, this maleimide-based compound showed > 1 µM activity in the cellular context. In this study, we have chosen a late-stage modification of an unselective macrocyclic scaffold. The synthesis of macrocyclic compounds is often tedious as it usually requires several protecting groups as well as macrocyclization. However, this is often the last step of the reaction, which makes systematic SAR lengthy and more challenging. By introducing an urea motif, which is a frequent structural element in kinase inhibitors, we were able to rapidly synthesize a macrocyclic series and comprehensively characterize its *in vitro* and *in cellulo* activity. By obtaining a co-crystal structure of **23** with STK10, we were able to identify a unique, previously undescribed binding conformation for STK10. Compound **23** showed excellent selectivity towards approximately 90 kinases as well as towards closely related kinases within the STE20 subfamily such as STK3/4 and especially towards the closest relative SLK. The back-pocket of STK10 is highly flexible and can be exploited for inhibitor design.[28] We propose that targeting this unique conformation of STK10 might open a new way for the development of selective STK10 inhibitors. This conformational selective design strategy can be used to further increase the affinity of the compound and lead to a potent tool compound to study the biology of STK10.

## ABBREVIATIONS

ACN: acetonitrile
DCM: dichloromethane
DIAD: diisopropyl azodicarboxylate
DIPEA: *N,N*-diisopropylethylamine
DMF: dimethylformamide
DPPA: diphenyl phosphorazidate
DRAK DAP: kinase-related apoptosis-inducing protein kinase
DSF: differential scanning fluorimetry
MST: mammalian sterile 20-like serine/threonine
NBS: *N*-Bromosuccinimide
rt: room temperature
SLK: STE20-like kinase
SPR: surface plasmon resonance
STK: serine/threonine protein kinase
TBAF: tetrabutylammonium fluoride
TBDMS: *tert*-butyldimethylsilyl chloride
TEA: triethylamine
TFA: trifluoroacetic acid
TK: tyorsine kinase
TKL: tyrosine kinase-like

## ACKNOWLEDGMENTS

The authors are grateful for support by the Structural Genomics Consortium (SGC), a registered charity (no. 1097737) that receives funds from Bayer AG, Boehringer Ingelheim, Bristol Myers Squibb, Genentech, Genome Canada through Ontario Genomics Institute [OGI-196], EU/EFPIA/OICR/McGill/KTH/Diamond Innovative Medicines Initiative 2 Joint Undertaking [EUbOPEN grant 875510], Janssen, Merck KGaA (aka EMD in Canada and U.S.), Pfizer, and Takeda. FAG, MPS, AK and SK would like to acknowledge funding from the German Cancer Consortium (DKTK) at the German Cancer Research Center (DKFZ). MPS is funded by the Deutsche Forschungsgemeinschaft (DFG, German Research Foundation), CRC1430 (Project-ID 424228829).

## AUTHOR CONTRIBUTIONS

Manuscript and figures were prepared by MD, LNT, FAG and edited by TH and SK. MD, LNT, JAA, CGK, MPS, TEM, LMW, AK, JG, CL, LE, BTB and MS contributed experimental data. Scientific supervision by FAG, TH, KS, SM, SK.

