## Supplemental Information for "Un-LOK-ing a new approach for conformational selective targeting of STK10 (LOK)"

#### Supporting Information

Martina Dettenhöfer[1,2‡], Laura Nadine Tandara[1,2‡], Jennifer Alisa Amrhein[1,2], Christian Georg Kurz[1,2], Martin Peter Schwalm[1,2,3], Theresa Elisabeth Mensing[1,2], Laurenz Maximilian Wahl[1,2], Andreas Krämer[1,2,3], Joshua Gerninghaus[1,2], Christopher Lenz[1,2], Lewis Elson[1,2], Benedict-Tilman Berger[1,2], Martin Schröder[1,2], Krishna Saxena[1,2], Susanne Müller[1,2], Stefan Knapp[1,2,3], Francesco Aleksy Greco[1,2,3*], Thomas Hanke[1,2*]

[1]Institute of Pharmaceutical Chemistry, Goethe University Frankfurt, Max-von-Laue-Str. 9, 60438 Frankfurt am Main, Germany

[2]Structural Genomics Consortium, Buchmann Institute for Molecular Life Sciences, Goethe-University Frankfurt, Max-von-Laue-Str. 15, 60438 Frankfurt am Main, Germany

[3]German Cancer Consortium (DKTK), German Cancer Research Center (DKFZ), DKTK Site Frankfurt-Mainz, 69120 Heidelberg, Germany

**Figure S1:** A “late-stage” modification approach of **4** (ODS2004070) by introduction of diverse back pocket motifs, resulting in a series of 20 urea-based derivatives.

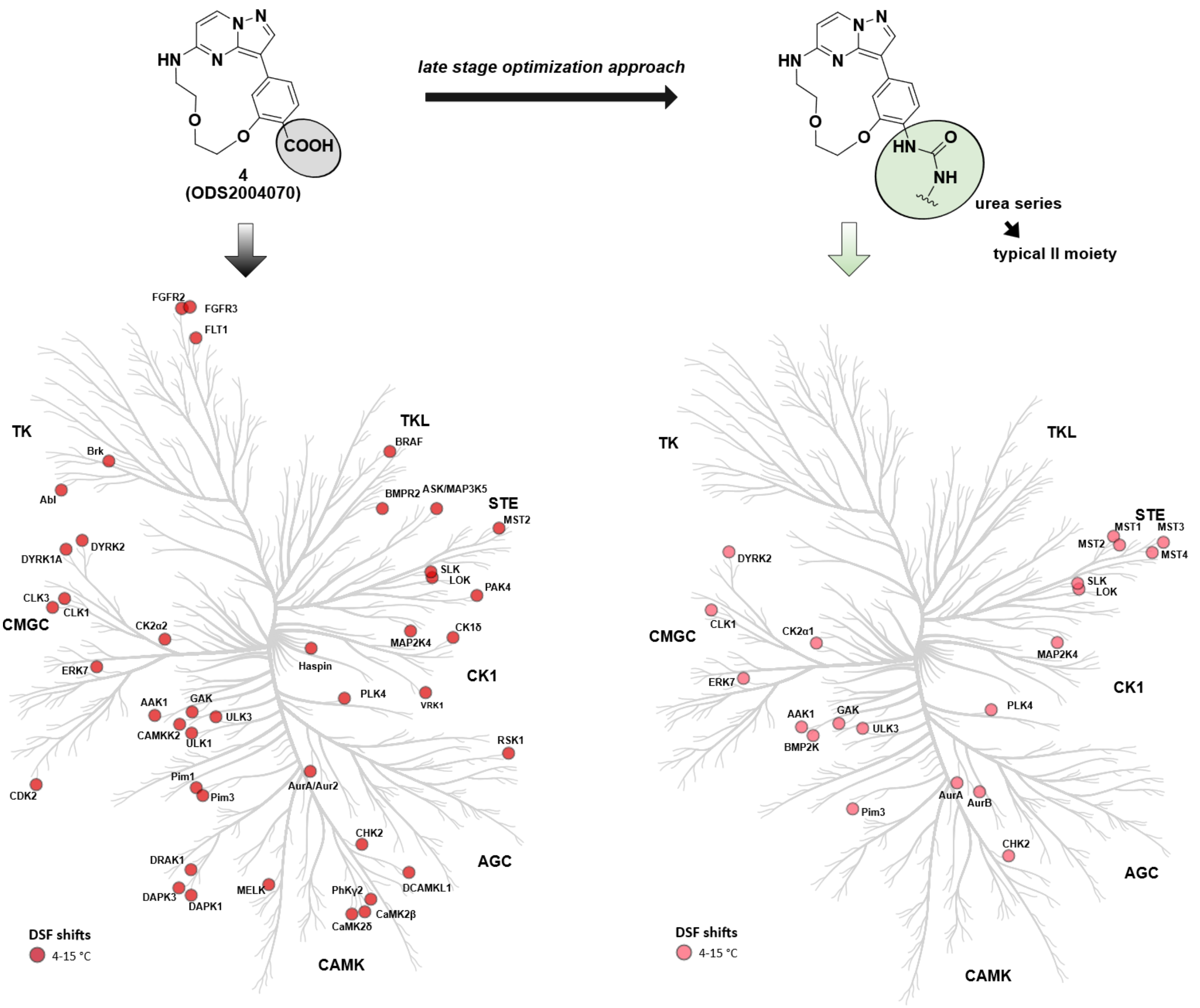

**Figure S2:** Binding mode of **23** to STK10, disrupting the catalytic β3-lysine (K65) and αC-glutamate (E81).

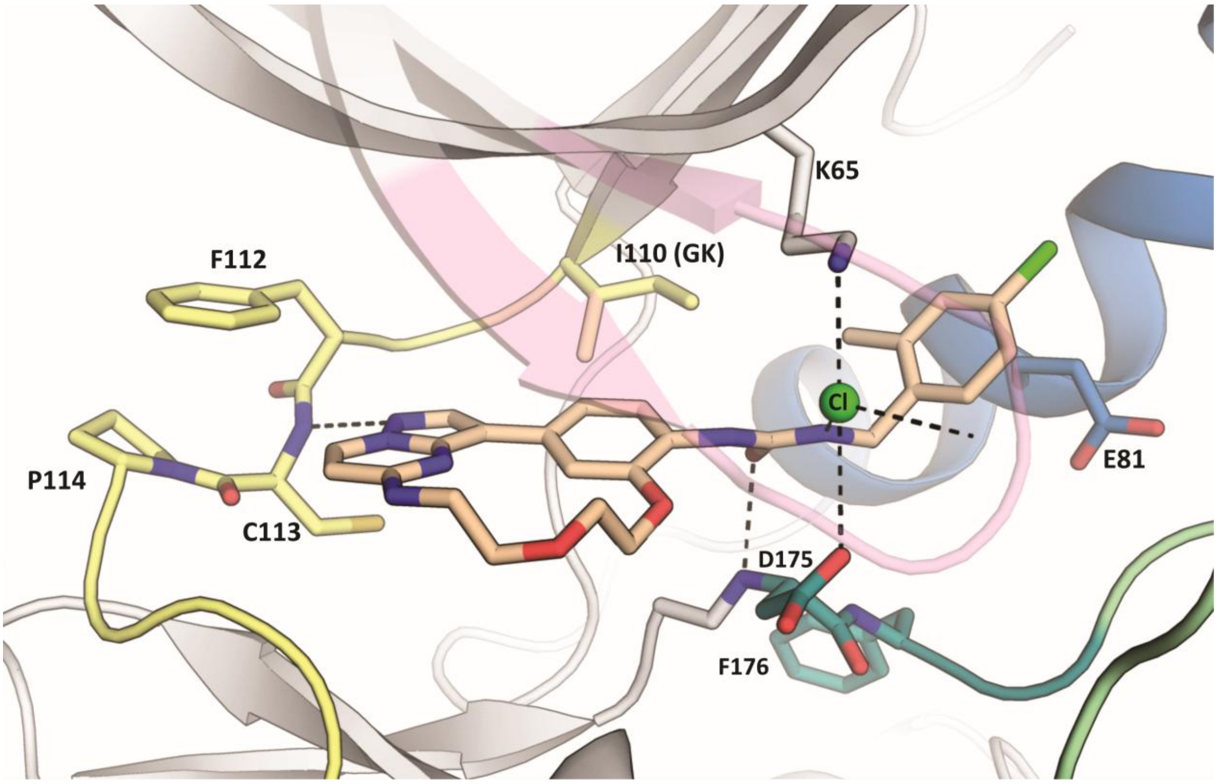

**Table S1:** Selectivity of **16-21, 23-24 and 26-35** measured in in-house Tm assay.

| 16 | | 17 | | 18 | | 19 | | 20 | | 21 | | 23 | | 24 | | 26 | |
|---|---|---|---|---|---|---|---|---|---|---|---|---|---|---|---|---|---|
| targets | ΔTm | targets | ΔTm | targets | ΔTm | targets | ΔTm | targets | ΔTm | targets | ΔTm | targets | ΔTm | targets | ΔTm | targets | ΔTm |
| BMP2KA | 12,2 | BMP2KA | 9,2 | **STK10** | 9,4 | **STK10** | 10,0 | **STK10** | 9,6 | BMP2KA | 7,5 | **STK10** | 11,1 | CHEK2A | 8,6 | **STK10** | 9,2 |
| AAK1 | 8,3 | **STK10** | 8,5 | BMP2KA | 8,7 | BMP2KA | 9,8 | STK3 | 8,0 | **STK10** | 7,3 | STK3 | 9,1 | **STK10** | 8,3 | BMP2KA | 9,1 |
| STK6A | 7,5 | AAK1 | 7,0 | STK3 | 8,0 | STK6A | 7,1 | STK6A | 7,4 | AAK1 | 7,1 | STK4A | 7,4 | AURKBA | 7,7 | AURKBA | 7,8 |
| **STK10** | 6,9 | STK6A | 6,7 | STK6A | 7,3 | AAK1 | 7,1 | BMP2KA | 6,7 | STK3 | 6,9 | AAK1 | 6,5 | BMP2KA | 6,7 | CHEK2A | 7,2 |
| UKL3A | 6,8 | STK3 | 6,2 | AAK1 | 5,5 | STK3 | 6,7 | AAK1 | 6,3 | STK6A | 6,9 | BMP2KA | 5,9 | GAKA | 5,9 | STK6A | 6,4 |
| CHEK2A | 6,5 | MAPK15A | 6,0 | STK17AA | 5,1 | MAP2K4A | 5,7 | SLK | 5,8 | CDK2A | 5,9 | CHEK2A | 5,7 | STK6A | 5,4 | GAKA | 6,0 |
| GAKA | 6,1 | CHEK2A | 5,3 | SLK | 4,7 | CDK2A | 5,6 | MAP2K4A | 3,9 | SLK | 5,7 | STK6A | 5,6 | AAK1A | 4,0 | STK17AA | 5,2 |
| STK3 | 5,9 | MAP2K4A | 5,0 | UKL3A | 4,6 | STK4A | 5,5 | STK4A | 3,6 | MAP2K4A | 5,4 | MAP2K4A | 4,8 | STK3A | 3,8 | AAK1A | 5,0 |
| MAP2K4A | 5,2 | SLK | 4,8 | STK4A | 4,5 | SLK | 5,2 | CHEK2A | 3,6 | CHEK2A | 5,0 | ULK3A | 3,7 | ULK3A | 3,7 | STK3A | 4,1 |
| FGFR3A | 5,0 | GAKA | 4,8 | CDK2A | 4,3 | ULK3A | 4,6 | PIM3A | 3,3 | STK4A | 4,9 | SLK | 3,5 | STK17BA | 3,6 | PIM3A | 4,0 |
| STK17AA | 4,9 | CLK1A | 4,5 | GAKA | 4,1 | CHEK2A | 4,5 | PIM1 | 3,0 | UKL3A | 4,6 | STK17BA | 3,3 | MAP2K4A | 3,2 | ULK3A | 3,9 |
| STK4A | 4,9 | STK4A | 4,1 | MAP2K4A | 4,1 | GAKA | 4,4 | K2A2 | 3,0 | K2A2 | 4,2 | CDK2A | 3,1 | SLK | 2,9 | FGFR2A | 3,5 |
| MAPK15A | 4,9 | STK17BA | 4,1 | STK39A | 4,0 | FGFR2A | 3,7 | PLK4A | 2,8 | GAKA | 3,9 | MERTK | 3,1 | STK4A | 2,8 | BMPR2A | 3,4 |
| FGFR2A | 4,8 | K2A2 | 4,0 | FLT1A | 3,7 | K2A2 | 3,6 | STK17BA | 2,8 | PIM3A | 3,9 | GAKA | 2,8 | PIM3A | 2,8 | PLK4A | 3,3 |
| FLT1A | 4,7 | PLK4A | 3,9 | FGFR3A | 3,6 | FLT1A | 3,4 | CAMK1GA | 2,7 | CAMKK2B | 3,5 | PLK4A | 2,8 | TTKA | 2,6 | SLK | 3,2 |
| BMPR2A | 4,4 | CAMKK2B | 3,3 | CHEK2A | 3,6 | CAMKK2B | 3,4 | CDK2A | 2,5 | PIM1 | 2,8 | K2A2 | 2,8 | BMPR2A | 2,3 | MAP2K4A | 3,2 |
| SLK | 4,4 | UKL3A | 3,2 | FGFR2A | 3,3 | STK39A | 2,6 | PAK1A | 2,5 | CLK1A | 2,8 | PCTK1A | 2,8 | STK17AA | 2,3 | STK17BA | 3,1 |
| PLK4A | 4,1 | CDK2A | 2,9 | PIM1 | 3,2 | MERTK | 2,5 | GAKA | 2,4 | MAP3K5A | 2,7 | CLK1A | 2,6 | PLK4A | 2,3 | STK4A | 3,1 |
| CDK2A | 4,1 | CASKA | 2,8 | PLK4A | 3,2 | PLK4A | 2,5 | UKL3A | 2,3 | CASKA | 2,7 | DYRK1AA | 2,6 | CAMK2DA | 2,2 | MAPK14A | 2,7 |
| DYRK1AA | 3,8 | BMPR2A | 2,6 | CAMKK2B | 3,1 | PIM3A | 2,4 | MERTK | 2,2 | MERTK | 2,7 | STK39A | 2,5 | FGFR2A | 2,2 | MAPK15HSD | 2,4 |
| CAMKK2B | 3,8 | PCTK1A | 2,5 | K2A2 | 3,1 | BMPR2A | 2,3 | MAP3K5A | 2,2 | STK17BA | 2,7 | FGFR3A | 2,5 | EPHA7A | 2,0 | CAMK2DA | 2,4 |
| DYRK2A | 3,7 | PIM3A | 2,4 | PIM3A | 3,0 | CASKA | 2,2 | ABL1 | 2,2 | PLK4A | 2,6 | CASKA | 2,5 | MAPK14A | 2,0 | CDK2A | 2,3 |
| CLK1A | 3,4 | MELKA | 2,4 | CLK1A | 2,9 | CLK1A | 2,1 | CLK1A | 2,1 | SRCA | 2,6 | MARK3A | 2,4 | MARK4A | 1,9 | DYRK2A | 2,3 |
| DAPK3 | 3,4 | STK39A | 2,4 | BMPR2A | 2,9 | SRCA | 2,1 | MARK4A | 2,0 | FLT1A | 2,6 | PIM1 | 2,3 | PAK4A | 1,9 | SRCA | 2,2 |
| STK39A | 3,0 | STK17AA | 2,3 | DYRK2A | 2,8 | STK17AA | 2,0 | RIOK1A | 1,9 | MARK4A | 2,6 | PIM3A | 2,2 | BMXA | 1,8 | CLK1A | 2,1 |
| FESA | 3,0 | MARK4A | 2,2 | SRCA | 2,6 | MARK4A | 2,0 | STK39A | 1,9 | STK39A | 2,4 | MST4A | 2,1 | SRCA | 1,7 | EPHA7A | 2,0 |
| SRCA | 2,5 | FLT1A | 2,1 | STK17BA | 2,4 | ABL1 | 1,8 | OSR1A | 1,8 | MELKA | 2,4 | CAMK1DA | 2,1 | BRPF1B | 1,6 | TTKA | 1,9 |
| FGFR1B | 2,4 | FGFR3A | 2,1 | MAPK15A | 2,3 | MAPK15A | 1,8 | BRD4 | 1,8 | FGFR2A | 2,3 | PKMYT1A | 2,1 | NEK7A | 1,6 | FGFR1B | 1,9 |
| PIM3A | 2,4 | SRCA | 1,8 | TTKA | 2,2 | PIM1 | 1,8 | CASKA | 1,7 | ABL1 | 2,3 | PAK1A | 2,0 | DCAMKL1A | 1,6 | MARK4A | 1,9 |
| K2A2 | 2,3 | TTKA | 1,8 | CASKA | 2,1 | FGFR3A | 1,7 | MARK3A | 1,7 | BRD4 | 2,0 | CAMK1GA | 2,0 | CDK2A | 1,6 | PAK4A | 1,8 |
| CASKA | 2,2 | ABL1 | 1,7 | FGFR1B | 2,0 | FGFR1B | 1,5 | MAPK10A | 1,6 | STK17AA | 2,0 | MAPK9A | 2,0 | OSR1A | 1,6 | AKT3A | 1,8 |
| MELKA | 2,0 | FGFR2A | 1,7 | MERTK | 1,8 | GPRK5A | 1,5 | STK17AA | 1,6 | CAMK1GA | 2,0 | BRD4 | 2,0 | EPHA5A | 1,4 | DCAMKL1A | 1,6 |
| MAP3K5A | 2,0 | MERTK | 1,6 | ABL1 | 1,8 | MELKA | 1,5 | CAMK2BA | 1,6 | EPHA7A | 1,9 | FLT1A | 2,0 | MAP3K5A | 1,4 | CASKA | 1,2 |
| PCTK1A | 1,8 | PIM1 | 1,5 | OSR1A | 1,7 | TTKA | 1,4 | BMX | 1,6 | PKMYT1A | 1,9 | FGFR2A | 1,9 | BRD4A | 1,4 | DAPK3A | 1,2 |
| BMX | 1,7 | BRD4 | 1,3 | MAP3K5A | 1,6 | PKMYT1A | 1,4 | PKMYT1A | 1,6 | BMPR2A | 1,8 | CAMKK2B | 1,9 | CAMK1DA | 1,3 | OSR1A | 1,2 |
| MAPK9A | 1,7 | GPRK5A | 1,3 | BRD4 | 1,5 | BMX | 1,4 | MAPK9A | 1,5 | TTKA | 1,8 | TTKA | 1,8 | MST4A | 1,2 | EPHA5A | 1,1 |
| MARK4A | 1,7 | RPS6KA1A | 1,2 | PKMYT1A | 1,5 | MAP3K5A | 1,4 | FLT1A | 1,4 | MAPK15A | 1,8 | BMX | 1,8 | FGFR1B | 1,2 | MELKA | 1,1 |
| TTKA | 1,5 | BMX | 1,2 | MELKA | 1,5 | PCTK1A | 1,3 | FGFR3A | 1,4 | OSR1A | 1,6 | ABL1 | 1,7 | PAK1A | 1,1 | CAMK1DA | 1,0 |
| MST3A | 1,5 | DYRK2A | 1,1 | BMX | 1,4 | PHKG2A | 1,2 | GPRK5A | 1,3 | FGFR3A | 1,6 | STK38LA | 1,7 | TIF1A | 1,0 | PCTK1A | 1,0 |
| RPS6KA1A | 1,4 | GSK3BB | 1,0 | MARK4A | 1,4 | STK17BA | 1,2 | MAPK15A | 1,3 | MARK3A | 1,5 | CAMK2DA | 1,7 | RPS6KA5A | 1,0 | BRPF1B | 1,0 |
| MERTK | 1,4 | PKMYT1A | 1,0 | RIOK1A | 1,4 | RIOK1A | 1,1 | TTKA | 1,2 | GPRK5A | 1,4 | WNK1A | 1,6 | VRK1A | 1,0 | PIM1A | 1,0 |
| MAP2K6A | 1,4 | MAP3K5A | 1,0 | PCTK1A | 1,3 | MAP2K6A | 1,1 | SRCA | 1,2 | MAPK10A | 1,4 | MARK4A | 1,5 | TLK1A | 1,0 | MAP3K5A | 0,9 |
| CAMK2BA | 1,3 | MAP2K6A | 0,9 | CAMK4A | 1,3 | MST4A | 1,1 | TOPKA | 1,2 | BMX | 1,4 | SRCA | 1,5 | CLK1A | 1,0 | NEK7A | 0,9 |
| STK17BA | 1,3 | CAMK1GA | 0,9 | DAPK3 | 1,3 | CAMK4A | 1,0 | CAMKK2B | 1,2 | MAP2K6A | 1,3 | TOPKA | 1,5 | STK39A | 1,0 | CAMKK2B | 0,9 |
| MST4A | 1,3 | BRAFA | 0,8 | MAP2K6A | 1,2 | BRD4 | 1,0 | PDK4A | 1,1 | STK38LA | 1,2 | BMPR2A | 1,5 | CAMK4A | 0,8 | TLK1A | 0,9 |
| PHKG2A | 1,3 | MARK3A | 0,8 | DYRK1AA | 1,1 | PAK1A | 0,9 | BMPR2A | 1,1 | MAP2K1A | 1,2 | CSNK1DA | 1,2 | MARK3A | 0,8 | CSNK2A2A | 0,9 |
| DAPK1 | 1,3 | PDK4A | 0,8 | STK38LA | 1,1 | DAPK3 | 0,9 | PCTK1A | 1,1 | PDK4A | 1,1 | PAK4A | 1,2 | PCTK1A | 0,7 | STK39A | 0,9 |
| ABL1 | 1,2 | MST4A | 0,7 | GPRK5A | 1,1 | CAMK1GA | 0,9 | MAP2K6A | 1,1 | MST4A | 1,1 | FGFR1B | 1,1 | CSNK1EA | 0,7 | EPHB3A | 0,9 |
| PIM1 | 1,2 | DCAMKL1A | 0,7 | PDK4A | 1,0 | BRAFA | 0,8 | CAMK1DA | 1,0 | FGFR1B | 1,1 | MELKA | 1,1 | MAPK1A | 0,7 | MST4A | 0,8 |
| GPRK5A | 1,2 | OSR1A | 0,7 | CAMK1GA | 0,8 | MAPK10A | 0,8 | EPHA7A | 1,0 | RPS6KA1A | 1,1 | MAPK10A | 1,0 | CDC42BPAA | 0,7 | MAPK1A | 0,8 |
| GSK3BB | 1,1 | DAPK3 | 0,7 | EPHA7A | 0,8 | PDK4A | 0,8 | CDKL1A | 0,9 | DAPK3 | 1,1 | MAP3K5A | 1,0 | MAPK15HSD | 0,7 | DYRK1AA | 0,8 |
| STK38LA | 0,8 | STK38LA | 0,7 | MST4A | 0,8 | MST3A | 0,7 | MELKA | 0,9 | PCTK1A | 1,1 | EPHA5A | 1,0 | CAMKK2B | 0,7 | CAMK4A | 0,8 |
| BRAFA | 0,8 | MST3A | 0,7 | CAMK2BA | 0,8 | OSR1A | 0,7 | CAMK2DA | 0,9 | PAK1A | 1,0 | EPHA7A | 1,0 | EPHB3A | 0,7 | CSNK1EA | 0,7 |
| EPHB3A | 0,8 | PAK1A | 0,6 | PAK1A | 0,7 | STK38LA | 0,6 | FGFR2A | 0,9 | PAK4A | 0,9 | CAMK2BA | 0,9 | MAP2K6A | 0,6 | MAP2K6A | 0,7 |
| DCAMKL1A | 0,8 | CAMK4A | 0,6 | MARK3A | 0,7 | MAPK13A | 0,6 | FGFR1B | 0,9 | PHKG2A | 0,9 | NEK1A | 0,9 | MAP2K1A | 0,6 | PAK1A | 0,7 |
| EPHA2 | 0,8 | CSNK1DA | 0,6 | MAPK13A | 0,7 | GSK3BB | 0,6 | RPS6KA1A | 0,9 | CAMK4A | 0,9 | AKT3A | 0,9 | CASKA | 0,6 | RPS6KA5A | 0,7 |
| CSNK1DA | 0,7 | PHKG2A | 0,6 | MAPK1A | 0,7 | CAMK1DA | 0,6 | PAK4A | 0,8 | VRK1A | 0,9 | EPHB3A | 0,9 | MAPK8B | 0,6 | GSG2A | 0,6 |
| MARK3A | 0,7 | FGFR1B | 0,6 | RPS6KA1A | 0,7 | RPS6KA1A | 0,4 | MAPK14A | 0,8 | CAMK2DA | 0,8 | GPRK5A | 0,8 | PIM1A | 0,5 | ABL1A | 0,6 |
| BRD4 | 0,7 | MAPK13A | 0,5 | MST3A | 0,7 | EPHA7A | 0,4 | PHKG2A | 0,8 | CAMK2BA | 0,8 | MAPK15A | 0,8 | CSNK2A2A | 0,5 | BRD4A | 0,5 |
| PKMYT1A | 0,6 | WNK1A | 0,5 | EPHA2 | 0,6 | ULK1A | 0,4 | MAPK1A | 0,8 | CAMK1DA | 0,8 | OSR1A | 0,8 | PKMYT1A | 0,5 | GSK3BB | 0,5 |
| ULK1A | 0,6 | EPHA2 | 0,5 | MAPK10A | 0,6 | FESA | 0,4 | MAP2K1A | 0,8 | DYRK1AA | 0,7 | DYRK2A | 0,8 | MAPK10A | 0,5 | MAPK8B | 0,5 |
| CAMK4A | 0,5 | TAF1A | 0,5 | BRAFA | 0,6 | CLK3A | 0,4 | STK38LA | 0,7 | EPHA5A | 0,7 | MAPK1A | 0,8 | CDKL1A | 0,5 | VRK1A | 0,5 |
| OSR1A | 0,5 | RIOK1A | 0,5 | TOPKA | 0,5 | CSNK1DA | 0,4 | NEK1A | 0,7 | DYRK2A | 0,7 | VRK1A | 0,8 | DYRK2A | 0,5 | MAPK10A | 0,5 |
| GSG2A | 0,5 | TOPKA | 0,4 | MAPK9A | 0,5 | EPHA5A | 0,3 | DMPK1A | 0,7 | NEK1A | 0,7 | RPS6KA1A | 0,8 | FESA | 0,5 | CDKL1A | 0,5 |
| TOPKA | 0,4 | CDKL1A | 0,4 | CAMK2DA | 0,5 | CDKL1A | 0,3 | EPHA5A | 0,7 | TOPKA | 0,6 | STK17AA | 0,7 | DYRK1AA | 0,4 | HIPK2HSF | 0,4 |
| TAF1A | 0,4 | CAMK2DA | 0,4 | TAF1A | 0,5 | DMPK1A | 0,3 | CDC42BPAA | 0,7 | CLK3A | 0,6 | DCAMKL1A | 0,7 | DAPK3A | 0,4 | TIF1A | 0,4 |
| CLK3A | 0,3 | EPHA7A | 0,4 | PHKG2A | 0,4 | MAPK9A | 0,3 | TAF1A | 0,6 | MST3A | 0,6 | CAMK4A | 0,6 | MAP2K7A | 0,3 | MSSK1A | 0,4 |
| DMPK1A | 0,3 | MAPK9A | 0,4 | EPHA5A | 0,4 | DYRK2A | 0,2 | SRPK1A | 0,6 | MAPK13A | 0,6 | RIOK1A | 0,6 | DMPK1A | 0,3 | FESA | 0,3 |
| CAMK2DA | 0,2 | EPHB3A | 0,3 | CSNK1DA | 0,4 | MARK3A | 0,2 | GSK3BB | 0,6 | FESA | 0,6 | MAP2K1A | 0,6 | AKT3A | 0,3 | RPS6KA1A | 0,3 |
| PAK1A | 0,1 | ULK1A | 0,3 | CAMK1DA | 0,4 | TOPKA | 0,2 | ULK1A | 0,6 | MAPK14A | 0,5 | PDK4A | 0,6 | EPHB1A | 0,2 | BMXA | 0,3 |
| EPHA5A | 0,1 | MAPK1A | 0,3 | CLK3A | 0,3 | SRPK1A | 0,2 | DYRK1AA | 0,6 | BRAFA | 0,5 | TAF1A | 0,6 | EPHA4A | 0,2 | DMPK1A | 0,3 |
| NEK1A | 0,1 | CAMK2BA | 0,3 | VRK1A | 0,3 | EPHA2 | 0,2 | DYRK2A | 0,6 | EPHA2 | 0,5 | CDC42BPAA | 0,6 | CLK3A | 0,2 | MAP2K1A | 0,3 |
| RIOK1A | 0,1 | NEK1A | 0,2 | DAPK1 | 0,3 | VRK1A | 0,1 | DCAMKL1A | 0,6 | CDC42BPAA | 0,4 | PHKG2A | 0,5 | CSNK1DA | 0,2 | EPHA4A | 0,3 |
| MAPK13A | 0,1 | MAPK10A | 0,2 | MAPK14A | 0,3 | PAK4A | 0,1 | DAPK3 | 0,5 | TAF1A | 0,4 | BRAFA | 0,5 | MSSK1A | 0,2 | EPHA2A | 0,3 |
| SRPK1A | 0,0 | FESA | 0,2 | ULK1A | 0,3 | MAPK1A | 0,1 | MST4A | 0,5 | DCAMKL1A | 0,4 | EPHA2 | 0,5 | EPHA2A | 0,2 | PKMYT1A | 0,3 |
| AKT3A | 0,0 | CDC42BPAA | 0,1 | NEK1A | 0,2 | MAPK14A | 0,1 | FESA | 0,5 | SRPK1A | 0,4 | DMPK1A | 0,5 | HIPK2HSF | 0,2 | EPHB1A | 0,3 |
| CAMK1DA | 0,0 | CLK3A | 0,1 | EPHB3A | 0,2 | MAP2K1A | 0,1 | RPS6KA6A | 0,5 | DAPK1 | 0,3 | FESA | 0,5 | RPS6KA1A | 0,1 | CDC42BPAA | 0,2 |
| WNK1A | 0,0 | DAPK1 | 0,1 | MAP2K1A | 0,2 | DCAMKL1A | 0,0 | CSNK1DA | 0,4 | MAPK1A | 0,3 | CDKL1A | 0,4 | MAPKAPK2A | 0,1 | MARK3A | 0,2 |
| MAPK1A | -0,1 | VRK1A | 0,1 | PAK4A | 0,1 | CDC42BPAA | 0,0 | MAPK13A | 0,4 | MAPK9A | 0,3 | MAP2K6A | 0,4 | TAF1A | 0,1 | DAPK1A | 0,2 |
| EPHA7A | -0,1 | EPHA5A | 0,1 | DMPK1A | 0,1 | DAPK1 | -0,1 | CAMK4A | 0,4 | DMPK1A | 0,2 | GSK3BB | 0,4 | MELKA | 0,0 | MAP2K7A | 0,1 |
| CDKL1A | -0,1 | SRPK1A | 0,0 | WNK1A | 0,09 | CAMK2BA | -0,1 | AKT3A | 0,4 | GSK3BB | 0,2 | CLK3A | 0,4 | GSG2A | 0,0 | GPRK5A | 0,1 |
| MAP2K1A | -0,1 | PAK4A | 0,0 | FESA | 0,1 | CAMK2DA | -0,1 | WNK1A | 0,4 | ULK1A | 0,2 | MAPK13A | 0,3 | GPRK5A | 0,0 | TAF1A | 0,1 |
| PAK4A | -0,2 | MAPK14A | 0,0 | GSK3BB | 0,0 | EPHB3A | -0,1 | BRAFA | 0,4 | GSG2A | 0,1 | ULK1A | 0,3 | ULK1A | 0,0 | CLK3A | 0,1 |
| CDC42BPAA | -0,2 | DMPK1A | -0,1 | GSG2A | 0,0 | GSG2A | -0,2 | VRK1A | 0,4 | CDKL1A | 0,0 | MST3A | 0,3 | SRPK1A | -0,2 | MAPKAPK2A | 0,1 |
| CAMK1GA | -0,3 | MAP2K1A | -0,2 | CDC42BPAA | 0,0 | NEK1A | -0,2 | EPHA2 | 0,4 | CSNK1DA | 0,0 | MAPK14A | 0,3 | CAMK2BA | -0,2 | ULK1A | 0,1 |
| MAPK10A | -0,3 | AKT3A | -0,3 | DCAMKL1A | -0,1 | WNK1A | -0,2 | CLK3A | 0,3 | RIOK1A | -0,1 | SRPK1A | 0,2 | GSK3BB | -0,2 | CAMK2BA | 0,0 |
| VRK1A | -0,4 | CAMK1DA | -0,3 | CDKL1A | -0,1 | RPS6KA6A | -0,2 | DAPK1 | 0,2 | WNK1A | -0,2 | DAPK1 | 0,1 | DAPK1A | -0,3 | SRPK1A | 0,0 |
| MAPK14A | -0,6 | GSG2A | -0,3 | RPS6KA6A | -0,3 | AKT3A | -0,4 | EPHB3A | 0,1 | AKT3A | -0,3 | GSG2A | -0,1 | BRAFA | -0,3 | CSNK1DA | -0,1 |
| PDK4A | -0,7 | RPS6KA6A | -0,3 | SRPK1A | -0,3 | TAF1A | -0,4 | MST3A | 0,0 | EPHB3A | -0,4 | DAPK3 | -0,3 | MAPK13A | -0,4 | MAPK13A | -0,2 |
| RPS6KA6A | -1,2 | DYRK1AA | -1,3 | AKT3A | -0,4 | DYRK1AA | -1,0 | GSG2A | 0,0 | RPS6KA6A | -0,7 | RPS6KA6A | -0,3 | ABL1A | -0,4 | NEK1A | -0,4 |
| | | | | | | | | | | | | | | NEK1A | -0,4 | BRAFA | -0,6 |
| | | | | | | | | | | | | | | MAPK9A | -1,2 | NEK2A | -1,2 |
| | | | | | | | | | | | | | | NEK2A | -1,3 | NQO2A | -1,5 |
| | | | | | | | | | | | | | | NQO2A | -1,9 | MAPK9A | -1,6 |
| | | | | | | | | | | | | | | MST3A | -4,3 | MST3A | -2,0 |

| 27 | | 28 | | 29 | | 30 | | 31 | | 32 | | 33 | | 34 | | 35 | |
|---|---|---|---|---|---|---|---|---|---|---|---|---|---|---|---|---|---|
| targets | ΔTm | targets | ΔTm | targets | ΔTm | targets | ΔTm | targets | ΔTm | targets | ΔTm | targets | ΔTm | targets | ΔTm | targets | ΔTm |
| **STK10** | 10,2 | **STK10** | 9,9 | BMP2KA | 8,3 | BMP2KA | 9,8 | BMP2KA | 12,0 | BMP2KA | 10,9 | BMP2KA | 10,2 | BMP2KA | 9,1 | BMP2KA | 11,8 |
| CHEK2A | 9,9 | AURKBA | 9,0 | **STK10** | 7,5 | STK6A | 7,4 | STK6A | 8,0 | AURKBA | 7,9 | STK6A | 9,9 | STK6A | 7,7 | AAK1 | 7,7 |
| AURKBA | 8,6 | BMP2KA | 7,7 | AURKBA | 6,6 | AURKBA | 7,3 | AURKBA | 7,6 | STK6A | 7,6 | AURKBA | 9,1 | AURKBA | 7,1 | FLT1A | 4,4 |
| BMP2KA | 7,2 | CHEK2A | 7,7 | CHEK2A | 5,8 | **STK10** | 7,1 | CHEK2A | 7,0 | **STK10** | 7,6 | STK17AA | 7,6 | CHEK2A | 6,3 | GAKA | 3,8 |
| STK6A | 7,0 | STK6A | 7,6 | STK6A | 5,6 | AAK1A | 6,3 | **STK10** | 6,7 | CHEK2A | 6,8 | **STK10** | 6,7 | STK17AA | 6,0 | **STK10** | 3,7 |
| GAKA | 5,4 | GAKA | 5,1 | AAK1A | 5,5 | STK17AA | 5,8 | AAK1A | 6,7 | AAK1A | 6,3 | CHEK2A | 6,5 | AAK1A | 5,9 | K2A2 | 3,6 |
| MAP2K4A | 5,1 | STK3A | 5,1 | GAKA | 4,5 | CHEK2A | 5,1 | GAKA | 6,2 | GAKA | 6,0 | AAK1A | 6,4 | GAKA | 5,7 | BMPR2A | 3,5 |
| STK17BA | 5,1 | MAP2K4A | 5,0 | ULK3A | 4,2 | GAKA | 4,7 | ULK3A | 6,1 | ULK3A | 5,9 | GAKA | 6,2 | **STK10** | 5,6 | STK17BA | 3,4 |
| STK3A | 4,7 | STK17BA | 4,8 | BMPR2A | 4,0 | ULK3A | 4,4 | STK17AA | 5,1 | STK17AA | 5,3 | PLK4A | 5,5 | BMPR2A | 5,1 | STK3 | 3,3 |
| PIM3A | 4,6 | AAK1A | 4,7 | PIM3A | 3,9 | BMPR2A | 4,3 | PLK4A | 4,9 | STK3A | 4,7 | PIM3A | 5,1 | PLK4A | 4,7 | SLK | 3,3 |
| ULK3A | 4,5 | STK4A | 4,3 | STK3A | 3,4 | PIM3A | 4,1 | PIM3A | 4,9 | PLK4A | 4,6 | CLK1A | 4,7 | PIM3A | 4,4 | PLK4A | 3,0 |
| SLK | 4,2 | ULK3A | 4,3 | MAPK15HSD | 3,3 | STK3A | 3,7 | BMPR2A | 4,6 | BMPR2A | 4,4 | ULK3A | 4,5 | DYRK2A | 4,3 | FGFR3A | 3,0 |
| STK17AA | 4,0 | PIM3A | 4,3 | STK17AA | 3,1 | PLK4A | 3,5 | STK3A | 4,4 | STK4A | 4,4 | STK3A | 3,9 | ULK3A | 4,1 | CAMKK2B | 2,9 |
| PLK4A | 3,9 | EPHA7A | 3,5 | PLK4A | 3,0 | MAP2K4A | 3,2 | MAP2K4A | 4,3 | MAP2K4A | 4,0 | DYRK2A | 3,9 | STK3A | 3,6 | CLK1A | 2,9 |
| AAK1A | 3,9 | CDK2A | 3,3 | FGFR2A | 2,9 | DYRK2A | 3,1 | STK4A | 4,1 | SLK | 3,6 | STK4A | 3,8 | MAP2K4A | 3,3 | DAPK3 | 2,8 |
| STK4A | 3,8 | PLK4A | 3,3 | CLK1A | 2,8 | FGFR2A | 3,1 | MAPK15HSD | 4,1 | CDK2A | 3,4 | FGFR2A | 3,7 | CLK1A | 3,1 | UKL3A | 2,7 |
| TTKA | 3,5 | BMPR2A | 3,2 | MAP2K4A | 2,8 | STK4A | 3,0 | SLK | 3,4 | STK17BA | 3,4 | BMPR2A | 3,7 | FGFR2A | 2,8 | STK6A | 2,4 |
| BMXA | 3,0 | MAPK14A | 3,1 | STK4A | 2,6 | SRCA | 2,7 | CDK2A | 3,3 | EPHA7A | 2,9 | MAP2K4A | 3,6 | STK4A | 2,7 | PIM1 | 2,4 |
| EPHA7A | 3,0 | SLK | 3,1 | SLK | 2,5 | CLK1A | 2,5 | SRCA | 3,2 | FGFR2A | 2,8 | MAPK15HSD | 3,3 | STK17BA | 2,7 | MELKA | 2,2 |
| BMPR2A | 2,8 | TTKA | 3,1 | CAMK2DA | 2,1 | CDK2A | 2,5 | FGFR2A | 3,1 | CAMK2DA | 2,7 | CAMKK2B | 3,2 | CAMK2DA | 2,2 | MAPK15A | 2,0 |
| FGFR2A | 2,7 | MARK4A | 2,7 | FESA | 2,0 | CAMK2DA | 2,3 | CAMK2DA | 2,8 | CLK1A | 2,7 | MAP3K5A | 2,8 | MAPK15HSD | 2,1 | DYRK2A | 1,8 |
| MARK4A | 2,7 | BMXA | 2,7 | SRCA | 2,0 | MAPK15HSD | 2,3 | CLK1A | 2,8 | SRCA | 2,6 | GSG2A | 2,8 | EPHA7A | 1,9 | MAP2K4A | 1,8 |
| MAPK14A | 2,6 | CAMK2DA | 2,7 | MARK4A | 1,9 | SLK | 2,1 | STK17BA | 2,6 | MAPK15HSD | 2,5 | CDK2A | 2,7 | DCAMKL1A | 1,9 | GSK3BB | 1,7 |
| CAMK2DA | 2,5 | DCAMKL1A | 2,7 | NEK7A | 1,7 | CAMKK2B | 2,0 | CSNK2A2A | 2,5 | MARK4A | 2,5 | STK17BA | 2,7 | TTKA | 1,8 | MAP3K5A | 1,7 |
| SRCA | 2,5 | PAK4A | 2,4 | CDK2A | 1,7 | STK17BA | 1,8 | PIM1A | 2,0 | TTKA | 2,4 | SRCA | 2,6 | DAPK3A | 1,8 | STK17AA | 1,7 |
| ABL1A | 2,4 | STK17AA | 2,4 | STK17BA | 1,7 | TTKA | 1,6 | MARK4A | 2,0 | MAPK14A | 2,4 | SLK | 2,4 | SLK | 1,6 | PIM3A | 1,6 |
| MAPK15HSD | 2,4 | FGFR2A | 2,3 | CAMKK2B | 1,5 | PAK4A | 1,5 | PCTK1A | 1,9 | CSNK2A2A | 2,2 | CAMK2DA | 2,4 | SRCA | 1,4 | STK39A | 1,6 |
| DCAMKL1A | 2,3 | CLK1A | 2,3 | DYRK2A | 1,5 | EPHA7A | 1,5 | DYRK2A | 1,9 | PAK4A | 2,0 | GPRK5A | 2,2 | PAK4A | 1,4 | FGFR2A | 1,5 |
| CLK1A | 2,3 | NEK7A | 2,3 | FGFR1B | 1,4 | ABL1A | 1,5 | TTKA | 1,8 | MAP3K5A | 2,0 | MARK4A | 2,2 | MAPK14A | 1,3 | CDK2A | 1,4 |
| OSR1A | 2,1 | SRCA | 2,3 | TTKA | 1,4 | MARK4A | 1,5 | EPHA7A | 1,8 | PCTK1A | 2,0 | DCAMKL1A | 2,2 | MELKA | 1,3 | ABL1 | 1,4 |
| PAK4A | 2,1 | BRPF1B | 2,2 | PAK4A | 1,3 | MAP3K5A | 1,5 | CASKA | 1,8 | CAMKK2B | 1,8 | PAK4A | 2,2 | FESA | 1,3 | GSG2A | 1,3 |
| NEK7A | 2,0 | OSR1A | 2,2 | GSK3BB | 1,2 | FGFR1B | 1,4 | CAMKK2B | 1,8 | DYRK2A | 1,8 | MELKA | 2,1 | ABL1A | 1,3 | RIOK1A | 1,2 |
| CDK2A | 1,9 | BRD4A | 2,1 | CASKA | 1,1 | FESA | 1,4 | PAK4A | 1,7 | CASKA | 1,7 | DAPK3A | 2,1 | CSNK2A2A | 1,2 | BRAFA | 1,2 |
| BRPF1B | 1,9 | MAP3K5A | 2,1 | EPHA7A | 1,1 | GSK3BB | 1,3 | ABL1A | 1,7 | PIM1A | 1,7 | TTKA | 2,0 | OSR1A | 1,2 | PCTK1A | 1,0 |
| MST4A | 1,9 | CAMK1DA | 2,1 | MAPK8B | 1,1 | PCTK1A | 1,3 | MELKA | 1,7 | OSR1A | 1,7 | FGFR1B | 1,9 | FGFR1B | 1,2 | STK4A | 0,9 |
| BRD4A | 1,8 | MST4A | 2,1 | PIM1A | 1,0 | MELKA | 1,3 | MAP3K5A | 1,6 | ABL1A | 1,7 | EPHA7A | 1,9 | GSK3BB | 1,1 | GPRK5A | 0,8 |
| CAMK1DA | 1,7 | EPHA5A | 1,9 | MAP3K5A | 1,0 | MAPK8B | 1,2 | AKT3A | 1,6 | DCAMKL1A | 1,6 | PCTK1A | 1,7 | CASKA | 1,0 | MARK4A | 0,8 |
| CASKA | 1,6 | PAK1A | 1,8 | MELKA | 1,0 | DAPK3A | 1,2 | FGFR1B | 1,5 | BRPF1B | 1,5 | CSNK2A2A | 1,7 | STK39A | 1,0 | CHEK2A | 0,8 |
| EPHA5A | 1,6 | CAMKK2B | 1,8 | PCTK1A | 1,0 | OSR1A | 1,2 | MAPK14A | 1,5 | STK39A | 1,4 | MAPK1A | 1,6 | CDK2A | 1,0 | PKMYT1A | 0,8 |
| CAMKK2B | 1,5 | DYRK1AA | 1,6 | OSR1A | 0,9 | DCAMKL1A | 1,2 | DCAMKL1A | 1,4 | MELKA | 1,4 | GSK3BB | 1,6 | EPHA5A | 1,0 | FGFR1B | 0,7 |
| PAK1A | 1,5 | TIF1A | 1,6 | MAPK14A | 0,9 | STK39A | 1,2 | BMXA | 1,4 | CAMK1DA | 1,4 | PIM1A | 1,6 | MAPK8B | 0,9 | CLK3A | 0,7 |
| STK39A | 1,5 | TLK1A | 1,6 | DCAMKL1A | 0,9 | NEK7A | 1,0 | OSR1A | 1,3 | NEK7A | 1,3 | ABL1A | 1,5 | MARK4A | 0,9 | DAPK1 | 0,7 |
| MAP3K5A | 1,4 | FGFR1B | 1,5 | STK39A | 0,8 | PIM1A | 1,0 | DAPK3A | 1,2 | FGFR1B | 1,2 | OSR1A | 1,5 | NEK7A | 0,9 | RPS6KA1A | 0,7 |
| MARK3A | 1,4 | RPS6KA5A | 1,5 | ABL1A | 0,7 | MAPK14A | 0,9 | STK39A | 1,2 | MST4A | 1,2 | MAPK14A | 1,4 | MAP3K5A | 0,8 | SRCA | 0,7 |
| EPHB3A | 1,4 | MARK3A | 1,4 | MAP2K6A | 0,7 | MAP2K6A | 0,9 | GSG2A | 1,2 | DAPK3A | 1,2 | EPHA5A | 1,4 | MAP2K6A | 0,8 | DCAMKL1A | 0,6 |
| RPS6KA5A | 1,3 | EPHB3A | 1,4 | MAPK1A | 0,7 | CASKA | 0,9 | MAP2K6A | 1,2 | EPHA5A | 1,2 | NEK7A | 1,4 | CAMK1DA | 0,8 | EPHB3A | 0,6 |
| TIF1A | 1,3 | STK39A | 1,3 | CSNK1EA | 0,7 | CAMK4A | 0,8 | EPHA5A | 1,2 | BRD4A | 1,2 | STK39A | 1,2 | MAPK1A | 0,7 | CAMK2BA | 0,6 |
| FGFR1B | 1,3 | DAPK3A | 1,2 | EPHA5A | 0,6 | CSNK2A2A | 0,8 | CAMK1DA | 1,2 | TLK1A | 1,2 | MST4A | 1,2 | MAP2K1A | 0,7 | EPHA2 | 0,6 |
| TLK1A | 1,3 | DYRK2A | 1,2 | MAP2K1A | 0,6 | EPHA5A | 0,7 | MAPK8B | 1,1 | BMXA | 1,1 | BRPF1B | 1,2 | CAMKK2B | 0,6 | MARK3A | 0,5 |
| DAPK3A | 1,2 | MAP2K7A | 1,1 | CAMK4A | 0,6 | GSG2A | 0,7 | RPS6KA1A | 1,1 | RPS6KA5A | 1,1 | MAP2K6A | 1,1 | CAMK4A | 0,6 | CSNK1DA | 0,5 |
| CAMK4A | 1,2 | MAP2K6A | 1,1 | GSG2A | 0,6 | TLK1A | 0,7 | CAMK4A | 1,1 | MAP2K6A | 1,0 | CAMK1DA | 1,1 | PIM1A | 0,6 | PDK4A | 0,5 |
| MAP2K6A | 1,2 | MAPK10A | 1,1 | RPS6KA1A | 0,5 | CSNK1EA | 0,7 | TLK1A | 1,1 | PAK1A | 1,0 | CAMK4A | 1,0 | RPS6KA5A | 0,6 | CAMK2DA | 0,5 |
| PCTK1A | 1,1 | PKMYT1A | 1,1 | TIF1A | 0,5 | RPS6KA1A | 0,6 | DMPK1A | 1,0 | GSG2A | 1,0 | MAPK8B | 0,9 | PCTK1A | 0,5 | MAPK13A | 0,4 |
| PIM1A | 1,1 | PCTK1A | 1,0 | MSSK1A | 0,5 | MAP2K1A | 0,6 | GSK3BB | 1,0 | CAMK4A | 0,9 | AKT3A | 0,9 | DAPK1A | 0,5 | STK38LA | 0,4 |
| MAP2K7A | 1,0 | FESA | 1,0 | DAPK3A | 0,5 | MAPK1A | 0,6 | BRPF1B | 1,0 | AKT3A | 0,8 | DYRK1AA | 0,9 | BRPF1B | 0,5 | MAP2K6A | 0,4 |
| DYRK1AA | 1,0 | PIM1A | 1,0 | HIPK2HSF | 0,5 | GPRK5A | 0,5 | MST4A | 0,9 | RPS6KA1A | 0,8 | TLK1A | 0,9 | EPHB3A | 0,5 | MST3A | 0,4 |
| CDC42BPAA | 1,0 | MAP2K1A | 1,0 | GPRK5A | 0,4 | MAPK10A | 0,5 | GPRK5A | 0,9 | MAPK10A | 0,8 | CASKA | 0,9 | HIPK2HSF | 0,4 | WNK1A | 0,4 |
| MAPK1A | 0,9 | CAMK4A | 1,0 | TLK1A | 0,4 | ULK1A | 0,5 | MAPK1A | 0,9 | MAPK8B | 0,8 | EPHB3A | 0,8 | MSSK1A | 0,4 | MERTK | 0,4 |
| DYRK2A | 0,8 | ABL1A | 0,9 | CAMK1DA | 0,4 | EPHA2A | 0,4 | EPHB3A | 0,8 | FESA | 0,8 | MAP2K1A | 0,7 | TLK1A | 0,4 | NEK1A | 0,4 |
| FESA | 0,7 | MAPK1A | 0,9 | EPHB3A | 0,4 | MSSK1A | 0,4 | FESA | 0,8 | GPRK5A | 0,7 | MAPK10A | 0,7 | MST4A | 0,4 | MAPK9A | 0,4 |
| PKMYT1A | 0,7 | MELKA | 0,9 | EPHA2A | 0,4 | HIPK2HSF | 0,4 | RPS6KA5A | 0,8 | MAP2K1A | 0,7 | BRD4A | 0,7 | AKT3A | 0,4 | TTKA | 0,3 |
| CSNK2A2A | 0,7 | CDC42BPAA | 0,9 | ULK1A | 0,4 | BMXA | 0,4 | MSSK1A | 0,7 | MAPK1A | 0,7 | DMPK1A | 0,7 | RPS6KA1A | 0,4 | BMX | 0,3 |
| MAPK10A | 0,7 | GSG2A | 0,9 | MAPK10A | 0,3 | DMPK1A | 0,4 | MAPK10A | 0,7 | EPHB3A | 0,6 | RPS6KA5A | 0,7 | CSNK1EA | 0,4 | MAPK1A | 0,3 |
| GSG2A | 0,7 | MAPK8B | 0,9 | CSNK2A2A | 0,3 | CAMK1DA | 0,4 | CDKL1A | 0,6 | DMPK1A | 0,6 | CSNK1EA | 0,6 | EPHA2A | 0,3 | ULK1A | 0,2 |
| MAP2K1A | 0,7 | GPRK5A | 0,7 | CDC42BPAA | 0,3 | EPHB3A | 0,3 | MAP2K1A | 0,6 | DYRK1AA | 0,6 | TIF1A | 0,6 | EPHA4A | 0,3 | DMPK1A | 0,2 |
| MAPK8B | 0,5 | ULK1A | 0,6 | BRD4A | 0,3 | VRK1A | 0,3 | BRD4A | 0,6 | GSK3BB | 0,6 | CDKL1A | 0,5 | PAK1A | 0,3 | SRPK1A | 0,2 |
| MSSK1A | 0,5 | CDKL1A | 0,6 | DMPK1A | 0,3 | TIF1A | 0,3 | ULK1A | 0,6 | TIF1A | 0,5 | FESA | 0,5 | CDKL1A | 0,2 | TAF1A | 0,1 |
| ULK1A | 0,4 | VRK1A | 0,6 | EPHA4A | 0,3 | RPS6KA5A | 0,3 | CSNK1EA | 0,6 | ULK1A | 0,5 | EPHB1A | 0,5 | GPRK5A | 0,2 | CASKA | 0,0 |
| VRK1A | 0,4 | HIPK2HSF | 0,6 | AKT3A | 0,3 | EPHA4A | 0,3 | NEK7A | 0,6 | CDKL1A | 0,5 | PAK1A | 0,5 | CDC42BPAA | 0,1 | AKT3A | 0,0 |
| CDKL1A | 0,4 | MAPK15HSD | 0,5 | RPS6KA5A | 0,2 | CDKL1A | 0,3 | HIPK2HSF | 0,5 | HIPK2HSF | 0,5 | CLK3A | 0,5 | GSG2A | 0,1 | CAMK4A | 0,0 |
| MELKA | 0,4 | EPHB1A | 0,5 | MAPKAPK2A | 0,2 | MST4A | 0,3 | PAK1A | 0,5 | MAP2K7A | 0,5 | RPS6KA1A | 0,5 | EPHB1A | 0,1 | CAMK1GA | 0,0 |
| EPHA4A | 0,3 | EPHA4A | 0,5 | MST4A | 0,2 | PAK1A | 0,2 | MAP2K7A | 0,5 | MARK3A | 0,5 | EPHA2A | 0,5 | MAPKAPK2A | 0,1 | OSR1A | 0,0 |
| EPHB1A | 0,3 | CASKA | 0,5 | SRPK1A | 0,2 | CSNK1DA | 0,2 | EPHA2A | 0,4 | MSSK1A | 0,4 | DAPK1A | 0,5 | MAPK10A | 0,1 | VRK1A | -0,1 |
| GPRK5A | 0,2 | CSNK2A2A | 0,5 | CSNK1DA | 0,2 | AKT3A | 0,2 | EPHA4A | 0,4 | EPHA4A | 0,4 | HIPK2HSF | 0,5 | BMXA | 0,1 | PAK1A | -0,1 |
| DMPK1A | 0,2 | CLK3A | 0,4 | CDKL1A | 0,2 | EPHB1A | 0,2 | EPHB1A | 0,4 | DAPK1A | 0,4 | EPHA4A | 0,4 | MARK3A | 0,0 | PHKG2A | -0,1 |
| CLK3A | 0,2 | NEK2A | 0,4 | CLK3A | 0,2 | CLK3A | 0,2 | DAPK1A | 0,3 | PKMYT1A | 0,4 | MSSK1A | 0,4 | CLK3A | 0,0 | FESA | -0,2 |
| RPS6KA1A | 0,2 | MAPKAPK2A | 0,4 | MAP2K7A | 0,1 | MAPKAPK2A | 0,2 | PKMYT1A | 0,3 | EPHB1A | 0,3 | BMXA | 0,3 | MAP2K7A | 0,0 | MAP2K1A | -0,2 |
| HIPK2HSF | 0,2 | AKT3A | 0,4 | EPHB1A | 0,1 | CDC42BPAA | 0,2 | DYRK1AA | 0,3 | EPHA2A | 0,3 | VRK1A | 0,3 | VRK1A | 0,0 | CDKL1A | -0,2 |
| EPHA2A | 0,2 | DAPK1A | 0,4 | PAK1A | 0,1 | BRPF1B | 0,2 | MAPKAPK2A | 0,2 | CSNK1EA | 0,3 | MARK3A | 0,3 | DMPK1A | 0,0 | MAPK14A | -0,3 |
| DAPK1A | 0,1 | MSSK1A | 0,3 | VRK1A | 0,0 | DAPK1A | 0,1 | VRK1A | 0,2 | CLK3A | 0,2 | ULK1A | 0,2 | BRD4A | -0,1 | RPS6KA6A | -0,3 |
| MAPKAPK2A | 0,1 | CAMK2BA | 0,3 | MAPK13A | 0,02 | SRPK1A | 0,1 | MARK3A | 0,2 | VRK1A | 0,2 | PKMYT1A | 0,2 | ULK1A | -0,1 | PAK4A | -0,3 |
| TAF1A | 0,1 | TAF1A | 0,3 | TAF1A | 0,0 | BRD4A | 0,1 | TIF1A | 0,2 | TAF1A | 0,2 | CAMK2BA | 0,1 | TIF1A | -0,1 | TOPKA | -0,4 |
| CSNK1EA | 0,0 | DMPK1A | 0,3 | BRPF1B | -0,1 | MAP2K7A | 0,1 | CSNK1DA | 0,1 | MAPKAPK2A | 0,2 | MAPKAPK2A | 0,1 | CSNK1DA | -0,1 | BRD4 | -0,4 |
| SRPK1A | -0,1 | GSK3BB | 0,2 | DAPK1A | -0,1 | PKMYT1A | 0,0 | SRPK1A | 0,1 | CDC42BPAA | 0,1 | CDC42BPAA | 0,1 | PKMYT1A | -0,1 | MST4A | -0,4 |
| MAPK9A | -0,1 | RPS6KA1A | 0,2 | BMXA | -0,1 | TAF1A | -0,1 | CDC42BPAA | 0,0 | SRPK1A | 0,0 | TAF1A | 0,1 | DYRK1AA | -0,1 | EPHA5A | -0,5 |
| CAMK2BA | -0,1 | EPHA2A | 0,1 | MARK3A | -0,1 | MAPK13A | -0,1 | CLK3A | 0,0 | CAMK2BA | 0,0 | CSNK1DA | 0,1 | SRPK1A | -0,1 | CDC42BPAA | -0,6 |
| AKT3A | -0,1 | SRPK1A | 0,0 | PKMYT1A | -0,1 | BRAFA | -0,2 | TAF1A | 0,0 | BRAFA | -0,2 | SRPK1A | 0,0 | TAF1A | -0,3 | MAPK10A | -0,7 |
| GSK3BB | -0,2 | NEK1A | 0,0 | CAMK2BA | -0,2 | CAMK2BA | -0,2 | CAMK2BA | -0,1 | CSNK1DA | -0,2 | MAP2K7A | 0,0 | CAMK2BA | -0,3 | CAMK1DA | -1,2 |
| NEK1A | -0,4 | BRAFA | 0,0 | NEK1A | -0,3 | NEK1A | -0,2 | NEK1A | -0,4 | MAPK13A | -0,3 | NEK1A | -0,1 | MAPK13A | -0,3 | EPHA7A | -1,4 |
| BRAFA | -0,4 | CSNK1EA | -0,2 | NEK2A | -0,3 | MARK3A | -0,3 | BRAFA | -0,5 | NEK1A | -0,4 | MAPK13A | -0,3 | NEK1A | -0,4 | DYRK1AA | -1,5 |
| NEK2A | -0,4 | MAPK9A | -0,2 | BRAFA | -0,4 | DYRK1AA | -0,4 | NQO2A | -0,6 | MAPK9A | -0,6 | BRAFA | -0,6 | NQO2A | -0,5 | | |
| MAPK13A | -0,5 | MAPK13A | -0,5 | DYRK1AA | -0,5 | NEK2A | -0,5 | MAPK13A | -1,3 | NEK2A | -1,3 | MAPK9A | -1,2 | BRAFA | -0,5 | | |
| CSNK1DA | -0,5 | CSNK1DA | -0,5 | NQO2A | -0,5 | NQO2A | -0,5 | NEK2A | -1,3 | NQO2A | -2,8 | NEK2A | -1,3 | NEK2A | -1,3 | | |
| MST3A | -3,4 | NQO2A | -2,7 | MAPK9A | -2,1 | MAPK9A | -1,9 | MAPK9A | -1,3 | PIM3A | -4,8 | NQO2A | -1,5 | MAPK9A | -2,0 | | |
| NQO2A | -4,1 | MST3A | -4,5 | FECHHSB | -3,1 | FECHHSB | -3,8 | MST3A | -5,3 | MST3A | -6,4 | MST3A | -5,3 | MST3A | -7,8 | | |
| | | | | MST3A | -7,7 | MST3A | -5,1 | | | | | | | | | | |

**Table S2:** Data collection and Refinement Statistics.

| **Data collection** | **STK10 / compound 23** |
|---|---|
| Beamline | X06DA/PXI SLS |
| Wavelength (Å) | 1.000041 |
| Space group | C 1 2 1 |
| Cell dimensions | |
| *a, b, c (Å)* | 93.3, 61.84, 56.81 |
| α, β, γ (°) | 90.00, 102.79, 90.00 |
| Resolution (Å)* | 40.18-1.90 (1.94-1.90) |
| unique observations* | 24697 (1600) |
| *Rmeas** | 0.05 (0.275) |
| Completeness (%)* | 99.1 (99.6) |
| Multiplicity* | 3.8 (4.0) |
| mean I/σI* | 11.6 (3.8) |
| CC1/2* | 0.998 (0.967) |
| **Refinement** | |
| *Rwork / Rfree* | 19.95 / 23.68 |
| No. of atoms | 2390 (2186 protein) |
| overall B-factors (Å$^2$) | 41.0 |
| Rms deviations | |
| Bond lengths (Å) | 0.008 |
| Bond angles (°) | 1.447 |
| Ramachandran (%) | |
| outlier | 0.4 |
| **Protein Data Bank entry** | **7QGP** |

*Values for the highest resolution shell are shown in parentheses.

#### Experimental Section

**DSF-based selectivity screening against a curated kinase library ($T_m$ Panel).** The assay was performed as previously described.[1,2] Briefly, recombinant protein kinase domains at a concentration of 2 µM were mixed with 10 µM compound in a buffer containing 20 mM HEPES, pH 7.5, and 500 mM NaCl. SYPRO Orange (5000×, Invitrogen) was added as a fluorescence probe (1 µl per mL). Subsequently, temperature-dependent protein unfolding profiles were measured using the QuantStudio™ 5 realtime PCR machine (Thermo Fisher). Excitation and emission filters were set to 465 nm and 590 nm, respectively. The temperature was raised with a step rate of 3 °C per minute. Data points were analysed with the internal software (Thermal Shift SoftwareTM Version 1.4, Thermo Fisher) using the Boltzmann equation to determine the inflection point of the transition curve.

**NanoBRET Target Engagement assay.** The assay was performed as described previously.[3] In brief, STK10, STK3, STK4 and SLK were obtained as plasmids cloned in frame with terminal NanoLuc-fusion (gift from Promega). Plasmids were transfected into human embryonic kidney 293T cells [American Type Culture Collection (ATCC), CRL-3216; RRID: CVCL_0063] using FuGENE 4 K (Promega, E5911), and proteins were allowed to express for 20 hours. Serially diluted inhibitor and K10 Tracer (Promega, N2642) at the Tracer $K_D$ concentration taken from TracerDB (tracerdb.org)[4] were pipetted into white 384-well plates (Greiner, 781207) using an ECHO acoustic dispenser (Labcyte). The corresponding protein-transfected cells were added and reseeded at a density of $2 \times 10^5$ cells/ml after trypsinization and resuspending in Opti-MEM without phenol red (Life Technologies). The system was allowed to equilibrate for 2 hours at 37°C/5% $CO_2$ before bioluminescence resonance energy transfer (BRET) measurements. To measure BRET, NanoBRET NanoGlo Substrate + extracellular NanoLuc Inhibitor (Promega, N2540) was added as per the manufacturer's protocol, and filtered luminescence was measured on a PHERAstar plate reader (BMG Labtech) equipped with a luminescence filter pair [450-nm BP filter (donor) and 610-nm LP filter (acceptor)]. Competitive displacement data were then graphed using

GraphPad Prism 9 software (www.graphpad.com/, RRID:SCR_002798) using a normalized three-parameter curve fit with the following equation: $Y = 100/[1 + 10(X - \log IC50)]$.

**PhosphoSens Assay.** The kinase activity of STK10 and SLK was assessed using the PhosphoSense® kinase as with a SOX-based substrate peptide (AQT0505, Assay Quant Technologies). Both proteins were used at a concentration of 25 nM. The ATP concentrations were determined by ATP titration experiments, which were used to measure the dissociation constant (KD) of ATP. The ATP concentrations were therefore set at 150 µM and 160 µM for the SLK and STK10 experiments, respectively. A serial dilution of each test compound was prepared across 11 concentrations ranging from 40 µM to 0.4 nM. These were dispensed in quadruplicate into white 384-well plates (Greiner 781207) using an ECHO 550 acoustic dispenser (Labcyte). For each compound, two wells served as 0% control (lacking both protein and compound) and two wells as 100% control (lacking only the compound).

Purified SLK protein was diluted to a final concentration of 25 nM in reaction buffer [50 mM HEPES (pH 7.5), 10 mM $MgCl_2$, 150 mM NaCl, 1% glycerol, 1 mM DTT, 0.2 mg/ml bovine serum albumin (BSA), 0.01% Tween 20, and 4 µM AQT0505]. Then, 10 µL of this solution was added to each well using a Mantis liquid dispenser (Formulatrix). The 0% control wells received 10 µL of reaction buffer without protein. Subsequently, 15 nL of 100 mM ATP was added to each well via ECHO. Plates were centrifuged at 1500 × g for 2 minutes and incubated at room temperature for 1 hour.

For STK10, the purified protein was diluted to 50 nM in a double concentrated reaction buffer [100 mM HEPES (pH 7.5), 20 mM $MgCl_2$, 300 mM NaCl, 2% glycerol, 2 mM DTT, 0.4 mg/mL BSA, 0.02% Tween 20, and 320 µM ATP]. A volume of 5 µl of this solution was pipetted into each well using a multichannel pipette (Eppendorf). Plates were incubated at 37 °C for 1 hour. For substrate addition, 5 µL of 8 µM AQT0505 peptide was added to each well using the Mantis dispenser. Plates were then centrifuged at 1500 × g for 2 minutes and incubated at room temperature for an additional hour.

Fluorescence of the SOX-peptide was measured after excitation at 360 nm and emission at 487 nm using a PHERAstar plate reader (BMG Labtech). $IC_{50}$ values were calculated by nonlinear regression of log[inhibitor] versus normalized response using GraphPad Prism 8 (www.graphpad.com, RRID:SCR_002798).

**SPR.** Surface plasmon resonance (SPR) experiments were conducted using a Biacore T200 instrument equipped with a Series S Sensor Chip NTA (Cytiva) at 25 °C. His-STK10 was immobilized at a flow rate of 10 µL/min on flow channels (FC) 2 while FC1 served as a blank reference without protein. Nickel loading of the chip was achieved by injecting 500 µM $NiCl_2$. To activate the chip surface for covalent coupling, 483 mM EDC and 10 mM NHS were mixed in a 1:1 (v/v) ratio and injected. Subsequently, 100 nM His-STK10 was injected over FC2 yielding an immobilization level of 1500 RU. The surface was deactivated, and nickel was stripped by injecting 1 M ethanolamine and 350 mM EDTA. HBS-P+ containing 1 mM TCEP was used as both the immobilization buffer and protein-dilution buffer, while HBS-P+ with 1 mM TCEP and 2% DMSO was used as the running buffer for analyte-protein interactions.

Compound 23 was measured in duplicates using Multi-Cycle Kinetics, with 10 concentrations ranging from 0.01 µM to 5 µM injected across FC1 and FC2 at 30 µL/min for 150 s. Interaction data was FC1-referenced, blank-subtracted, solvent-corrected, and analyzed using Biacore T200 Evaluation Software version 3.2.1. To determine $K_D$ affinity values, data fitting was performed using a 1:1 Langmuir kinetic model.

**Protein expression and purification.** STK10 and the STK10 surface entropy reduction mutant STK10K272A (residues R18-E317) with a TEV-cleavable N-terminal His6-tag were purified in exactly the same manner. Plamids were transformed in BL21(DE3) cells and grown in Terrific Broth medium containing 50 mg/mL kanamycin. Protein expression was induced at an OD600 of 2 by using 0.5 mM isopropyl-thio-galactopyranoside (IPTG) at 18 C for 12 hours. Cells expressing the protein were lysed in lysis buffer containing 50 mM HEPES pH 7.5, 500 mM NaCl, 25 mM imidazole, 5% glycerol, and 0.5 mM Tris(2 carboxyethyl)phosphine (TCEP) by sonication. After centrifugation, the supernatant was loaded onto a Nickel-Sepharose column equilibrated with 30 mL lysis buffer. The column was washed with 60 mL lysis buffer. Proteins were eluted by an imidazole step gradient (50, 100, 200, 300 mM). Fractions containing protein were pooled together and dialyzed overnight using 1 L of final buffer (25 mM HEPES pH 7.5, 300 mM NaCl, 0.5 mM TCEP) at 4 C. Additionally, TEV protease was added (protein:TEV 1:20 molar ratio) to remove the tag. Next day the protein solution was loaded onto 2 mL Nickel-Sepharose column beads to remove the TEV protease and uncleaved Tag. The combined flow through fraction and the wash fraction (25 mM imidazole) containing the protein were concentrated to approximately 4-5 mL and loaded onto Superdex 75 16/60 Hi-Load gel filtration column equilibrated with final buffer. The protein was concentrated to approximately 10 mg/mL.

**Crystallization.** STK10 K272A (10 mg/mL) was co-crystallized with **23** (final conc. approx. 500 µM) using the sitting-drop vapor diffusion method by mixing protein and well solutions in 2:1, 1:1, and 1:2 ratios (Drop size 200 nL) at 4 °C. The reservoir solution contained 13% PEG 1.000 and 10% PEG 8.000.

**Data collection, structure solution and refinement.** Diffraction data were collected at beamline X06DA (Villigen, CH) at a wavelength of 1.0 Å at 100 K. The reservoir solution supplemented with 20% ethylene glycol was used as cryoprotectant. Data were processed using XDS[5] and scaled with aimless[6]. The PDB structure with the accession code 6EIM was used as an initial search MR model using the program PHASER[7]. The final model was built manually using Coot[8] and refined with REFMAC5[9], which are all part of the CCP4 suite[10]. Data collection and refinement statistics are summarized in **Table S2**. Dictionary file for the ligand was generated using the Grade Web Server (http://grade.globalphasing.org).

**Chemistry.** The synthetic routes of compounds will be outlined in the following and the analytical data can be found in the Supporting Information. All commercial chemicals were purchased from BLD Pharm, TCI, Merck, and Enamine with a purity ≥95 % and were used without further purification. The solvents with an analytical grade were obtained from VWR Chemicals and Merck and all dry solvents from Acros Organics. All reactions were carried out under an argon atmosphere. The thin layer chromatography was done with silica gel on aluminium foils (60 Å pore diameter) obtained from Macherey-Nagel and visualized with ultraviolet light (λ = 254 and 365 nm). The purification of the compounds was performed by flash chromatography using puriFlash XS 420 device with a UV-VIS multiwave detector (200 – 400 nm) from Interchim. Pre-packed normal-phase PF-SIHP and pre-packed PF-C18HP reverse phase columns with particle sizes of 15 and 30 µm (Interchim) were used for purification. Preparative purification by HPLC was carried out on an Agilent 1260 Infinity II device using an Eclipse XDB-C18 (Agilent, 21.2 x 250mm, 7µm) reversed phase column. A gradient was used with 0.1% TFA in water (A) and 0.1% TFA in acetonitrile (B) (flow rate 21 mL/min), as a mobile phase. DMSO was evaporated using a SpeedVac Concentrator (ThermoFischer Scientific, Model#: SPD121P-230) coupled to a Savant RVT5105 Refrigerated Vapor Trap (ThermoFischer Scientific) and a VLP80 vacuum pump (ThermoScientific). Nuclear magnetic resonance spectroscopy (NMR) was performed with AV400, AV500, AV600 MHz spectrometers from Bruker. Chemical shifts (δ) are reported in parts per million (ppm). DMSO-$d_6$ was used as solvent, and the spectra were calibrated to the solvent signal: 2.50 ppm (1H NMR) or 39.52 ppm (13C NMR). Coupling constants (*J*) were reported in hertz (Hz) and multiplicities were designated as followed: s (singlet), d (doublet), dd (doublet of doublet), t (triplet), dt (doublet of triplets), td (triplet of doublets), ddd (doublet of doublet of doublet), q (quartet), m (multiplet). Mass spectra were measured on a Surveyor MSQ device from ThermoFisher measuring in the positive- or negative-ion mode. Final compounds were additionally characterized by HRMS using a MALDI LTQ Orbitrap XL from ThermoScientific and a microOTOF-Q ESI source. The purity of the final compounds was determined by HPLC using an Agilent 1260 Infinity II device with a 1260 DAD HS detector (G7117C; 254 nm, 280 nm, 310 nm) and a LC/MSD device (G6125B, ESI pos. 100–1000). The compounds were analyzed on a Poroshell 120 EC-C18 (Agilent, 3 × 150 mm, 2.7 µm) reversed phase column using 0.1 % formic acid in water (A) and 0.1 % formic acid in acetonitrile (B) as a mobile phase. The following gradient was used: Method 1 (M1): 0 min: 5% B - 2 min: 80% B – 5 min: 95% B - 7 min: 95% B (flow rate of 0.6 mL/min). Method 2 (M2): 0 min: 5% B – 2.8 min: 75% B – 7.2

min: 100% B - 10 min: 100% B (flow rate of 0.6 mL/min). UV-detection was performed at 254, 280 and 320 nm and all compounds used for further biological characterization showed a purity ≥95% unless stated otherwise.

**Synthesis of 3-Bromo-5-chloropyrazolo[1,5-*a*]pyrimidine** (**6**). 5-Chloropyrazolo[1,5-*a*]pyrimidine (**5**) (20.00 g, 1.0 eq, 130.23 mmol) was suspended in acetonitrile (250 mL) and *N*-bromosuccinimide (25.50 g, 1.1 eq, 143.26 mmol) was added. After 1 h at room temperature, the reaction was quenched with water. The resulting precipitate was filtered off, washed with water, and dried to afford the product as a bright yellow solid (28.90 g, 96%). $^{1}H$ NMR (500 MHz, DMSO-$d_6$): δ 9.22 (d, J = 7.2 Hz, 1H), 8.44 (s, 1H), 7.23 (d, J = 7.2 Hz, 1H) ppm. $^{13}C$ NMR (126 MHz, DMSO-$d_6$) δ 151.25, 145.68, 143.83, 139.04, 110.07, 82.90 ppm. MS-ESI m/z $[M+H]^+$: calcd 233.92, found 233.90.

**Synthesis of 2-[2-({3-Bromopyrazolo[1,5-*a*] pyrimidine-5-yl}amino) ethoxy]ethan-1-ol** (**7**). (**6**) (3.45 g, 14.85 mmol, 1.0 eq) was dissolved in acetonitrile (35 mL) and *N*,*N*-diisopropylethylamine (2.88 g, 22.26 mmol, 1.5 eq) and 2-(2-aminoethoxy)ethanol (1.87 g, 17.81 mmol, 1.2 eq) were added. The reaction mixture was stirred for 16 h at 85 °C after which time the solvent was removed *in vacuo*. The residue was taken up in ethyl acetate and washed with water and brine. The aqueous phase was extracted with ethyl acetate (3x). The combined organic phases were dried over $MgSO_4$, filtered over cotton, and purified by flash column chromatography on silica gel using cyclohexane/ethyl acetate as an eluent to afford the product as a light yellow solid (3.57 g, 80%). $^{1}H$ NMR (400 MHz, DMSO-$d_6$) δ 8.45 (d, J = 7.5 Hz, 1H), 7.87 (s, 1H), 7.71 (t, J = 5.3 Hz, 1H), 6.34 (d, J = 7.6 Hz, 1H), 4.59 (t, J = 5.3 Hz, 1H), 3.62 – 3.57 (m, 2H), 3.57 – 3.48 (m, 5H), 3.50 – 3.45 (m, 2H) ppm. $^{13}C$ NMR (101 MHz, DMSO-$d_6$) δ 156.14, 145.05, 143.15, 135.36, 100.94, 77.15, 72.18, 68.57, 60.21 ppm. MS-ESI m/z $[M+H]^+$: calcd 301.02, found 301.05.

**Synthesis of 3-bromo-5-(8,8,9,9-tetramethyl-4,7-dioxa-1-aza-8-siladecan-1-yl)pyrazolo-[1,5-*a*]pyrimidine** (**8**). (**7**) (3.57 g, 11.85 mmol, 1.0 eq), *tert*-butyldimethylsilyl chloride (2.68 g, 17.78 mmol, 1.2 eq) and imidazole (2.02 g, 29.64 mmol, 2.5 eq) were solved in DMF (40 mL) and stirred for 1 h at room temperature after which time the solvent was removed *in vacuo*.

The residue was taken up in ethyl acetate and washed with water and brine. The aqueous phase was extracted with ethyl acetate (3x). The combined organic phases were dried over $MgSO_4$, filtered over cotton, and purified by flash column chromatography on silica gel using cyclohexane/ethyl acetate as an eluent to afford the product as a yellow oil (4.29 g, 87%). $^1H$ NMR (400 MHz, DMSO-$d_6$) δ 8.44 (d, J = 7.6 Hz, 1H), 7.86 (s, 1H), 7.72 (t, J = 5.5 Hz, 1H), 6.33 (d, J = 7.5 Hz, 1H), 3.72 – 3.67 (m, 2H), 3.64 – 3.59 (m, 2H), 3.55 – 3.49 (m, 4H), 0.83 (s, 9H), 0.01 (s, 6H) ppm. $^{13}C$ NMR (101 MHz, DMSO-$d_6$) δ 156.14, 145.04, 143.12, 135.32, 100.96, 77.14, 71.68, 68.67, 62.25, 25.79, 17.96, -5.29 ppm. MS-ESI m/z $[M+H]^+$: calcd 415.11, found 415.15.

**Synthesis of *tert*-butyl *N*-{3-bromopyrazolo[1,5-*a*]pyrimidine-5-yl}-N-(2-{2-[(*tert*-butyl-dimethylsilyl)oxy]ethoxy}ethyl)carbamate** (**9**). Triethylamine (1.25 g, 12.39 mmol, 1.2 eq) was added to a solution of (**8**) (4.29 g, 10.32 mmol), di-*tert*-butyl dicarbonate (5.63 g, 25.81 mmol, 2.5 eq) and 4-dimethylaminopyridine (0.025 g, 0.21 mmol, 0.02 eq) in THF (60 mL). The reaction mixture was stirred for 16 h at 75 °C after which time the solvent was removed *in vacuo*. The residue was taken up in ethyl acetate and washed with water and brine. The aqueous phase was extracted with ethyl acetate (3x). The combined organic phases were dried over $MgSO_4$, filtered over cotton, and purified by flash column chromatography on silica gel using cyclohexane/ethyl acetate as an eluent to afford the product as a yellow oil (5.00 g, 94%). $^1H$ NMR (400 MHz, DMSO-$d_6$) δ 8.95 (d, J = 7.7 Hz, 1H), 8.24 (s, 1H), 7.42 (d, J = 7.8 Hz, 1H), 4.18 (t, J = 6.1 Hz, 2H), 3.69 (t, J = 6.1 Hz, 2H), 3.58 (t, J = 5.0 Hz, 2H), 3.46 (t, J = 5.0 Hz, 2H), 1.51 (s, 9H), 0.78 (s, 9H), -0.06 (s, 6H) ppm. $^{13}C$ NMR (101 MHz, DMSO-$d_6$) δ 153.81, 152.78, 144.56, 143.06, 136.46, 105.16, 82.40, 81.16, 71.73, 68.20, 62.30, 45.31, 27.62, 25.70, 17.88, -5.43 ppm.

**Synthesis of Methyl 2-hydroxy-4(4,4,5,5-tetramethyl-1,3,2-dioxaborolan-2-yl)benzoate** (**11**). (**10**) (1.00 g, 4.33 mmol, 1.0 eq), [1,1′-Bis-(diphenylphosphino)-ferrocen]-dichloro-palladium(II) (158 mg, 216 µmol, 0.05 eq), potassium acetate (1.27 g, 13.0 mmol, 3.0 eq) and Bis(pinacolato)diboron (1.21 g, 4.76 mmol, 1.1 eq) were dissolved in 1,4-dioxane (25 mL) and degassed with Argon. The reaction mixture was stirred for 3 h at 100 °C after which time the solvent was removed *in vacuo*. The residue was taken up in ethyl acetate and washed with water and brine. The aqueous phase was extracted with ethyl acetate (3x). The combined organic phases were dried over $MgSO_4$, filtered over cotton, and purified by flash column chromatography on silica gel using cyclohexane/ethyl acetate as an eluent to afford the product as a colourless solid (903 mg, 75%). $^1$H NMR (400 MHz, DMSO-$d_6$) 10.30 (s, 1H), 7.76 (d, J = 7.7 Hz, 1H), 7.23 – 7.17 (m, 2H), 3.88 (s, 3H), 1.30 (s, 12H) ppm. $^{13}$C NMR (101 MHz, DMSO-$d_6$) δ 168.73, 158.82, 129.44, 124.63, 122.96, 115.70, 84.09, 52.42, 24.60 ppm. MS-ESI m/z $[M+H]^+$: calcd 279.13, found 279.10.

**Synthesis of Methyl 4-[5-(2,2,3,3,13,13-hexamethyl-11-oxo-4,7,12-trioxa-10-aza-3-silatetra decan-10-yl)pyrazolo[1,5-*a*]pyrimidine-3-yl]-2-hydroxybenzoate** (**12**). (**9**) (1.00 g, 1.94 mmol, 1.0 eq), (**11**) (701 mg, 2.52 mmol, 1.3 eq), potassium phosphate (1.03 g, 4.85 mmol, 2.5 eq), XPhos (46 mg, 97 µmol, 0.05 eq) and XPhos Pd G2 (86 mg, 97 µmol, 0.05 eq) were dissolved in a mixture of 1,4-dioxane and water (15 mL, 4:1 ratio). The reaction mixture was stirred for 1 h at 85 °C after which time the solvent was removed *in vacuo*. The residue was taken up in ethyl acetate and washed with water and brine. The aqueous phase was extracted with ethyl acetate (3x). The combined organic phases were dried over MgSO4, filtered over cotton, and purified by flash column chromatography on silica gel using cyclohexane/ethyl acetate as an eluent to afford the product as a yellow solid (961 mg, 84%). $^1$H NMR (400 MHz, DMSO-$d_6$) δ 10.63 (s, 1H), 8.99 (d, J = 7.8 Hz, 1H), 8.75 (s, 1H), 7.80 (d, J = 8.3 Hz, 1H), 7.71 (d, J = 1.5 Hz, 1H), 7.69 (dd, J = 8.3, 1.7 Hz, 1H), 7.49 (d, J = 7.8 Hz, 1H), 4.24 (t, J = 6.2 Hz, 2H), 3.77 (t, J = 6.2 Hz, 2H), 3.61 (t, J = 5.0 Hz, 2H), 3.48 (t, J = 5.0 Hz, 2H), 1.53 (s, 9H), 0.76 (s, 9H), -0.08 (s, 6H) ppm. $^{13}$C NMR (101 MHz, DMSO-$d_6$) δ 169.48, 160.87, 153.80, 152.81, 143.80, 143.10, 139.85, 136.58, 130.08, 116.37, 112.74, 109.27, 106.26, 104.74, 82.51, 71.89, 68.22, 62.34, 52.34, 45.79, 27.65, 25.68, 17.86, -5.47 ppm. MS-ESI m/z $[M+H]^+$: calcd 587.28, found 587.10.

**Synthesis of Methyl 4-(5-{[(tert-butoxy)carbonyl][2-(2-hydroxyethoxy)ethyl]amino} pyrazolo[1,5-*a*]pyrimidine-3-yl)-2-hydroxybenzoate** (**13**). (**12**) (950 mg, 1.62 mmol, 1.0 eq) was dissolved in tetrahydrofuran (15 mL) and a 1 M solution of tetra-butylammonium fluoride in tetrahydrofuran (720 mg, 2.75 mmol) was added. The reaction mixture was stirred for 2 h at room temperature after which time the solvent was removed *in vacuo*. The reaction mixture was stirred for 16 h at 85 °C after which time the solvent was removed in vacuo. The residue was taken up in ethyl acetate and washed with water and brine. The aqueous phase was extracted with ethyl acetate (3x). The combined organic phases were dried over MgSO4, filtered over cotton, and purified by flash column chromatography on silica gel using cyclohexane/ethyl acetate as an eluent to afford the product as a light yellow solid (610 mg, 80%). $^{1}$H NMR (400 MHz, DMSO-$d_6$) δ 10.61 (s, 1H), 8.99 (d, J = 7.8 Hz, 1H), 8.76 (s, 1H), 7.81 (d, J = 8.3 Hz, 1H), 7.71 (d, J = 1.6 Hz, 1H), 7.69 (dd, J = 8.3, 1.6 Hz, 1H), 7.50 (d, J = 7.8 Hz, 1H), 4.53 (s, 1H), 4.22 (t, J = 6.3 Hz, 2H), 3.77 (t, J = 6.3 Hz, 2H), 3.51 – 3.40 (m, 4H), 1.53 (s, 9H) ppm. $^{13}$C NMR (101 MHz, DMSO-$d_6$) δ 169.44, 160.84, 153.85, 152.83, 143.80, 143.10, 139.80, 136.60, 130.17, 116.36, 112.70, 109.34, 106.20, 104.75, 82.54, 72.25, 67.99, 60.22, 52.35, 45.83, 27.65 ppm. MS-ESI m/z $[M+H]^+$: calcd 473.20, found 473.10.

**Synthesis of 9-(*tert*-butyl) 2$^4$-methyl (1$^3$*Z*,1$^4$*E*)-3,6-dioxa-9-aza-1(3,5)-pyrazolo[1,5-*a*] pyrimidina-2(1,3)-benzenacyclononaphane-2$^4$,9-dicarboxylate** (**14**). Triphenylphosphine (666 mg, 2.54 mmol, 6 eq) was solved in toluene (100 mL) and a solution of diisopropyl azodicarbonate (514 mg, 2.54 mmol, 6.0 eq) in toluene (25 mL) was added dropwise and the reaction mixture was stirred for 10 min at room temperature. Then a solution (**13**) (200 mg, 423 µmol, 1.0 eq) in toluene (15 mL) was added dropwise. The reaction mixture was stirred for 16 h at 90 °C after which time the solvent was removed *in vacuo*. The residue was taken up in ethyl acetate and washed with water and brine. The aqueous phase was extracted with ethyl acetate (3x). The combined organic phases were dried over MgSO4, filtered over cotton, and purified by flash column chromatography on silica gel using cyclohexane/ethyl acetate as an eluent to afford the product as a light yellow solid (167 mg, 87%). $^{1}$H NMR (500 MHz, DMSO-$d_6$) δ 9.01 (d, J = 7.8 Hz, 1H), 8.75 (d, J = 1.5 Hz, 1H), 8.71 (s, 1H), 7.69 (d, J = 8.1 Hz,

1H), 7.55 (d, J = 7.8 Hz, 1H), 7.43 (dd, J = 8.1, 1.5 Hz, 1H), 4.39 (t, J = 6.0 Hz, 2H), 4.09 (t, J = 6.6 Hz, 2H), 3.92 (t, J = 6.7 Hz, 2H), 3.88 (t, J = 6.0 Hz, 2H), 3.78 (s, 3H), 1.54 (s, 9H) ppm. $^{13}C$ NMR (126 MHz, DMSO-$d_6$) δ 165.79, 158.86, 153.41, 152.44, 143.08, 142.98, 137.60, 136.52, 131.24, 116.61, 112.64, 106.35, 104.16, 82.78, 67.51, 66.80, 66.57, 51.69, 46.57, 27.66 ppm. MS-ESI m/z $[M+H]^+$: calcd 455.19, found 455.15.

**Synthesis of Methyl ($1^3$*Z*,$1^4$*E*)-3,6-dioxa-9-aza-1(3,5)-pyrazolo[1,5-*a*]-pyrimidina-2(1,3)-benzenacyclononaphane-$2^4$-carboxylate** (**15**). (**14**) (82 mg, 180 µmol, 1.0 eq) was dissolved in dichloromethane (5 mL) and TFA (0.5 mL) was added. The reaction mixture was stirred for 16 h at room temperature after which time the solvent was removed *in vacuo*. The residue was taken up in ethyl acetate and washed with saturated sodium bicarbonate solution. The aqueous phase was extracted with ethyl acetate (3x). The combined organic phases were dried over MgSO4, filtered over cotton and the solvent was removed *in vacuo* to afford the product as a colourless solid (40 mg, 63%). $^1H$ NMR (400 MHz, DMSO-$d_6$) δ 8.83 (d, J = 1.5 Hz, 1H), 8.56 (d, J = 7.6 Hz, 1H), 8.39 (s, 1H), 7.93 (t, J = 5.3 Hz, 1H), 7.69 (d, J = 8.1 Hz, 1H), 7.28 (dd, J = 8.2, 1.3 Hz, 1H), 6.34 (d, J = 7.6 Hz, 1H), 4.40 – 4.33 (m, 2H), 4.03 – 3.95 (m, 2H), 3.90 – 3.83 (m, 2H), 3.76 (s, 3H), 3.58 – 3.48 (m, 2H) ppm. $^{13}C$ NMR (101 MHz, DMSO-$d_6$) δ 165.66, 158.09, 156.18, 145.33, 141.79, 139.04, 135.80, 131.60, 115.63, 114.82, 110.46, 103.40, 100.20, 65.36, 65.29, 63.88, 51.55 ppm. MS-ESI m/z $[M+H]^+$: calcd 355.13, found 355.10. HPLC: $t_R$ = 4.023 min (M1), purity ≧ 95% (UV: 254/280 nm).

**Synthesis of ($1^3$*Z*,$1^4$*E*)-3,6-dioxa-9-aza-1(3,5)-pyrazolo[1,5-*a*]pyrimidina-2(1,3)-benzena cyclononaphane-$2^4$-carboxylic acid** (**4**). (**15**) (60 mg, 169 µmol, 1.0 eq) was dissolved in a mixture of methanol and water (5 mL, 4:1 ratio) and lithium hydroxide monohydrate (35 mg, 847 mmol, 5.0 eq) was added. The reaction mixture was stirred for 16 h at 55 °C after which time the solvent was removed *in vacuo*. The residue was taken up in water and acidified with 4 M hydrochloric acid (pH=1). The resulting precipitate was filtered off, washed with water, and dried to afford the product as a colourless solid (55 mg, 87%). $^1H$ NMR (400 MHz, DMSO-$d_6$) δ 12.24 (bs, 1H), 8.81 (d, J = 1.3 Hz, 1H), 8.56 (d, J = 7.5 Hz, 1H), 8.38 (s, 1H), 8.03 (t, J = 5.3 Hz, 1H), 7.70 (d, J = 8.0 Hz, 1H), 7.27 (dd, J = 8.1, 1.3 Hz, 1H), 6.37 (d, J = 7.4 Hz, 1H), 4.43

– 4.33 (m, 2H), 3.99 (dd, J = 9.5, 5.9 Hz, 2H), 3.87 (dd, J = 9.1, 5.9 Hz, 2H), 3.56 – 3.49 (m, 2H) ppm. $^{13}C$ NMR (101 MHz, DMSO-$d_6$) δ 166.69, 158.07, 156.16, 145.29, 141.74, 138.71, 135.76, 131.81, 115.79, 115.61, 110.41, 103.48, 100.21, 65.34, 65.25, 63.92, 39.52 ppm. MS-ESI m/z $[M+H]^+$: calcd 341.12, found 341.10. HPLC: $t_R$ = 3.442 min (M1), purity ≧ 95% (UV: 254/280 nm).

**Synthesis of 1-((1$^3$*Z*,1$^4$*E*)-3,6-dioxa-9-aza-1(3,5)-pyrazolo[1,5-*a*]-pyrimidina-2(1,3) benzena cyclononaphane-2$^4$-yl)-3-benzylurea** (**16**). (**4**) (50 mg, 0.13 mmol, 1.0 eq) was dissolved in acetonitrile (10 mL) and triethylamine (27 mg, 0.26 mmol, 2.0 eq) and diphenylphosphoryl azide (47 mg, 0.17 mmol, 1.3 eq) were added. The mixture was stirred for 1.5 h at room temperature under argon atmosphere. Afterwards it was stirred under reflux for 1.5 h and benzylamine (21 mg, 0.20 mmol, 1.5 eq) was added. After another 1.5 h stirred under reflux, the solvent was removed *in vacuo*. The residue was taken up with ethyl acetate and was washed with water and brine. The organic phase was dried over $MgSO_4$ and purified by flash column chromatography on silica gel using water/acetonitrile as an eluent to afford the product as a white solid (23 mg, 35%). $^1H$ NMR (500 MHz, DMSO-$d_6$) δ 8.67 (d, J = 1.8 Hz, 1H), 8.51 (d, J = 7.5 Hz, 1H), 8.24 (s, 1H), 8.04 (d, J = 8.4 Hz, 1H), 7.95 (s, 1H), 7.76 (t, J = 5.3 Hz, 1H), 7.36 – 7.28 (m, 4H), 7.28 – 7.20 (m, 2H), 7.12 (dd, J = 8.4, 1.7 Hz, 1H), 6.28 (d, J = 7.6 Hz, 1H), 4.39 – 4.33 (m, 2H), 4.29 (d, J = 5.0 Hz, 2H), 4.02 – 3.97 (m, 2H), 3.91 – 3.85 (m, 2H), 3.55 – 3.48 (m, 2H) ppm. $^{13}C$ NMR (126 MHz, DMSO-$d_6$) δ 155.41, 155.19, 145.98, 144.19, 140.80, 140.29, 135.63, 128.36, 127.18, 126.77, 126.42, 126.13, 118.42, 116.04, 109.61, 104.38, 99.70, 65.12, 64.98, 63.75, 42.75 ppm. MS-ESI m/z $[M+H]^+$: calcd 445.19, found 445.20. HRMS *m/z* for $[C_{24}H_{24}N_6O_3+H]^+$: calcd 445.19827, found 445.19666. HPLC: $t_R$ = 3.840 min (M1), purity ≧ 95% (UV: 254/280 nm).

**Synthesis of 1-((1$^3$*Z*,1$^4$*E*)-3,6-dioxa-9-aza-1(3,5)-pyrazolo[1,5-*a*]-pyrimidina-2(1,3)benzena cyclononaphane-2$^4$-yl)-3-(2-chlorobenzyl)urea** (**17**). The title compound was prepared according to the procedure of (**16**), using (2-chlorophenyl) methanamine (28 mg, 0.20 mmol, 1.5 eq). The crude product was purified by flash column chromatography on silica gel using water/acetonitrile as an eluent to afford the product as a white solid (20 mg, 28%).

$^{1}H$ NMR (500 MHz, DMSO-$d_6$) δ 8.68 (d, J = 3.2 Hz, 1H), 8.51 (d, J = 7.5 Hz, 1H), 8.23 (s, 1H), 8.07 (s, 1H), 8.02 (d, J = 8.4 Hz, 1H), 7.76 (t, J = 5.0 Hz, 1H), 7.47 – 7.40 (m, 2H), 7.37 – 7.27 (m, 3H), 7.12 (d, J = 8.3 Hz, 1H), 6.27 (d, J = 7.5 Hz, 1H), 4.40 – 4.33 (m, 4H), 4.03 – 3.97 (m, 2H), 3.91 – 3.85 (m, 2H) ppm. $^{13}C$ NMR (126 MHz, DMSO-$d_6$) δ 155.42, 155.11, 146.06, 144.22, 140.81, 137.34, 135.62, 132.15, 129.16, 129.12, 128.67, 127.29, 126.56, 126.02, 118.47, 116.05, 109.64, 104.37, 99.71, 65.14, 65.01, 63.77, 40.73 ppm. MS-ESI m/z $[M+H]^+$: calcd 479.15, found 479.20. HRMS *m/z* for $[C_{24}H_{23}ClN_6O_3+H]^+$: calcd 479.15929, found 479.15793. HPLC: $t_R$ = 4.019 min (M1), purity ≧ 95% (UV: 254/280 nm).

**Synthesis of 1-((1$^3$*Z*,1$^4$*E*)-3,6-dioxa-9-aza-1(3,5)-pyrazolo[1,5-*a*]-pyrimidina-2(1,3)-benzena cyclononaphane-2$^4$-yl)-3-(3-chlorobenzyl)urea** (**18**). The title compound was prepared according to the procedure of (**16**), using (3-chlorophenyl) methanamine (28 mg, 0.20 mmol, 1.5 eq). The crude product was purified by flash column chromatography on silica gel using water/acetonitrile as an eluent to afford the product as a white solid (21 mg, 30%). $^{1}H$ NMR (500 MHz, DMSO-$d_6$) δ 8.68 (d, J = 1.8 Hz, 1H), 8.51 (d, J = 7.5 Hz, 1H), 8.24 (s, 1H), 8.02 (d, J = 8.3 Hz, 1H), 7.98 (s, 1H), 7.76 (t, J = 5.4 Hz, 1H), 7.39 – 7.34 (m, 2H), 7.34 – 7.24 (m, 3H), 7.12 (dd, J = 8.3, 1.7 Hz, 1H), 6.28 (d, J = 7.6 Hz, 1H), 4.37 (t, J = 7.7 Hz, 2H), 4.30 (d, J = 5.9 Hz, 2H), 4.03 – 3.96 (m, 2H), 3.91 – 3.85 (m, 2H), 3.55 – 3.48 (m, 2H) ppm. $^{13}C$ NMR (126 MHz, DMSO-$d_6$) δ 155.43, 155.21, 146.11, 144.22, 143.13, 140.82, 135.63, 133.02, 130.27, 126.86, 126.68, 126.62, 125.97, 125.81, 118.56, 116.06, 109.64, 104.36, 99.72, 65.13, 65.00, 63.75, 42.19 ppm. MS-ESI m/z $[M+H]^+$: calcd 479.15, found 479.15. HRMS *m/z* for $[C_{24}H_{23}ClN_6O_3+H]^+$: calcd 478.15147, found 478.14963. HPLC: $t_R$ = 3.990 min (M1), purity ≧ 95% (UV: 254/280 nm).

**Synthesis of 1-((1$^3$*Z*,1$^4$*E*)-3,6-dioxa-9-aza-1(3,5)-pyrazolo[1,5-*a*]pyrimidina-2(1,3)-benzena cyclononaphane-2$^4$-yl)-3-(4-chlorobenzyl)urea** (**19**). The title compound was prepared according to the procedure of (**16**), using (4-chlorophenyl) methanamine (28 mg, 0.20 mmol, 1.5 eq). The crude product was purified by flash column chromatography on silica gel using water/acetonitrile as an eluent to afford the product as a white solid (13 mg, 19%). $^{1}H$ NMR (600 MHz, DMSO-$d_6$) δ 8.68 (s, 1H), 8.51 (d, J = 7.5 Hz, 1H), 8.24 (s, 1H), 8.03 (d, J

= 8.3 Hz, 1H), 7.95 (s, 1H), 7.75 (t, J = 5.7 Hz, 1H), 7.42 – 7.30 (m, 4H), 7.26 (t, J = 6.1 Hz, 1H), 7.12 (d, J = 8.3 Hz, 1H), 6.28 (d, J = 7.6 Hz, 1H), 4.40 – 4.33 (m, 2H), 4.28 (d, J = 6.0 Hz, 2H), 4.02 – 3.84 (m, 4H), 3.56 – 3.48 (m, 2H) ppm. $^{13}$C NMR (151 MHz, DMSO-d$_6$) δ 155.37, 155.16, 146.06, 144.18, 140.74, 139.43, 135.54, 131.20, 128.94, 128.21, 126.53, 125.99, 118.50, 115.98, 109.67, 104.35, 99.65, 65.23, 65.07, 63.88, 42.03 ppm. MS-ESI m/z [M+H]$^+$: calcd 479.15, found 479.20. HRMS *m/z* for [$C_{24}H_{23}ClN_6O_3$+H]$^+$: calcd 479.15929, found 479.15805. HPLC: $t_R$ = 4.024 min (M1), purity ≧ 95% (UV: 254/280 nm).

**Synthesis of 1-((1$^3$*Z*,1$^4$*E*)-3,6-dioxa-9-aza-1(3,5)-pyrazolo[1,5-*a*]pyrimidina-2(1,3)-benzena cyclononaphane-2$^4$-yl)-3-(2,4-dichlorobenzyl)urea** (**20**). The title compound was prepared according to the procedure of (**16**), using 2,4-dichlorobenzylamine (35 mg, 0.20 mmol, 1.5 eq). The crude product was purified by flash column chromatography on silica gel using water/acetonitrile as an eluent to afford the product as a white solid (20 mg, 27%). $^1$H NMR (600 MHz, DMSO-d$_6$) δ 8.69 (d, J = 1.9 Hz, 1H), 8.51 (d, J = 7.7 Hz, 1H), 8.23 (s, 1H), 8.07 (s, 1H), 8.01 (d, J = 8.2 Hz, 1H), 7.76 (t, J = 5.5 Hz, 1H), 7.61 (d, J = 2.1 Hz, 1H), 7.47 – 7.39 (m, 2H), 7.32 (t, J = 6.0 Hz, 1H), 7.11 (d, J = 8.3 Hz, 1H), 6.28 (d, J = 7.5 Hz, 1H), 4.37 (t, J = 7.7 Hz, 2H), 4.34 (d, J = 5.8 Hz, 2H), 4.03 – 3.96 (m, 2H), 3.91 – 3.85 (m, 2H), 3.55 – 3.49 (m, 2H) ppm. $^{13}$C NMR (151 MHz, DMSO-d$_6$) δ 155.39, 155.05, 146.12, 144.18, 140.74, 136.60, 135.55, 132.95, 132.13, 130.34, 128.52, 127.35, 126.64, 125.88, 118.53, 115.99, 109.69, 104.31, 99.67, 65.26, 65.10, 63.90, 40.27 ppm. MS-ESI m/z [M+H]$^+$: calcd 513.11, found 513.20. HRMS *m/z* for [$C_{24}H_{22}Cl_2N_6O_3$+H]$^+$: calcd 514.12368, found 514.10914. HPLC: $t_R$ = 4.292 min (M1), purity ≧ 95% (UV: 254/280 nm).

**Synthesis of 1-((1$^3$*Z*,1$^4$*E*)-3,6-dioxa-9-aza-1(3,5)-pyrazolo[1,5-*a*]pyrimidina-2(1,3)-benzena cyclononaphane-2$^4$-yl)-3-(4-chloro-2-fluorobenzyl)urea** (**21**). The title compound was prepared according to the procedure of (**16**), using (4-chloro-2-fluorophenyl)methanamine (32 mg, 0.20 mmol, 1.5 eq). The crude product was purified by flash column chromatography on silica gel using water/acetonitrile as an eluent to afford the product as a white solid (23 mg, 32%). $^1$H NMR (600 MHz, DMSO-d$_6$) δ 8.68 (d, J = 1.8 Hz, 1H), 8.50 (d, J = 7.6 Hz, 1H), 8.23 (s, 1H), 8.02 – 7.98 (m, 2H), 7.76 (t, J = 5.4 Hz, 1H), 7.43 – 7.38 (m, 2H), 7.31 – 7.26 (m, 2H),

7.11 (dd, J = 8.4, 1.8 Hz, 1H), 6.28 (d, J = 7.6 Hz, 1H), 4.39 – 4.33 (m, 2H), 4.31 (d, J = 5.8 Hz, 2H), 4.02 – 3.97 (m, 2H), 3.91 – 3.85 (m, 2H), 3.55 – 3.48 (m, 2H) ppm. $^{13}$C NMR (151 MHz, DMSO-$d_6$) δ 159.92 (d, J = 248.0 Hz), 155.38, 155.04, 146.08, 144.17, 140.72, 135.54, 132.14 (d, J = 10.4 Hz), 130.77 (d, J = 5.3 Hz), 126.60, 126.29 (d, J = 14.9 Hz), 125.89, 124.56 (d, J = 3.7 Hz), 118.50, 115.98, 115.77, 115.60, 109.67, 104.31, 99.66, 65.24, 65.07, 63.88, 36.29 ppm. MS-ESI m/z $[M+H]^+$: calcd 497.14, found 497.15. HRMS *m/z* for $[C_{24}H_{22}ClFN_6O_3+H]^+$: calcd 497.14816, found 497.14987. HPLC: $t_R$ = 4.077 min (M1), purity ≧ 95% (UV: 254/280 nm).

**Synthesis of 1-((1$^3$*Z*,1$^4$*E*)-3,6-dioxa-9-aza-1(3,5)-pyrazolo[1,5-*a*]pyrimidina-2(1,3)-benzena cyclononaphane-2$^4$-yl)-3-(4-fluoro-2-methylbenzyl)urea** (**22**). The title compound was prepared according to the procedure of (**16**), using (4-fluoro-2-methylphenyl)methanamine (28 mg, 0.20 mmol, 1.5 eq). The crude product was purified by flash column chromatography on silica gel using water/acetonitrile as an eluent to afford the product as a white solid (13 mg, 21%). $^1$H NMR (400 MHz, DMSO-$d_6$) δ 8.67 (d, J = 1.8 Hz, 1H), 8.51 (d, J = 7.6 Hz, 1H), 8.23 (s, 1H), 8.04 (d, J = 8.3 Hz, 1H), 7.93 (s, 1H), 7.78 – 7.72 (m, 1H), 7.31 – 7.24 (m, 1H), 7.14 – 7.09 (m, 2H), 7.07 – 6.96 (m, 2H), 6.27 (d, J = 7.6 Hz, 1H), 4.39 – 4.32 (m, 2H), 4.24 (d, J = 5.6 Hz, 2H), 4.03 – 3.95 (m, 2H), 3.92 – 3.84 (m, 2H), 3.51 (q, J = 6.8 Hz, 2H), 2.32 (s, 3H) ppm. $^{13}$C NMR (126 MHz, DMSO-$d_6$) δ 161.08 (d, J = 241.9 Hz), 155.40, 155.01, 145.95, 144.18, 140.78, 138.43 (d, J = 7.9 Hz), 135.61, 134.10 (d, J = 2.9 Hz), 129.50 (d, J = 8.4 Hz), 126.40, 126.13, 118.32, 116.57, 116.36, 116.04, 112.27, 112.07, 109.63, 104.37, 99.69, 65.15, 65.01, 63.79, 18.56 ppm. MS-ESI m/z $[M+H]^+$: calcd 477.20, found 477.20. HRMS *m/z* for $[C_{25}H_{25}FN_6O_3+H]^+$: calcd 477.2045, found 477.2040. HPLC: $t_R$ = 3.957 min (M1), purity ≧ 95% (UV: 254/280 nm).

**Synthesis of 1-((1$^3$*Z*,1$^4$*E*)-3,6-dioxa-9-aza-1(3,5)-pyrazolo[1,5-*a*]pyrimidina-2(1,3)-benzena cyclononaphane-2$^4$-yl)-3-(4-chloro-2-methylbenzyl)urea** (**23**). The title compound was prepared according to the procedure of (**16**), using (4-chloro-2-methylphenyl)methanamine (31 mg, 0.20 mmol, 1.5 eq). The crude product was purified by flash column chromatography on silica gel using water/acetonitrile as an eluent to afford the product as a white solid (20 mg, 28%). $^{1}$H NMR (500 MHz, DMSO-$d_6$) δ 8.67 (d, J = 1.9 Hz, 1H), 8.51 (d, J = 7.6 Hz, 1H), 8.23 (s, 1H), 8.03 (d, J = 8.3 Hz, 1H), 7.96 (s, 1H), 7.76 (t, J = 5.3 Hz, 1H), 7.29 – 7.22 (m, 3H), 7.15 (t, J = 5.7 Hz, 1H), 7.12 (dd, J = 8.4, 1.7 Hz, 1H), 6.27 (d, J = 7.6 Hz, 1H), 4.39 – 4.33 (m, 2H), 4.25 (d, J = 5.6 Hz, 2H), 4.03 – 3.95 (m, 2H), 3.90 – 3.84 (m, 2H), 3.55 – 3.47 (m, 2H), 2.30 (s, 3H) ppm. $^{13}$C NMR (126 MHz, DMSO-$d_6$) δ 155.41, 155.04, 145.96, 144.20, 140.81, 138.13, 137.10, 135.63, 131.11, 129.47, 129.18, 126.45, 126.09, 125.62, 118.33, 116.05, 109.61, 104.37, 99.70, 65.12, 65.00, 63.74, 18.33 ppm. MS-ESI m/z $[M+H]^+$: calcd 493.17, found 493.15. HRMS *m/z* for $[C_{25}H_{25}ClN_6O_3+H]^+$: calcd 493.17494, found 493.17324. HPLC: $t_R$ = 4.927 min (M2), purity ≧ 95% (UV: 254/280 nm).

**Synthesis of 1-((1$^3$*Z*,1$^4$*E*)-3,6-dioxa-9-aza-1(3,5)-pyrazolo[1,5-*a*]pyrimidina-2(1,3)-benzena cyclononaphane-2$^4$-yl)-3-(4-bromo-2-methylbenzyl)urea** (**24**). The title compound was prepared according to the procedure of (**16**), using (4-bromo-2-methylphenyl)methanamine (13 mg, 66 umol, 1.5 eq). The crude product was purified by preparative HPLC to afford the product as a white solid (20 mg, 84%). $^{1}$H NMR (400 MHz, DMSO-$d_6$) δ 8.68 (s, 1H), 8.51 (d, J = 7.7 Hz, 1H), 8.23 (s, 1H), 8.03 (d, J = 8.2 Hz, 1H), 7.96 (s, 1H), 7.76 (s, 1H), 7.38 (d, J = 12.6 Hz, 2H), 7.27 – 7.07 (m, 3H), 6.28 (d, J = 7.7 Hz, 1H), 4.44 – 4.31 (m, 2H), 4.23 (s, 2H), 4.05 – 3.82 (m, 4H), 3.59 – 3.46 (m, 2H), 2.30 (s, 3H) ppm. $^{13}$C NMR (101 MHz, DMSO-$d_6$) δ 158.55, 158.18, 155.44, 155.07, 146.01, 144.22, 140.77, 138.49, 137.57, 135.62, 132.32, 129.51, 128.58, 126.47, 126.11, 119.67, 118.38, 116.79, 116.05, 113.90, 109.66, 104.39, 99.73, 65.19, 65.04, 63.82, 40.31, 18.25 ppm. MS-ESI m/z $[M+H]^+$: calcd 539.12, found 539.10. HRMS *m/z* for $[C_{25}H_{25}BrN_6O_3+H]^+$: calcd 537.1244, found 537.1227. HPLC: $t_R$ = 10.283 min (M3), purity = 94% (UV: 254/280 nm).

**Synthesis of 1-((1$^3$*Z*,1$^4$*E*)-3,6-dioxa-9-aza-1(3,5)-pyrazolo[1,5-*a*]pyrimidina-2(1,3)-benzena cyclononaphane-2$^4$-yl)-3-(2-methyl-4-(trifluoromethyl)benzyl)urea** (**25**). The title compound was prepared according to the procedure of (**16**), using (2-methyl-4-(trifluoromethyl)phenyl)methanamine (38 mg, 0.20 mmol, 1.5 eq). The crude product was purified by flash column chromatography on silica gel using water/acetonitrile as an eluent to afford the product as a white solid (10 mg, 14%). $^{1}$H NMR (400 MHz, DMSO-$d_6$) δ 8.68 (s, 1H),

8.51 (d, J = 7.4 Hz, 1H), 8.23 (s, 1H), 8.06 – 7.99 (m, 2H), 7.76 (t, J = 5.0 Hz, 1H), 7.55 (d, J = 7.0 Hz, 2H), 7.46 (d, J = 8.0 Hz, 1H), 7.26 (t, J = 5.7 Hz, 1H), 7.12 (d, J = 8.4 Hz, 1H), 6.28 (d, J = 7.5 Hz, 1H), 4.41 – 4.31 (m, 4H), 4.04 – 3.97 (m, 2H), 3.88 (t, J = 7.4 Hz, 2H), 3.56 – 3.47 (m, 2H), 2.39 (s, 3H) ppm. $^{13}$C NMR (101 MHz, DMSO-$d_6$) δ 155.41, 155.11, 146.01, 144.20, 143.01, 140.79, 136.91, 135.61, 127.65, 127.36 (q, J = 31.4 Hz), 126.53, 126.30 (q, J = 3.2 Hz), 126.03, 124.43 (q, J = 272.0 Hz), 122.60 (q, J = 3.9 Hz), 118.38, 116.04, 109.64, 104.35, 99.70, 65.16, 65.03, 63.79, 40.51, 18.40 ppm. MS-ESI m/z [M+H]$^+$: calcd 527.19, found 527.20. HRMS *m/z* for [$C_{26}H_{25}F_3N_6O_3$+H]$^+$: calcd 527.2013, found 527.2015. HPLC: $t_R$ = 5.001 min (M2), purity ≧ 95% (UV: 254/280 nm).

**Synthesis of 1-((1$^3$*Z*,1$^4$*E*)-3,6-dioxa-9-aza-1(3,5)-pyrazolo[1,5-*a*]pyrimidina-2(1,3)-benzena cyclononaphane-2$^4$-yl)-3-(5-fluoro-2-methylbenzyl)urea** (**26**). The title compound was prepared according to the procedure of (**16**), using (5-fluoro-2-methylphenyl)methanamine (18 mg, 0.13 mmol, 1.5 eq). The crude product was purified by preparative HPLC to afford the product as a white solid (5 mg, 11%). $^1$H NMR (400 MHz, DMSO-$d_6$) δ 8.68 (d, J = 1.8 Hz, 1H), 8.51 (d, J = 7.6 Hz, 1H), 8.23 (s, 1H), 8.03 (d, J = 8.3 Hz, 1H), 7.99 (s, 1H), 7.77 (t, J = 5.3 Hz, 1H), 7.23 – 7.16 (m, 2H), 7.12 (dd, J = 8.4, 1.7 Hz, 1H), 7.04 (dd, J = 10.1, 2.8 Hz, 1H), 6.98 (td, J = 8.5, 2.8 Hz, 1H), 6.28 (d, J = 7.6 Hz, 1H), 4.40 – 4.35 (m, 2H), 4.26 (d, J = 5.6 Hz, 2H), 4.02 – 3.98 (m, 2H), 3.90 – 3.86 (m, 2H), 3.54 – 3.49 (m, 2H), 2.26 (s, 3H) ppm. $^{13}$C NMR (101 MHz, DMSO-$d_6$) δ 155.41, 155.09, 146.06, 144.20, 140.78, 135.61, 131.43, 131.35, 126.55, 126.03, 118.46, 116.04, 113.75, 113.14, 112.94, 104.35, 99.70, 65.16, 65.00, 63.78, 50.98, 17.76 ppm. MS-ESI m/z [M+H]$^+$: calcd 477.20, found 477.20. HRMS *m/z* for [$C_{25}H_{25}FN_6O_3$+H]$^+$: calcd 477.2045, found 477.2029. HPLC: $t_R$ = 3.959 min (M1), purity ≧ 95% (UV: 254/280 nm).

**Synthesis of 1-((1$^3$*Z*,1$^4$*E*)-3,6-dioxa-9-aza-1(3,5)-pyrazolo[1,5-*a*]pyrimidina-2(1,3)-benzena cyclononaphane-2$^4$-yl)-3-(2-(trifluoromethyl)benzyl)urea** (**27**). The title compound was prepared according to the procedure of (**16**), using (2-(trifluoromethyl)phenyl)methanamine (23 mg, 0.13 mmol, 1.5 eq). The crude product was purified by preparative HPLC to afford the product as a white solid (14 mg, 31%). $^1$H NMR (400

MHz, DMSO-$d_6$) δ 8.69 (d, J = 1.8 Hz, 1H), 8.51 (d, J = 7.6 Hz, 1H), 8.24 (s, 1H), 8.09 (s, 1H), 8.03 (d, J = 8.4 Hz, 1H), 7.77 (t, J = 5.0 Hz, 1H), 7.73 – 7.66 (m, 2H), 7.61 (d, J = 7.7 Hz, 1H), 7.48 (t, J = 7.7 Hz, 1H), 7.32 (t, J = 6.0 Hz, 1H), 7.12 (dd, J = 8.4, 1.7 Hz, 1H), 6.28 (d, J = 7.6 Hz, 1H), 4.48 (d, J = 5.5 Hz, 2H), 4.41 – 4.34 (m, 2H), 4.03 – 3.97 (m, 2H), 3.92 – 3.85 (m, 2H), 3.55 – 3.48 (m, 2H) ppm. $^{13}$C NMR (101 MHz, DMSO-$d_6$) δ 158.45, 158.09, 155.43, 155.11, 146.10, 144.22, 140.80, 138.71, 135.62, 132.79, 129.12, 127.37, 126.63, 125.97, 125.73, 125.68, 123.18, 118.51, 116.04, 109.66, 104.35, 99.72, 65.17, 65.04, 63.80 ppm. MS-ESI m/z [M+H]$^+$: calcd 513.18, found 513.10. HRMS *m/z* for [$C_{25}H_{23}F_3N_6O_3$+H]$^+$: calcd 513.1857, found 513.1858. HPLC: $t_R$ = 4.224 min (M1), purity ≧ 95% (UV: 254/280 nm).

**Synthesis of 1-((1$^3$*Z*,1$^4$*E*)-3,6-dioxa-9-aza-1(3,5)-pyrazolo[1,5-*a*]pyrimidina-2(1,3)-benzena cyclononaphane-2$^4$-yl)-3-(4-(trifluoromethyl)benzyl)urea** (**28**). The title compound was prepared according to the procedure of (**16**), using (4-(trifluoromethyl)phenyl)methanamine (23 mg, 0.13 mmol, 1.5 eq). The crude product was purified by preparative HPLC to afford the product as a white solid (18 mg, 40%). $^1$H NMR (400 MHz, DMSO-$d_6$) δ 8.69 (d, J = 1.8 Hz, 1H), 8.51 (d, J = 7.5 Hz, 1H), 8.23 (s, 1H), 8.06 – 7.98 (m, 2H), 7.77 (t, J = 5.5 Hz, 1H), 7.70 (d, J = 8.0 Hz, 2H), 7.52 (d, J = 8.1 Hz, 2H), 7.36 (t, J = 5.8 Hz, 1H), 7.12 (dd, J = 8.4, 1.7 Hz, 1H), 6.28 (d, J = 7.5 Hz, 1H), 4.43 – 4.33 (m, 4H), 4.04 – 3.98 (m, 2H), 3.92 – 3.86 (m, 2H), 3.56 – 3.48 (m, 3H) ppm. $^{13}$C NMR (101 MHz, DMSO-$d_6$) δ 158.43, 158.07, 155.43, 155.25, 146.10, 145.45, 144.21, 140.78, 135.61, 127.70, 126.61, 125.97, 125.20, 123.05, 118.53, 116.03, 109.65, 104.35, 99.71, 65.17, 65.03, 63.79, 42.33, 34.27 ppm. MS-ESI m/z [M+H]$^+$: calcd 513.18, found 513.15. HRMS *m/z* for [$C_{25}H_{23}F_3N_6O_3$+H]$^+$: calcd 513.1857, found 513.1854. HPLC: $t_R$ = 9.748 min (M3), purity = 91% (UV: 254/280 nm).

**Synthesis of 1-((1$^3$*Z*,1$^4$*E*)-3,6-dioxa-9-aza-1(3,5)-pyrazolo[1,5-*a*]pyrimidina-2(1,3)-benzena cyclononaphane-2$^4$-yl)-3-(2-methoxybenzyl)urea** (**29**). The title compound was prepared according to the procedure of (**16**), using (2-methoxyphenyl)methanamine (18 mg, 0.13 mmol, 1.5 eq). The crude product was purified by preparative HPLC to afford the product as a white solid (10 mg, 25%). $^1$H NMR (400 MHz, DMSO-d$_6$) δ 8.67 (d, J = 1.8 Hz, 1H), 8.51 (d, J = 7.5 Hz, 1H), 8.23 (s, 1H), 8.02 (d, J = 8.4 Hz, 1H), 8.00 (s, 1H), 7.76 (t, J = 5.2 Hz, 1H), 7.24 (d, J = 7.5 Hz, 2H), 7.14 – 7.06 (m, 2H), 7.01 – 6.98 (m, 1H), 6.92 (t, J = 7.3 Hz, 1H), 6.28 (d, J = 7.6 Hz, 1H), 4.39 – 4.32 (m, 2H), 4.24 (d, J = 5.6 Hz, 2H), 4.02 – 3.96 (m, 2H), 3.90 – 3.85 (m, 2H), 3.82 (s, 3H), 3.54 – 3.48 (m, 2H) ppm. $^{13}$C NMR (101 MHz, DMSO-d$_6$) δ 156.81, 155.40, 155.17, 146.01, 144.18, 140.78, 135.61, 130.46, 128.14, 128.09, 127.66, 126.34, 126.22, 120.16, 118.46, 116.02, 114.02, 110.49, 109.62, 104.39, 99.69, 65.15, 64.99, 63.79, 55.33, 38.02 ppm. MS-ESI m/z [M+H]$^+$: calcd 475.20, found 475.20. HRMS *m/z* for [$C_{25}H_{26}N_6O_4$+H]$^+$: calcd 475.2088, found 475.2087. HPLC: $t_R$ = 3.853 min (M1), purity ≧ 95% (UV: 254/280 nm).

**Synthesis of 1-((1$^3$*Z*,1$^4$*E*)-3,6-dioxa-9-aza-1(3,5)-pyrazolo[1,5-*a*]pyrimidina-2(1,3)-benzena cyclononaphane-2$^4$-yl)-3-(3-methoxybenzyl)urea** (**30**). The title compound was prepared according to the procedure of (**16**), using (3-methoxyphenyl)methanamine (18 mg, 0.13 mmol, 1.5 eq). The crude product was purified by preparative HPLC to afford the product as a white solid (5 mg, 13%). $^1$H NMR (400 MHz, DMSO-d$_6$) δ 8.68 (d, J = 1.7 Hz, 1H), 8.51 (d, J = 7.6 Hz, 1H), 8.23 (s, 1H), 8.03 (d, J = 8.3 Hz, 1H), 7.94 (s, 1H), 7.76 (t, J = 5.2 Hz, 1H), 7.25 (t, J = 8.0 Hz, 1H), 7.21 (t, J = 5.8 Hz, 1H), 7.12 (dd, J = 8.4, 1.7 Hz, 1H), 6.90 – 6.85 (m, 2H), 6.84 – 6.79 (m, 1H), 6.28 (d, J = 7.6 Hz, 1H), 4.39 – 4.33 (m, 2H), 4.26 (d, J = 5.8 Hz, 2H), 4.02 – 3.96 (m, 2H), 3.91 – 3.84 (m, 2H), 3.74 (s, 3H), 3.56 – 3.48 (m, 2H) ppm. $^{13}$C NMR (101 MHz, DMSO-d$_6$) δ 159.35, 155.40, 155.17, 146.02, 141.90, 140.80, 135.61, 129.43, 126.45, 126.11, 119.31, 118.46, 116.04, 114.01, 112.85, 112.06, 109.64, 104.37, 99.69, 65.16, 65.01, 63.76, 55.00, 42.71 ppm. MS-ESI m/z [M+H]$^+$: calcd 475.20, found 475.20. HRMS *m/z* for [$C_{25}H_{26}N_6O_4$+H]$^+$: calcd 475.2088, found 475.2083. HPLC: $t_R$ = 3.788 min (M1), purity ≧ 95% (UV: 254/280 nm).

**Synthesis of 1-((1$^3$*Z*,1$^4$*E*)-3,6-dioxa-9-aza-1(3,5)-pyrazolo[1,5-*a*]pyrimidina-2(1,3)-benzena cyclononaphane-2$^4$-yl)-3-(4-methoxybenzyl)urea** (**31**). The title compound was prepared according to the procedure of (**16**), using (4-methoxyphenyl)methanamine (18 mg,

0.13 mmol, 1.5 eq). The crude product was purified by preparative HPLC to afford the product as a white solid (6 mg, 14%). $^1$H NMR (400 MHz, DMSO-$d_6$) δ 8.67 (s, 1H), 8.51 (d, J = 7.6 Hz, 1H), 8.23 (s, 1H), 8.04 (d, J = 8.4 Hz, 1H), 7.90 (s, 1H), 7.76 (t, J = 5.3 Hz, 1H), 7.22 (d, J = 8.5 Hz, 2H), 7.18 – 7.09 (m, 2H), 6.90 (d, J = 8.5 Hz, 2H), 6.28 (d, J = 7.6 Hz, 1H), 4.39 – 4.31 (m, 2H), 4.21 (d, J = 5.5 Hz, 2H), 4.04 – 3.94 (m, 2H), 3.92 – 3.83 (m, 2H), 3.73 (s, 3H), 3.58 – 3.47 (m, 2H) ppm. $^{13}$C NMR (101 MHz, DMSO-$d_6$) δ 158.21, 155.39, 155.13, 145.97, 144.18, 140.77, 135.60, 132.16, 128.53, 126.37, 126.16, 118.40, 116.02, 113.74, 109.62, 104.38, 99.69, 65.15, 64.99, 63.78, 55.06, 42.22, 40.43 ppm. MS-ESI m/z [M+H]$^+$: calcd 475.20, found 475.20. HRMS *m/z* for [$C_{25}H_{26}N_6O_4$+H]$^+$: calcd 475.2088, found 475.2088. HPLC: $t_R$ = 3.763 min (M1), purity ≧ 95% (UV: 254/280 nm).

**Synthesis of 1-((1$^3$*Z*,1$^4$*E*)-3,6-dioxa-9-aza-1(3,5)-pyrazolo[1,5-*a*]pyrimidina-2(1,3)-benzena cyclononaphane-2$^4$-yl)-3-(3-fluoro-4-methoxybenzyl)urea** (**32**). The title compound was prepared according to the procedure of (**16**), using (3-fluoro-4-methoxyphenyl)methanamine (21 mg, 0.13 mmol, 1.5 eq). The crude product was purified by preparative HPLC to afford the product as a white solid (11 mg, 26%). $^1$H NMR (400 MHz, DMSO-$d_6$) δ 8.68 (s, 1H), 8.51 (d, J = 7.6 Hz, 1H), 8.23 (s, 1H), 8.02 (d, J = 8.3 Hz, 1H), 7.93 (s, 1H), 7.78 (t, J = 4.9 Hz, 1H), 7.21 (t, J = 5.9 Hz, 1H), 7.16 – 7.05 (m, 4H), 6.28 (d, J = 7.6 Hz, 1H), 4.41 – 4.30 (m, 2H), 4.22 (d, J = 5.7 Hz, 2H), 4.04 – 3.94 (m, 2H), 3.93 – 3.84 (m, 2H), 3.81 (s, 3H), 3.56 – 3.49 (m, 2H) ppm. $^{13}$C NMR (101 MHz, DMSO-$d_6$) δ 155.41, 155.16, 151.33 (d, J = 243.7 Hz), 146.06, 144.20, 140.78, 135.60, 133.38 (d, J = 5.6 Hz), 126.52, 126.04, 123.29 (d, J = 3.3 Hz), 118.51, 116.03, 114.84, 114.66, 113.78 (d, J = 1.8 Hz), 109.64, 104.36, 99.70, 65.15, 65.01, 63.78, 56.03, 41.86 ppm. MS-ESI m/z [M+H]$^+$: calcd 493.19, found 493.20. HRMS *m/z* for [$C_{25}H_{25}FN_6O_4$+H]$^+$: calcd 493.1994, found 493.1987. HPLC: $t_R$ = 3.782 min (M1), purity ≧ 95% (UV: 254/280 nm).

**Synthesis of 1-((1$^3$*Z*,1$^4$*E*)-3,6-dioxa-9-aza-1(3,5)-pyrazolo[1,5-*a*]pyrimidina-2(1,3)-benzena cyclononaphane-2$^4$-yl)-3-(4-fluoro-3-methoxybenzyl)urea** (**33**). The title compound was prepared according to the procedure of (**16**), using (4-fluoro-3-

methoxyphenyl)methanamine (21 mg, 0.13 mmol, 1.5 eq). The crude product was purified by preparative HPLC to afford the product as a white solid (7 mg, 16%). $^1$H NMR (400 MHz, DMSO-$d_6$) δ 8.68 (d, J = 1.5 Hz, 1H), 8.51 (d, J = 7.6 Hz, 1H), 8.23 (s, 1H), 8.03 (d, J = 8.3 Hz, 1H), 7.94 (s, 1H), 7.78 (t, J = 5.5 Hz, 1H), 7.23 (t, J = 6.0 Hz, 1H), 7.19 – 7.08 (m, 3H), 6.90 – 6.83 (m, 1H), 6.28 (d, J = 7.6 Hz, 1H), 4.41 – 4.32 (m, 2H), 4.26 (d, J = 5.7 Hz, 2H), 4.03 – 3.95 (m, 2H), 3.91 – 3.86 (m, 2H), 3.83 (s, 3H), 3.13 – 3.04 (m, 2H) ppm. $^{13}$C NMR (126 MHz, DMSO-$d_6$) δ 155.41, 155.15, 146.04, 144.20, 140.77, 135.60, 126.49, 126.06, 119.25, 119.18, 118.49, 116.03, 115.66, 115.48, 112.87, 109.65, 104.36, 99.70, 65.16, 65.01, 63.78, 55.92, 45.62, 42.41 ppm. MS-ESI m/z [M+H]$^+$: calcd 493.19, found 493.15. HRMS *m/z* for [$C_{25}H_{25}FN_6O_4$+H]$^+$: calcd 493.1994, found 493.1985. HPLC: $t_R$ = 3.923 min (M1), purity ≧ 95% (UV: 254/280 nm).

**Synthesis of 1-((1$^3$*Z*,1$^4$*E*)-3,6-dioxa-9-aza-1(3,5)-pyrazolo[1,5-*a*]pyrimidina-2(1,3)-benzena cyclononaphane-2$^4$-yl)-3-((*S*)-1-phenylethyl)urea** (**34**). The title compound was prepared according to the procedure of (**16**), using (*S*)-1-phenylethan-1-amine (8 mg, 66 μmol, 1.5 eq). The crude product was purified by preparative HPLC to afford the product as a white solid (14 mg, 72%). $^1$H NMR (400 MHz, DMSO-$d_6$) δ 8.67 (d, J = 1.6 Hz, 1H), 8.50 (d, J = 7.6 Hz, 1H), 8.22 (s, 1H), 8.01 (d, J = 8.4 Hz, 1H), 7.88 (s, 1H), 7.75 (s, 1H), 7.36 – 7.31 (m, 4H), 7.31 – 7.26 (m, 1H), 7.25 – 7.20 (m, 1H), 7.09 (dd, J = 8.4, 1.6 Hz, 1H), 6.27 (d, J = 7.6 Hz, 1H), 4.81 (p, J = 7.2, 6.4 Hz, 1H), 4.43 – 4.32 (m, 2H), 4.06 – 3.81 (m, 4H), 3.57 – 3.46 (m, 2H), 1.37 (d, J = 6.9 Hz, 3H) ppm. $^{13}$C NMR (101 MHz, DMSO-$d_6$) δ 158.55, 158.18, 155.42, 154.33, 145.83, 145.42, 144.18, 140.75, 135.61, 128.33, 126.62, 126.21, 125.84, 118.09, 116.64, 116.03, 113.76, 109.60, 104.41, 99.71, 65.17, 65.03, 63.81, 48.61, 23.30 ppm. MS-ESI m/z [M+H]$^+$: calcd 459.20, found 459.15. HRMS *m/z* for [$C_{25}H_{26}N_6O_3$+H]$^+$: calcd 459.2139, found 459.2138. HPLC: $t_R$ = 4.059 min (M1), purity ≧ 95% (UV: 254/280 nm).

**Synthesis of *N*-((1$^3$*Z*,1$^4$*E*)-3,6-dioxa-9-aza-1(3,5)-pyrazolo[1,5-*a*]pyrimidina-2(1,3)-benzena cyclononaphane-2$^4$-yl)-4-methylpiperazine-1-carboxamide** (**35**). The title compound was prepared according to the procedure of (**16**), using 1-methylpiperazine (20 mg, 0.20 mmol, 1.5 eq). The crude product was purified by flash column chromatography on silica gel using water/acetonitrile as an eluent to afford the product as a white solid (24 mg, 37%). $^{1}H$ NMR (500 MHz, DMSO-$d_6$) δ 8.69 (d, J = 1.8 Hz, 1H), 8.52 (d, J = 7.5 Hz, 1H), 8.26 (s, 1H), 7.78 (t, J = 5.4 Hz, 1H), 7.57 (s, 1H), 7.56 (d, J = 8.2 Hz, 1H), 7.14 (dd, J = 8.2, 1.7 Hz, 1H), 6.28 (d, J = 7.6 Hz, 1H), 4.36 – 4.31 (m, 2H), 4.01 – 3.96 (m, 2H), 3.90 – 3.85 (m, 2H), 3.54 – 3.48 (m, 2H), 3.40 (t, J = 5.1 Hz, 4H), 2.31 (t, J = 5.0 Hz, 4H), 2.19 (s, 3H) ppm. $^{13}C$ NMR (126 MHz, DMSO-$d_6$) δ 155.54, 155.05, 149.16, 144.40, 140.96, 135.66, 128.81, 125.03, 123.13, 115.79, 109.86, 104.27, 99.79, 65.16, 65.06, 63.86, 54.45, 45.78, 43.58 ppm. MS-ESI m/z $[M+H]^+$: calcd 438.22, found 438.20. HRMS *m/z* for $[C_{22}H_{27}N_7O_3+H]^+$: calcd 437.21699, found 437.21768. HPLC: $t_R$ = 3.045 min (M2), purity ≧ 95% (UV: 254/280 nm).

#### Analytical data of compounds 6-9, 11-15, 4, 16-35

$^{1}$H and $^{13}$C {H} NMR and HPLC/MS (ESI) data of compound **6**

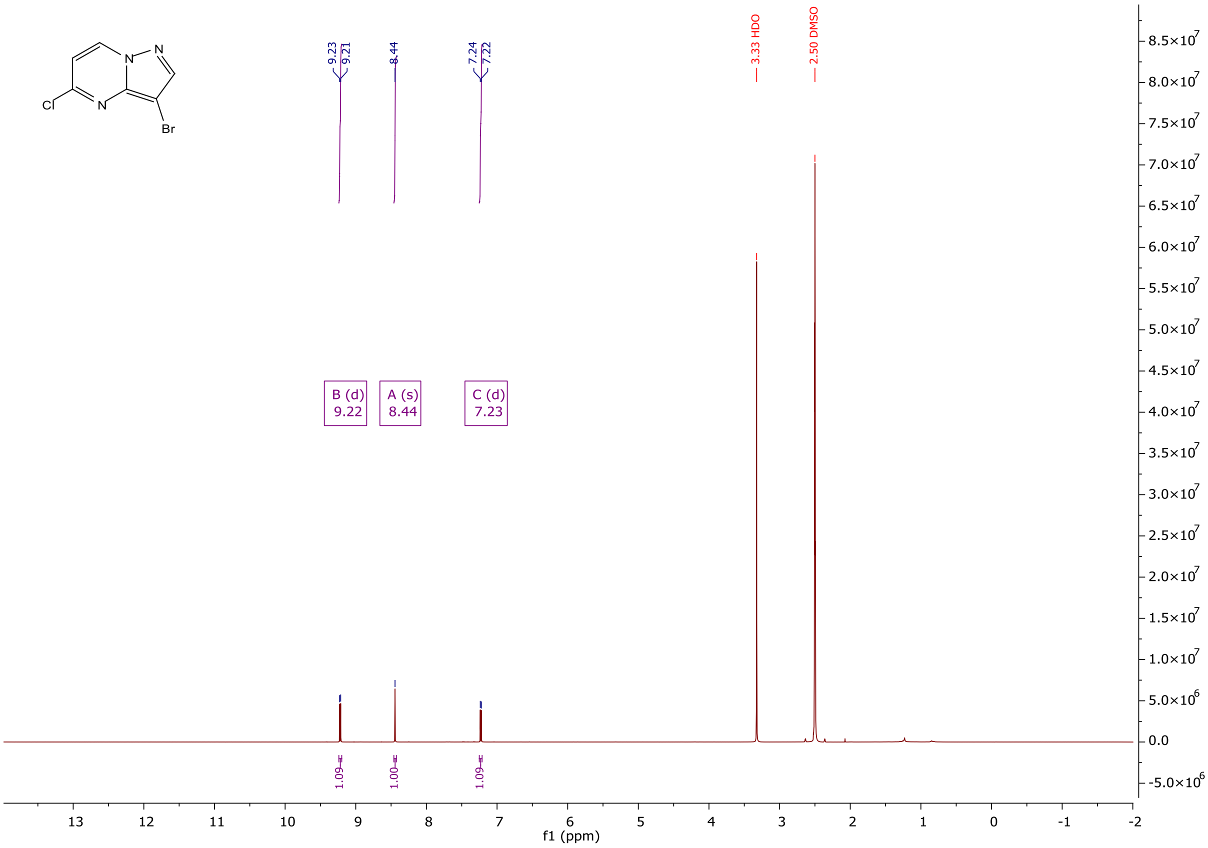

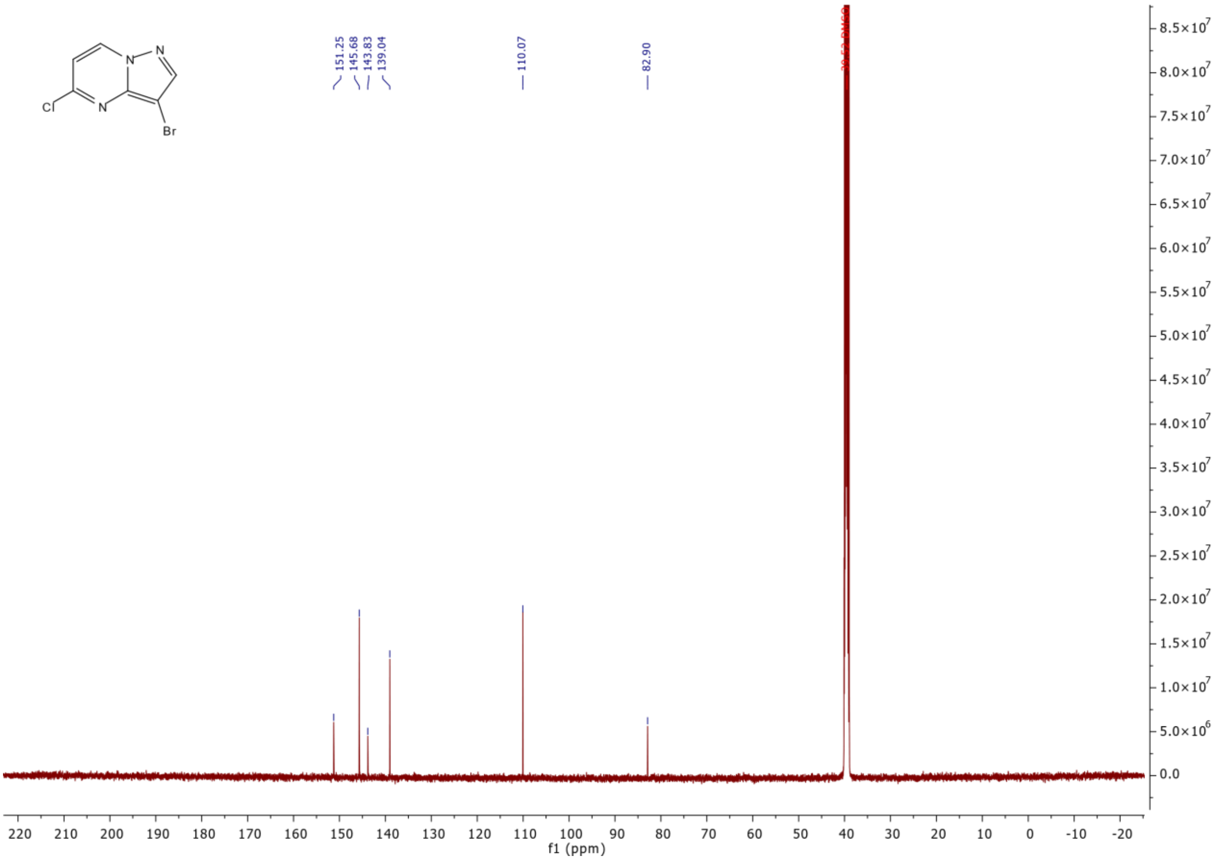

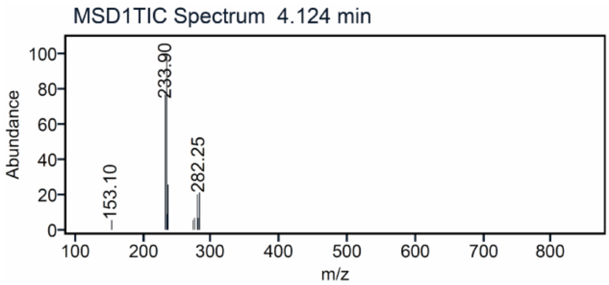

$^{1}H$ and $^{13}C$ {H} NMR and HPLC/MS (ESI) data of compound **7**

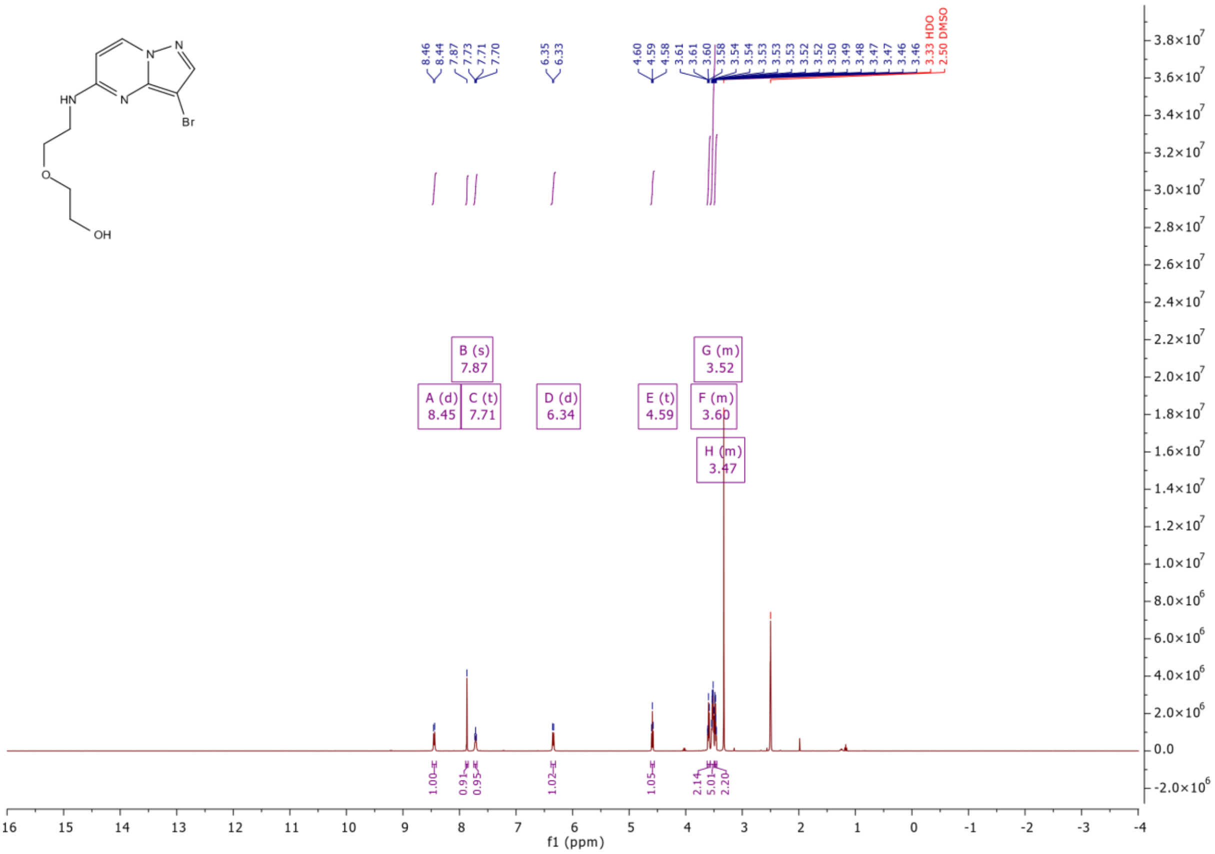

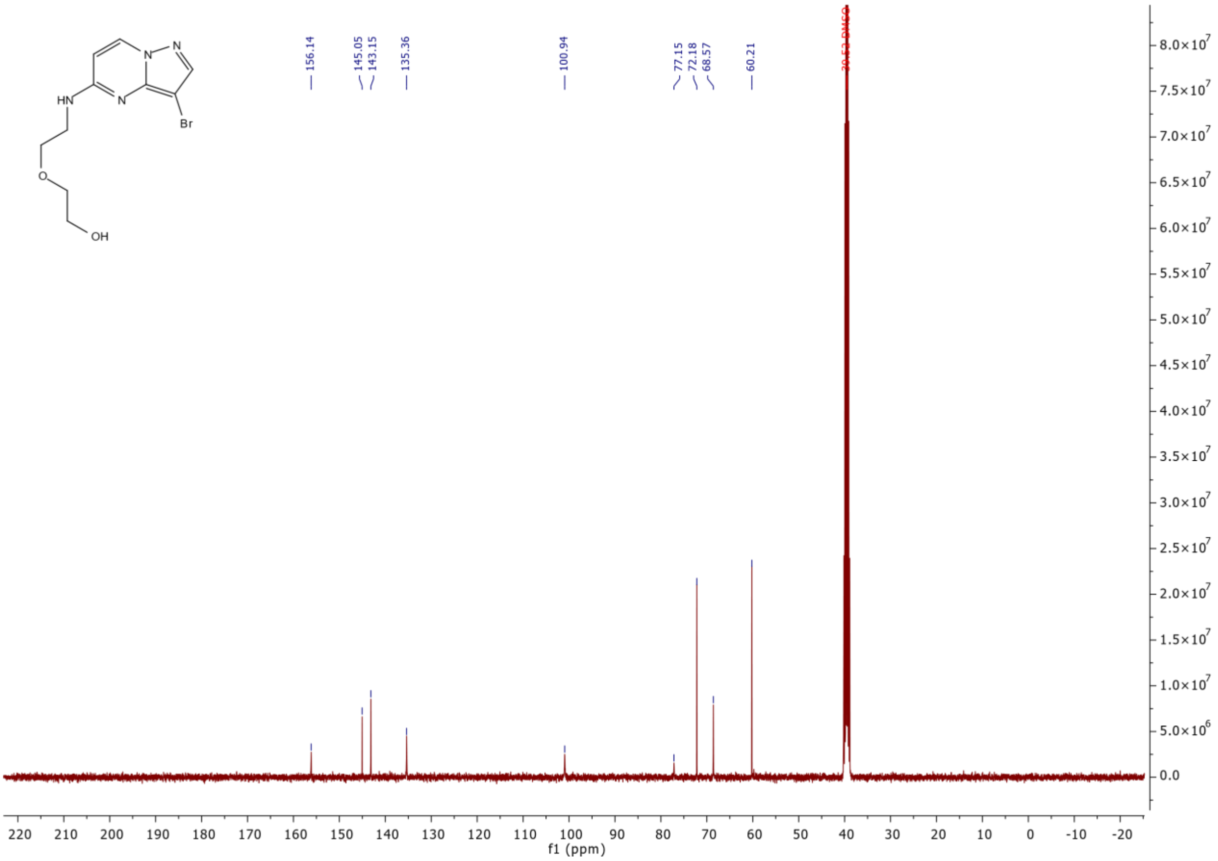

156.14
145.05
143.15
135.36
100.94
77.15
72.18
68.57
60.21
HN
N
N
N
Br
O
OH
f1 (ppm)

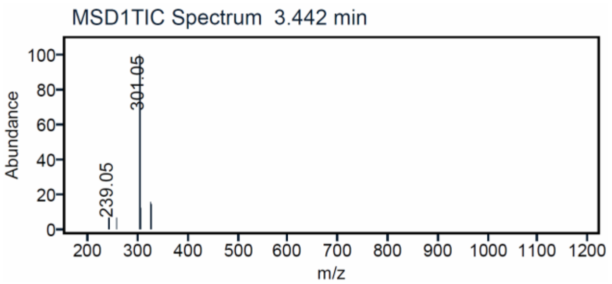

MSD1TIC Spectrum 3.442 min
Abundance
301.05
239.05
m/z

#### $^{1}H$ and $^{13}C$ {H} NMR and HPLC/MS (ESI) data of compound **8**

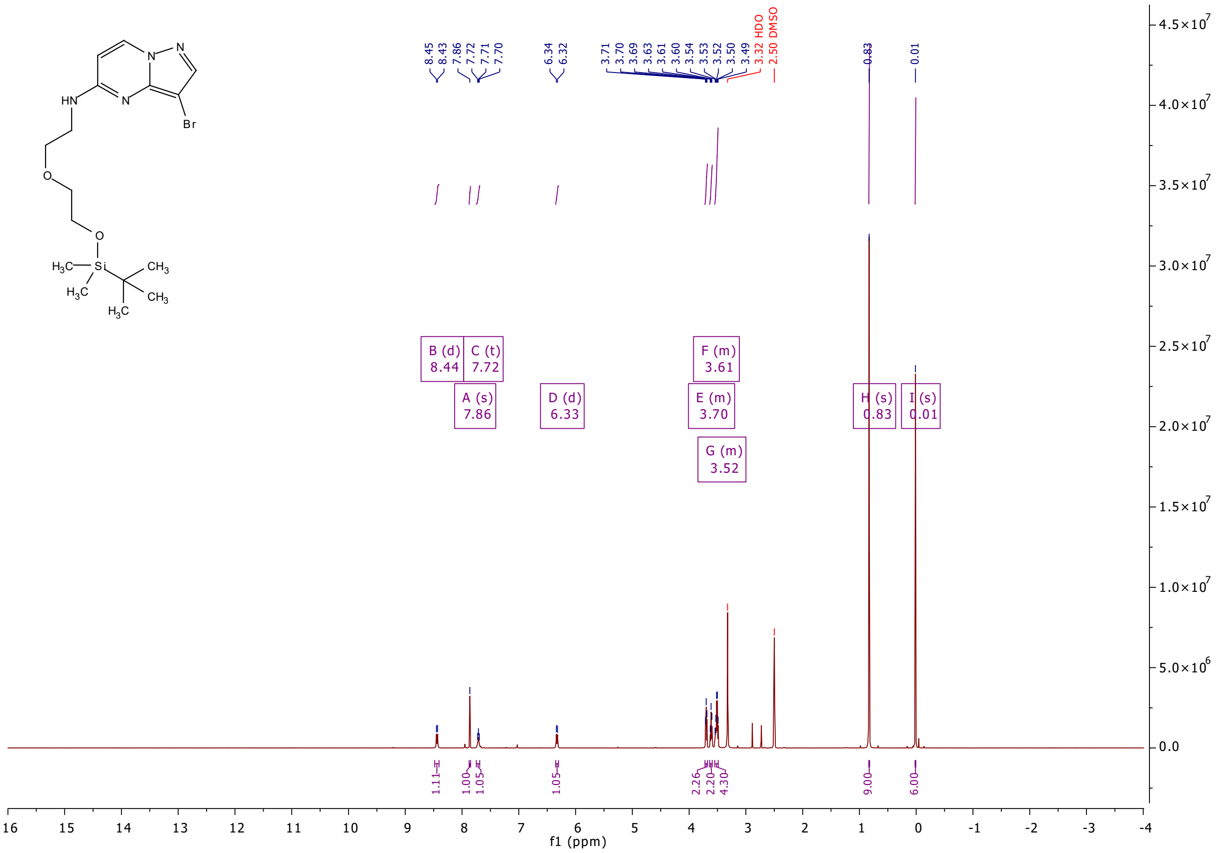

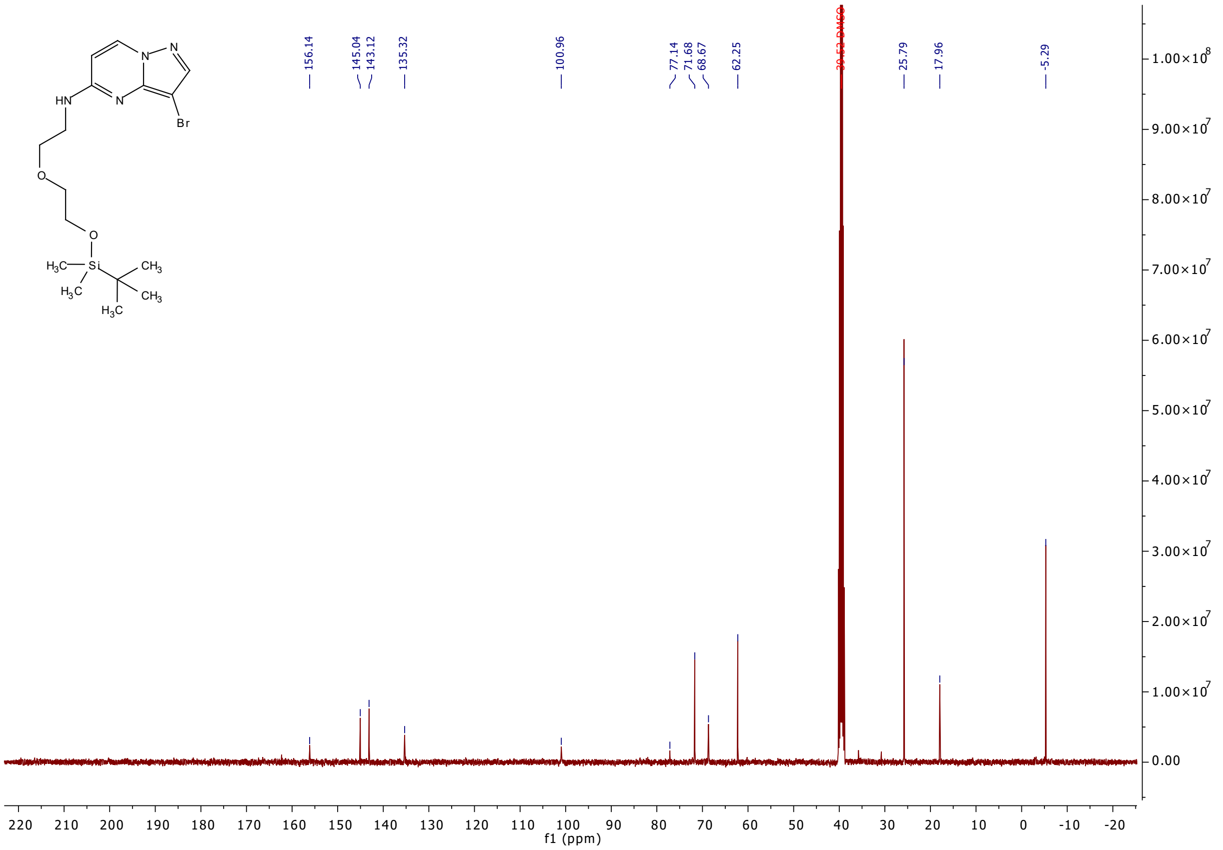

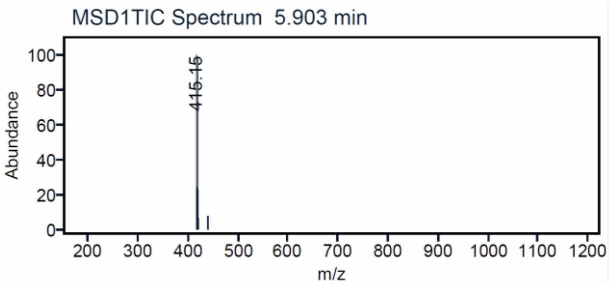

[1]H and [13]C {H} NMR and HPLC/MS (ESI) data of compound **9**

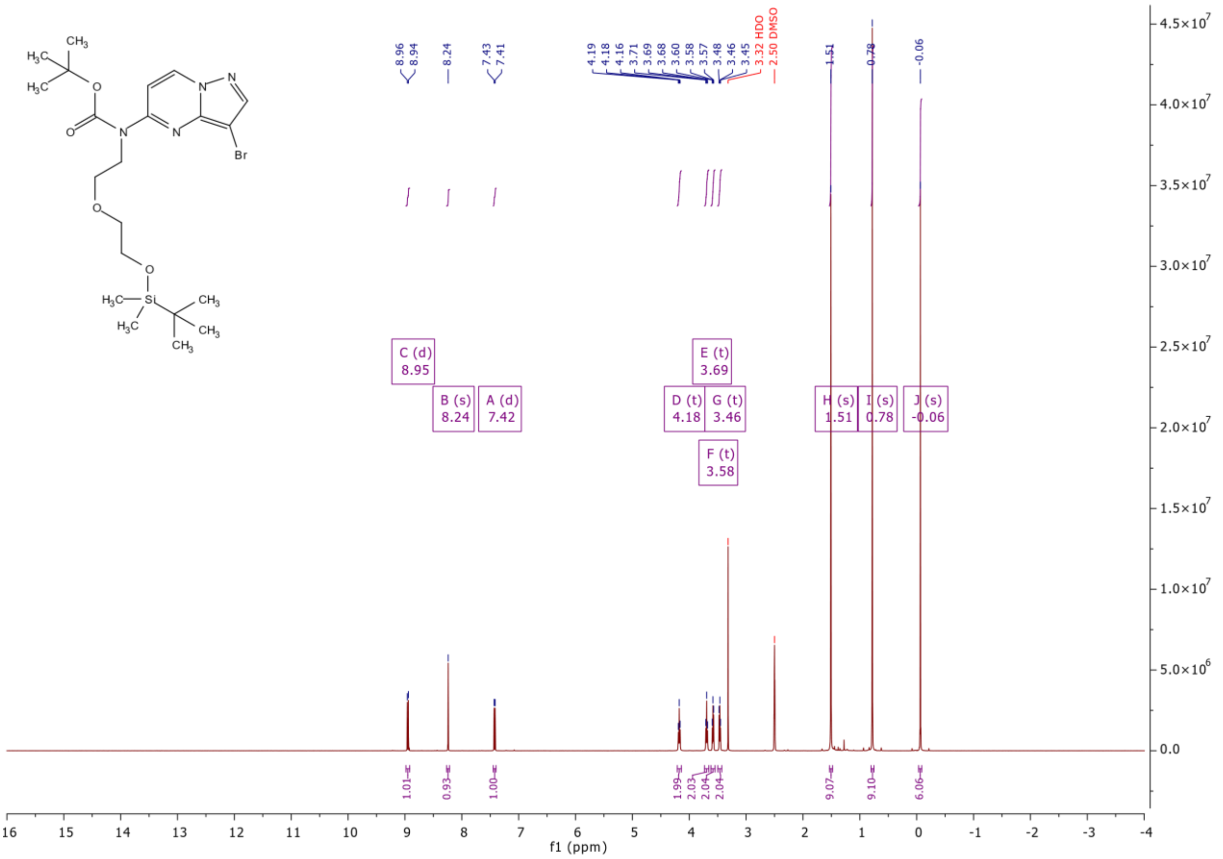

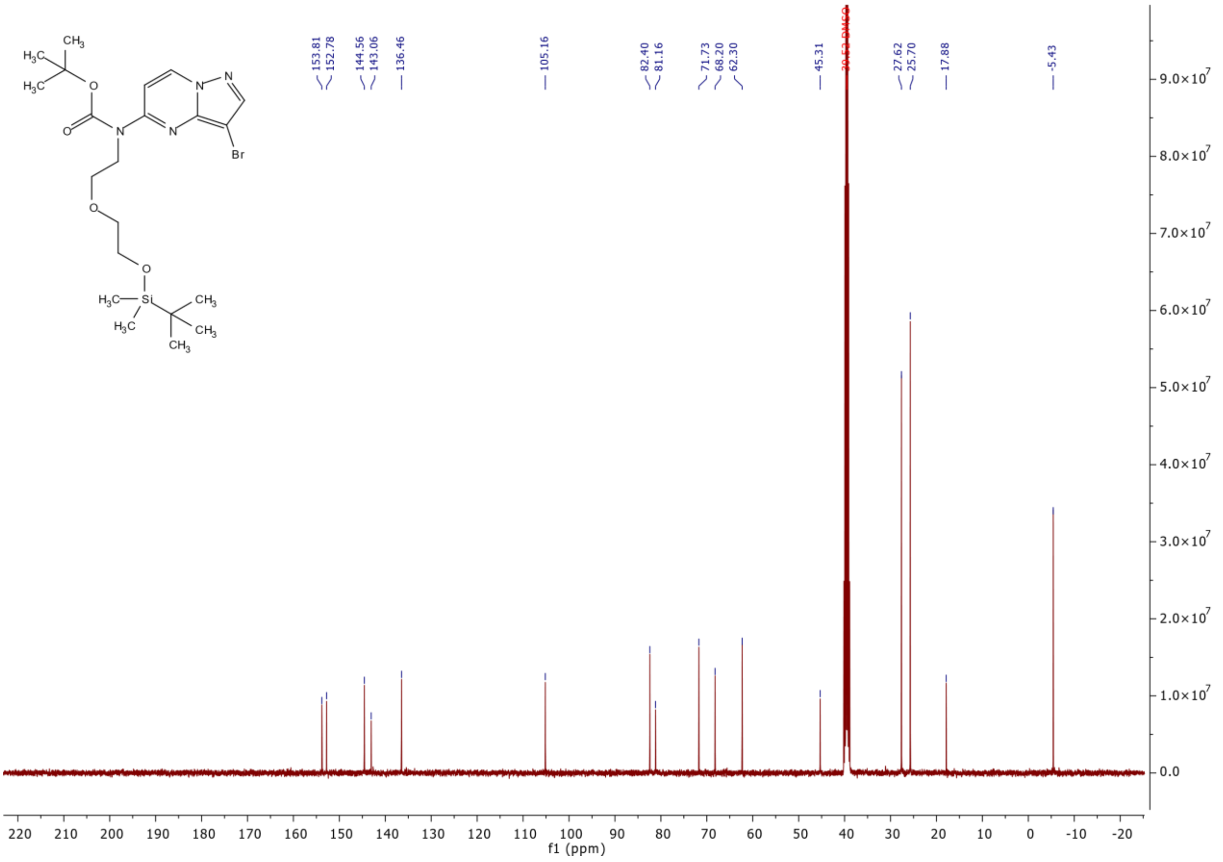

$^{1}$H and $^{13}$C {H} NMR and HPLC/MS (ESI) data of compound **11**

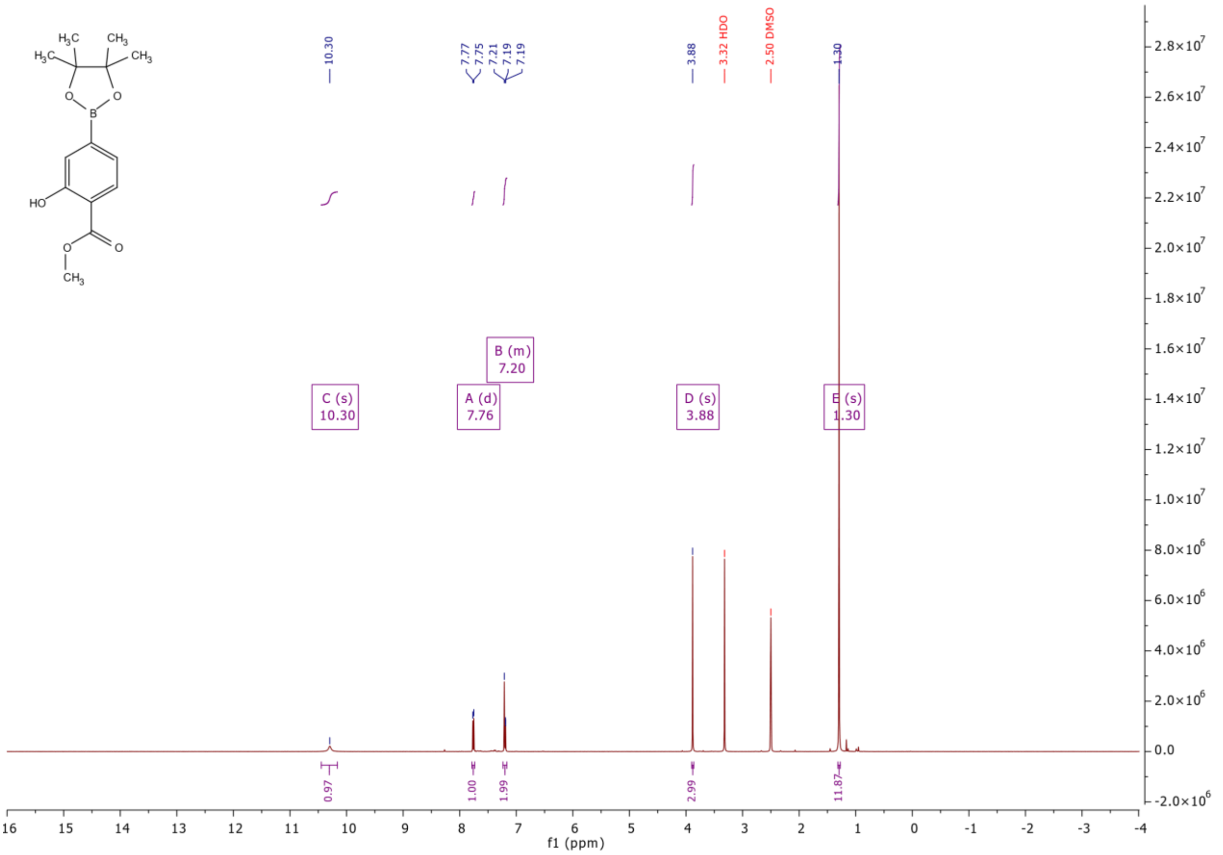

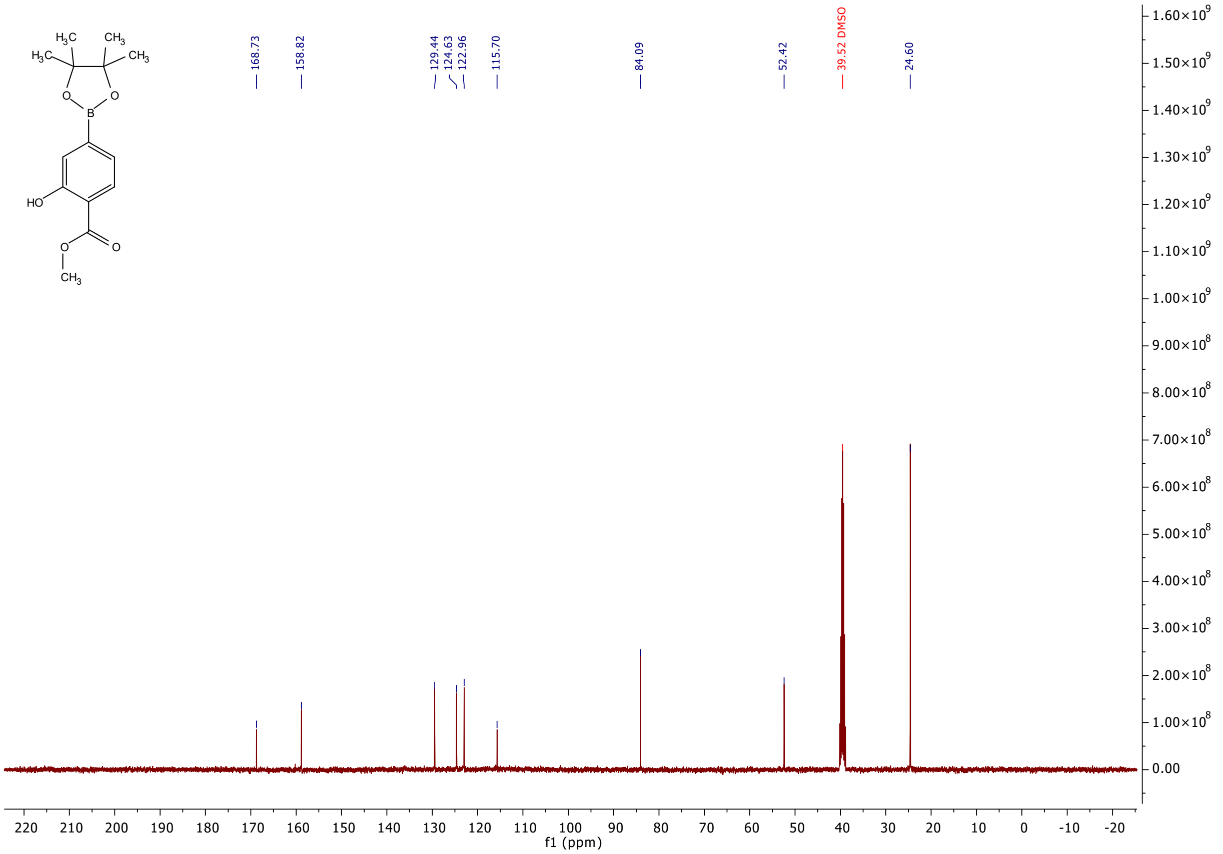

168.73
158.82
129.44
124.63
122.96
115.70
84.09
52.42
39.52 DMSO
24.60
f1 (ppm)

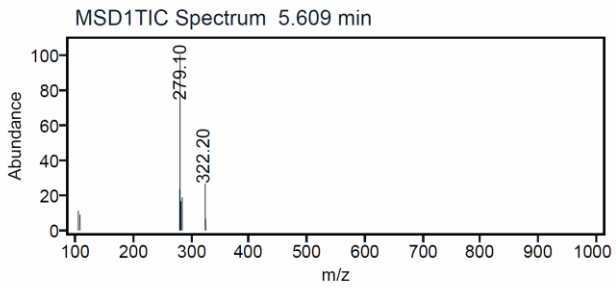

MSD1TIC Spectrum 5.609 min
Abundance
279.10
322.20
m/z

[1]H and [13]C {H} NMR and HPLC/MS (ESI) data of compound **12**

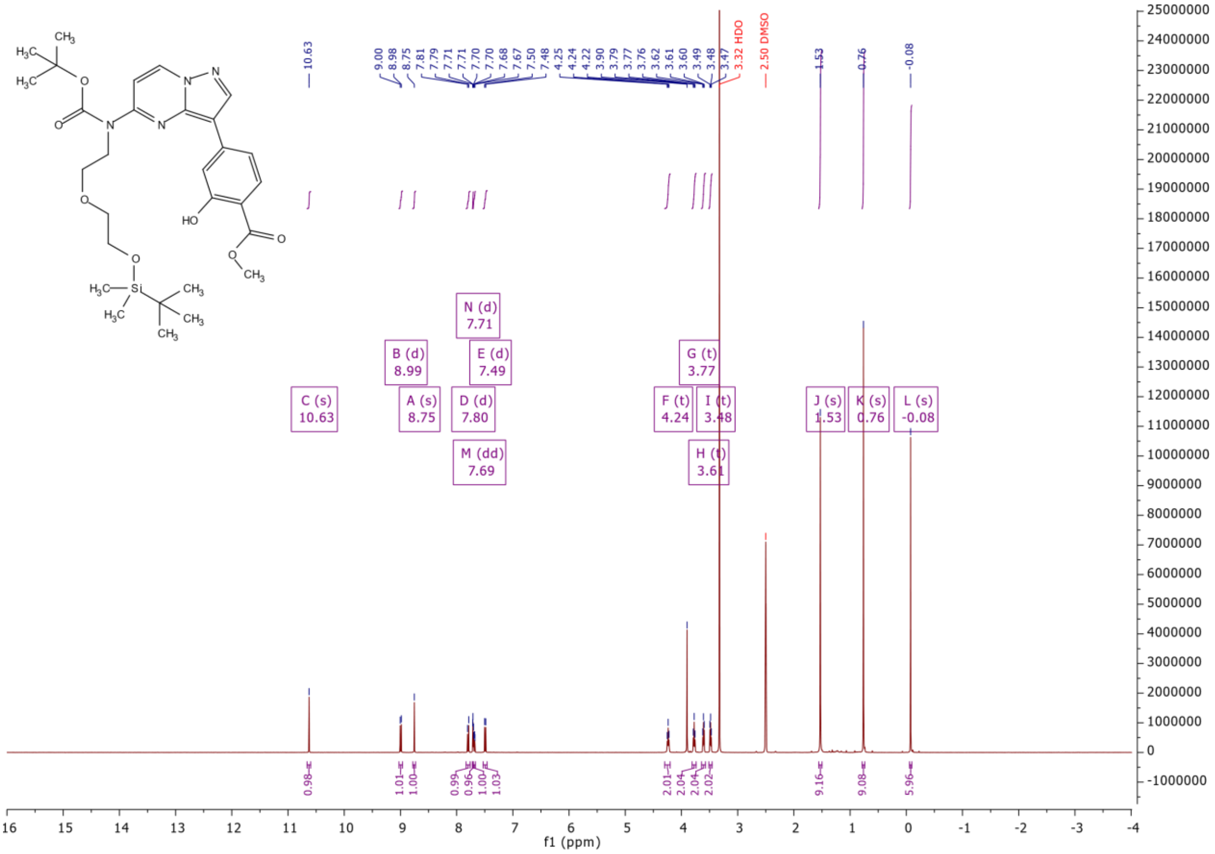

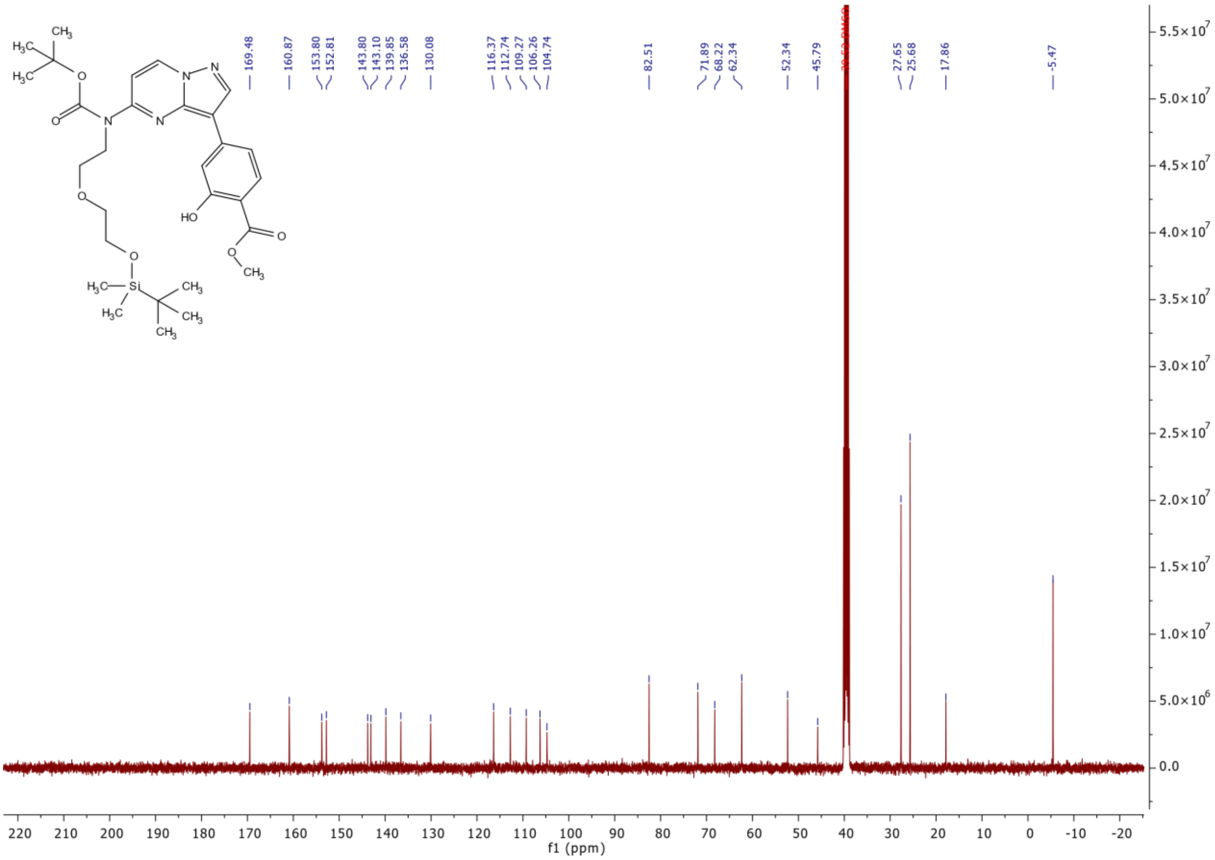

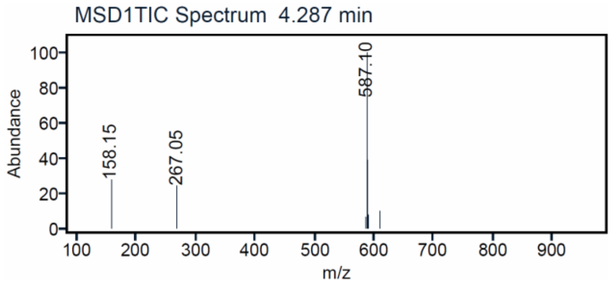

$^{1}H$ and $^{13}C$ {H} NMR and HPLC/MS (ESI) data of compound **13**

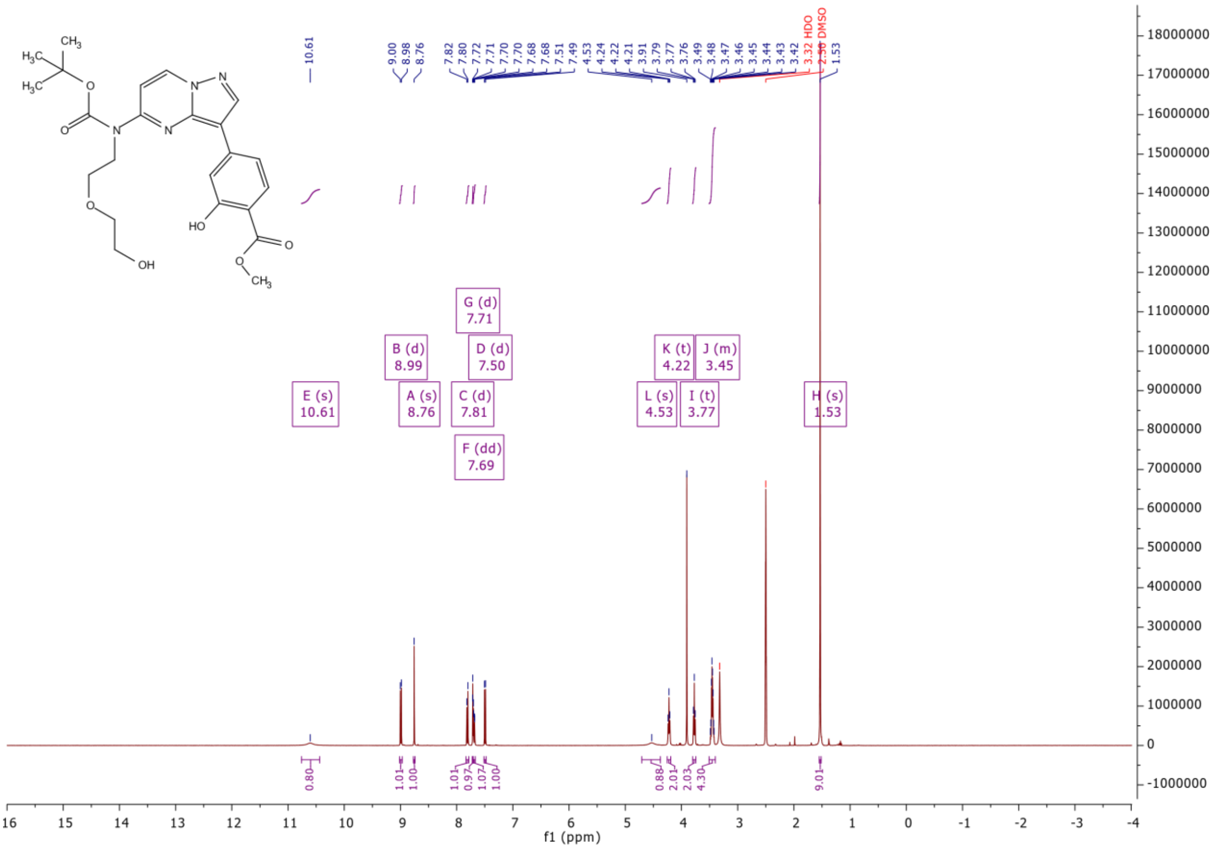

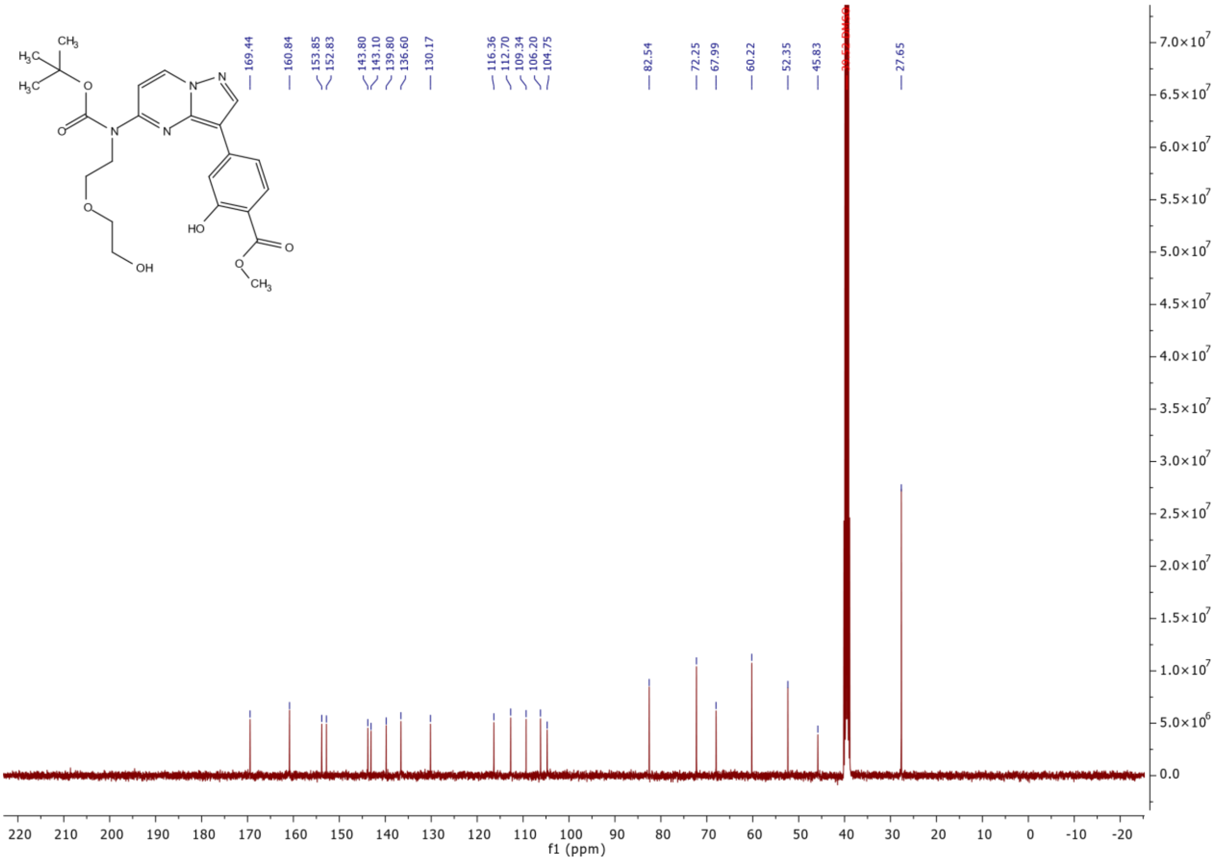

169.44
160.84
153.85
152.83
143.80
143.10
139.80
136.60
130.17
116.36
112.70
109.34
106.20
104.75
82.54
72.25
67.99
60.22
52.35
45.83
27.65
f1 (ppm)

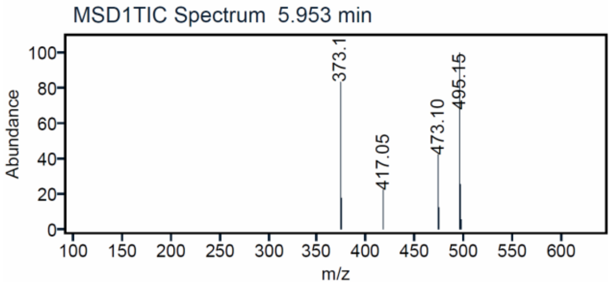

MSD1TIC Spectrum 5.953 min
Abundance
m/z
373.1
417.05
473.10
495.15

[1]H and [13]C {H} NMR and HPLC/MS (ESI) data of compound **14**

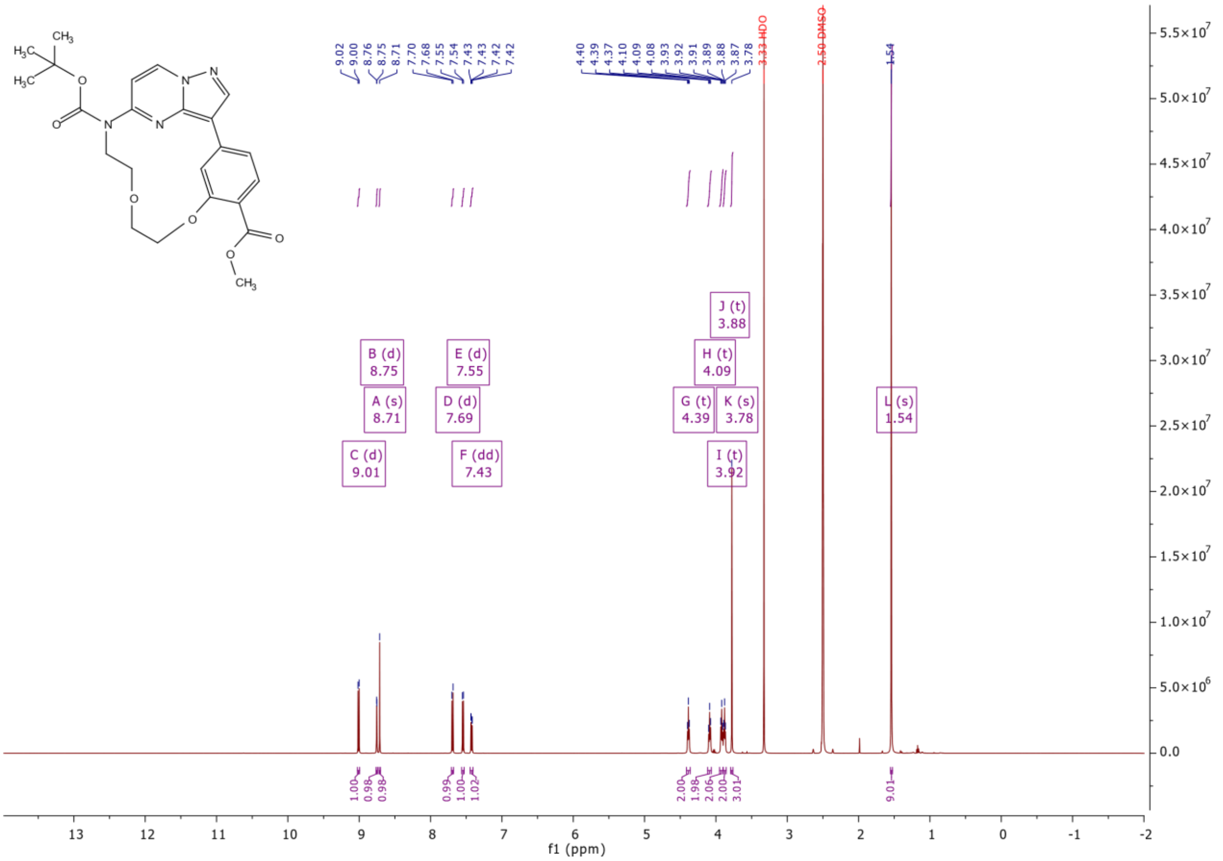

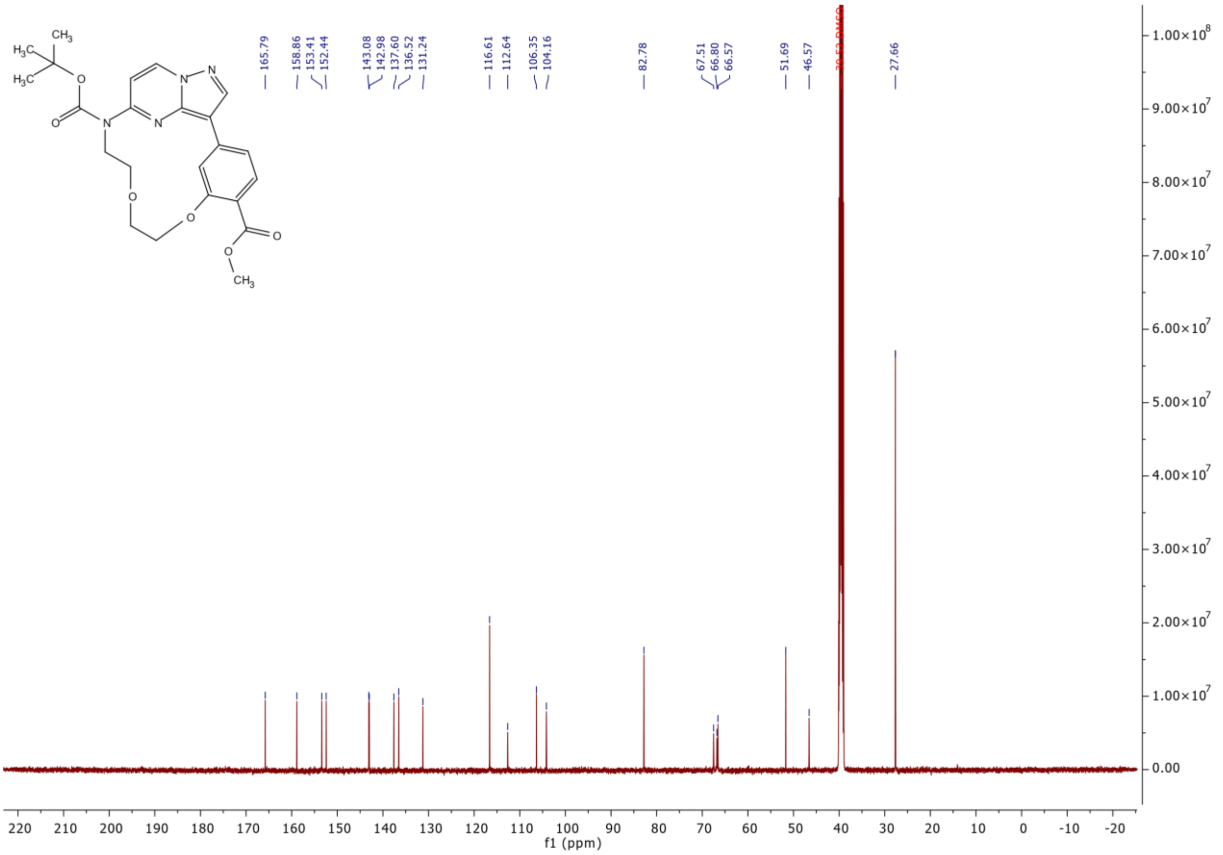

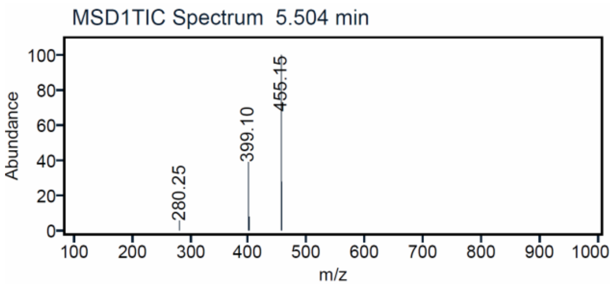

[1]H and [13]C {H} NMR and HPLC/MS (ESI) data of compound **15**

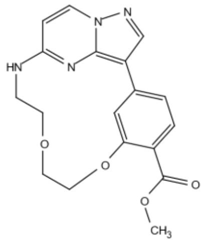

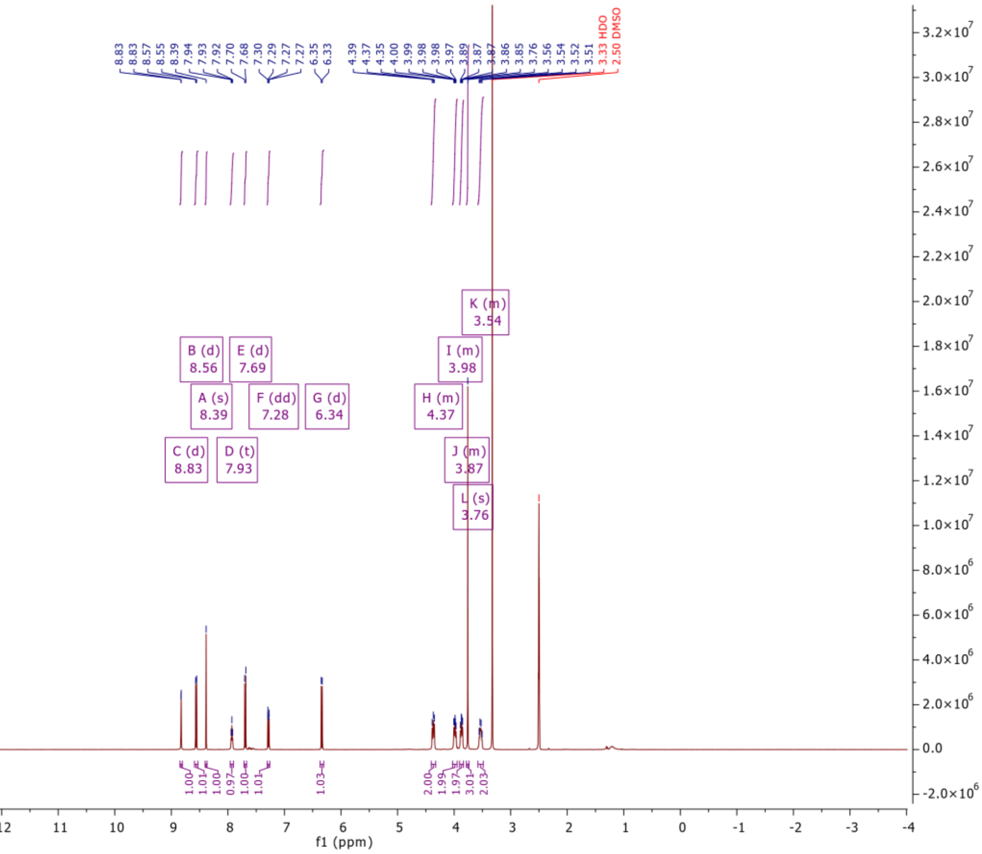

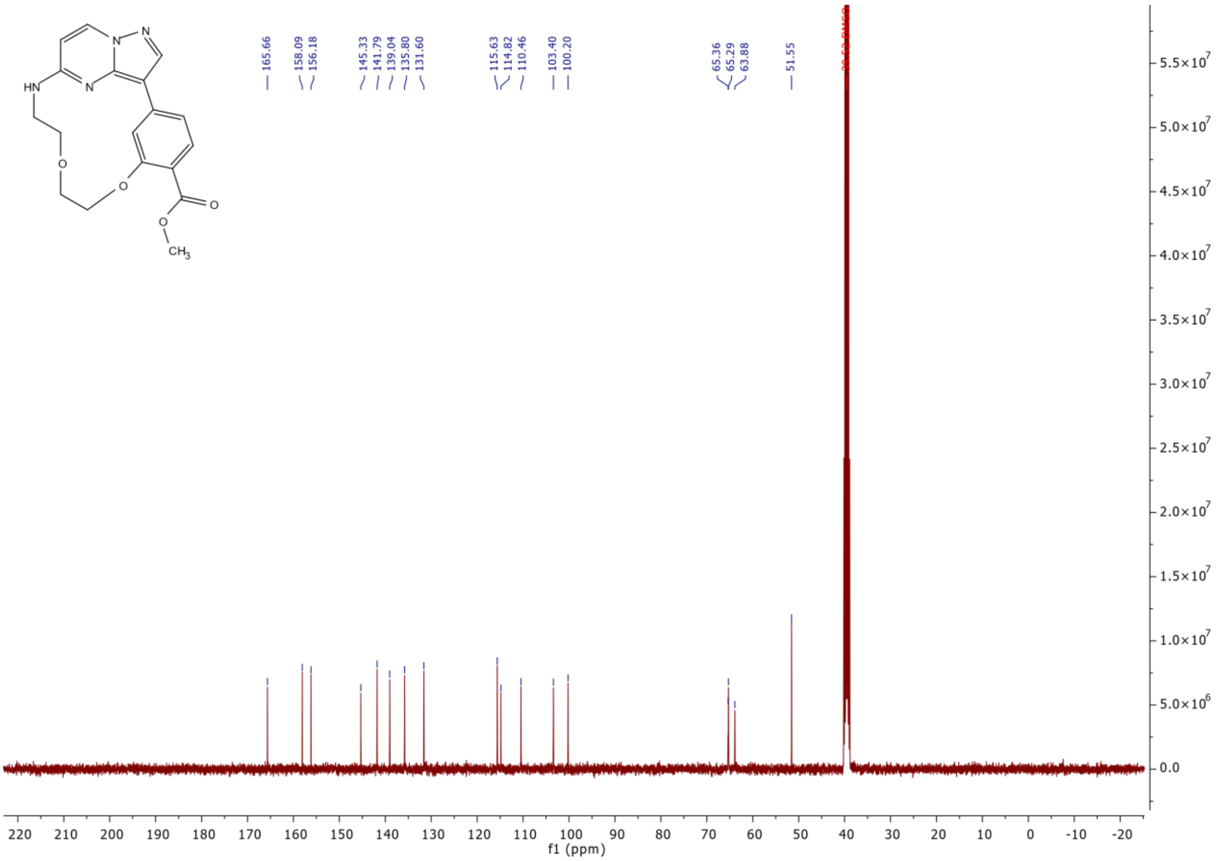
HN
N
N
N
O
O
O
O
CH3
165.66
158.09
156.18
145.33
141.79
139.04
135.80
131.60
115.63
114.82
110.46
103.40
100.20
65.36
65.29
63.88
51.55
f1 (ppm)

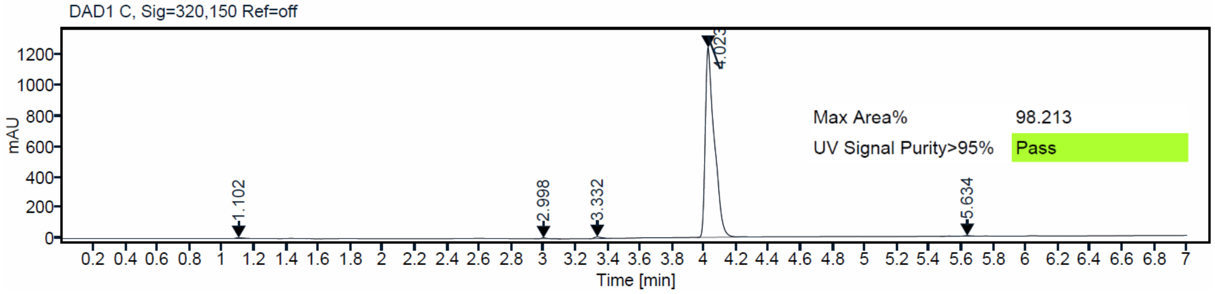
DAD1 C, Sig=320,150 Ref=off
mAU
Time [min]
1.102
2.998
3.332
4.023
5.634
Max Area% 98.213
UV Signal Purity>95% Pass

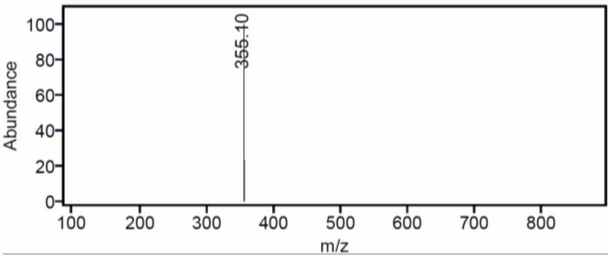
Abundance
m/z
355.10

[1]H and [13]C {H} NMR and HPLC/MS (ESI) data of compound **4**

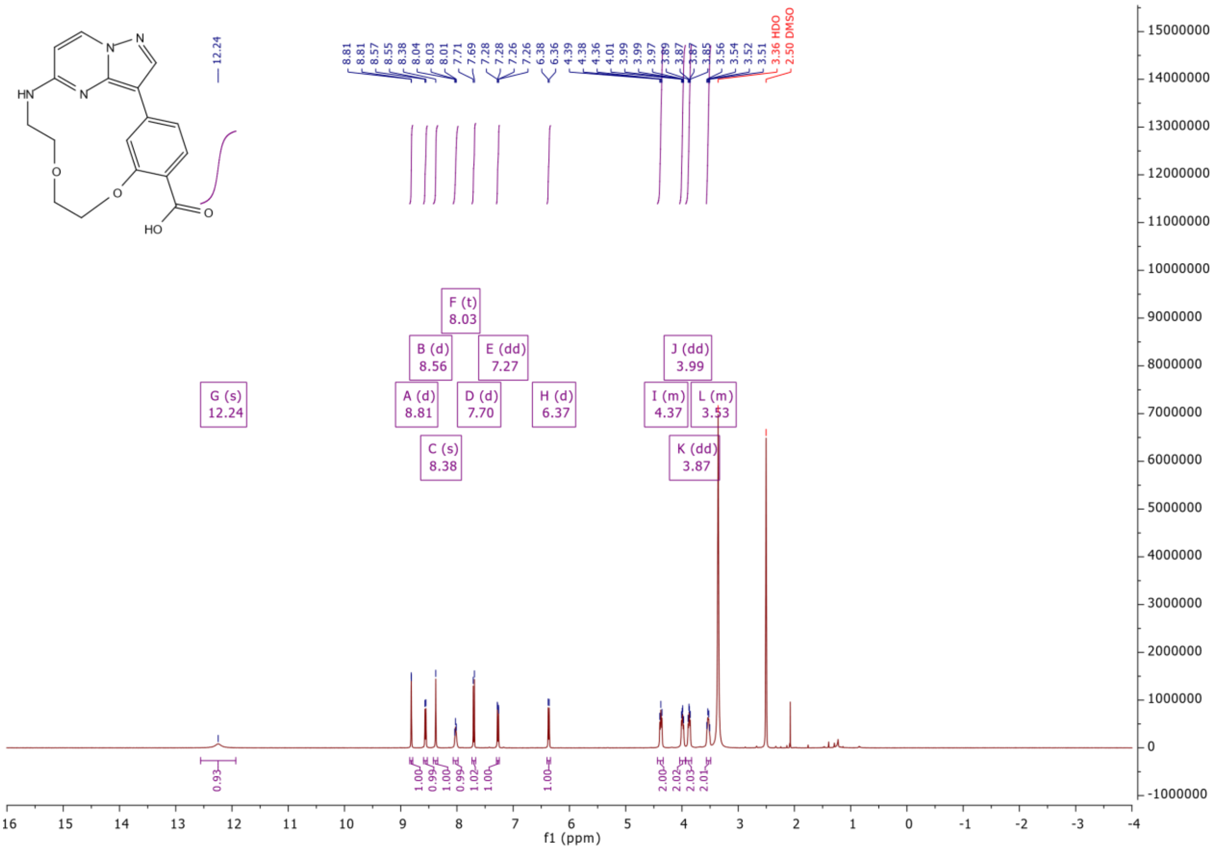

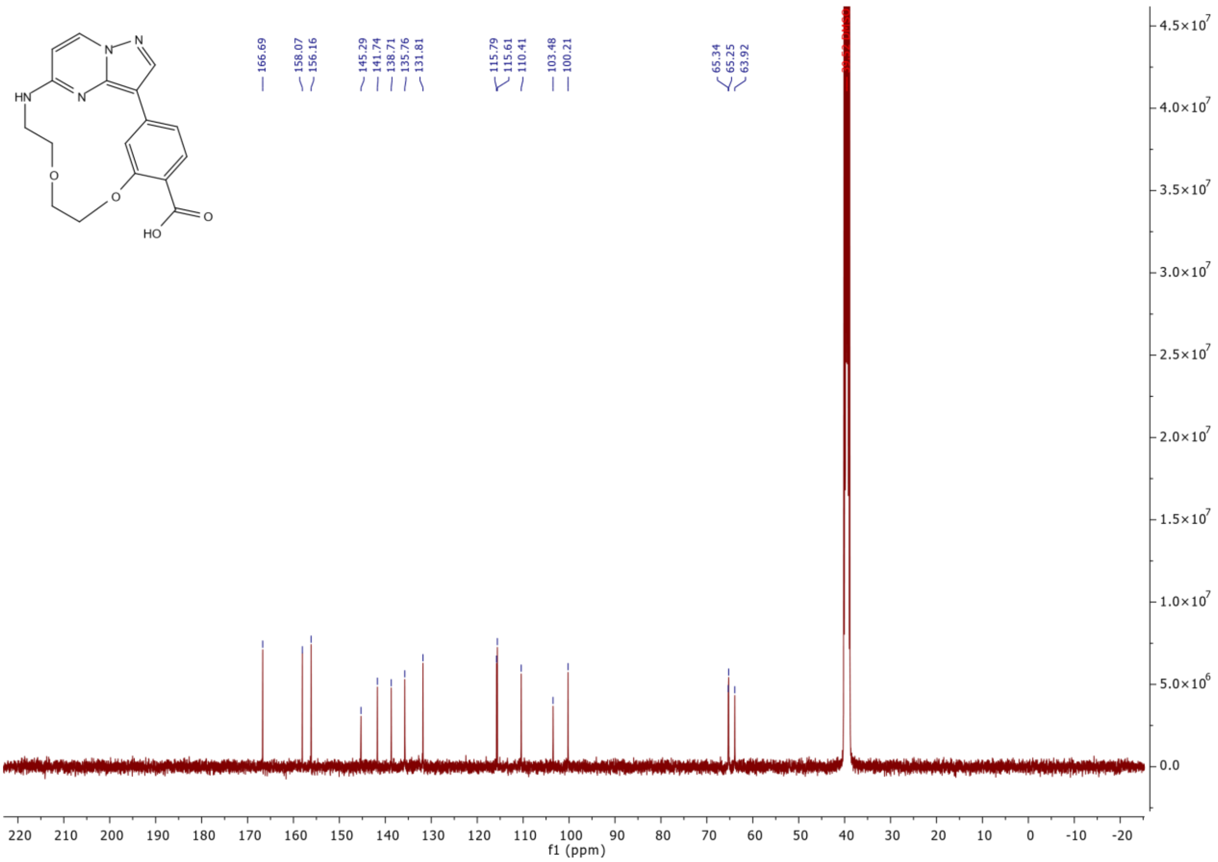

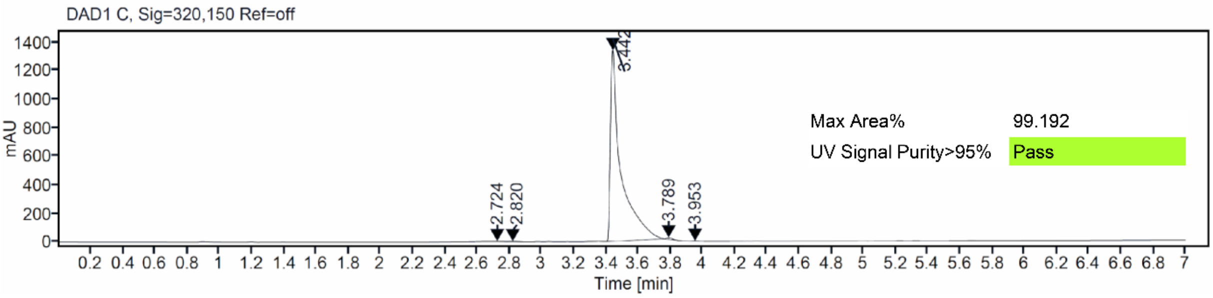

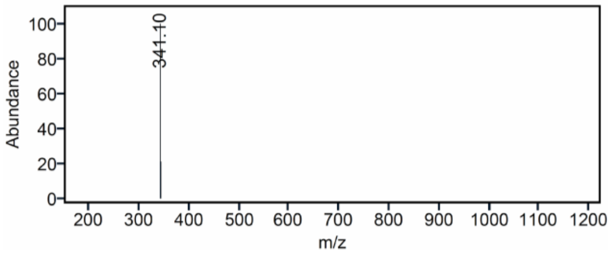

[1]H and [13]C {H} NMR, HPLC/MS (ESI) and HRMS data of compound **16**

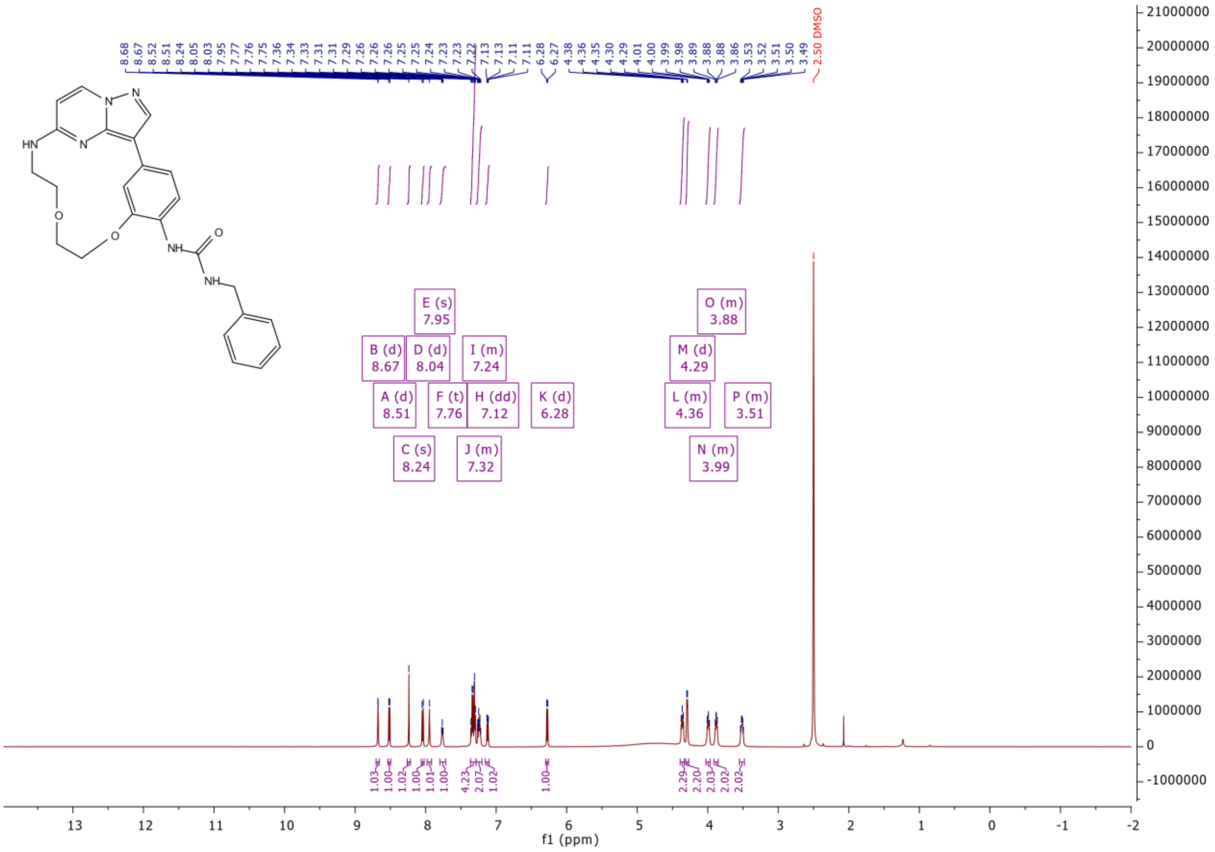

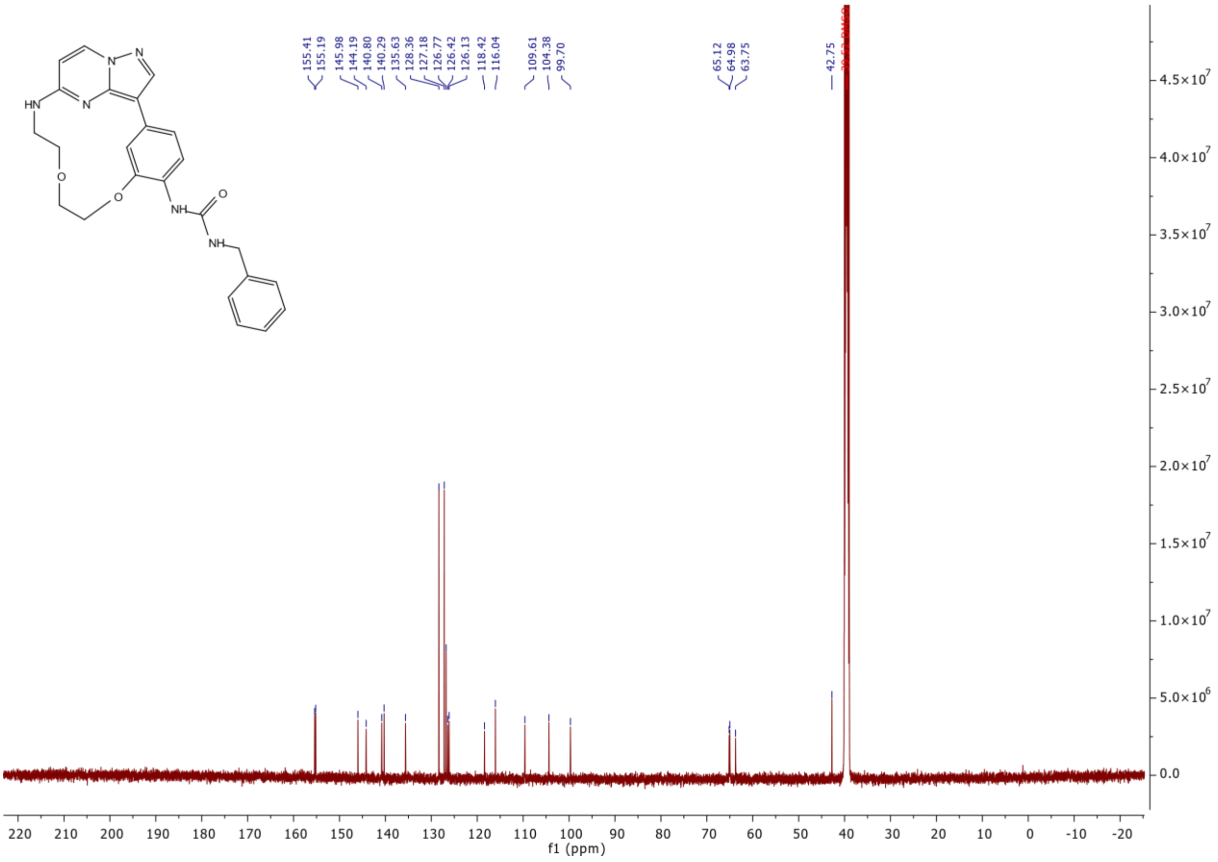
155.41
155.19
145.98
144.19
140.80
140.29
135.63
128.36
127.18
126.77
126.42
126.13
118.42
116.04
109.61
104.38
99.70
65.12
64.98
63.75
42.75
f1 (ppm)

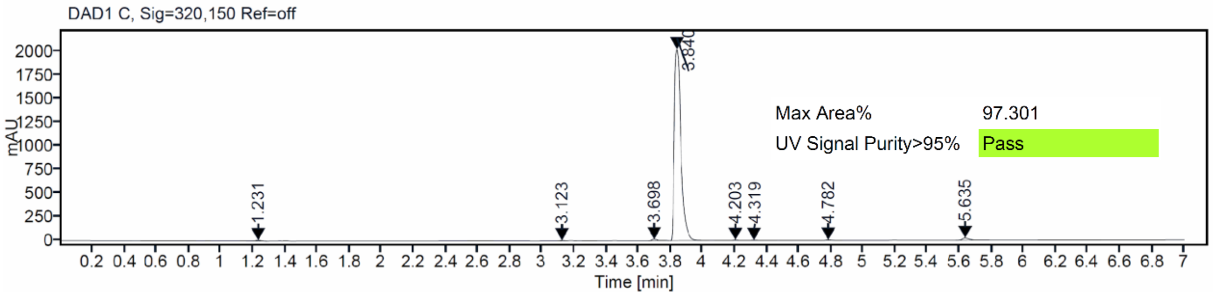
DAD1 C, Sig=320,150 Ref=off
Max Area% 97.301
UV Signal Purity>95% Pass
1.231
3.123
3.698
3.840
4.203
4.319
4.782
5.635
mAU
Time [min]

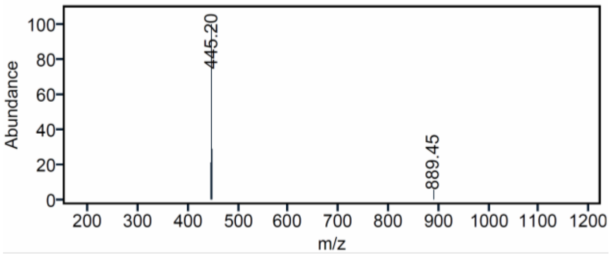
445.20
889.45
Abundance
m/z

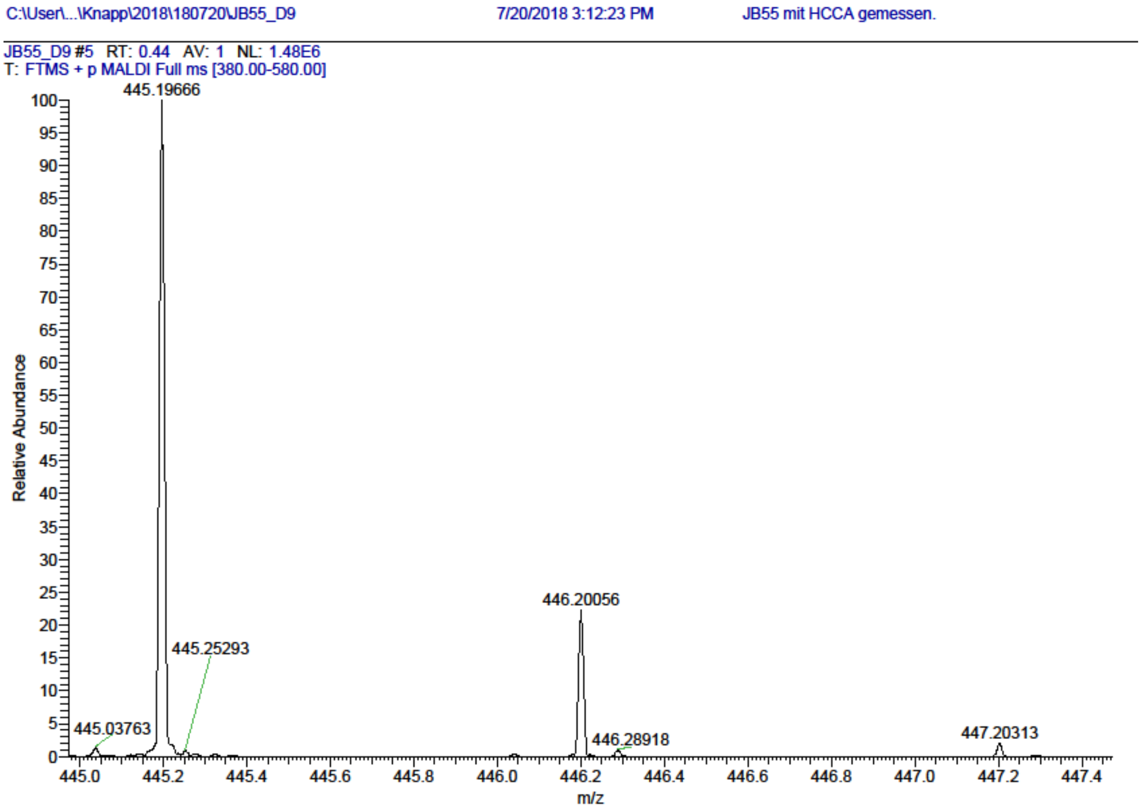

$^1$H and $^{13}$C {H} NMR, HPLC/MS (ESI) and HRMS data of compound **17**

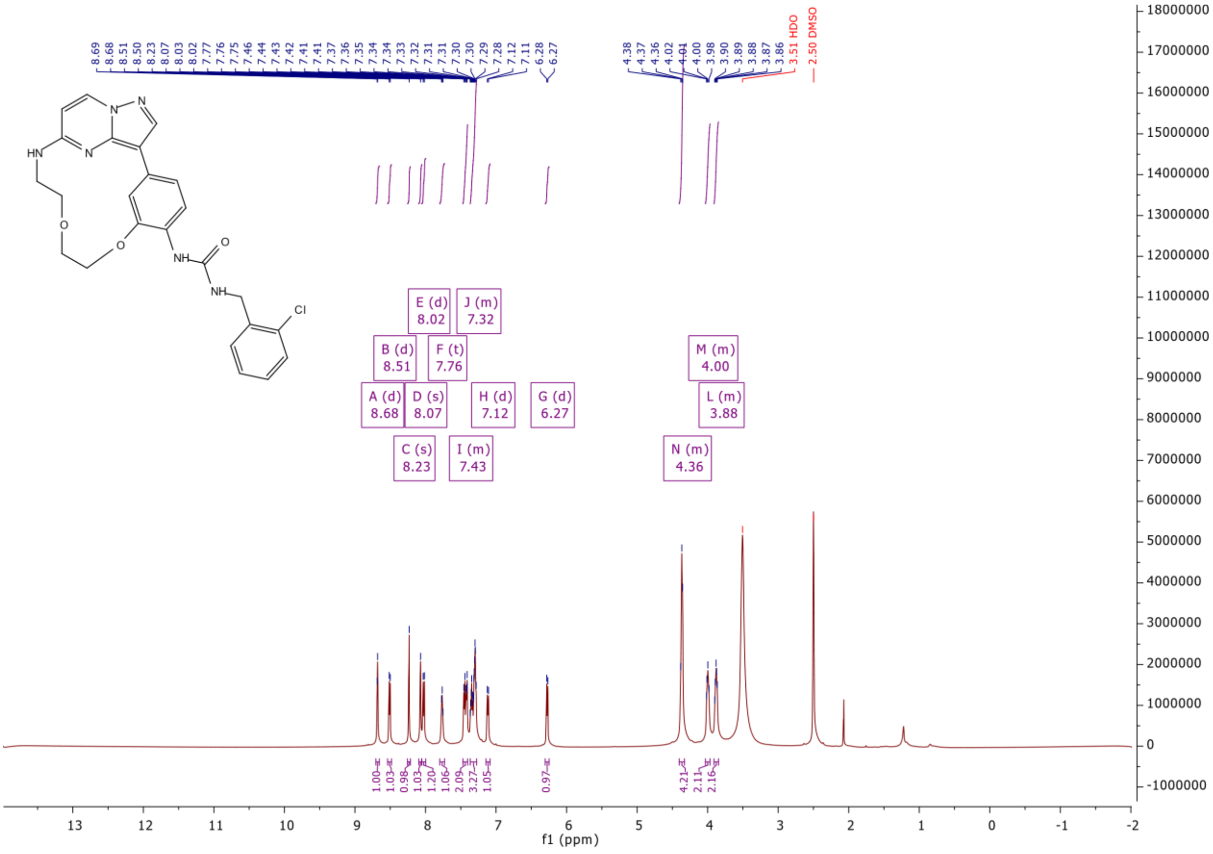

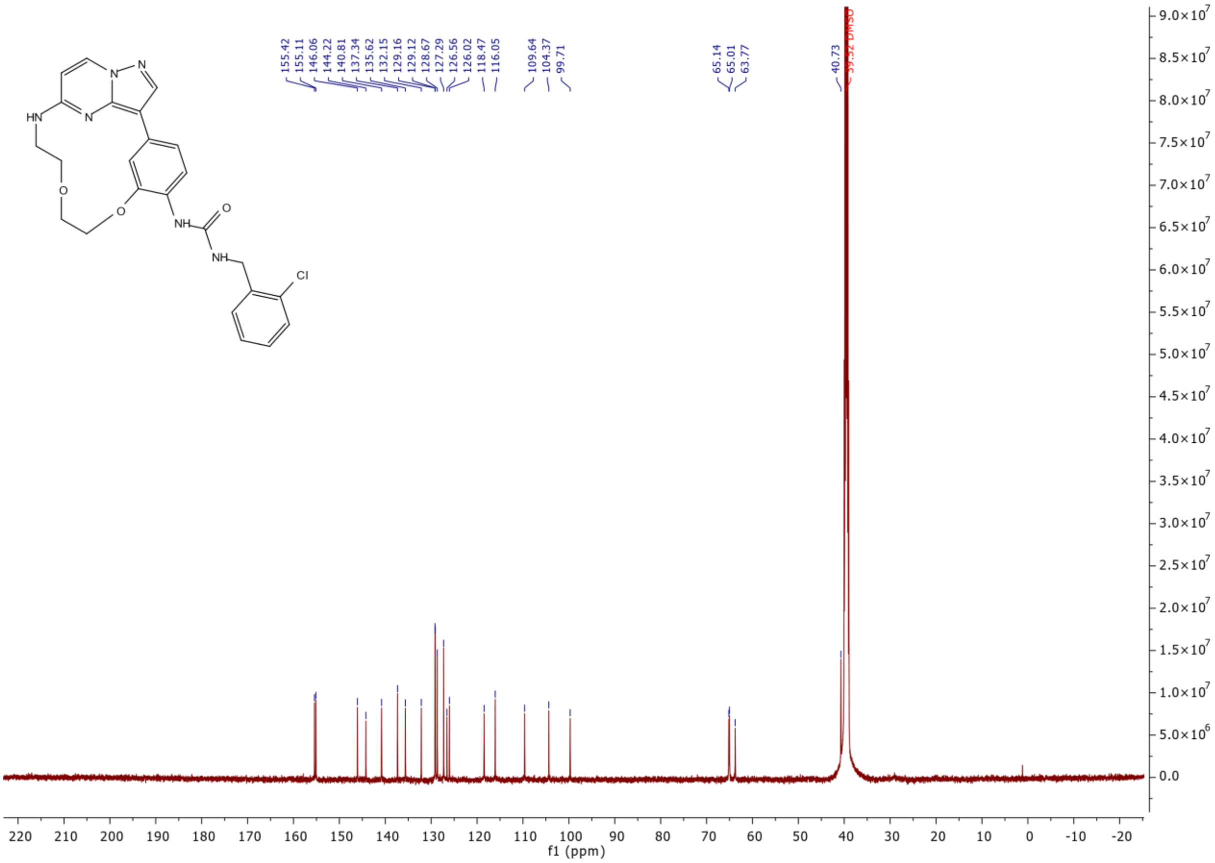
HN
N
N
N
O
O
NH
O
NH
Cl
155.42
155.11
146.06
144.22
140.81
137.34
135.62
132.15
129.16
129.12
128.67
127.29
126.56
126.02
118.47
116.05
109.64
104.37
99.71
65.14
65.01
63.77
40.73
39.52 DMSO
f1 (ppm)

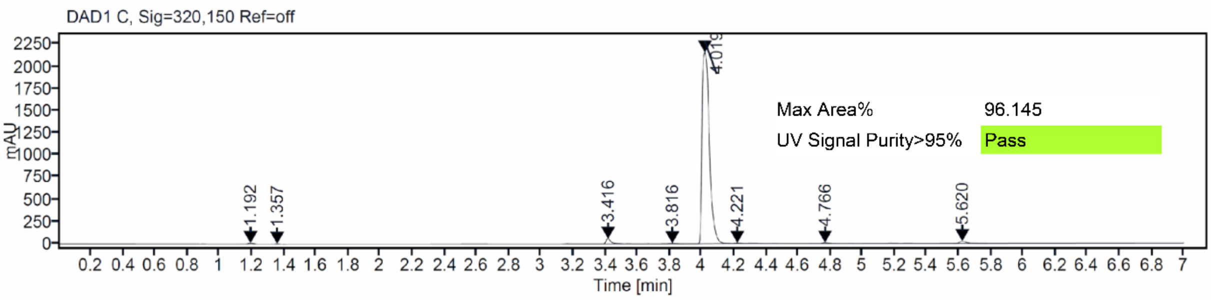
DAD1 C, Sig=320,150 Ref=off
mAU
4.019
1.192
1.357
3.416
3.816
4.221
4.766
5.620
Max Area% 96.145
UV Signal Purity>95% Pass
Time [min]

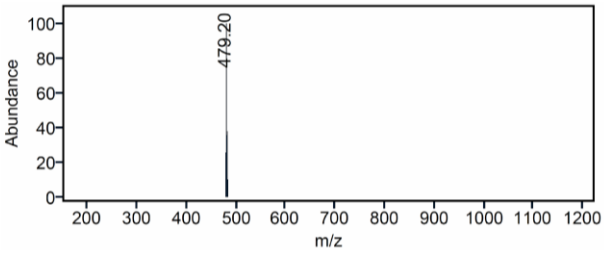
479.20
Abundance
m/z

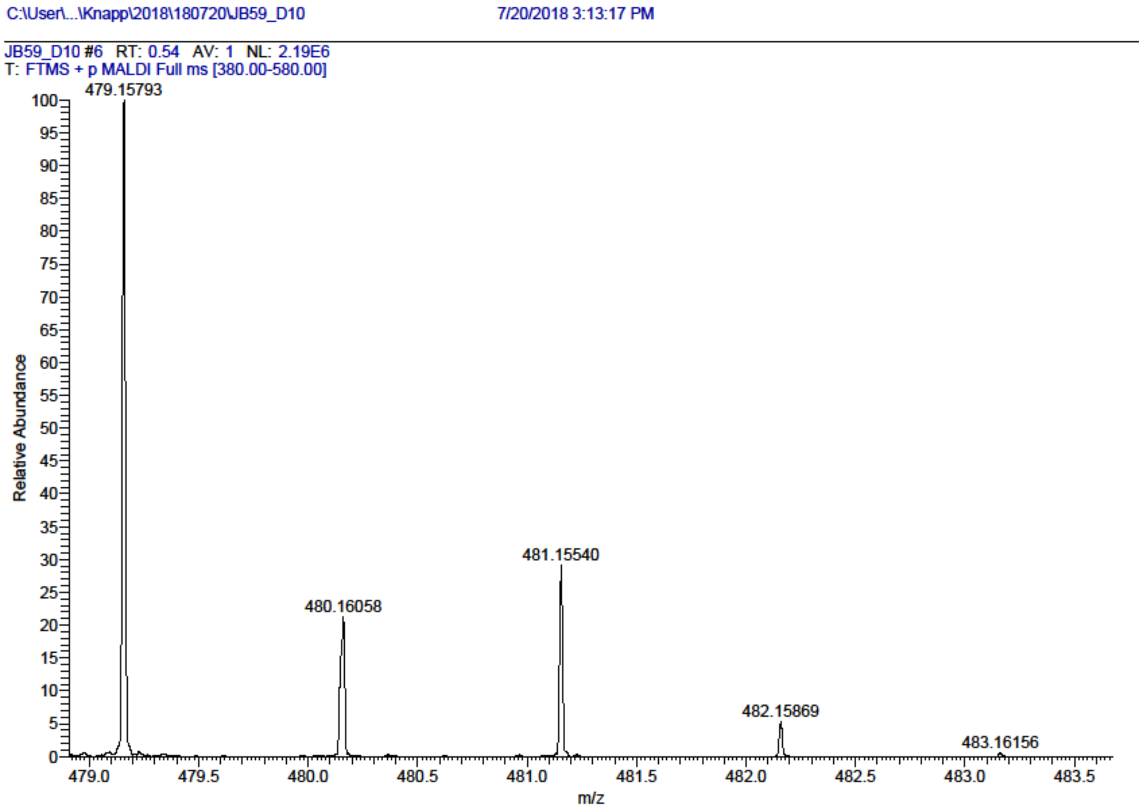

[1]H and [13]C {H} NMR, HPLC/MS (ESI) and HRMS data of compound **18**

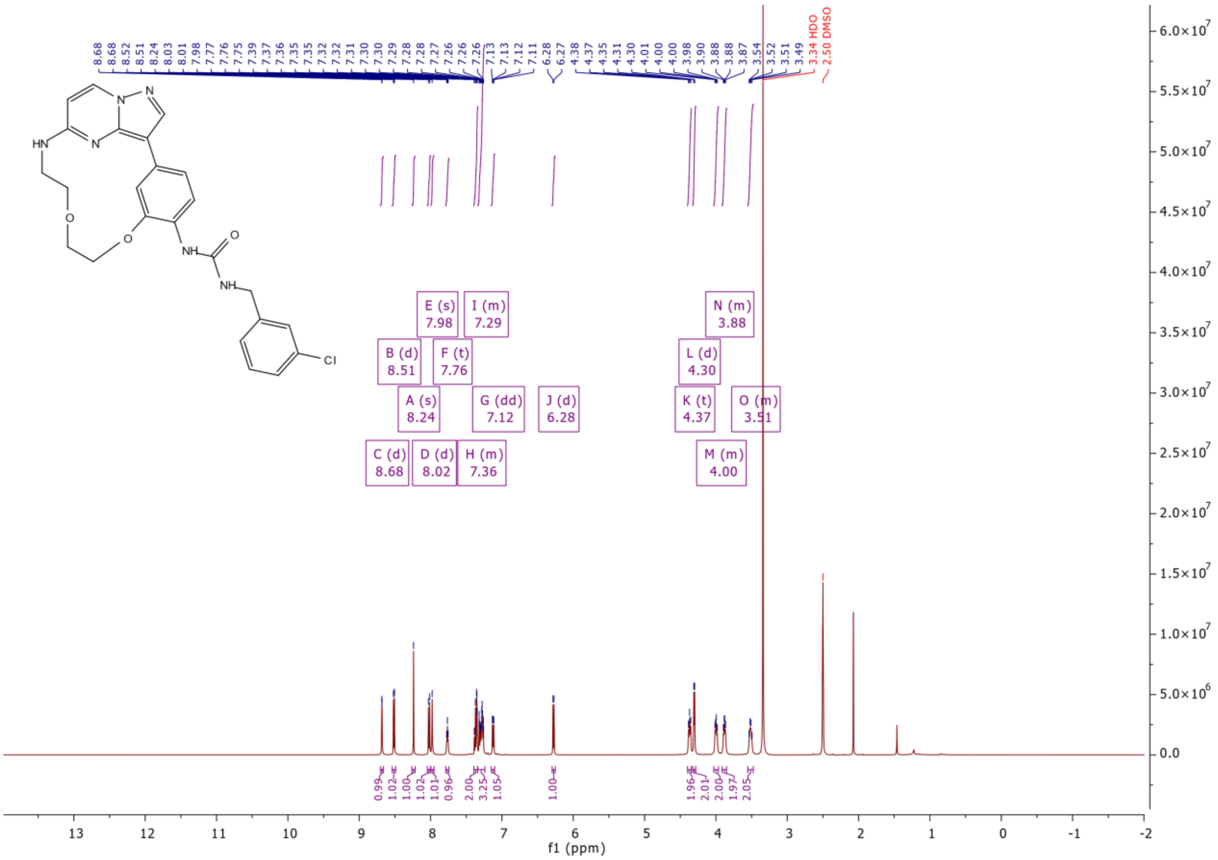

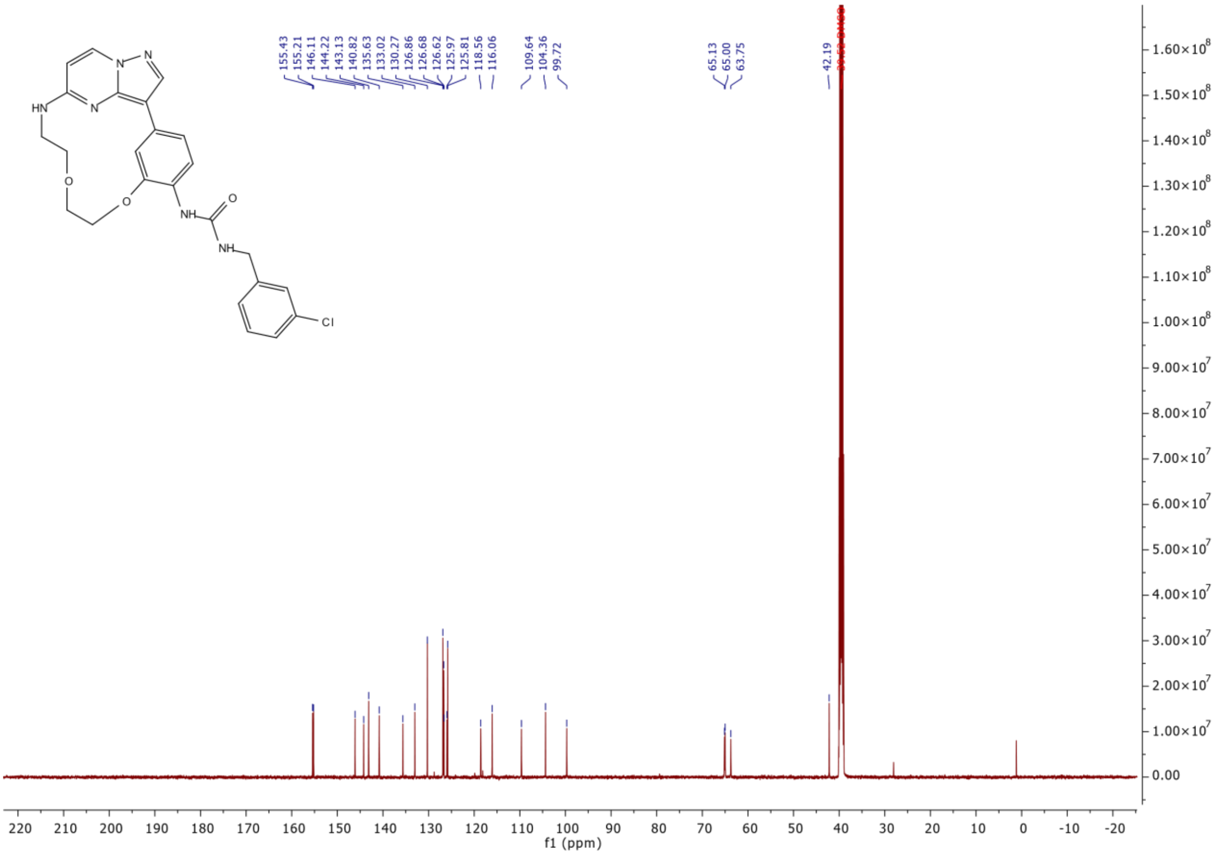
155.43
155.21
146.11
144.22
143.13
140.82
135.63
133.02
130.27
126.86
126.68
126.62
125.97
125.81
118.56
116.06
109.64
104.36
99.72
65.13
65.00
63.75
42.19
HN
N
N
N
O
O
NH
O
NH
Cl
f1 (ppm)

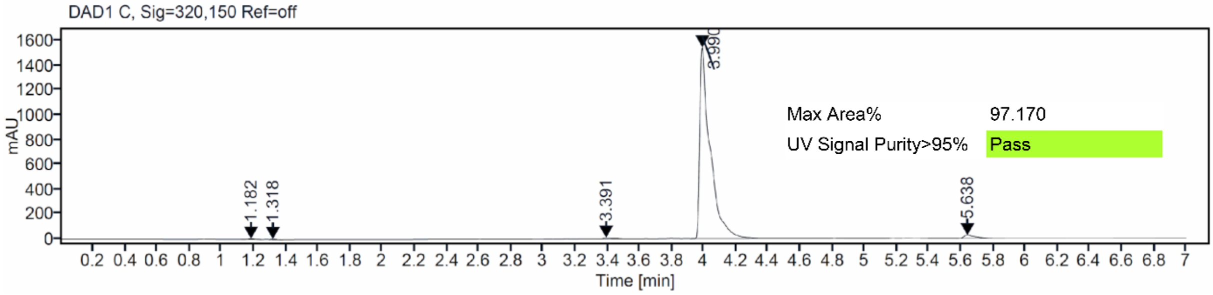
DAD1 C, Sig=320,150 Ref=off
mAU
Time [min]
1.182
1.318
3.391
3.990
5.638
Max Area% 97.170
UV Signal Purity>95% Pass

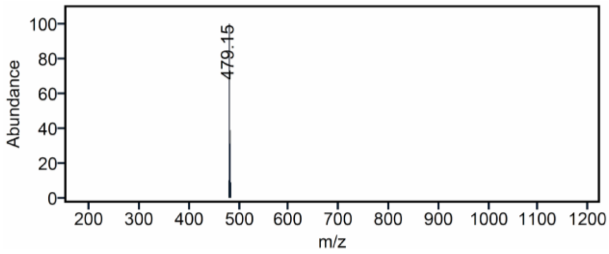
Abundance
m/z
479.15

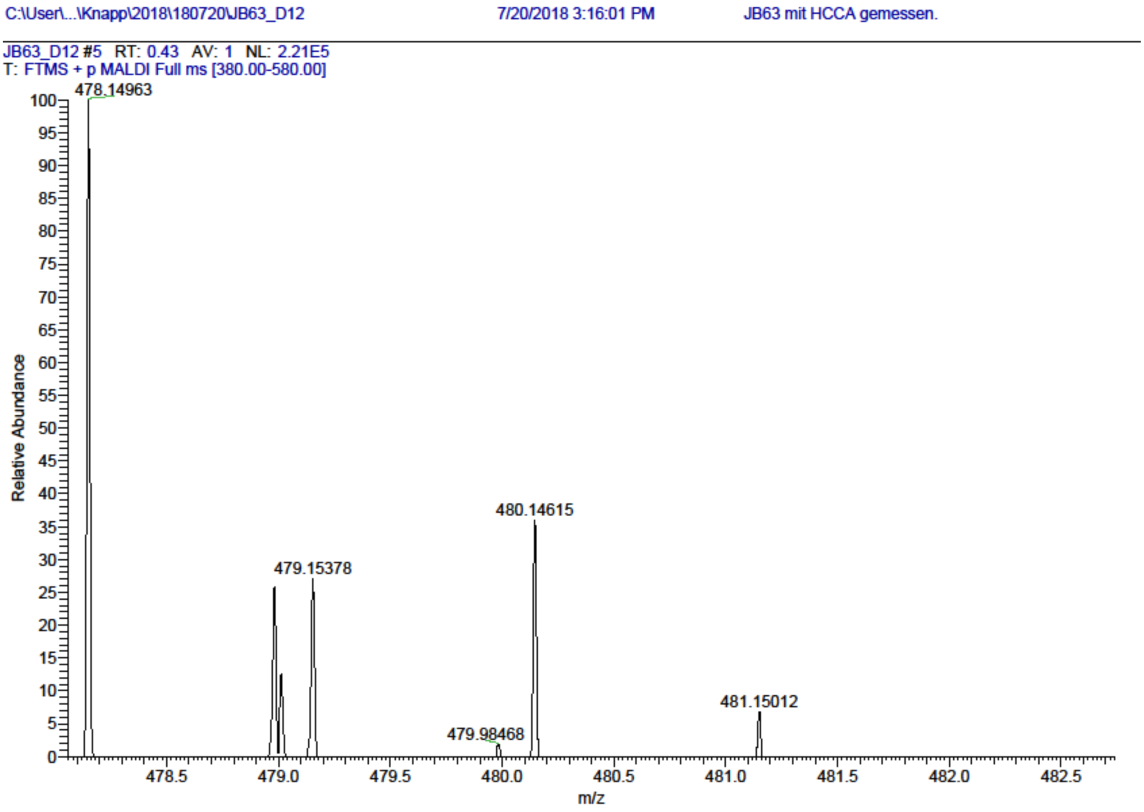

[1]H and [13]C {H} NMR, HPLC/MS (ESI) and HRMS data of compound **19**

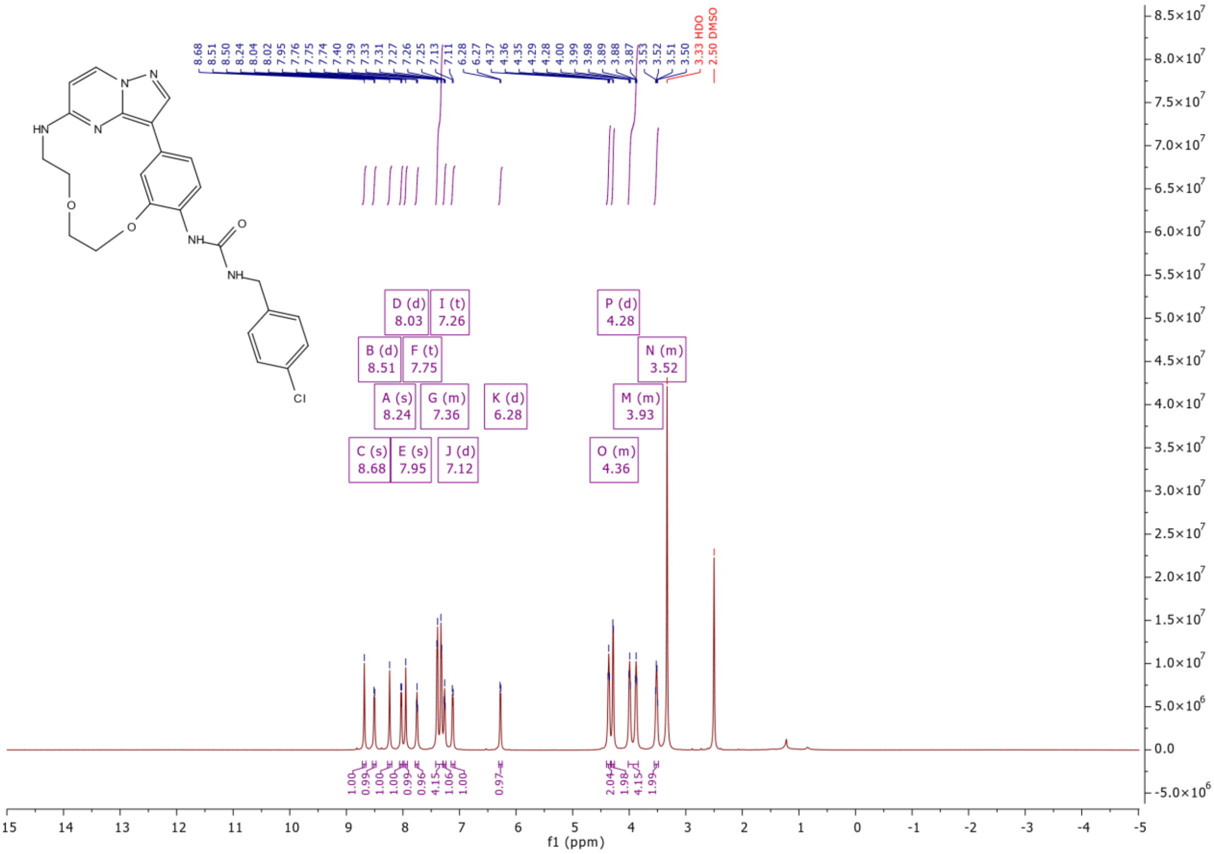

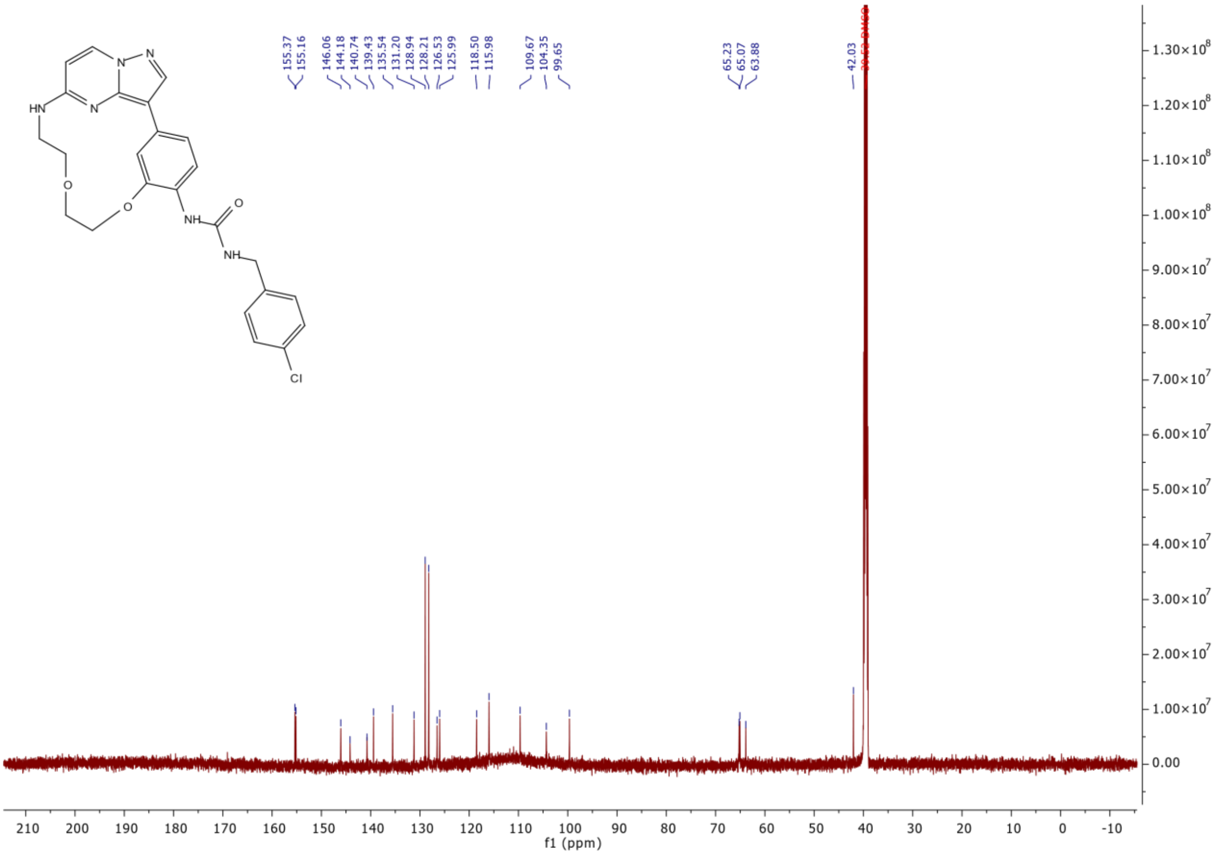
155.37
155.16
146.06
144.18
140.74
139.43
135.54
131.20
128.94
128.21
126.53
125.99
118.50
115.98
109.67
104.35
99.65
65.23
65.07
63.88
42.03
HN
N
N
N
O
O
NH
O
NH
Cl
f1 (ppm)

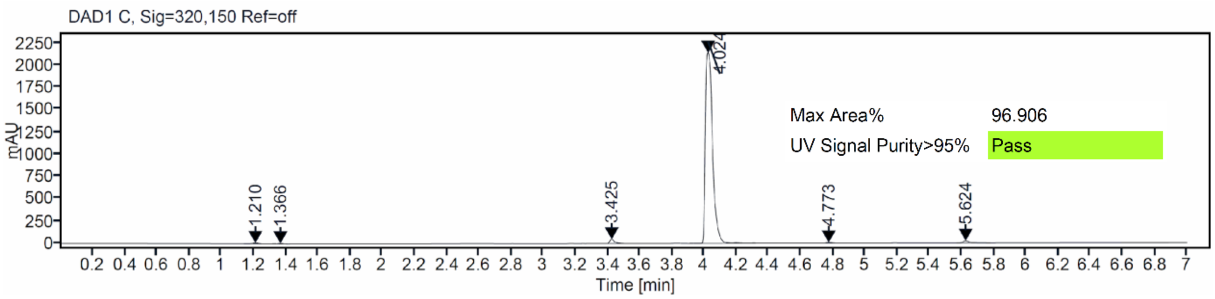
DAD1 C, Sig=320,150 Ref=off
mAU
1.210
1.366
3.425
4.024
4.773
5.624
Max Area%
96.906
UV Signal Purity>95%
Pass
Time [min]

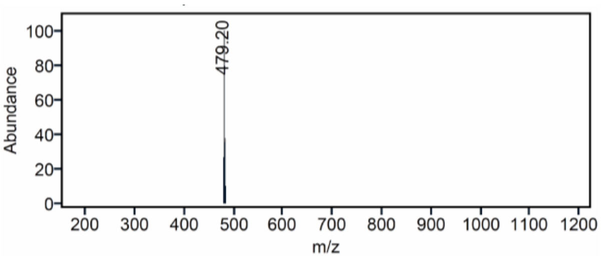
479.20
Abundance
m/z

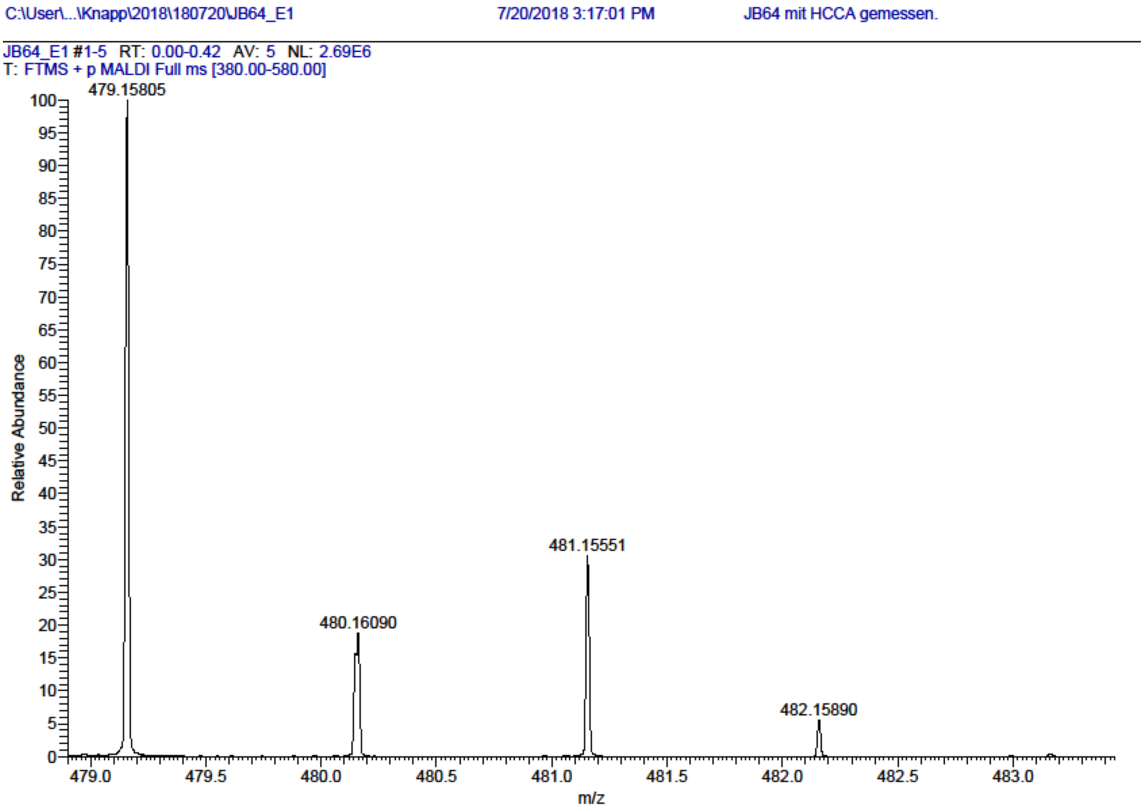

[1]H and [13]C {H} NMR, HPLC/MS (ESI) and HRMS data of compound **20**

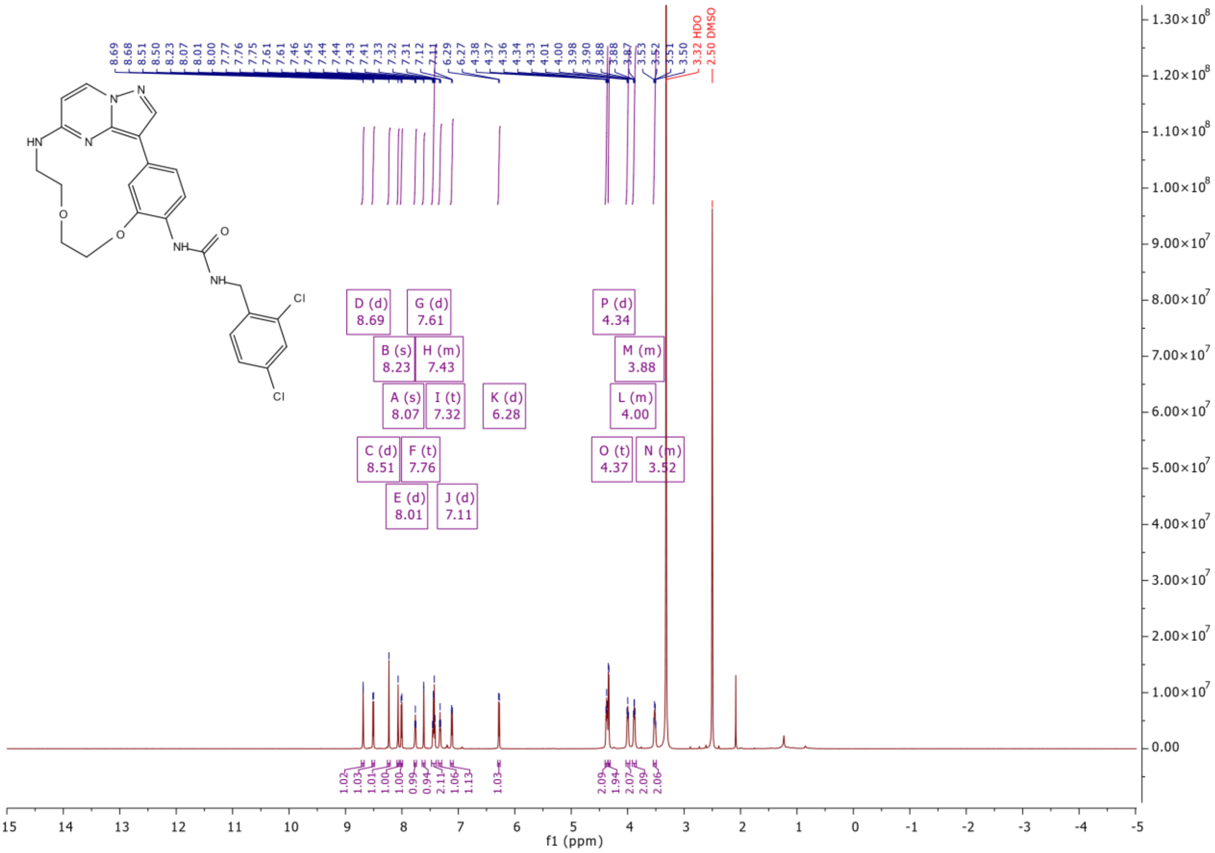

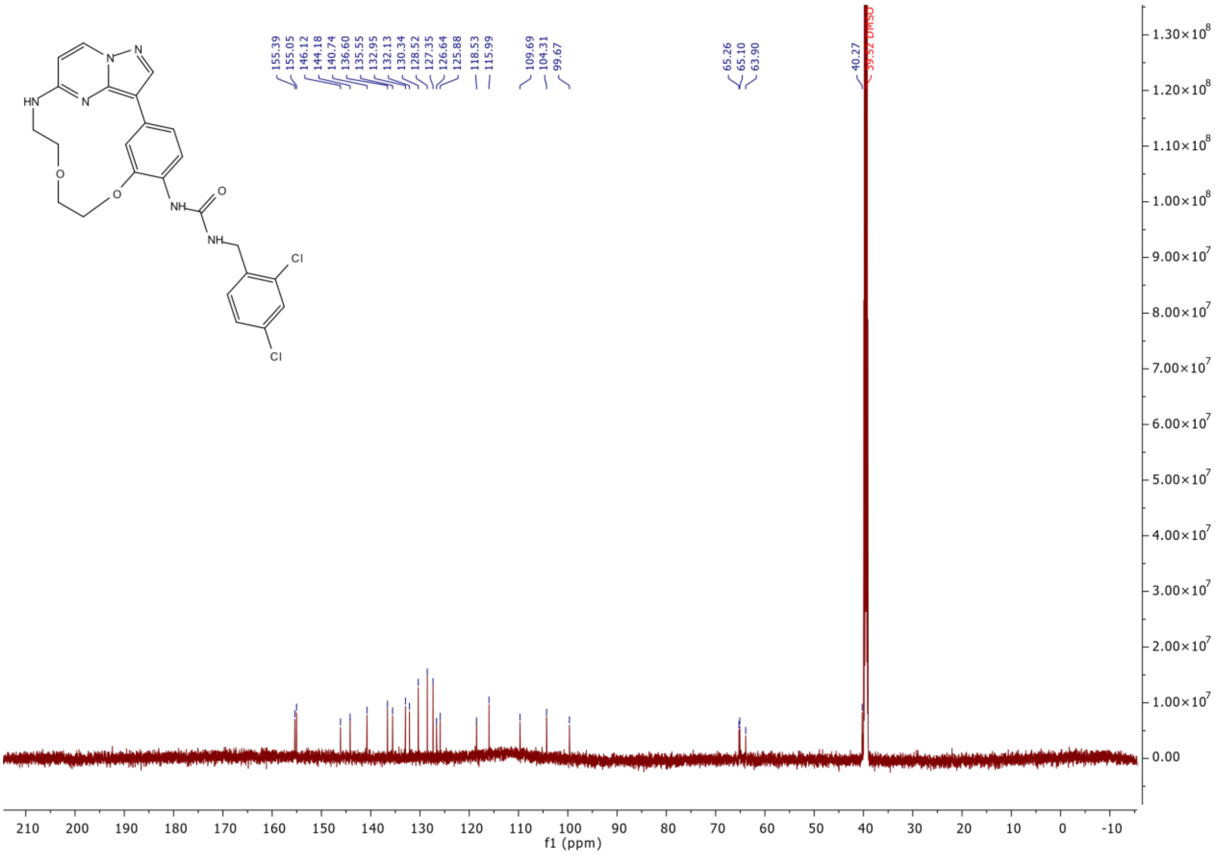

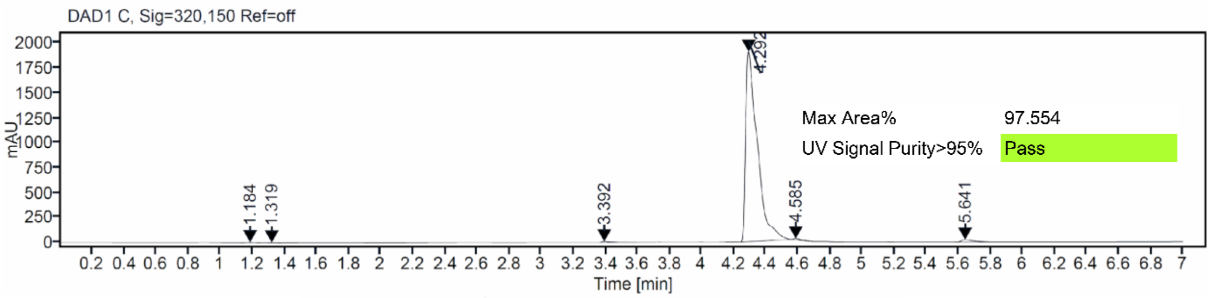
DAD1 C, Sig=320,150 Ref=off
Max Area% 97.554
UV Signal Purity>95% Pass
Time [min]
mAU

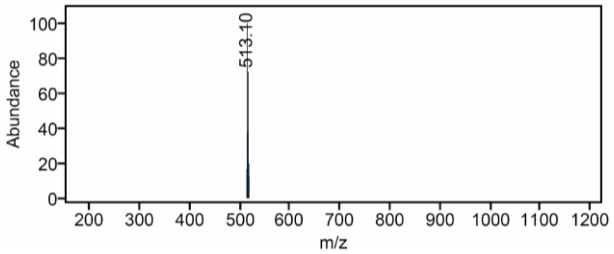
513.10
Abundance
m/z

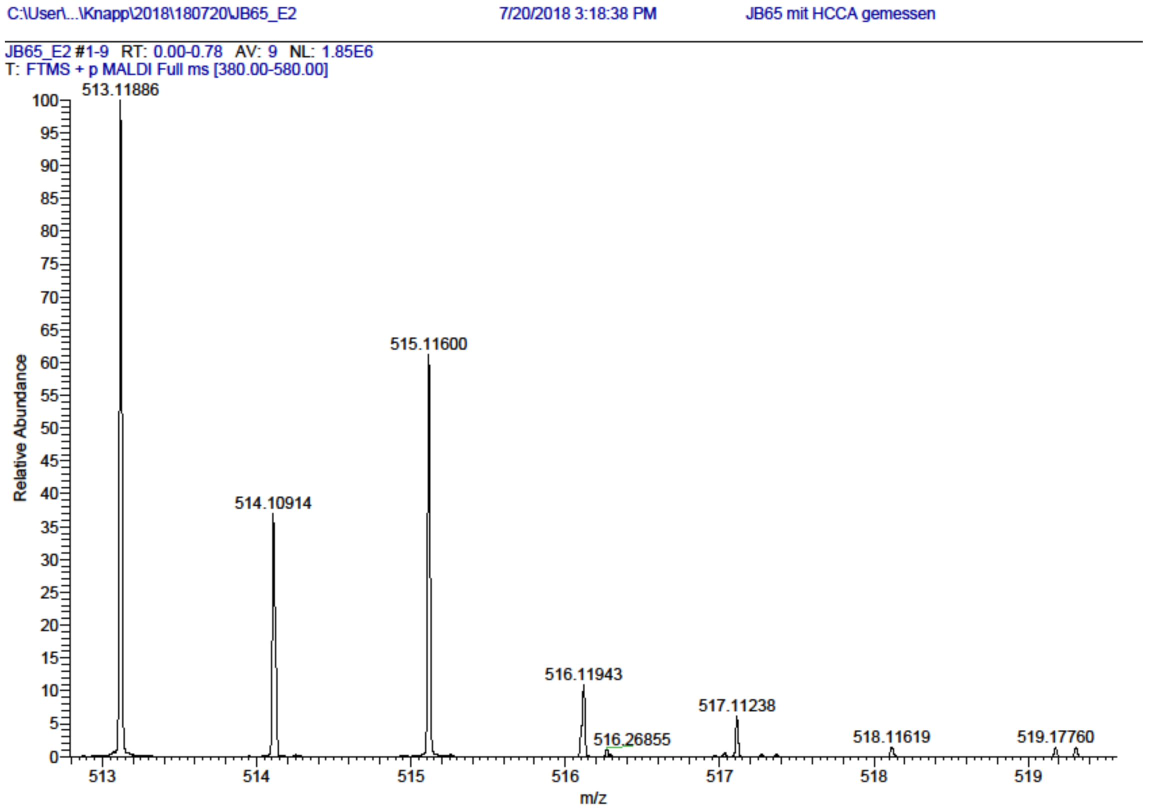

[1]H and [13]C {H} NMR, HPLC/MS (ESI) and HRMS data of compound **21**

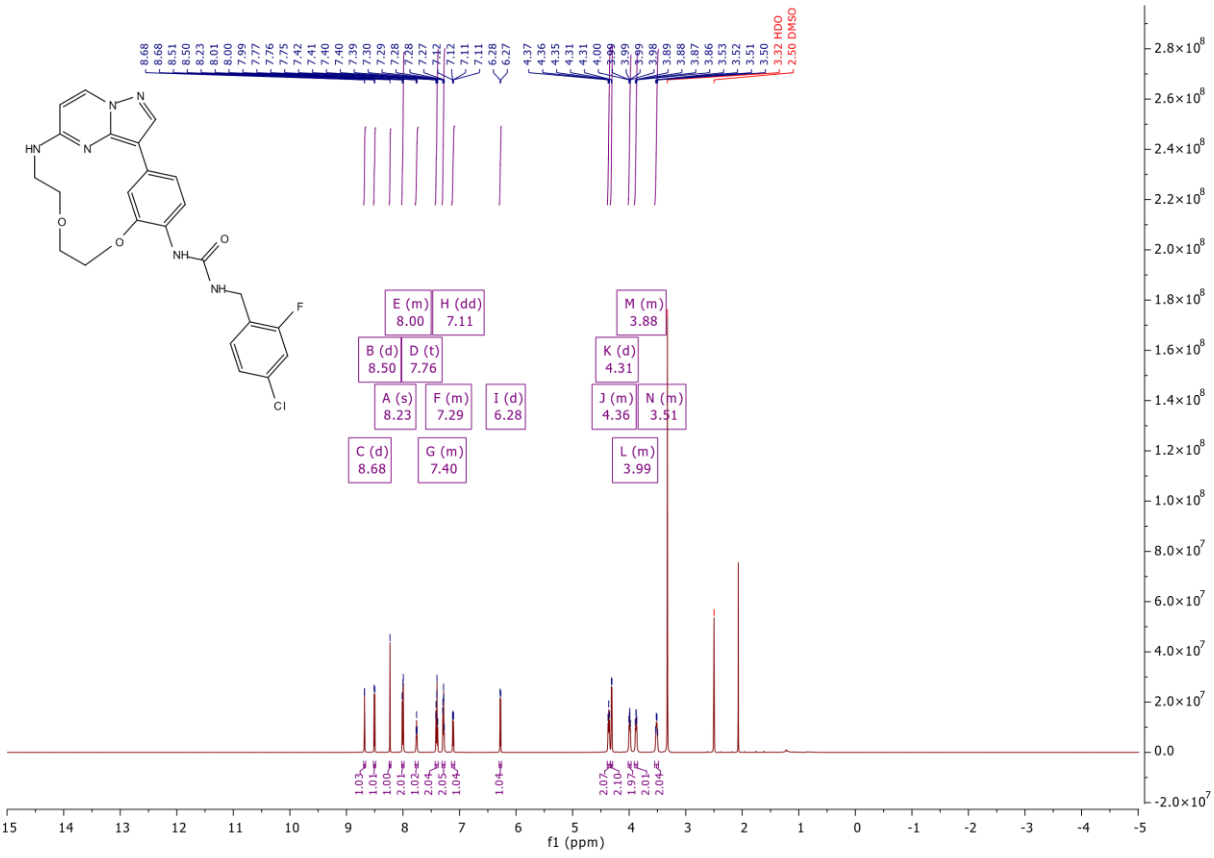

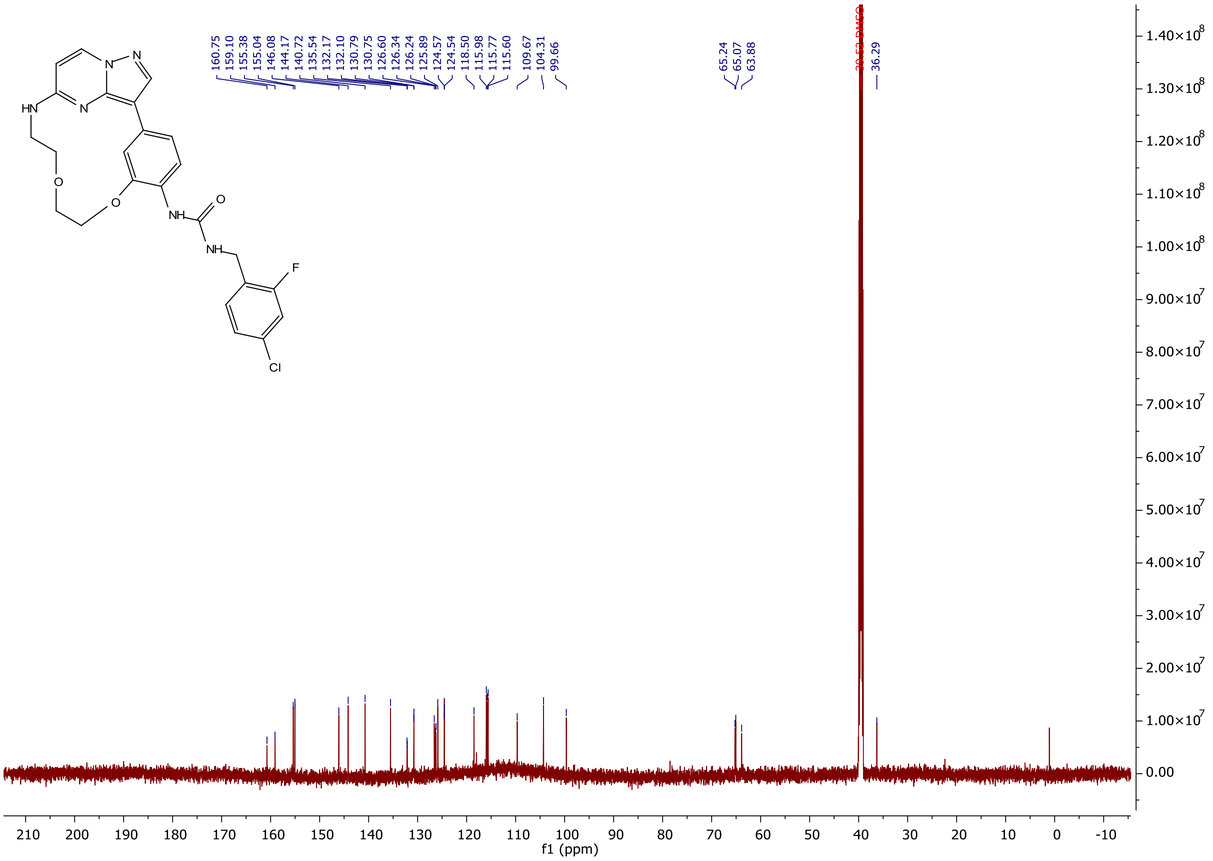

HN
N
N
N
O
O
NH
O
NH
F
Cl
160.75
159.10
155.38
155.04
146.08
144.17
140.72
135.54
132.17
132.10
130.79
130.75
126.60
126.34
126.24
125.89
124.57
124.54
118.50
115.98
115.77
115.60
109.67
104.31
99.66
65.24
65.07
63.88
36.29
f1 (ppm)

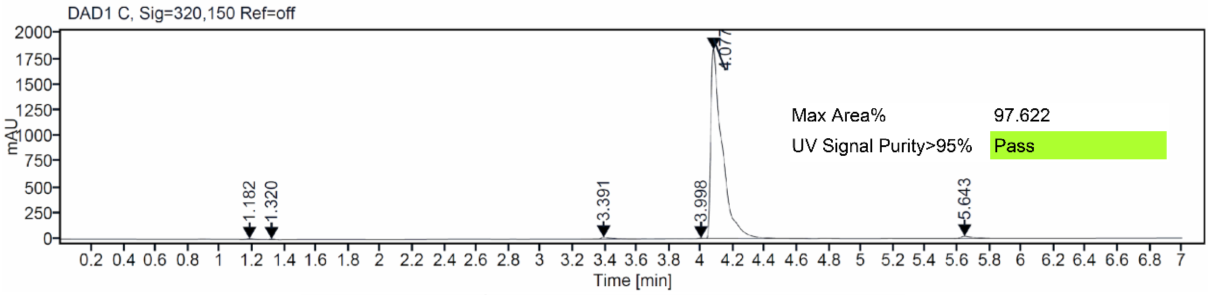

DAD1 C, Sig=320,150 Ref=off
mAU
Max Area% 97.622
UV Signal Purity>95% Pass
1.182
1.320
3.391
3.998
4.077
5.643
Time [min]

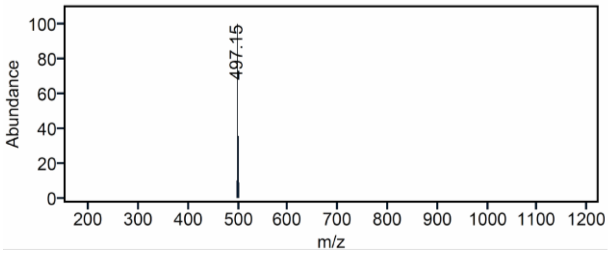

Abundance
497.15
m/z

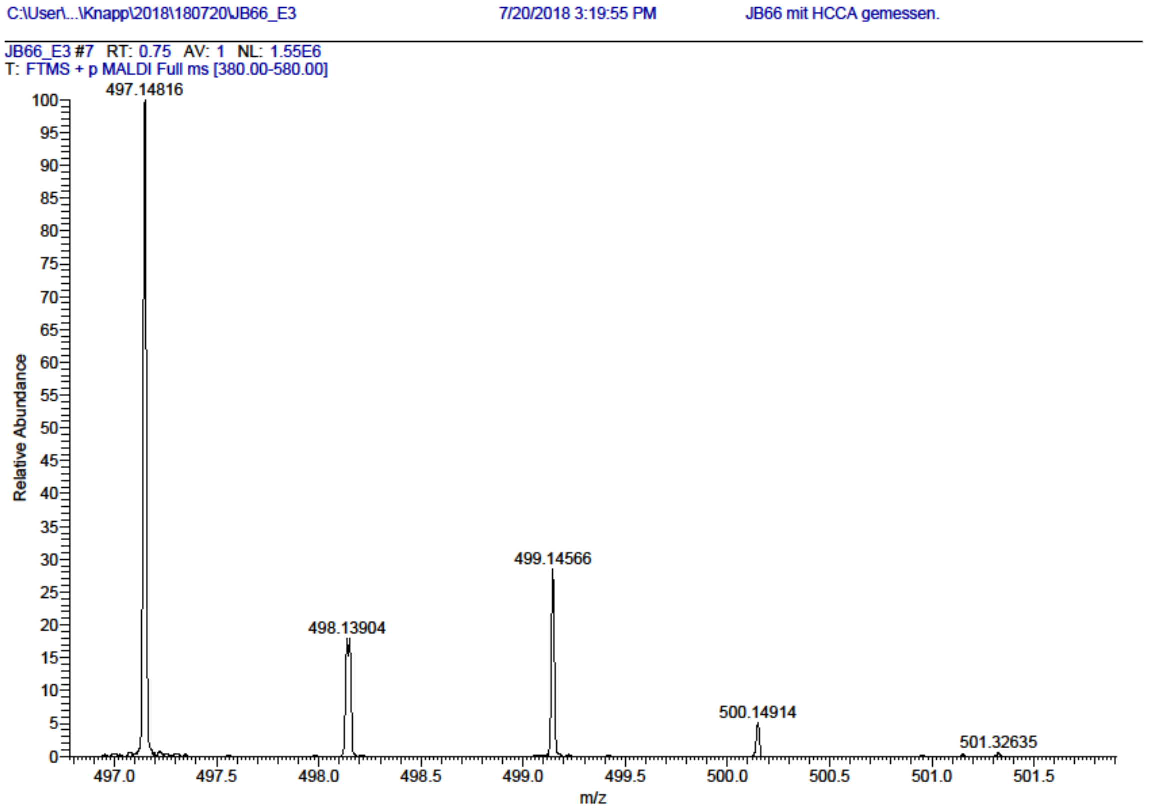

$^1H$ and $^{13}C$ {H} NMR, HPLC/MS (ESI) and HRMS data of compound **22**

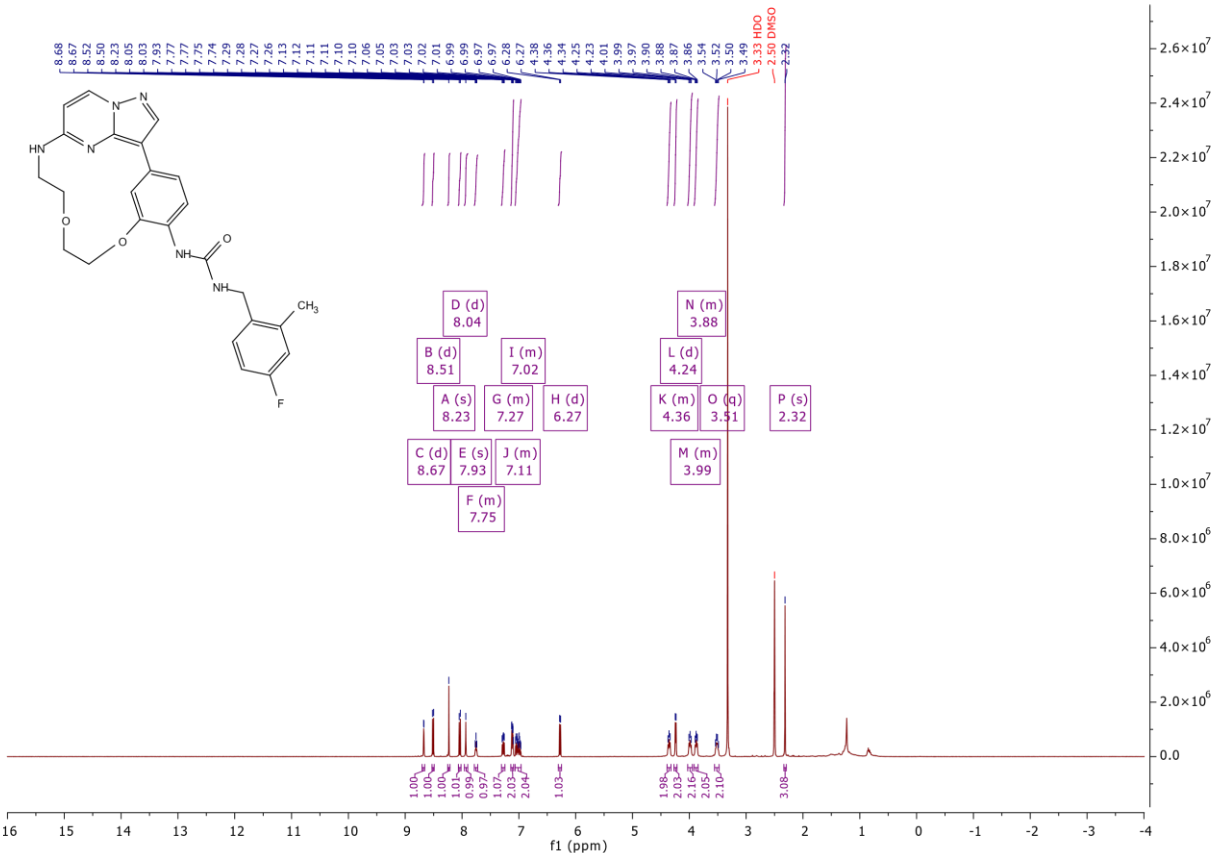

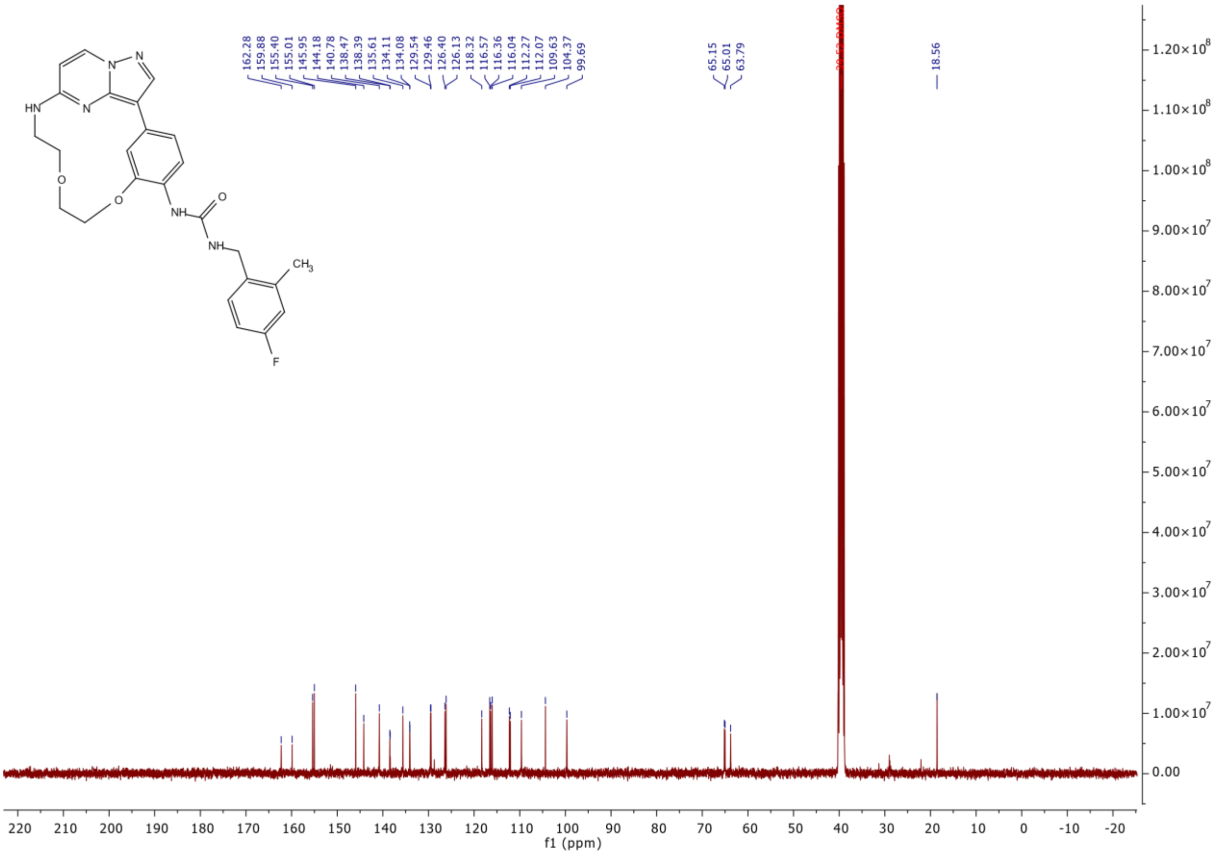

HN
N
N
N
O
O
NH
O
NH
CH3
F
162.28
159.88
155.40
155.01
145.95
144.18
140.78
138.47
138.39
135.61
134.11
134.08
129.54
129.46
126.40
126.13
118.32
116.57
116.36
116.04
112.27
112.07
109.63
104.37
99.69
65.15
65.01
63.79
18.56
f1 (ppm)

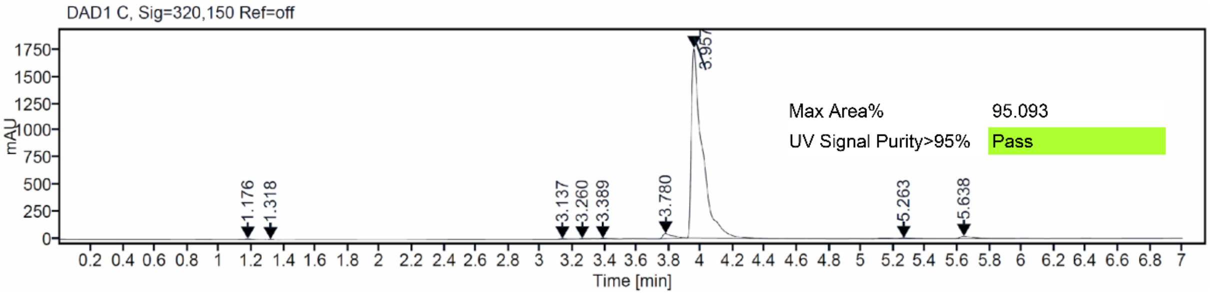

DAD1 C, Sig=320,150 Ref=off
mAU
Time [min]
1.176
1.318
3.137
3.260
3.389
3.780
3.957
5.263
5.638
Max Area% 95.093
UV Signal Purity>95% Pass

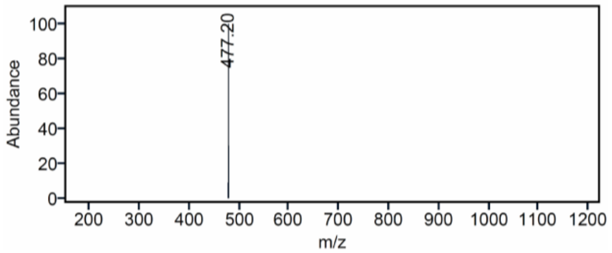

Abundance
477.20
m/z

### Mass Spectrum SmartFormula Report

**Analysis Info**

Acquisition Date 10/15/2024 4:21:57 PM

| | | | |
|---|---|---|---|
| Analysis Name | D:\Data\MS_Service\2024Q4\20241015_SKN_FG_000066-77\SKN_FG_000066_MD121_1.d | | |
| Method | Positive(100-1600)_Infusion_200uL-min.m | Operator | BDAL@DE |
| Sample Name | Test1 | Instrument | micrOTOF-Q 228888.10407 |
| Comment | | | |

**Acquisition Parameter**

| | | | | | |
|---|---|---|---|---|---|
| Source Type | ESI | Ion Polarity | Positive | Set Nebulizer | 1.8 Bar |
| Focus | Active | Set Capillary | 4000 V | Set Dry Heater | 280 °C |
| Scan Begin | 100 m/z | Set End Plate Offset | -500 V | Set Dry Gas | 8.0 l/min |
| Scan End | 1600 m/z | Set Collision Cell RF | 150.0 Vpp | Set Divert Valve | Waste |

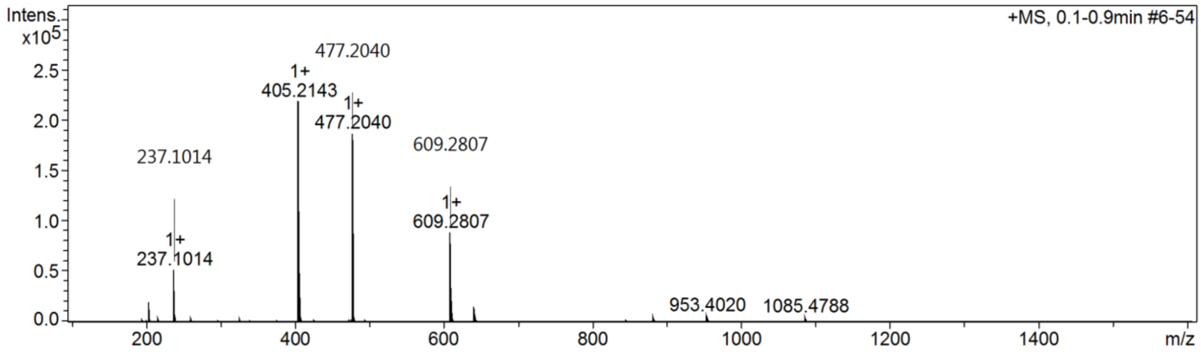

$^{1}H$ and $^{13}C$ {H} NMR, HPLC/MS (ESI) and HRMS data of compound **23**

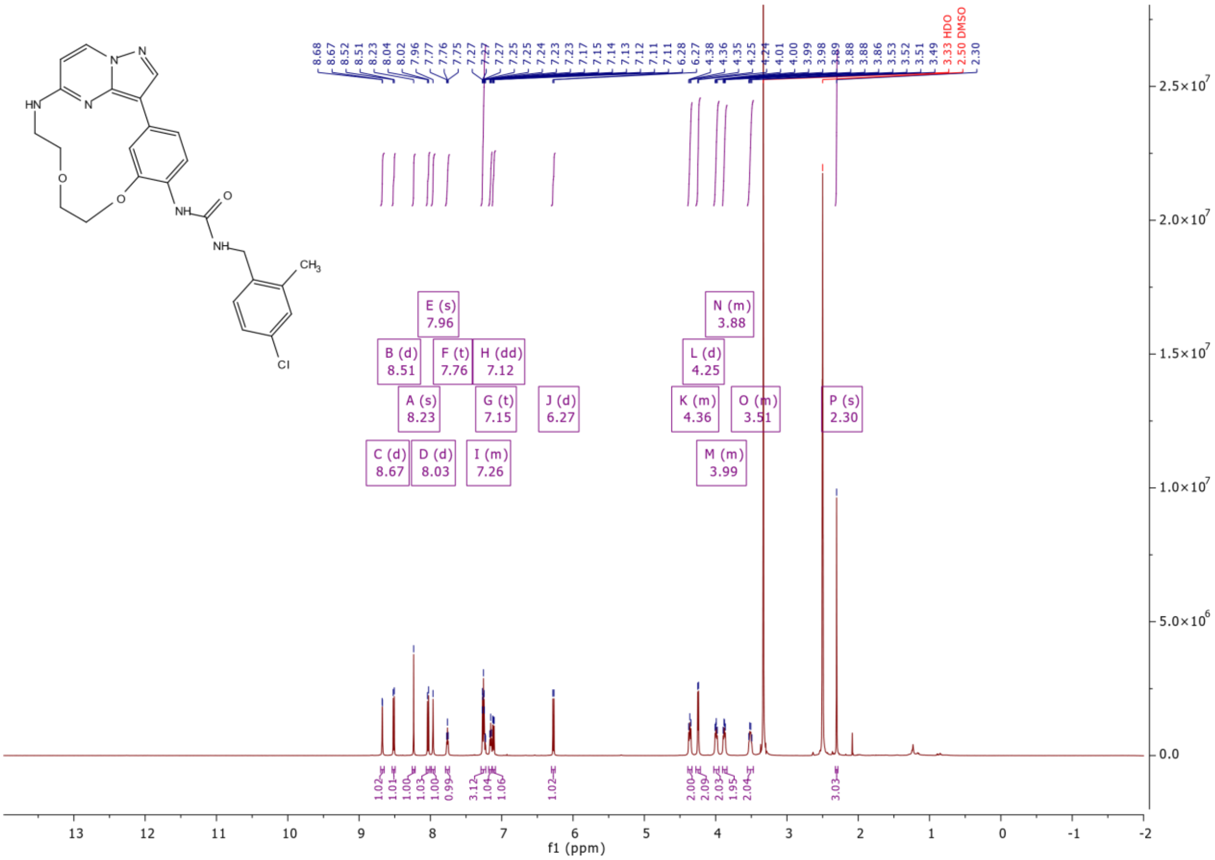

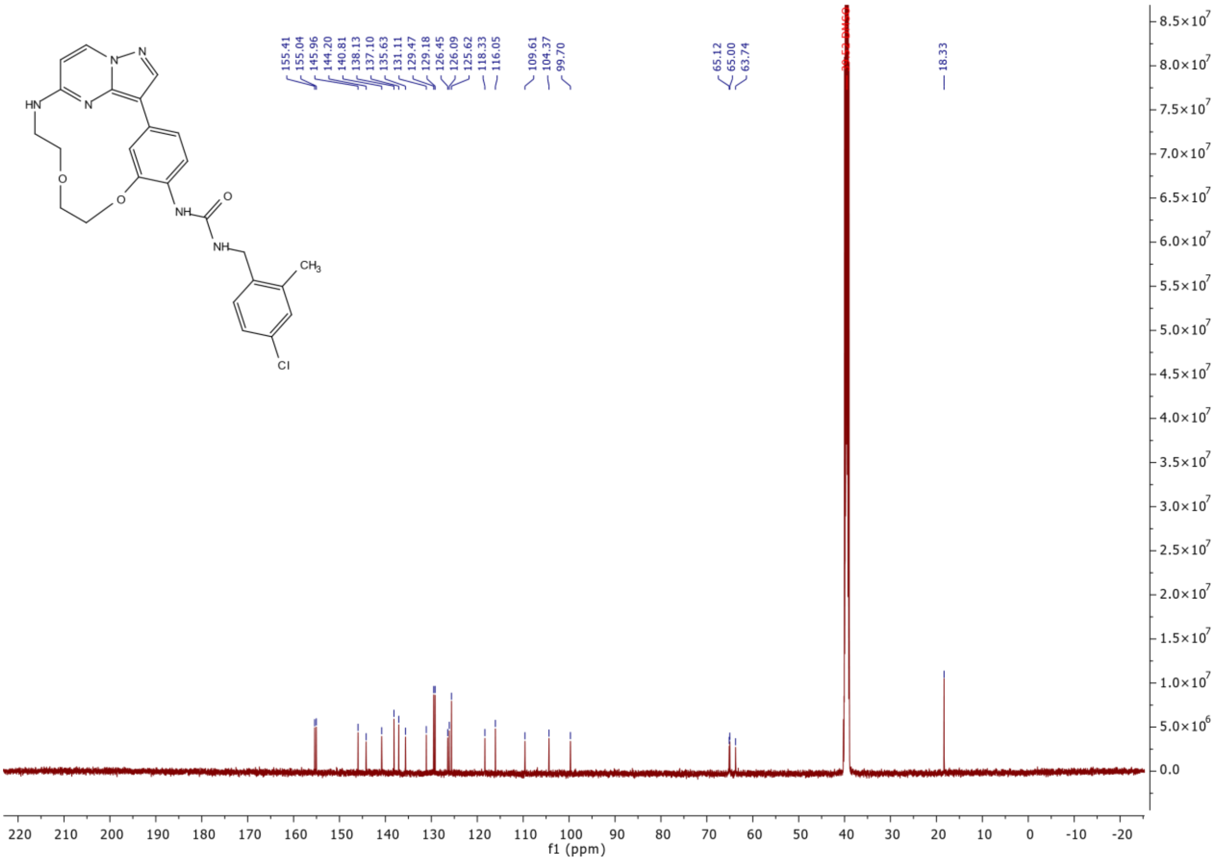
155.41
155.04
145.96
144.20
140.81
138.13
137.10
135.63
131.11
129.47
129.18
126.45
126.09
125.62
118.33
116.05
109.61
104.37
99.70
65.12
65.00
63.74
18.33
f1 (ppm)

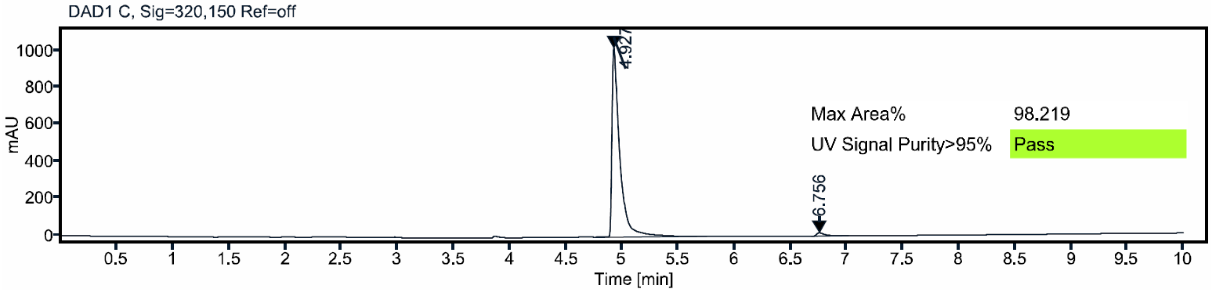
DAD1 C, Sig=320,150 Ref=off
mAU
4.927
6.756
Max Area% 98.219
UV Signal Purity>95% Pass
Time [min]

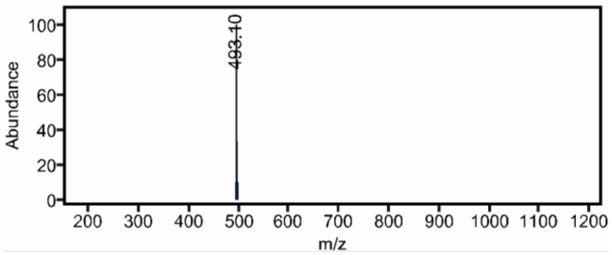
Abundance
493.10
m/z

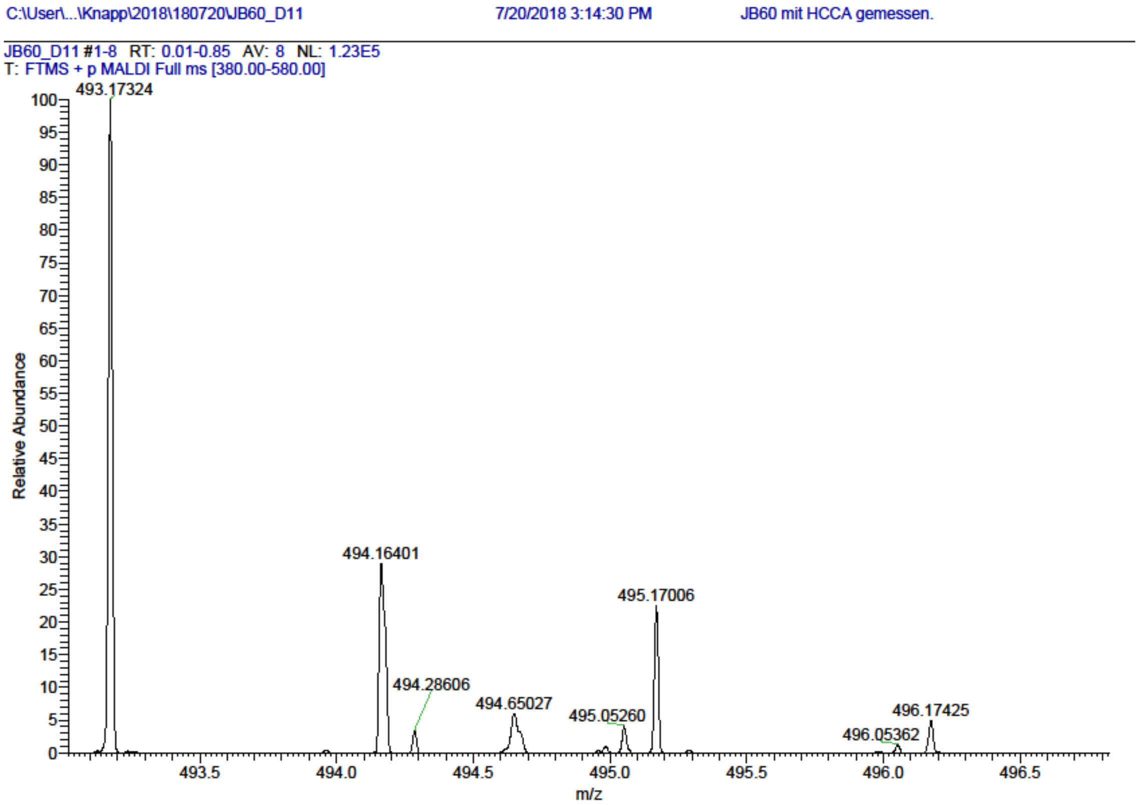

$^{1}H$ and $^{13}C$ {H} NMR, HPLC/MS (ESI) and HRMS data of compound **24**

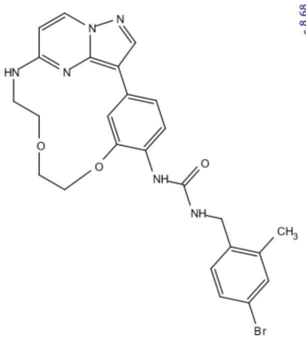

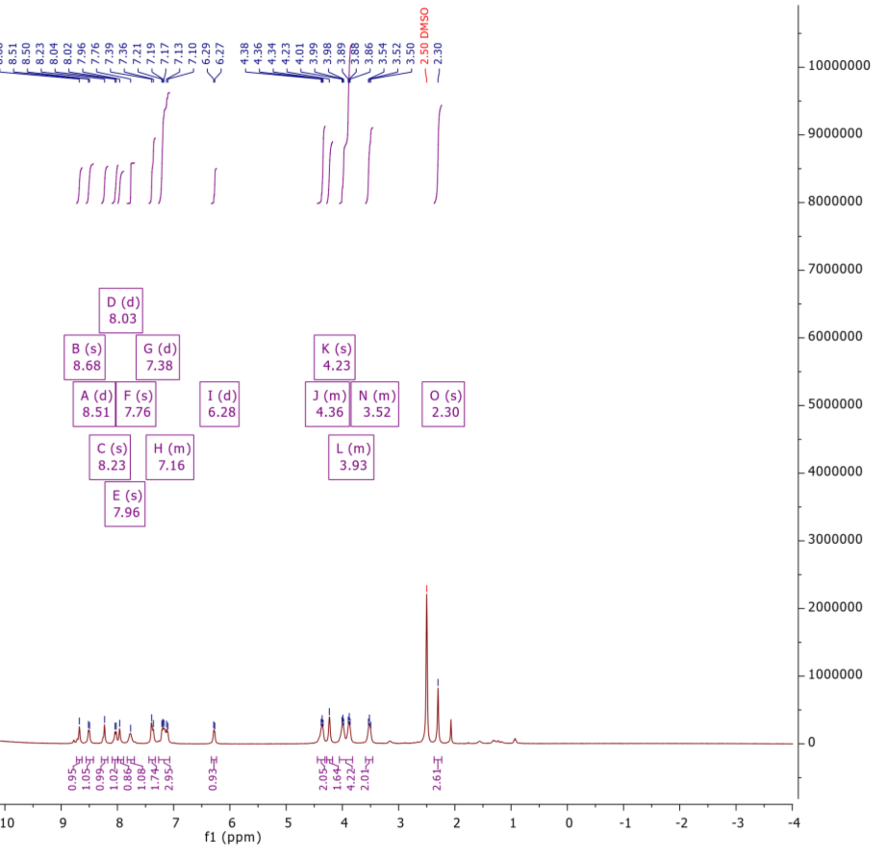

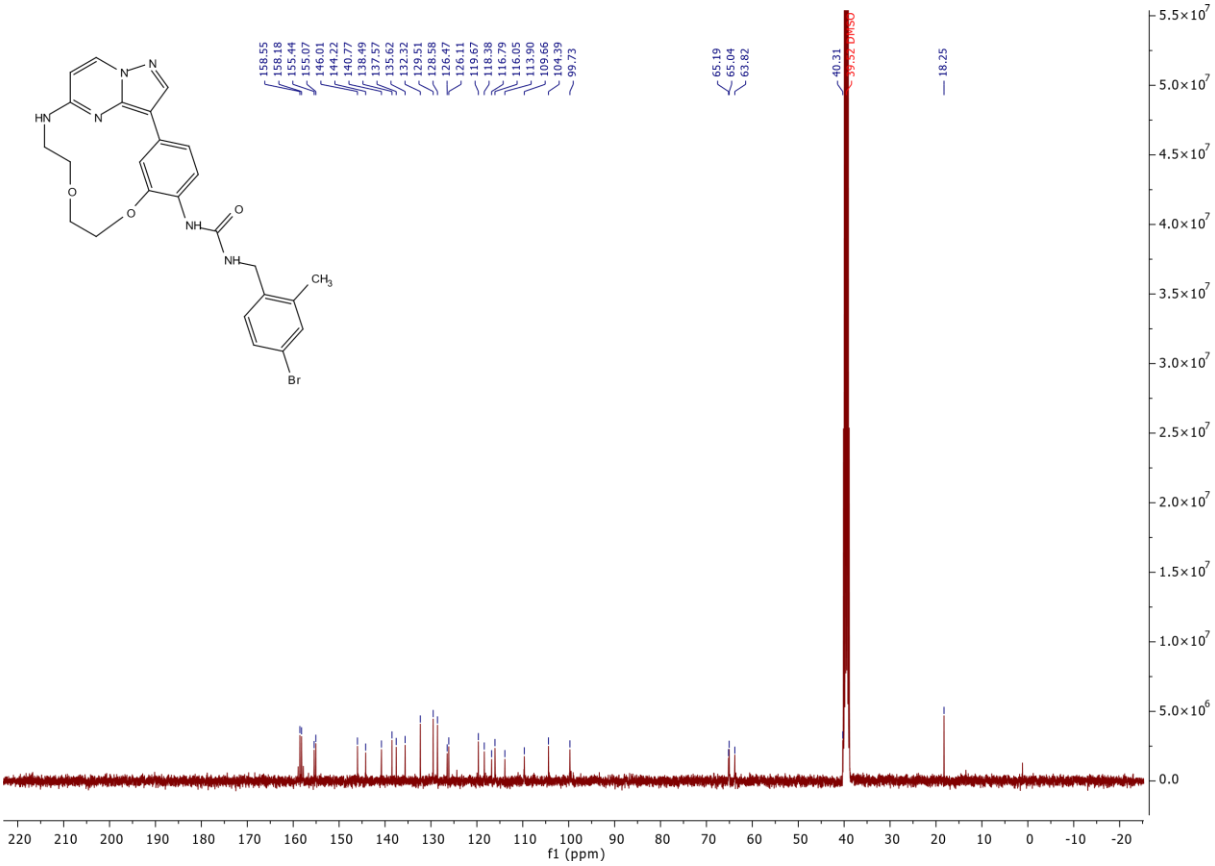

158.55
158.18
155.44
155.07
146.01
144.22
140.77
138.49
137.57
135.62
132.32
129.51
128.58
126.47
126.11
119.67
118.38
116.79
116.05
113.90
109.66
104.39
99.73
65.19
65.04
63.82
40.31
39.52 DMSO
18.25
f1 (ppm)

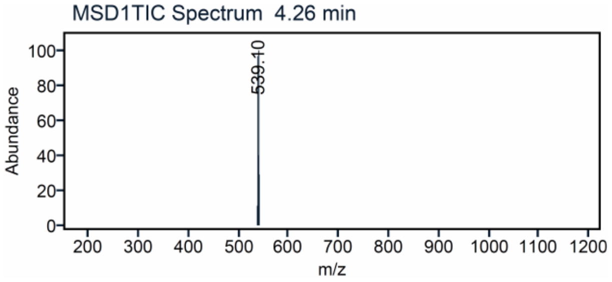

MSD1TIC Spectrum 4.26 min
539.10
Abundance
m/z

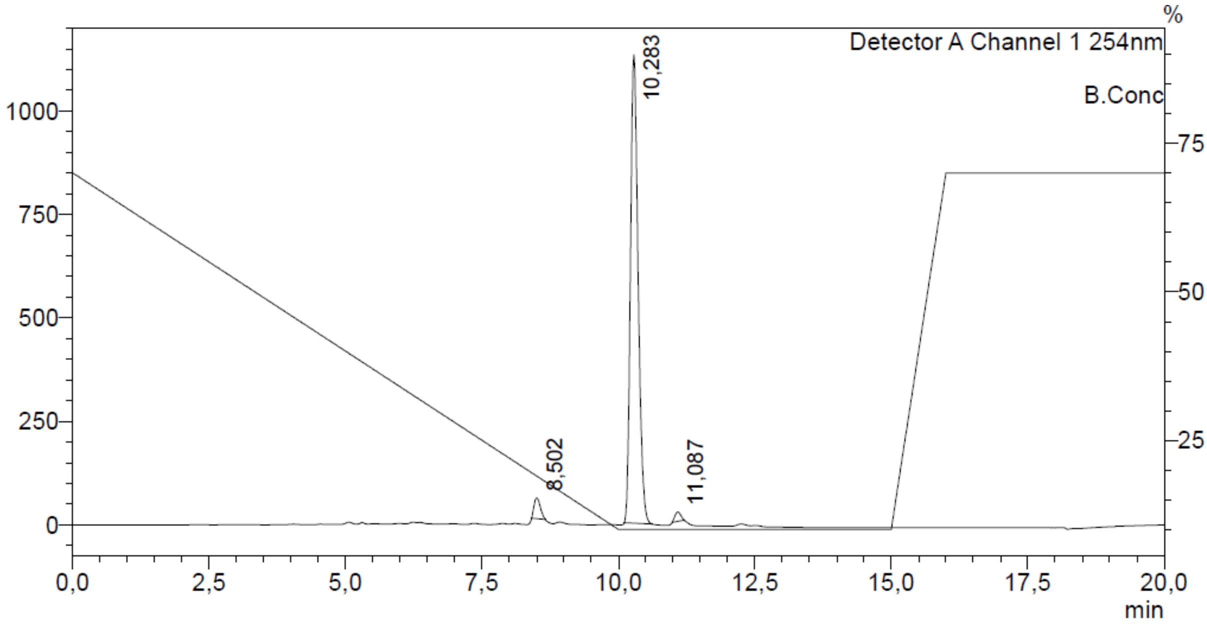

Detector A Channel 1 254nm
B.Conc
%
10,283
8,502
11,087
min

### Mass Spectrum SmartFormula Report

**Analysis Info**

| | | | |
|---|---|---|---|
| Analysis Name | D:\Data\MS_Service\2024Q4\20241015_SKN_FG_000066-77\SKN_FG_000077_TML79_1.d | Acquisition Date | 10/15/2024 6:00:29 PM |
| Method | Positive(100-1600)_Infusion_200uL-min.m | Operator | BDAL@DE |
| Sample Name | Test1 | Instrument | micrOTOF-Q 228888.10407 |
| Comment | | | |

**Acquisition Parameter**

| | | | | | |
|---|---|---|---|---|---|
| Source Type | ESI | Ion Polarity | Positive | Set Nebulizer | 1.8 Bar |
| Focus | Active | Set Capillary | 4000 V | Set Dry Heater | 280 °C |
| Scan Begin | 100 m/z | Set End Plate Offset | -500 V | Set Dry Gas | 8.0 l/min |
| Scan End | 1600 m/z | Set Collision Cell RF | 150.0 Vpp | Set Divert Valve | Waste |

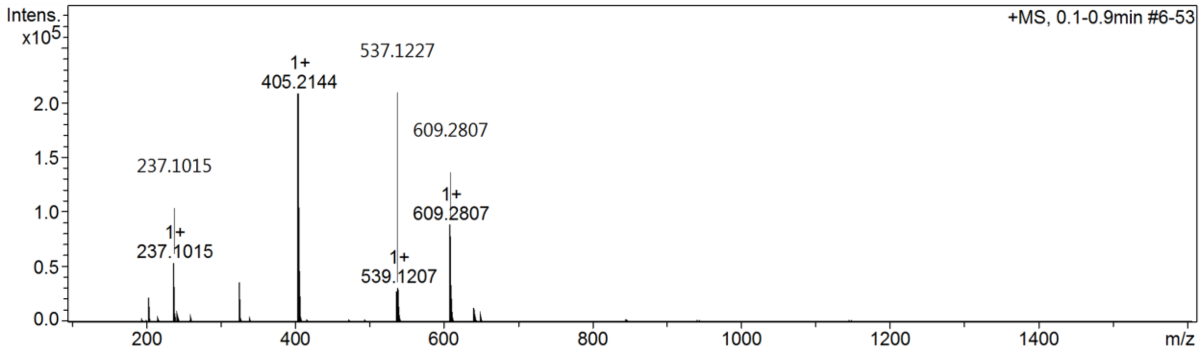

$^{1}H$ and $^{13}C$ {H} NMR, HPLC/MS (ESI) and HRMS data of compound **25**

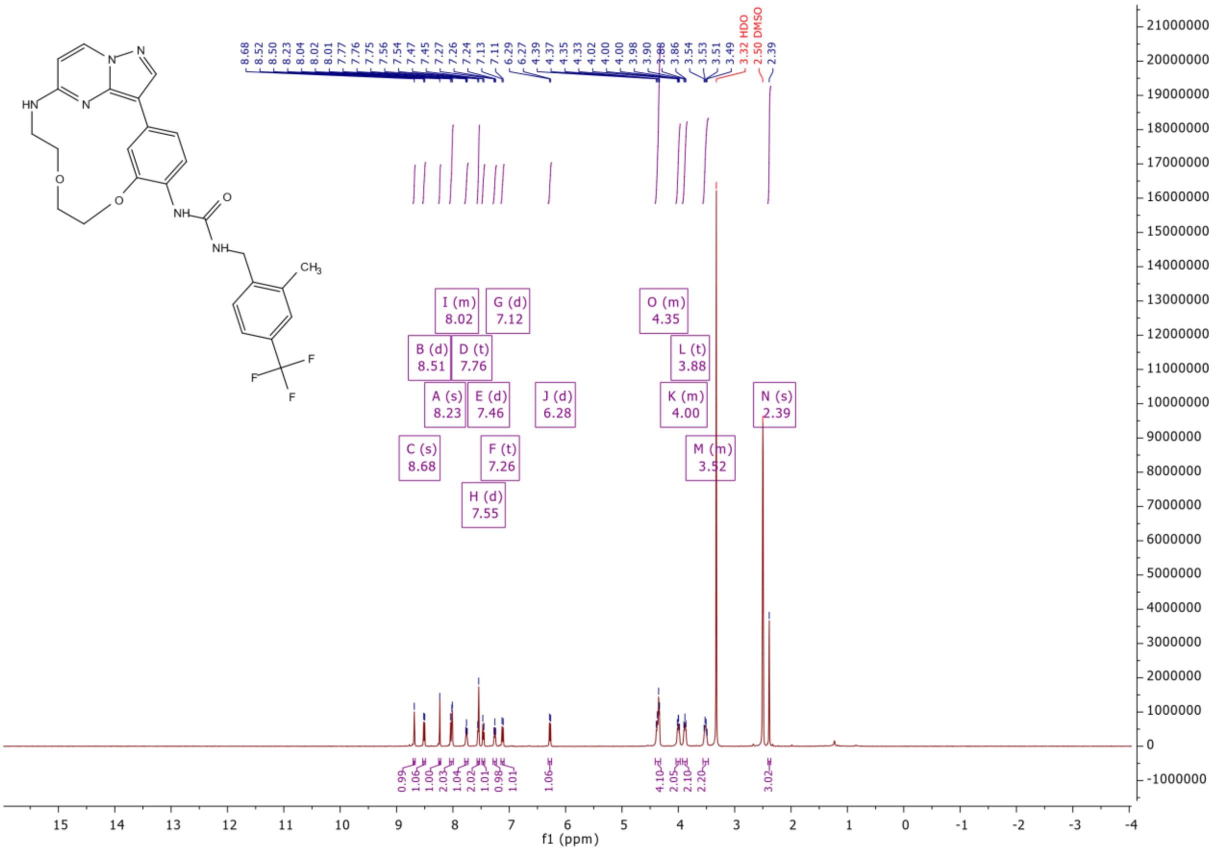

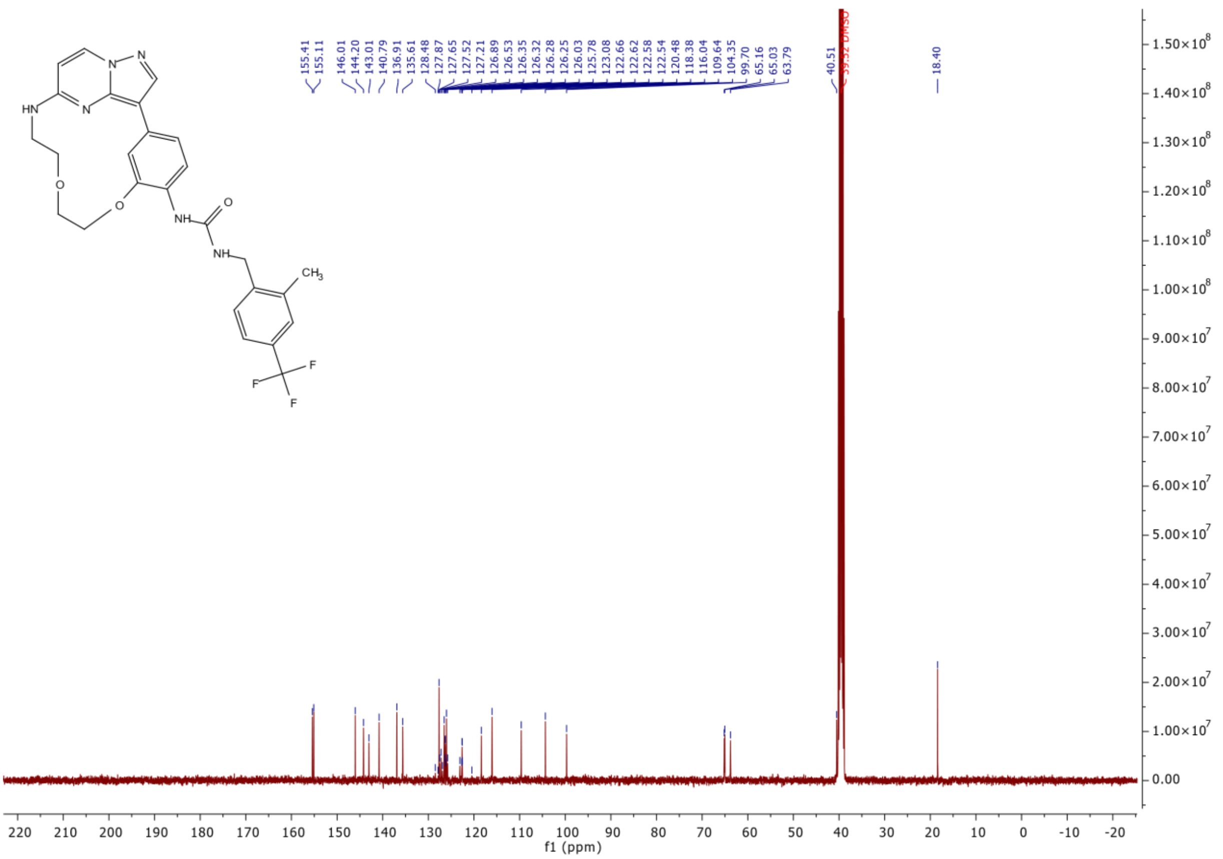
155.41
155.11
146.01
144.20
143.01
140.79
136.91
135.61
128.48
127.87
127.65
127.52
127.21
126.89
126.53
126.35
126.32
126.28
126.25
126.03
125.78
123.08
122.66
122.62
122.58
122.54
120.48
118.38
116.04
109.64
104.35
99.70
65.16
65.03
63.79
40.51
39.52 DMSO
18.40
f1 (ppm)

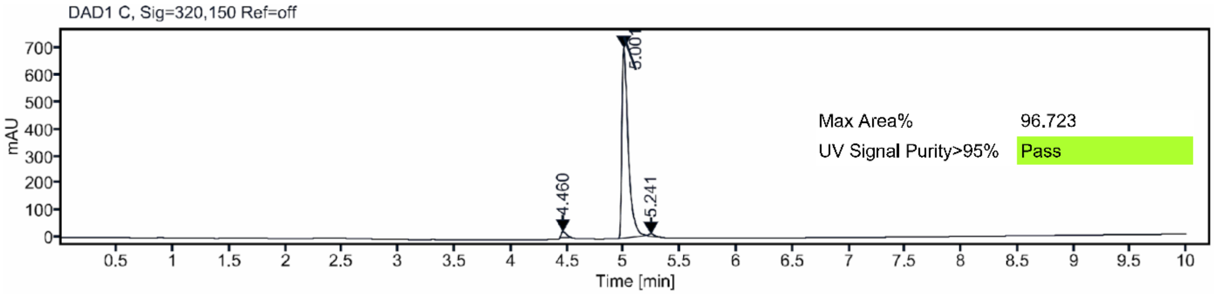
DAD1 C, Sig=320,150 Ref=off
mAU
Time [min]
5.001
4.460
5.241
Max Area%
96.723
UV Signal Purity>95%
Pass

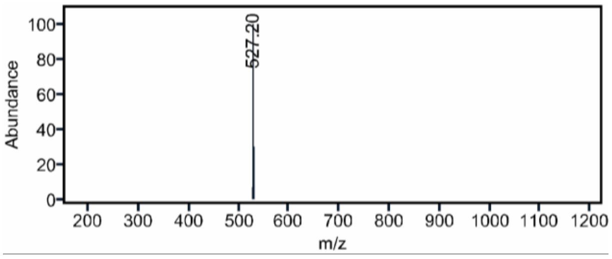
Abundance
m/z
527.20

### Mass Spectrum SmartFormula Report

**Analysis Info**

| | | | |
|---|---|---|---|
| Analysis Name | D:\Data\MS_Service\2024Q4\20241015_SKN_FG_000066-77\SKN_FG_000067_MD123_1.d | Acquisition Date | 10/15/2024 4:26:56 PM |
| Method | Positive(100-1600)_Infusion_200uL-min.m | Operator | BDAL@DE |
| Sample Name | Test1 | Instrument | micrOTOF-Q 228888.10407 |
| Comment | | | |

**Acquisition Parameter**

| | | | | | |
|---|---|---|---|---|---|
| Source Type | ESI | Ion Polarity | Positive | Set Nebulizer | 1.8 Bar |
| Focus | Active | Set Capillary | 4000 V | Set Dry Heater | 280 °C |
| Scan Begin | 100 m/z | Set End Plate Offset | -500 V | Set Dry Gas | 8.0 l/min |
| Scan End | 1600 m/z | Set Collision Cell RF | 150.0 Vpp | Set Divert Valve | Waste |

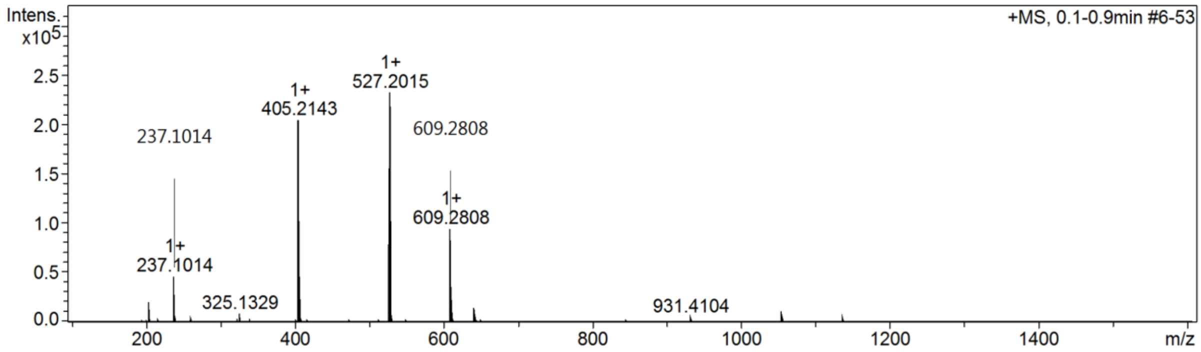

$^1$H and $^{13}$C {H} NMR, HPLC/MS (ESI) and HRMS data of compound **26**

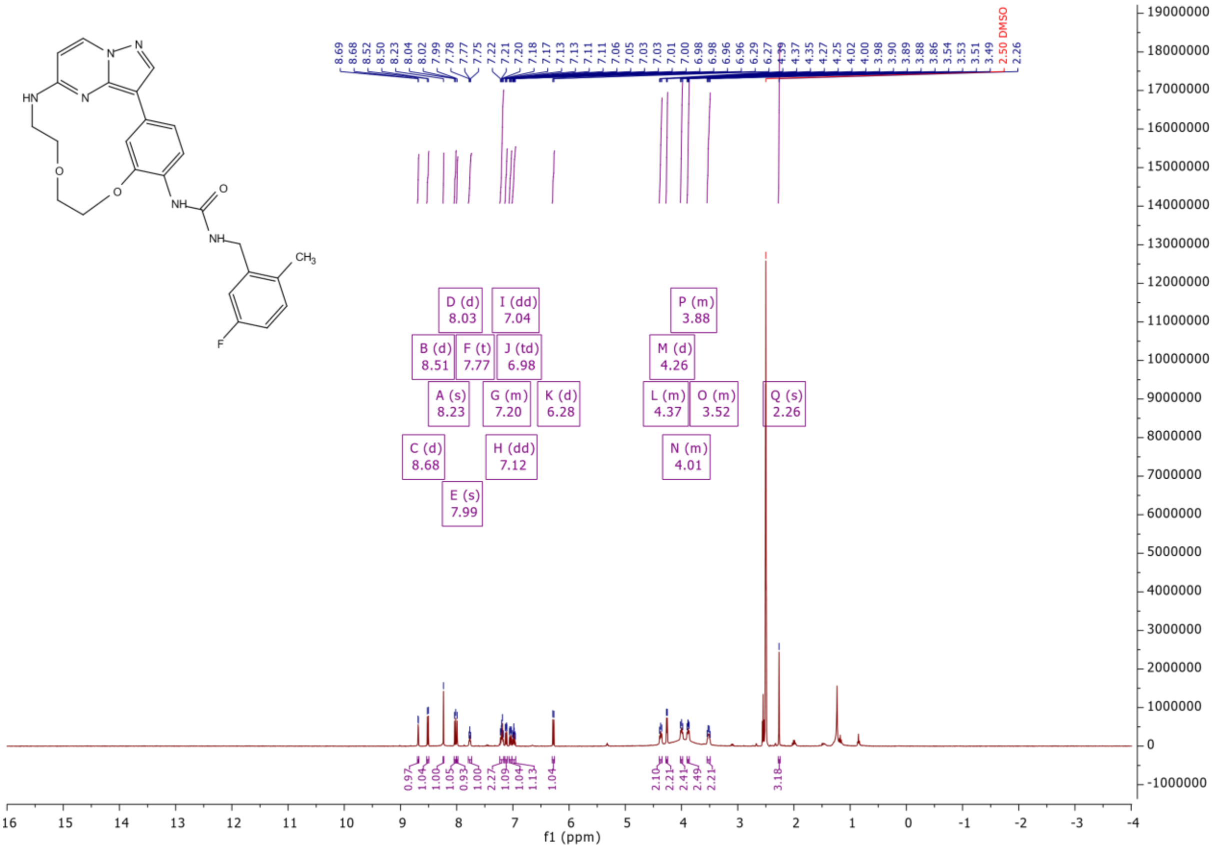

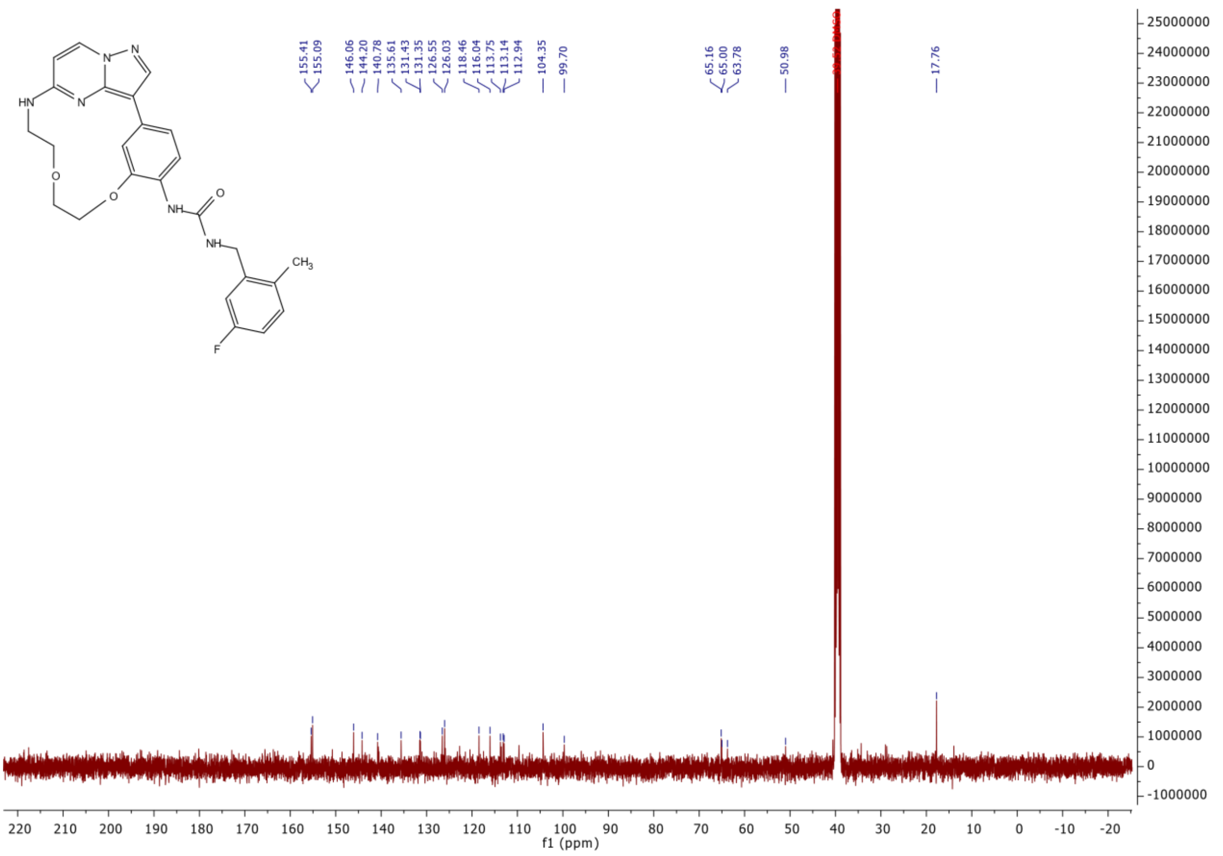
HN
N
N
N
O
O
NH
O
NH
CH3
F
155.41
155.09
146.06
144.20
140.78
135.61
131.43
131.35
126.55
126.03
118.46
116.04
113.75
113.14
112.94
104.35
99.70
65.16
65.00
63.78
50.98
17.76
f1 (ppm)

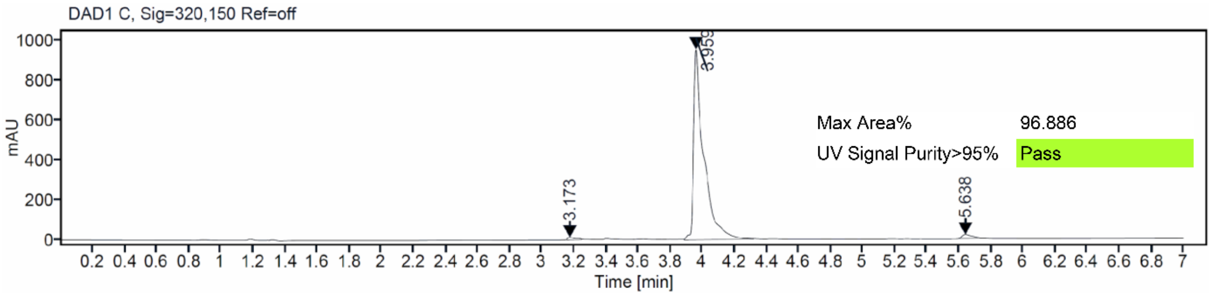
DAD1 C, Sig=320,150 Ref=off
mAU
3.173
3.959
5.638
Max Area% 96.886
UV Signal Purity>95% Pass
Time [min]

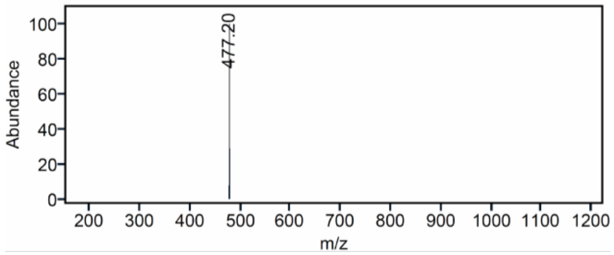
Abundance
477.20
m/z

### Mass Spectrum SmartFormula Report

**Analysis Info**

| | | | |
|---|---|---|---|
| Analysis Name | D:\Data\MS_Service\2024Q4\20241015_SKN_FG_000066-77\SKN_FG_000076_TML78_1.d | Acquisition Date | 10/15/2024 5:55:58 PM |
| Method | Positive(100-1600)_Infusion_200uL-min.m | Operator | BDAL@DE |
| Sample Name | Test1 | Instrument | micrOTOF-Q 228888.10407 |
| Comment | | | |

**Acquisition Parameter**

| | | | | | |
|---|---|---|---|---|---|
| Source Type | ESI | Ion Polarity | Positive | Set Nebulizer | 1.8 Bar |
| Focus | Active | Set Capillary | 4000 V | Set Dry Heater | 280 °C |
| Scan Begin | 100 m/z | Set End Plate Offset | -500 V | Set Dry Gas | 8.0 l/min |
| Scan End | 1600 m/z | Set Collision Cell RF | 150.0 Vpp | Set Divert Valve | Waste |

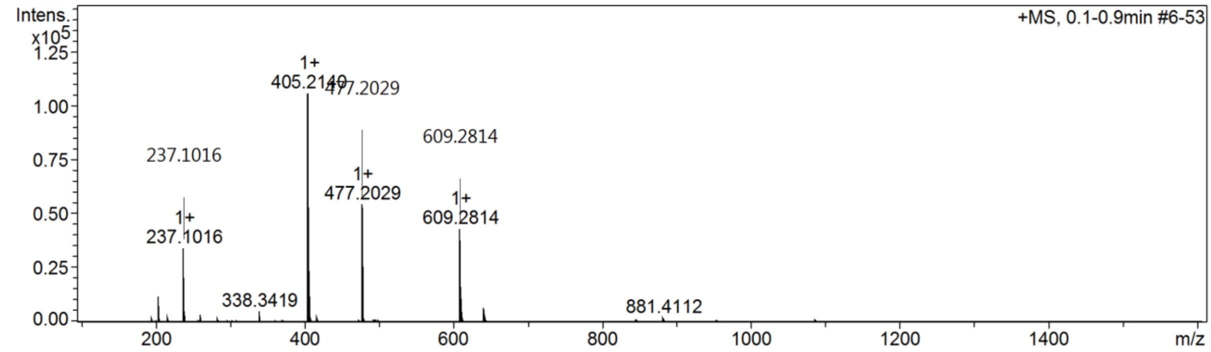

$^1$H and $^{13}$C {H} NMR, HPLC/MS (ESI) and HRMS data of compound **27**

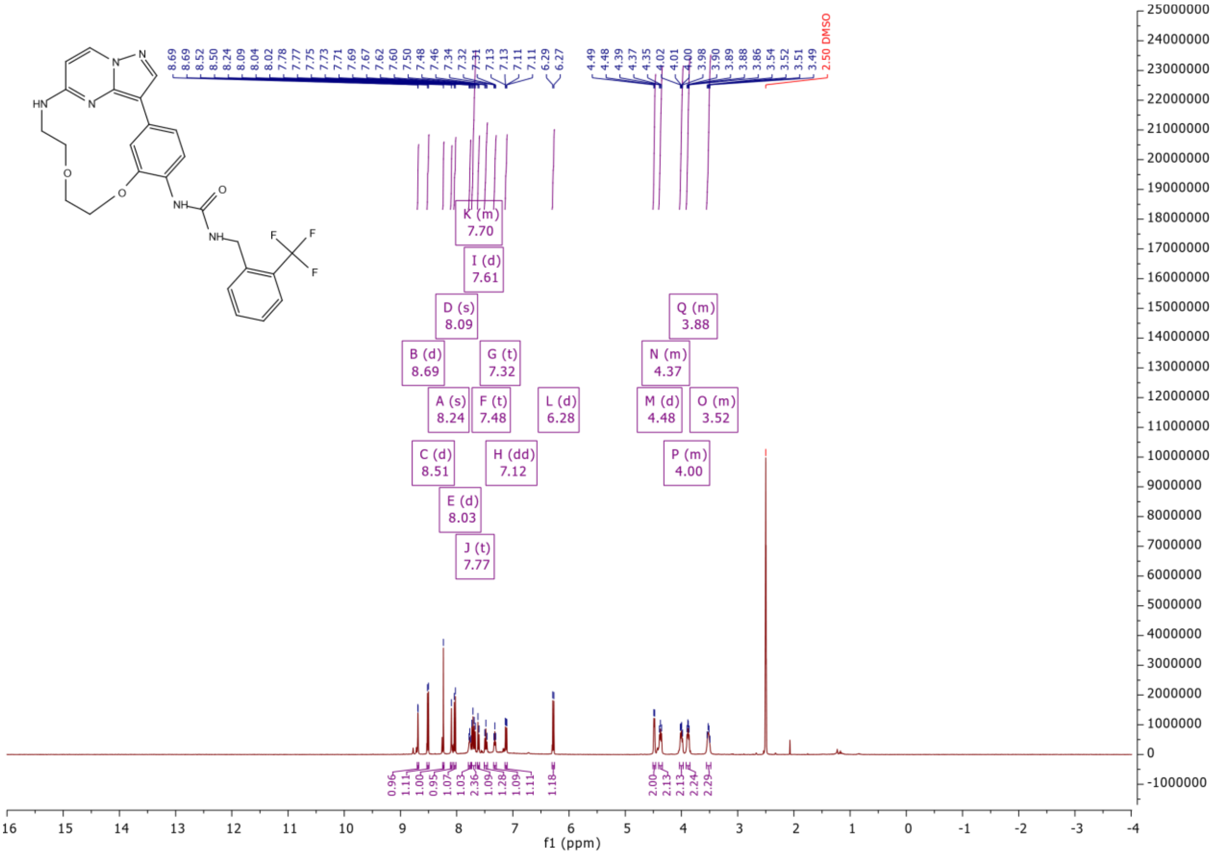

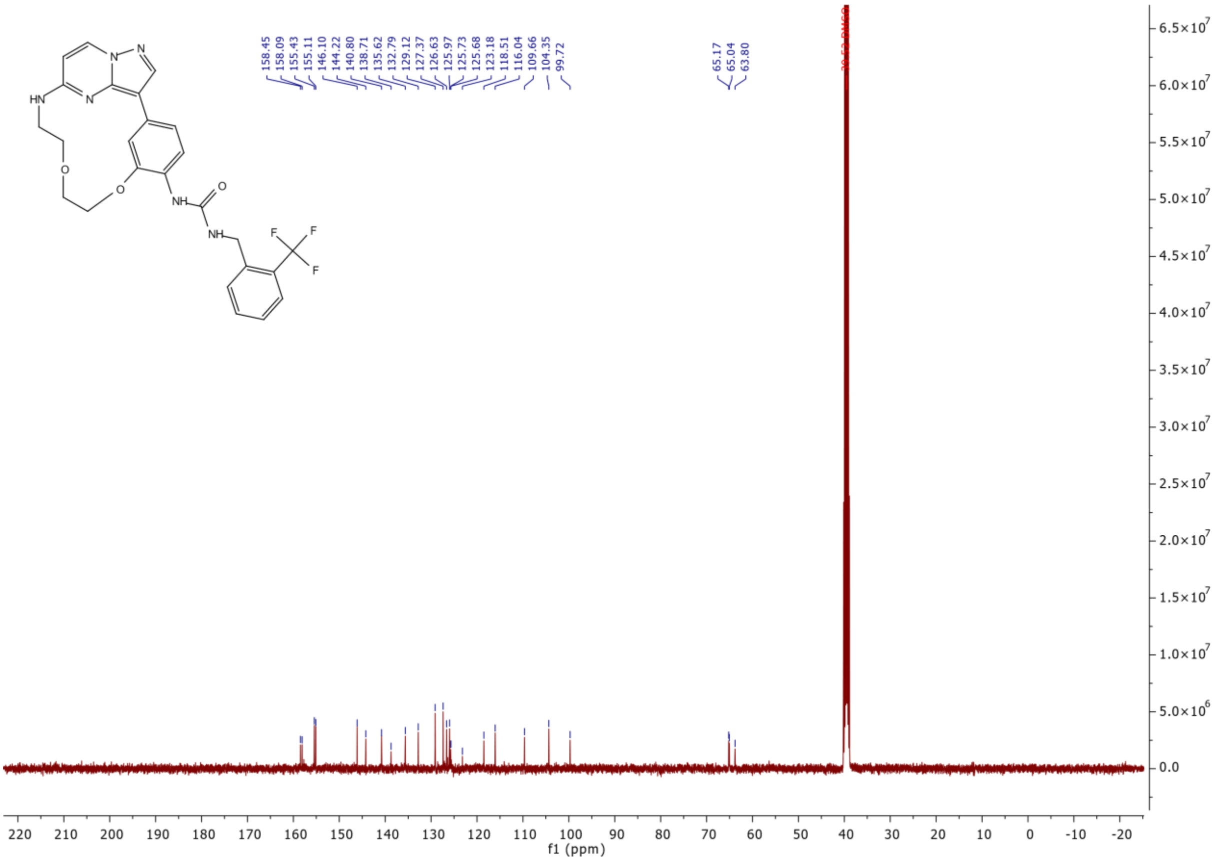

HN
N
N
N
O
O
NH
O
NH
F
F
F
158.45
158.09
155.43
155.11
146.10
144.22
140.80
138.71
135.62
132.79
129.12
127.37
126.63
125.97
125.73
125.68
123.18
118.51
116.04
109.66
104.35
99.72
65.17
65.04
63.80
f1 (ppm)

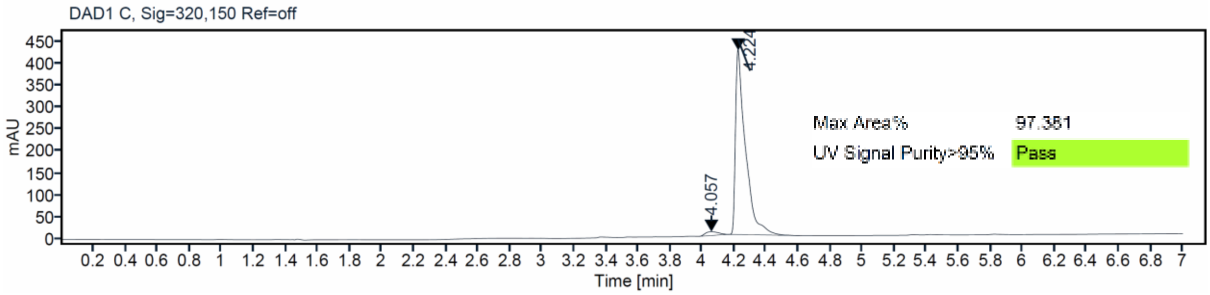

DAD1 C, Sig=320,150 Ref=off
mAU
4.057
4.224
Max Area%
97.381
UV Signal Purity>95%
Pass
Time [min]

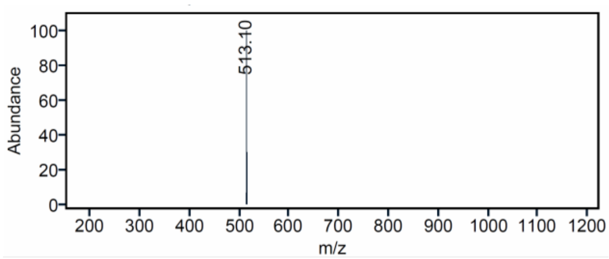

Abundance
513.10
m/z

### Mass Spectrum SmartFormula Report

**Analysis Info**

| | | | |
|---|---|---|---|
| Analysis Name | D:\Data\MS_Service\2024Q4\20241015_SKN_FG_000066-77\SKN_FG_000069_TML81_1.d | Acquisition Date | 10/15/2024 4:36:42 PM |
| Method | Positive(100-1600)_Infusion_200uL-min.m | Operator | BDAL@DE |
| Sample Name | Test1 | Instrument | micrOTOF-Q 228888.10407 |
| Comment | | | |

**Acquisition Parameter**

| | | | | | |
|---|---|---|---|---|---|
| Source Type | ESI | Ion Polarity | Positive | Set Nebulizer | 1.8 Bar |
| Focus | Active | Set Capillary | 4000 V | Set Dry Heater | 280 °C |
| Scan Begin | 100 m/z | Set End Plate Offset | -500 V | Set Dry Gas | 8.0 l/min |
| Scan End | 1600 m/z | Set Collision Cell RF | 150.0 Vpp | Set Divert Valve | Waste |

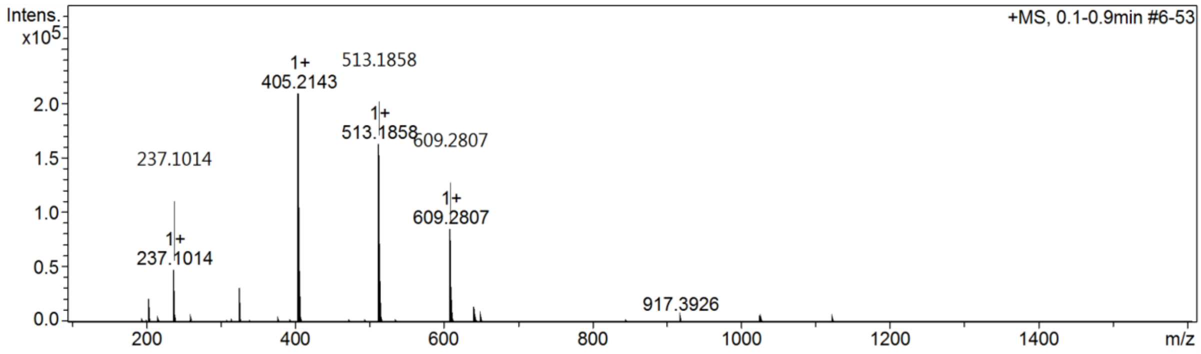

[1]H and [13]C {H} NMR, HPLC/MS (ESI) and HRMS data of compound **28**

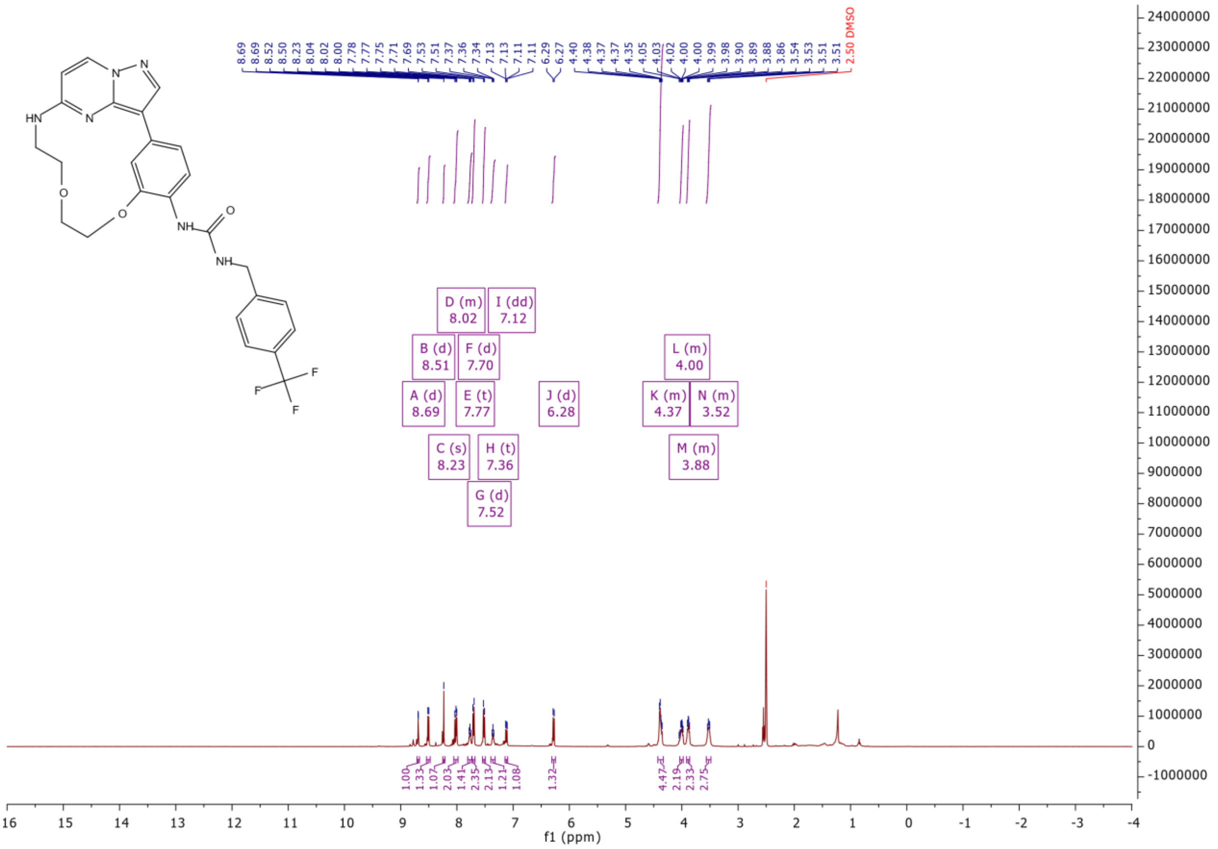

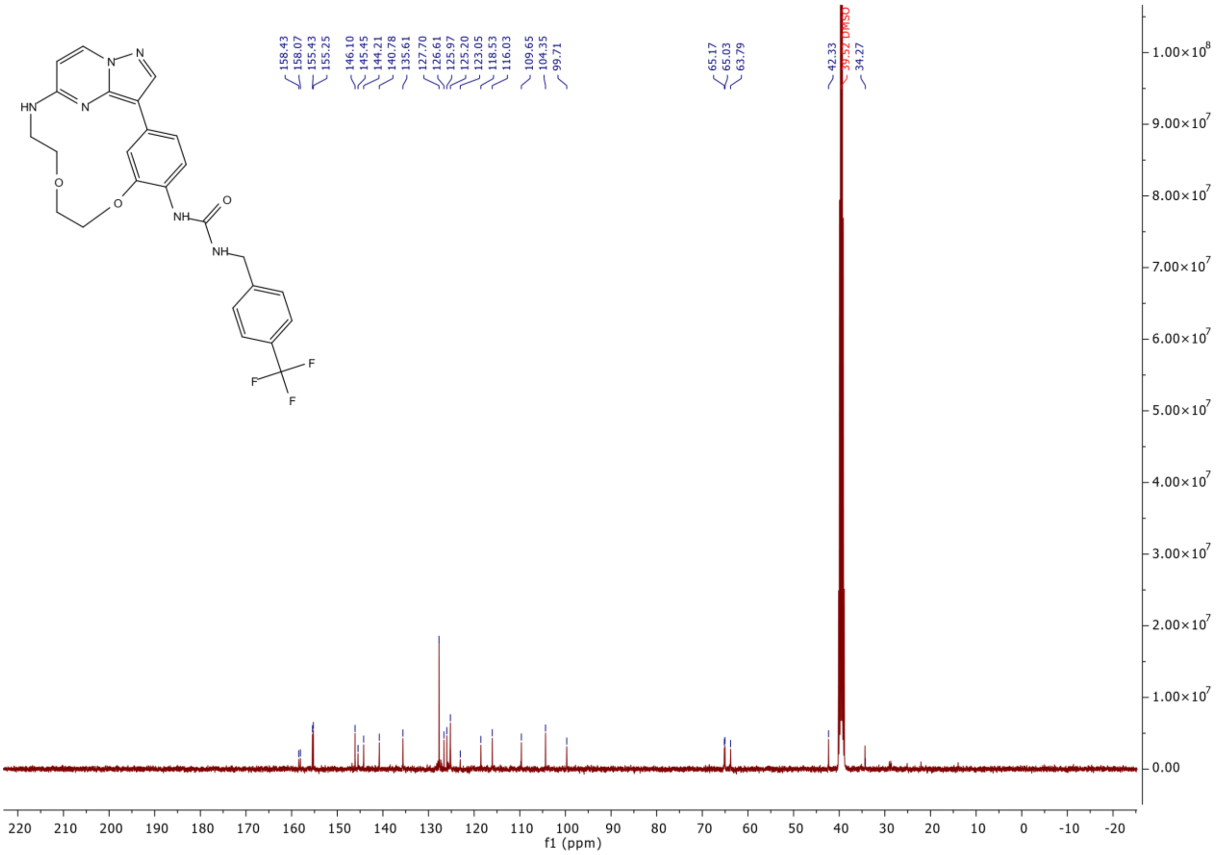
158.43
158.07
155.43
155.25
146.10
145.45
144.21
140.78
135.61
127.70
126.61
125.97
125.20
123.05
118.53
116.03
109.65
104.35
99.71
65.17
65.03
63.79
42.33
39.52 DMSO
34.27
f1 (ppm)

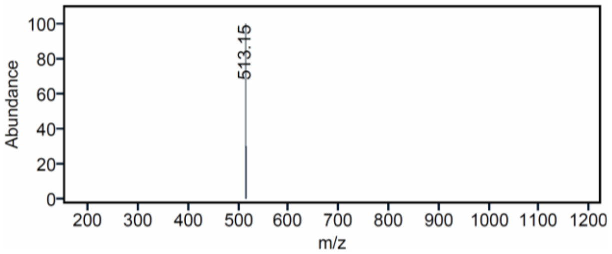
513.15
Abundance
m/z

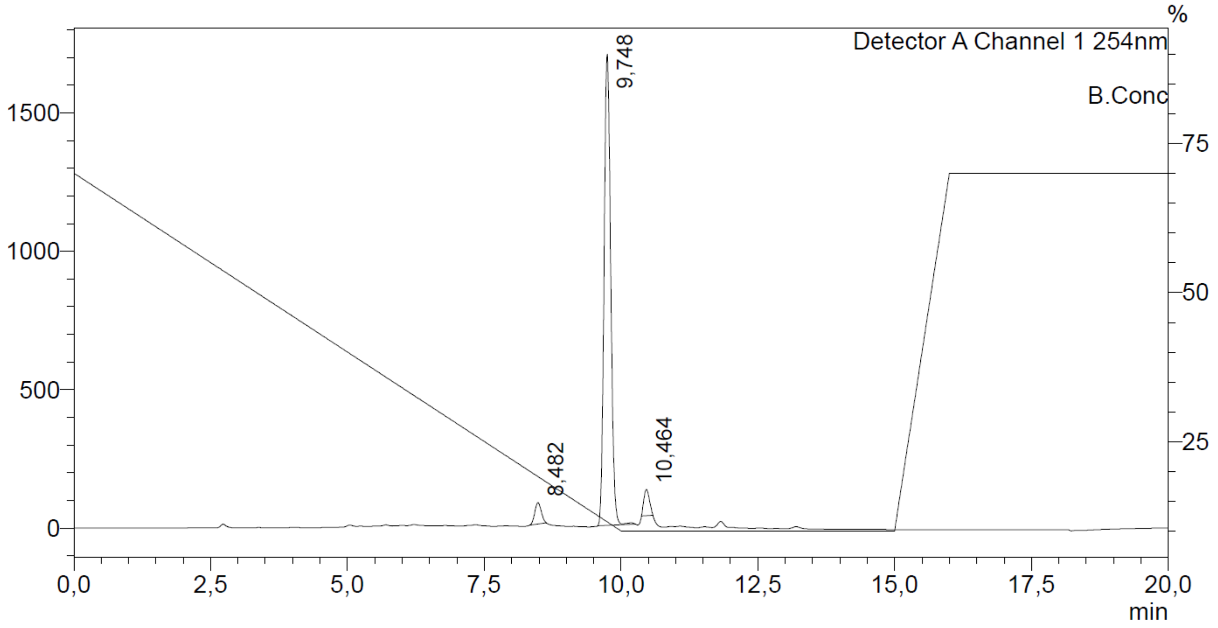
Detector A Channel 1 254nm
B.Conc
%
9,748
8,482
10,464
min

### Mass Spectrum SmartFormula Report

**Analysis Info** | Acquisition Date 10/15/2024 4:31:57 PM

| | | | |
|---|---|---|---|
| Analysis Name | D:\Data\MS_Service\2024Q4\20241015_SKN_FG_000066-77\SKN_FG_000068_TML80_1.d | | |
| Method | Positive(100-1600)_Infusion_200uL-min.m | Operator | BDAL@DE |
| Sample Name | Test1 | Instrument | micrOTOF-Q 228888.10407 |
| Comment | | | |

**Acquisition Parameter**

| | | | | | |
|---|---|---|---|---|---|
| Source Type | ESI | Ion Polarity | Positive | Set Nebulizer | 1.8 Bar |
| Focus | Active | Set Capillary | 4000 V | Set Dry Heater | 280 °C |
| Scan Begin | 100 m/z | Set End Plate Offset | -500 V | Set Dry Gas | 8.0 l/min |
| Scan End | 1600 m/z | Set Collision Cell RF | 150.0 Vpp | Set Divert Valve | Waste |

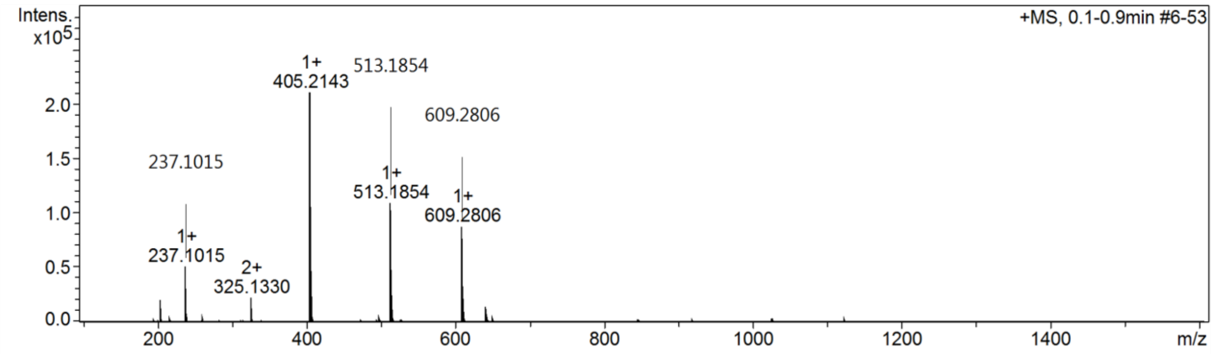

$^{1}H$ and $^{13}C$ {H} NMR, HPLC/MS (ESI) and HRMS data of compound **29**

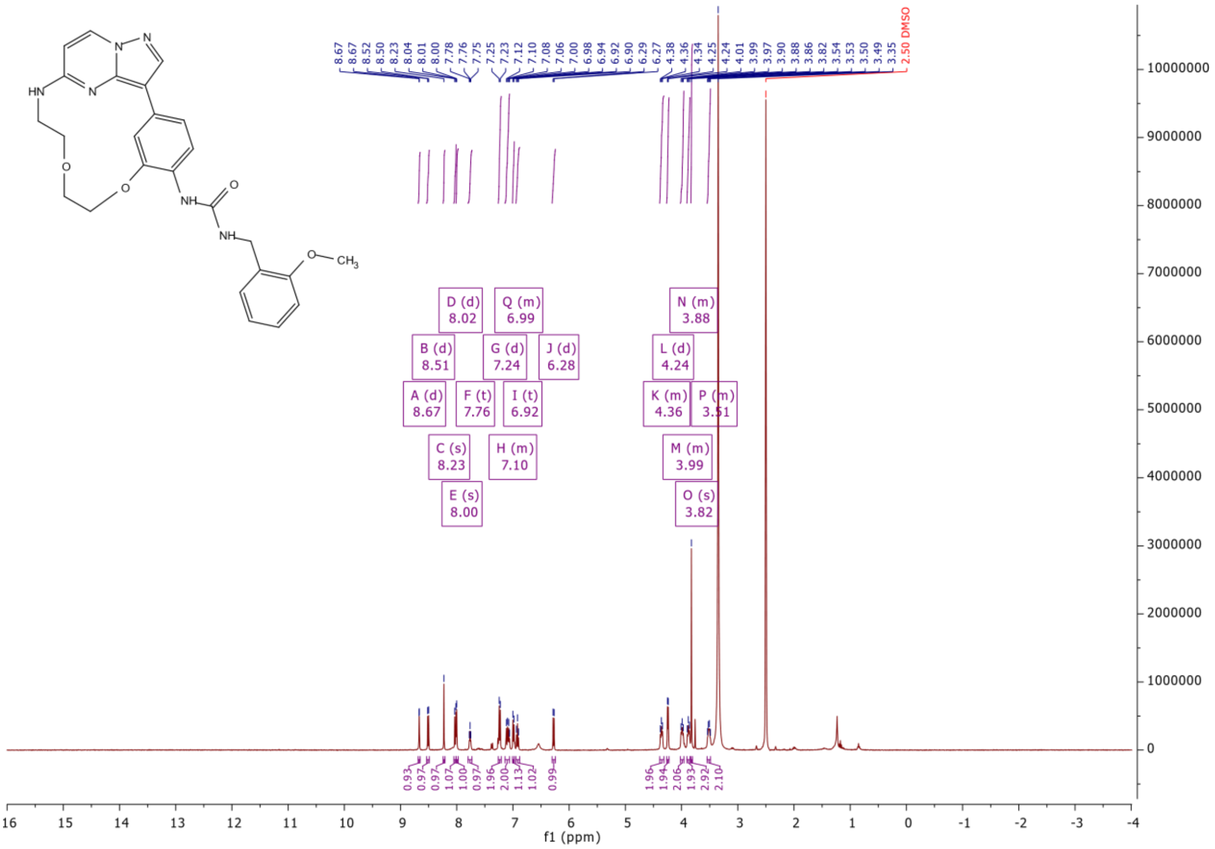

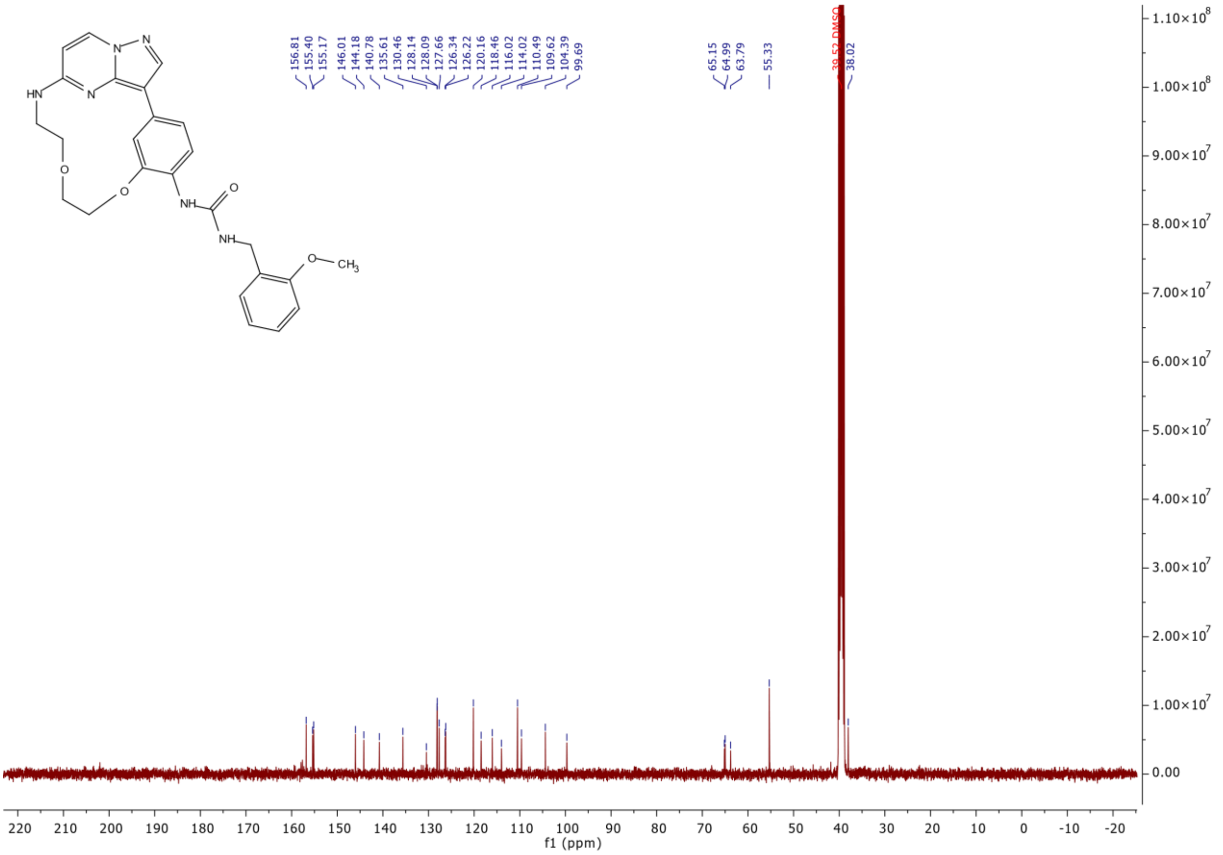

HN
N
N
N
O
O
NH
O
NH
O
CH3
156.81
155.40
155.17
146.01
144.18
140.78
135.61
130.46
128.14
128.09
127.66
126.34
126.22
120.16
118.46
116.02
114.02
110.49
109.62
104.39
99.69
65.15
64.99
63.79
55.33
39.52 DMSO
38.02
f1 (ppm)

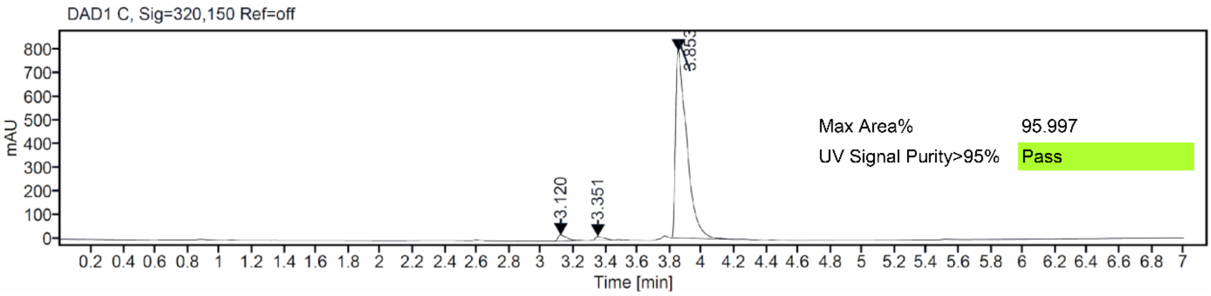

DAD1 C, Sig=320,150 Ref=off
mAU
3.120
3.351
3.853
Max Area% 95.997
UV Signal Purity>95% Pass
Time [min]

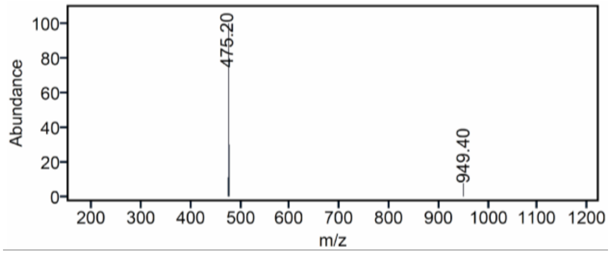

Abundance
475.20
949.40
m/z

### Mass Spectrum SmartFormula Report

**Analysis Info** Acquisition Date 10/15/2024 4:51:33 PM

| | | | |
|---|---|---|---|
| Analysis Name | D:\Data\MS_Service\2024Q4\20241015_SKN_FG_000066-77\SKN_FG_000072_TML62u_1.d | | |
| Method | Positive(100-1600)_Infusion_200uL-min.m | Operator | BDAL@DE |
| Sample Name | Test1 | Instrument | micrOTOF-Q 228888.10407 |
| Comment | | | |

**Acquisition Parameter**

| | | | | | |
|---|---|---|---|---|---|
| Source Type | ESI | Ion Polarity | Positive | Set Nebulizer | 1.8 Bar |
| Focus | Active | Set Capillary | 4000 V | Set Dry Heater | 280 °C |
| Scan Begin | 100 m/z | Set End Plate Offset | -500 V | Set Dry Gas | 8.0 l/min |
| Scan End | 1600 m/z | Set Collision Cell RF | 150.0 Vpp | Set Divert Valve | Waste |

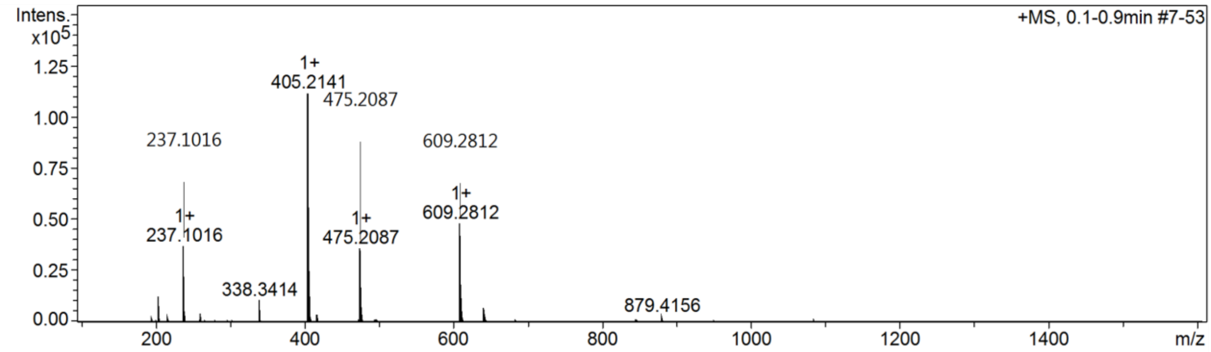

[1]H and [13]C {H} NMR, HPLC/MS (ESI) and HRMS data of compound **30**

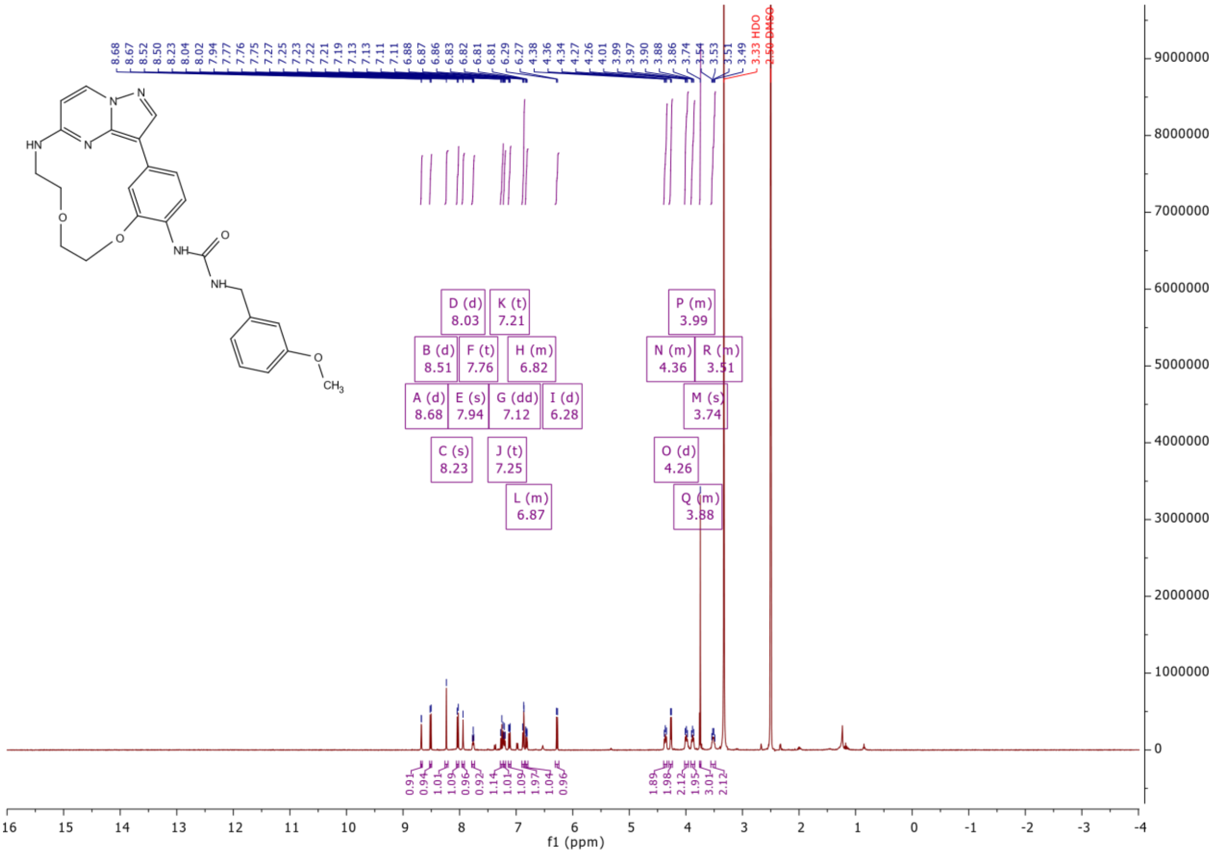

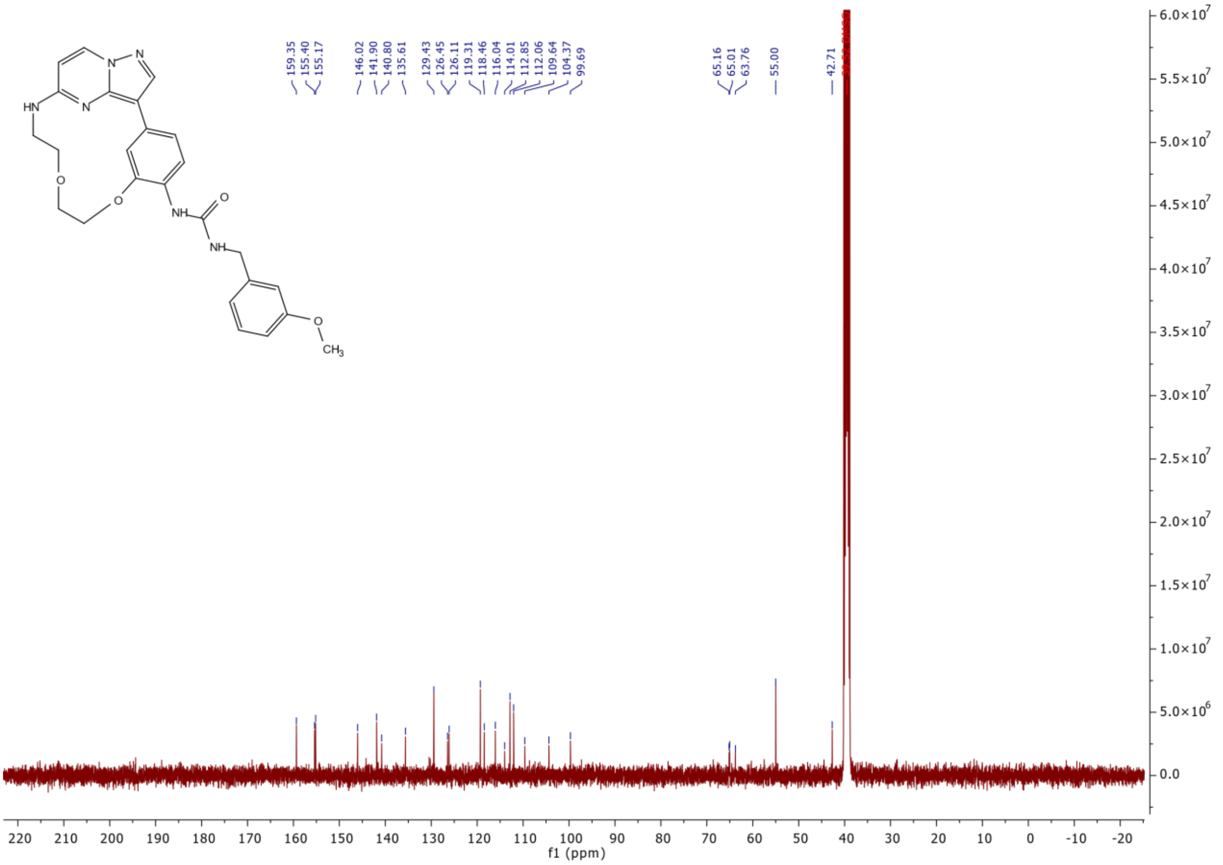
159.35
155.40
155.17
146.02
141.90
140.80
135.61
129.43
126.45
126.11
119.31
118.46
116.04
114.01
112.85
112.06
109.64
104.37
99.69
65.16
65.01
63.76
55.00
42.71
f1 (ppm)

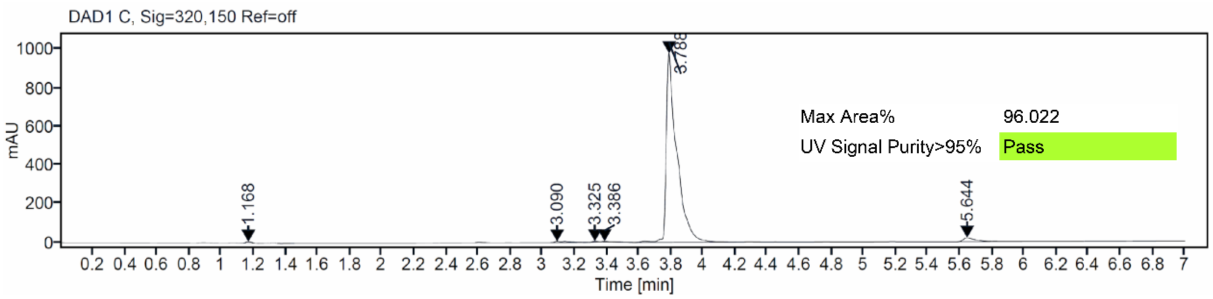
DAD1 C, Sig=320,150 Ref=off
mAU
1.168
3.090
3.325
3.386
3.788
5.644
Max Area% 96.022
UV Signal Purity>95% Pass
Time [min]

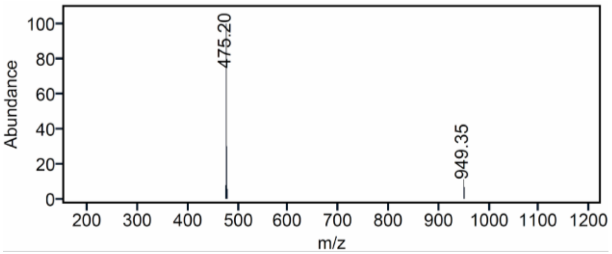
Abundance
475.20
949.35
m/z

### Mass Spectrum SmartFormula Report

**Analysis Info**

| | | | |
|---|---|---|---|
| Analysis Name | D:\Data\MS_Service\2024Q4\20241015_SKN_FG_000066-77\SKN_FG_000071_TML60u_1.d | Acquisition Date | 10/15/2024 4:47:03 PM |
| Method | Positive(100-1600)_Infusion_200uL-min.m | Operator | BDAL@DE |
| Sample Name | Test1 | Instrument | micrOTOF-Q 228888.10407 |
| Comment | | | |

**Acquisition Parameter**

| | | | | | |
|---|---|---|---|---|---|
| Source Type | ESI | Ion Polarity | Positive | Set Nebulizer | 1.8 Bar |
| Focus | Active | Set Capillary | 4000 V | Set Dry Heater | 280 °C |
| Scan Begin | 100 m/z | Set End Plate Offset | -500 V | Set Dry Gas | 8.0 l/min |
| Scan End | 1600 m/z | Set Collision Cell RF | 150.0 Vpp | Set Divert Valve | Waste |

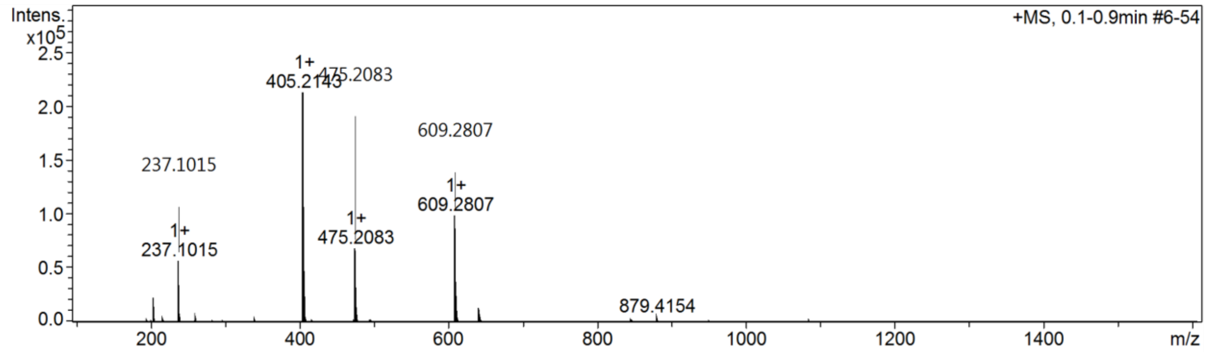

$^1$H and $^{13}$C {H} NMR, HPLC/MS (ESI) and HRMS data of compound **31**

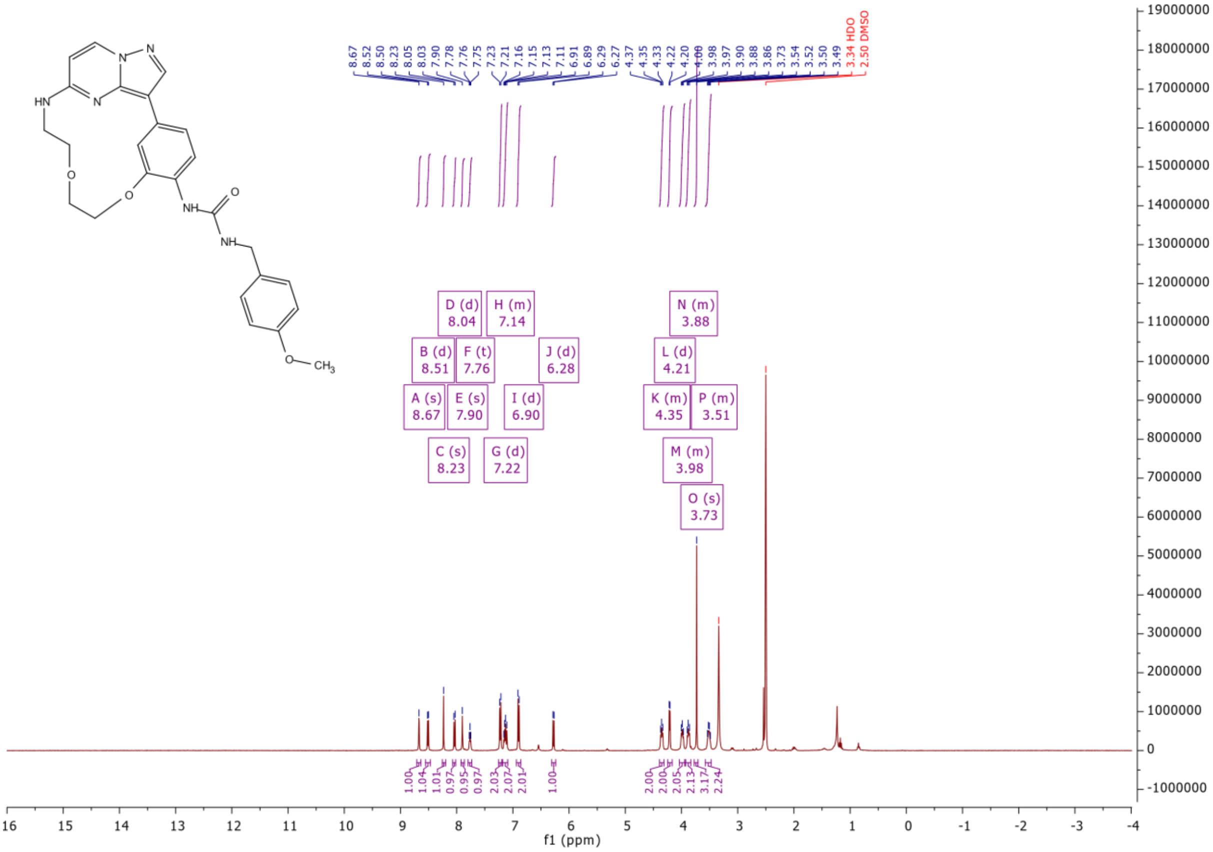

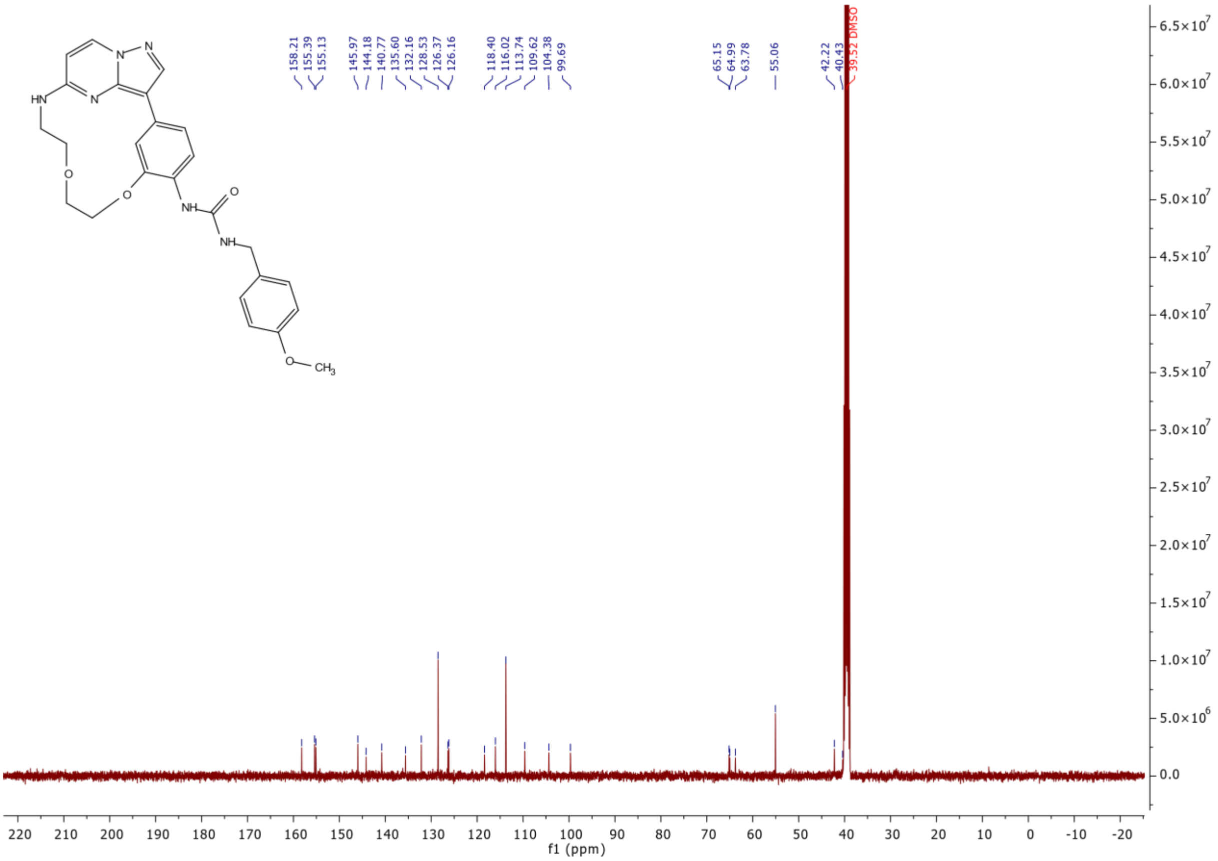

158.21
155.39
155.13
145.97
144.18
140.77
135.60
132.16
128.53
126.37
126.16
118.40
116.02
113.74
109.62
104.38
99.69
65.15
64.99
63.78
55.06
42.22
40.43
39.52 DMSO
f1 (ppm)

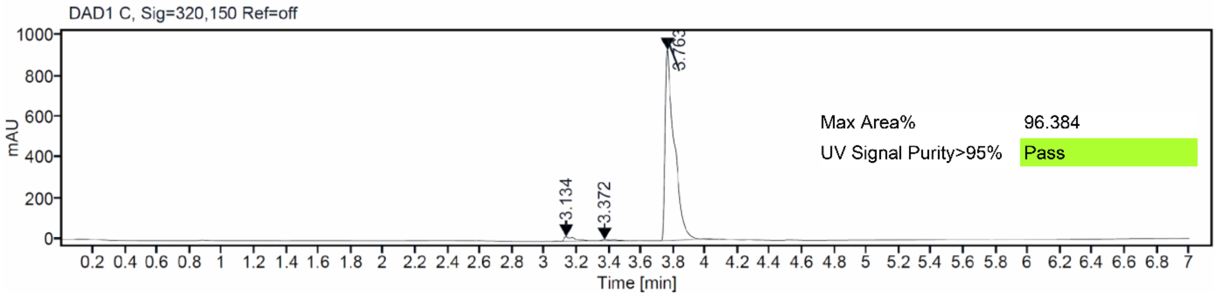

DAD1 C, Sig=320,150 Ref=off
mAU
Time [min]
3.134
3.372
3.763
Max Area% 96.384
UV Signal Purity>95% Pass

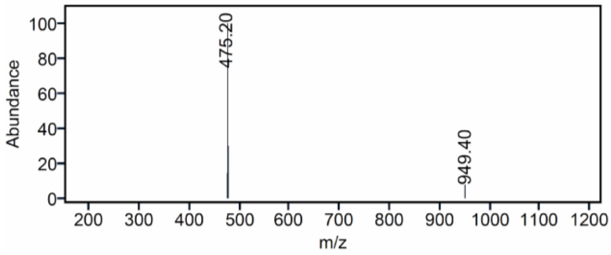

Abundance
m/z
475.20
949.40

### Mass Spectrum SmartFormula Report

**Analysis Info**

| | | | |
|---|---|---|---|
| Analysis Name | D:\Data\MS_Service\2024Q4\20241015_SKN_FG_000066-77\SKN_FG_000073_TMI.64u_1.d | Acquisition Date | 10/15/2024 5:41:45 PM |
| Method | Positive(100-1600)_Infusion_200uL-min.m | Operator | BDAL@DE |
| Sample Name | Test1 | Instrument | micrOTOF-Q 228888.10407 |
| Comment | | | |

**Acquisition Parameter**

| | | | | | |
|---|---|---|---|---|---|
| Source Type | ESI | Ion Polarity | Positive | Set Nebulizer | 1.8 Bar |
| Focus | Active | Set Capillary | 4000 V | Set Dry Heater | 280 °C |
| Scan Begin | 100 m/z | Set End Plate Offset | -500 V | Set Dry Gas | 8.0 l/min |
| Scan End | 1600 m/z | Set Collision Cell RF | 150.0 Vpp | Set Divert Valve | Waste |

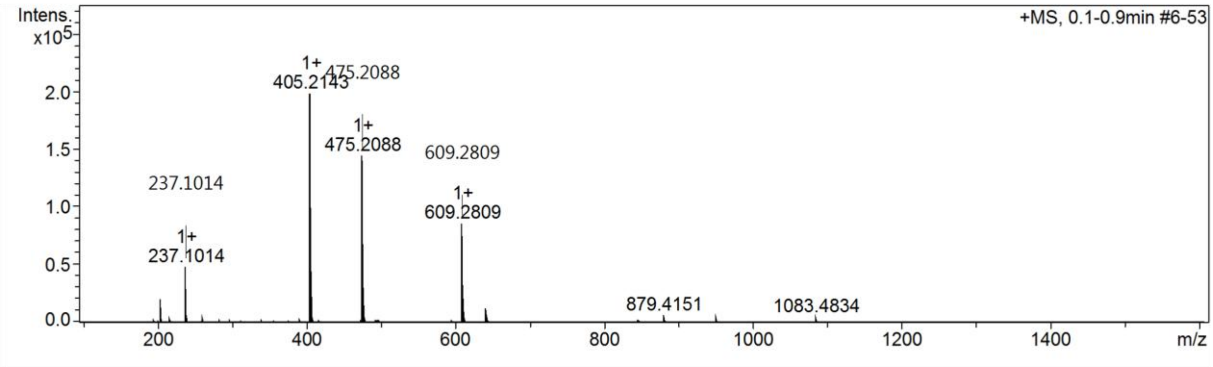

$^1$H and $^{13}$C {H} NMR, HPLC/MS (ESI) and HRMS data of compound **32**

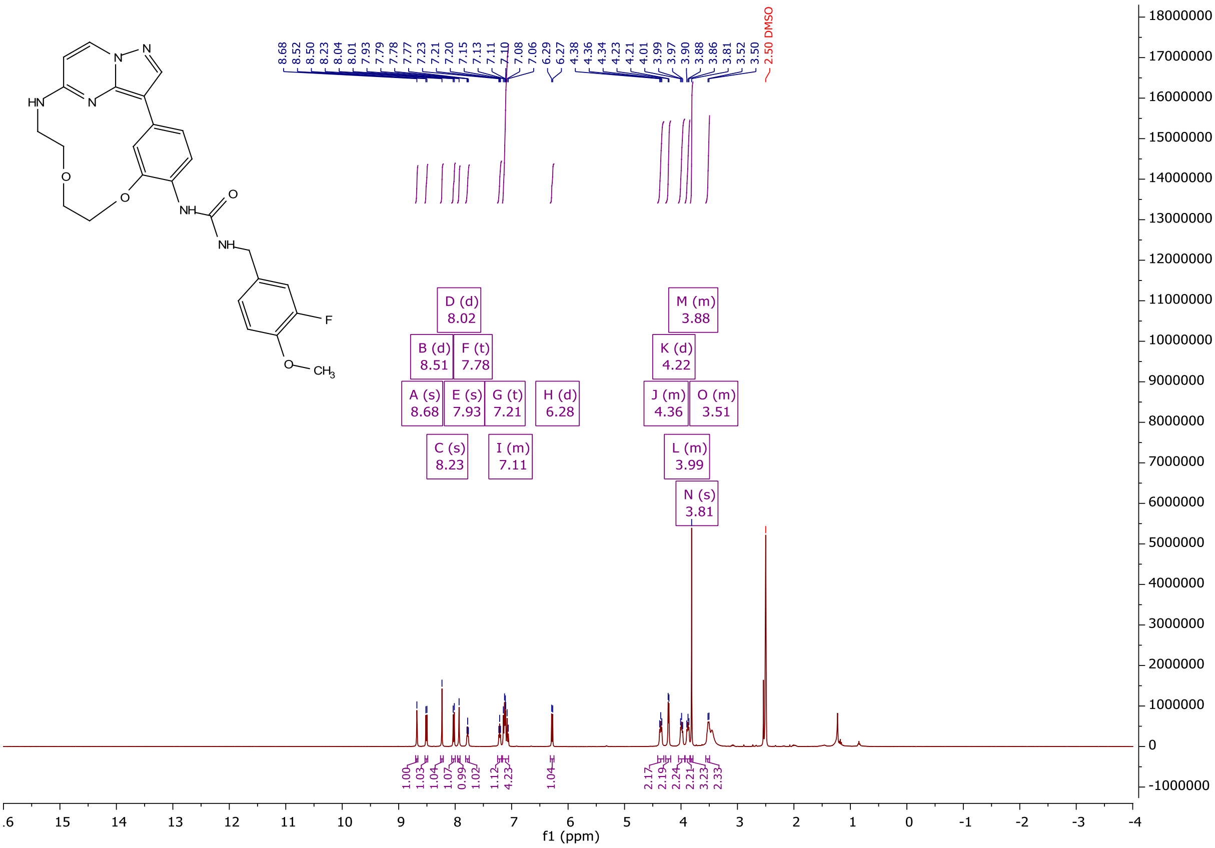

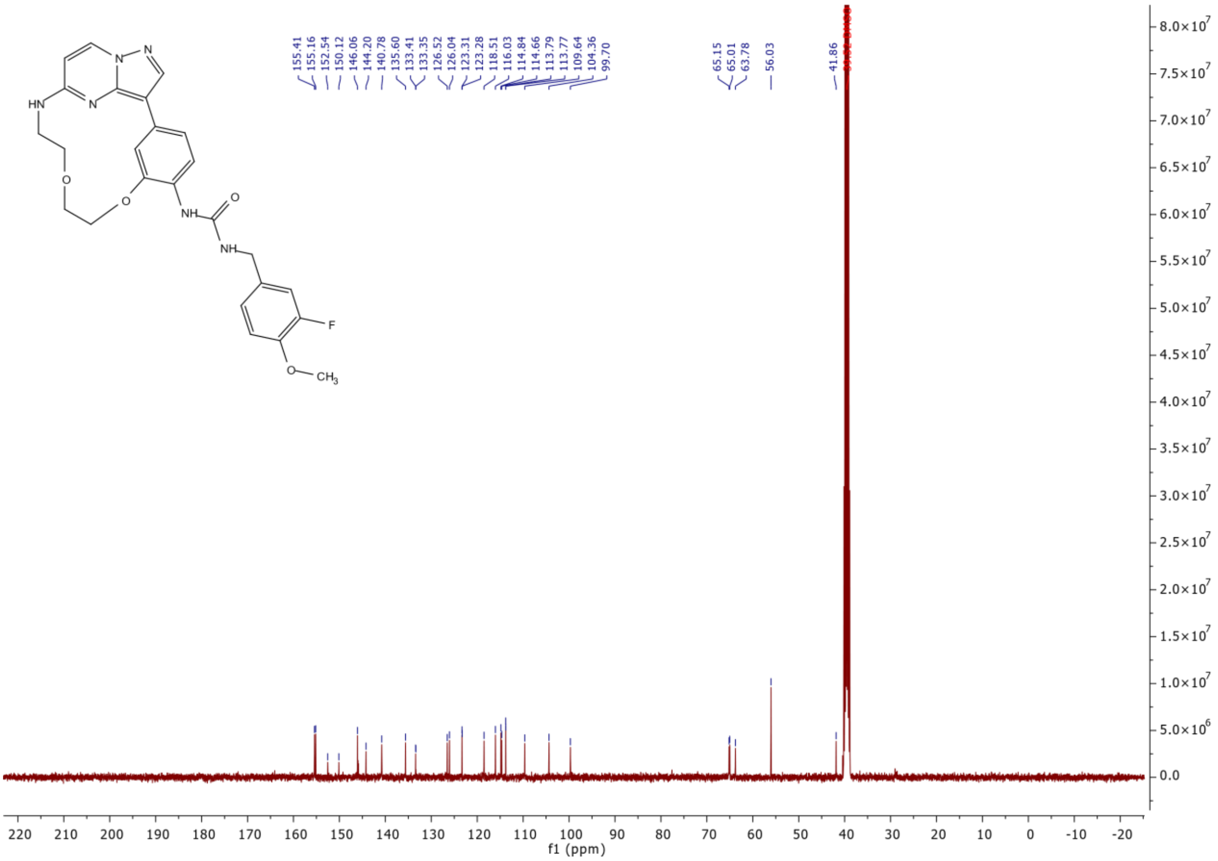
155.41
155.16
152.54
150.12
146.06
144.20
140.78
135.60
133.41
133.35
126.52
126.04
123.31
123.28
118.51
116.03
114.84
114.66
113.79
113.77
109.64
104.36
99.70
65.15
65.01
63.78
56.03
41.86
HN
N
N
N
O
O
NH
O
NH
F
O
CH3
f1 (ppm)

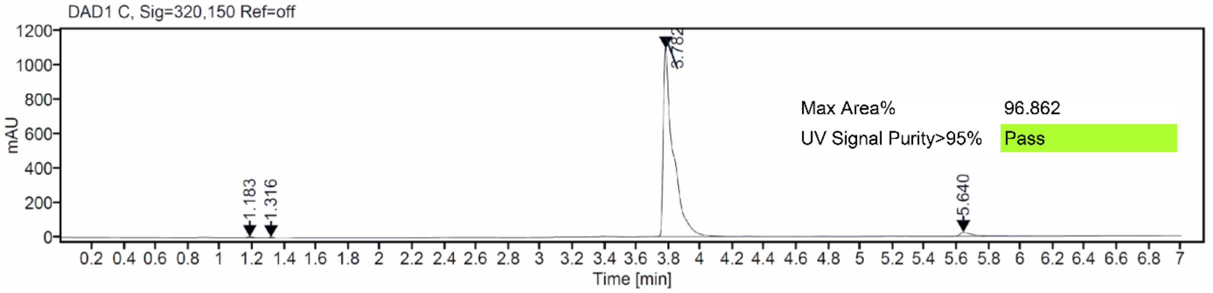
DAD1 C, Sig=320,150 Ref=off
mAU
1.183
1.316
3.782
5.640
Max Area% 96.862
UV Signal Purity>95% Pass
Time [min]

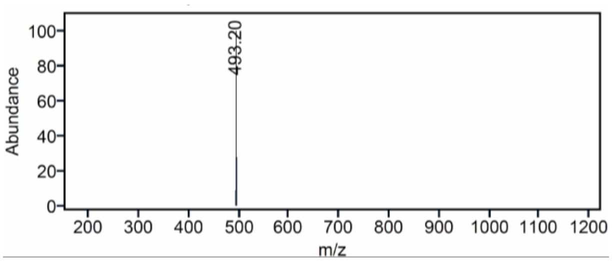
Abundance
493.20
m/z

### Mass Spectrum SmartFormula Report

#### Analysis Info

| | | | |
|---|---|---|---|
| Analysis Name | D:\Data\MS_Service\2024Q4\20241015_SKN_FG_000066-77\SKN_FG_000075_TML73u_1.d | Acquisition Date | 10/15/2024 5:51:26 PM |
| Method | Positive(100-1600)_Infusion_200uL-min.m | Operator | BDAL@DE |
| Sample Name | Test1 | Instrument | micrOTOF-Q 228888.10407 |
| Comment | | | |

#### Acquisition Parameter

| | | | | | |
|---|---|---|---|---|---|
| Source Type | ESI | Ion Polarity | Positive | Set Nebulizer | 1.8 Bar |
| Focus | Active | Set Capillary | 4000 V | Set Dry Heater | 280 °C |
| Scan Begin | 100 m/z | Set End Plate Offset | -500 V | Set Dry Gas | 8.0 l/min |
| Scan End | 1600 m/z | Set Collision Cell RF | 150.0 Vpp | Set Divert Valve | Waste |

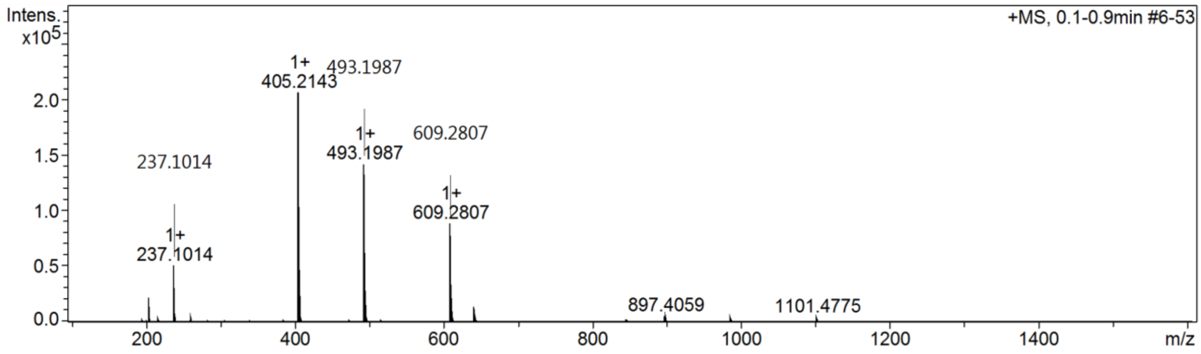

$^1$H and $^{13}$C {H} NMR, HPLC/MS (ESI) and HRMS data of compound **33**

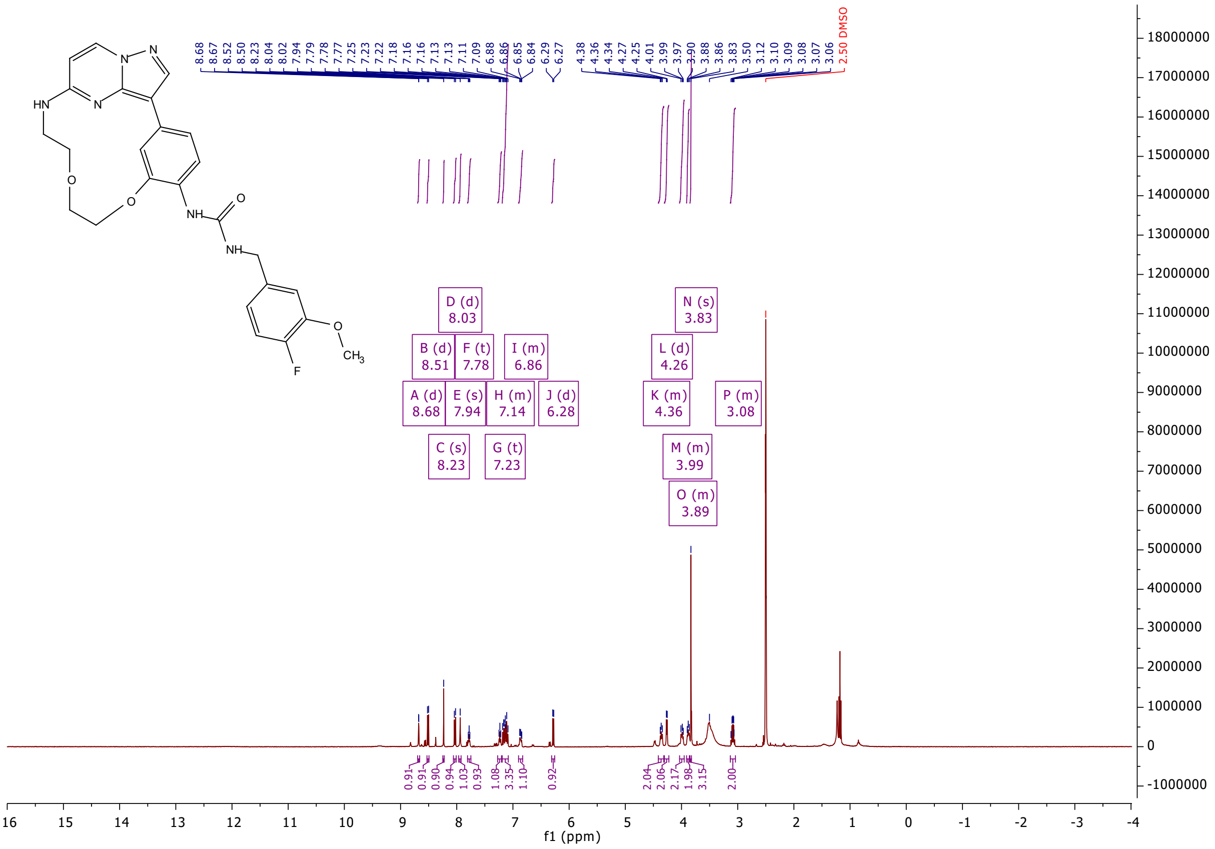

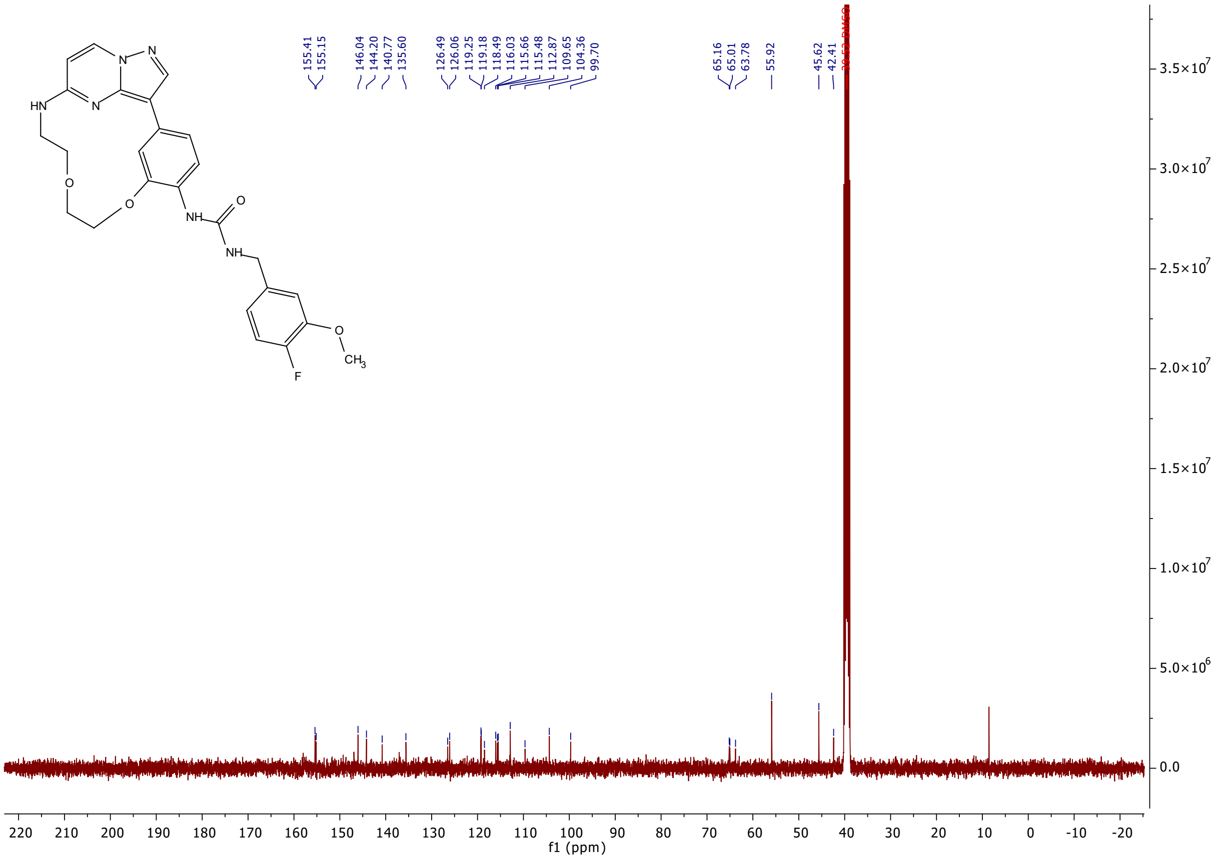
155.41
155.15
146.04
144.20
140.77
135.60
126.49
126.06
119.25
119.18
118.49
116.03
115.66
115.48
112.87
109.65
104.36
99.70
65.16
65.01
63.78
55.92
45.62
42.41
f1 (ppm)

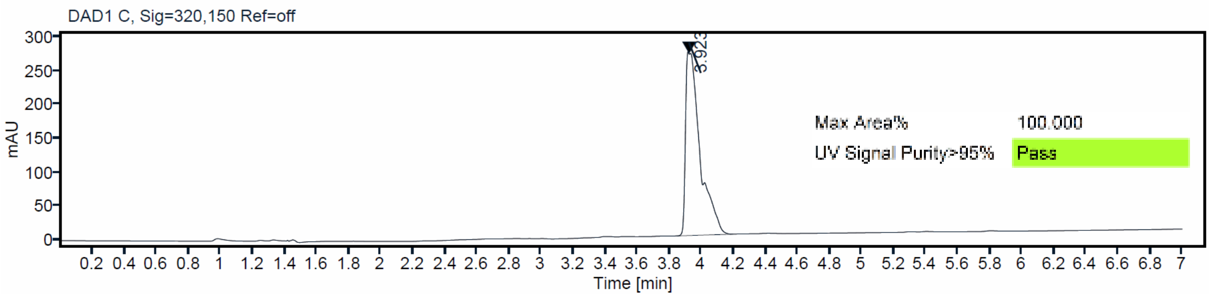
DAD1 C, Sig=320,150 Ref=off
mAU
3.923
Max Area%
100.000
UV Signal Purity>95%
Pass
Time [min]

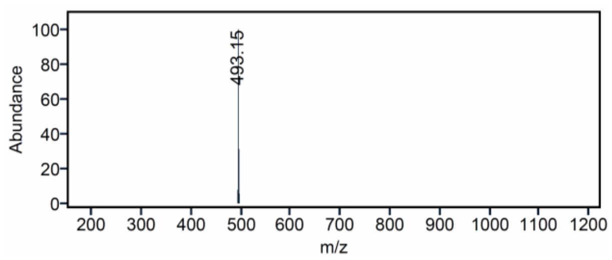
Abundance
493.15
m/z

### Mass Spectrum SmartFormula Report

**Analysis Info**

| | | | |
|---|---|---|---|
| | | Acquisition Date | 10/15/2024 5:46:55 PM |
| Analysis Name | D:\Data\MS_Service\2024Q4\20241015_SKN_FG_000066-77\SKN_FG_000074_TML72_1.d | | |
| Method | Positive(100-1600)_Infusion_200uL-min.m | Operator | BDAL@DE |
| Sample Name | Test1 | Instrument | micrOTOF-Q 228888.10407 |
| Comment | | | |

**Acquisition Parameter**

| | | | | | |
|---|---|---|---|---|---|
| Source Type | ESI | Ion Polarity | Positive | Set Nebulizer | 1.8 Bar |
| Focus | Active | Set Capillary | 4000 V | Set Dry Heater | 280 °C |
| Scan Begin | 100 m/z | Set End Plate Offset | -500 V | Set Dry Gas | 8.0 l/min |
| Scan End | 1600 m/z | Set Collision Cell RF | 150.0 Vpp | Set Divert Valve | Waste |

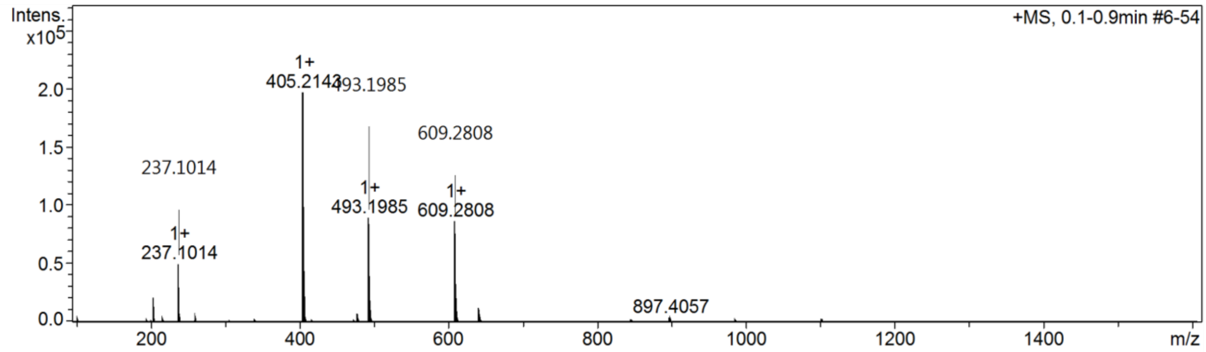

$^1$H and $^{13}$C {H} NMR, HPLC/MS (ESI) and HRMS data of compound **34**

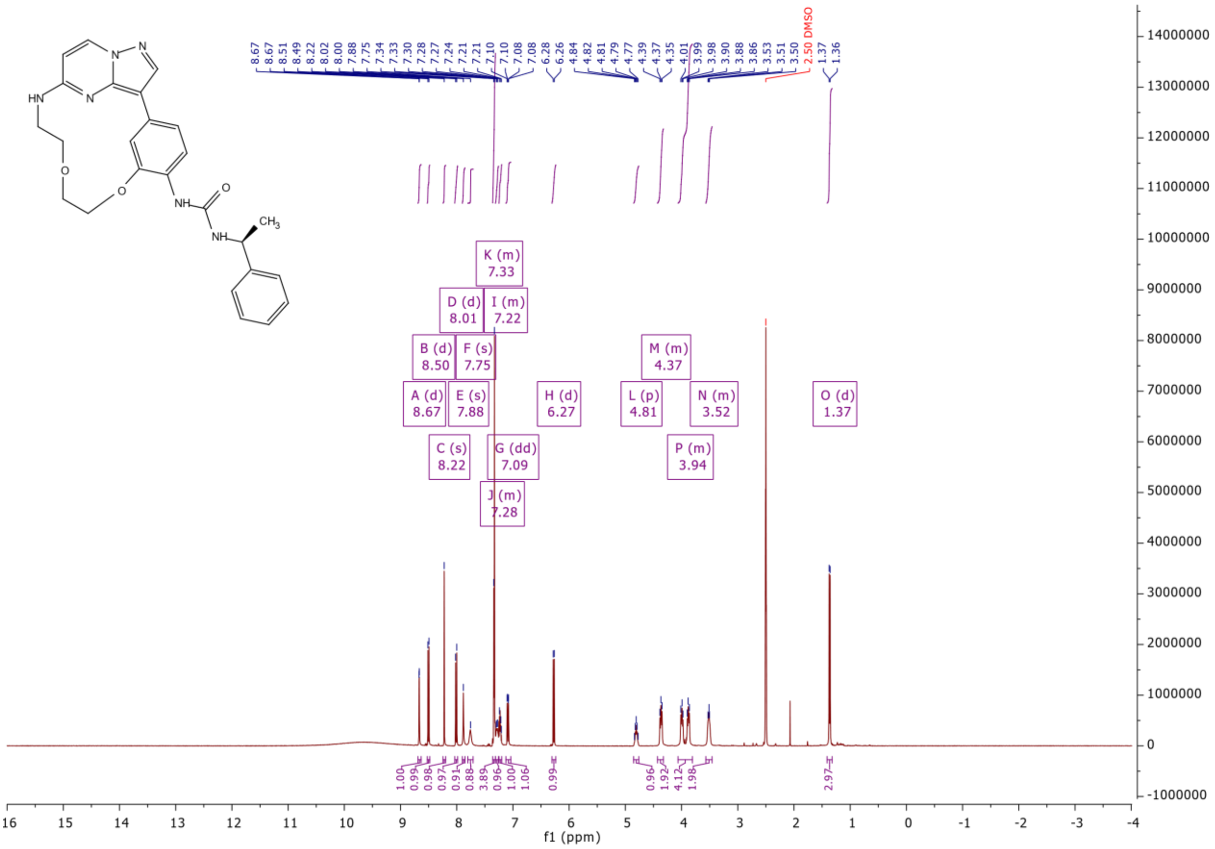

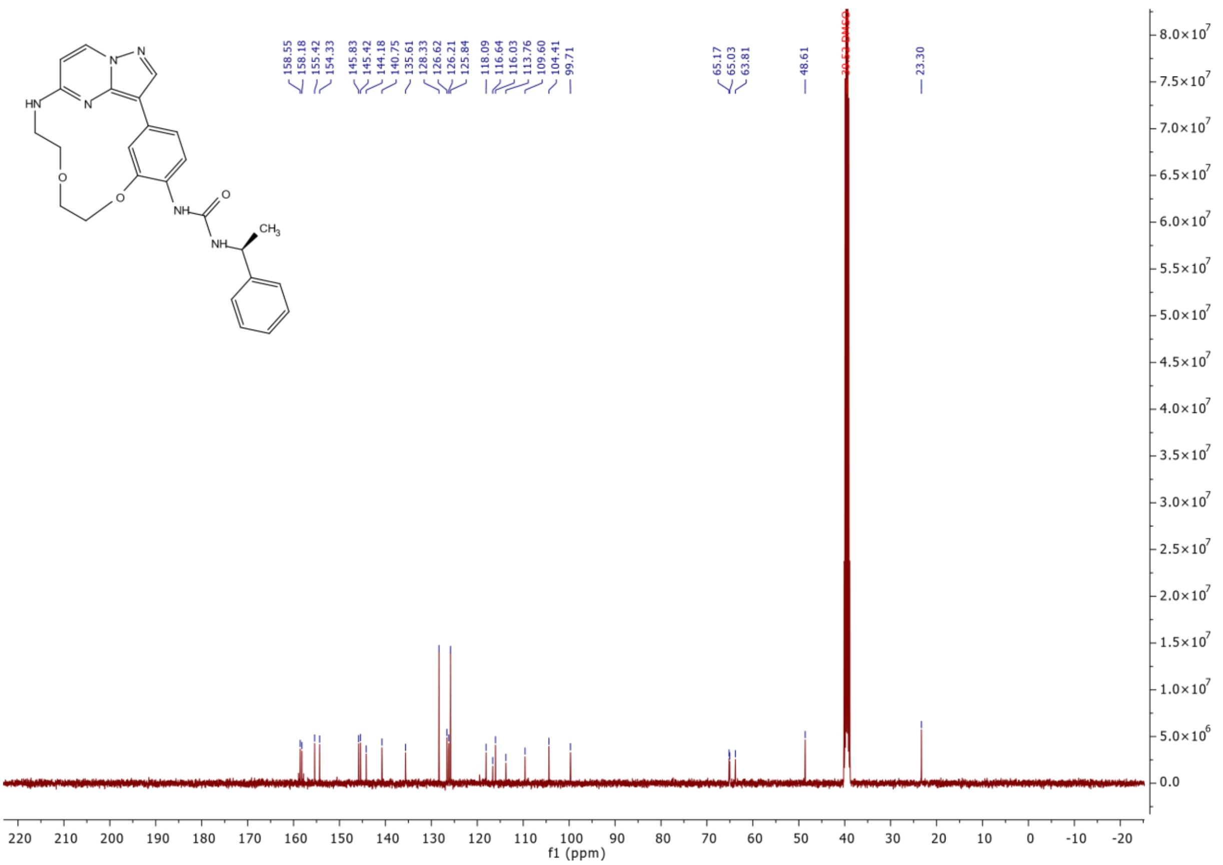
158.55
158.18
155.42
154.33
145.83
145.42
144.18
140.75
135.61
128.33
126.62
126.21
125.84
118.09
116.64
116.03
113.76
109.60
104.41
99.71
65.17
65.03
63.81
48.61
23.30
f1 (ppm)

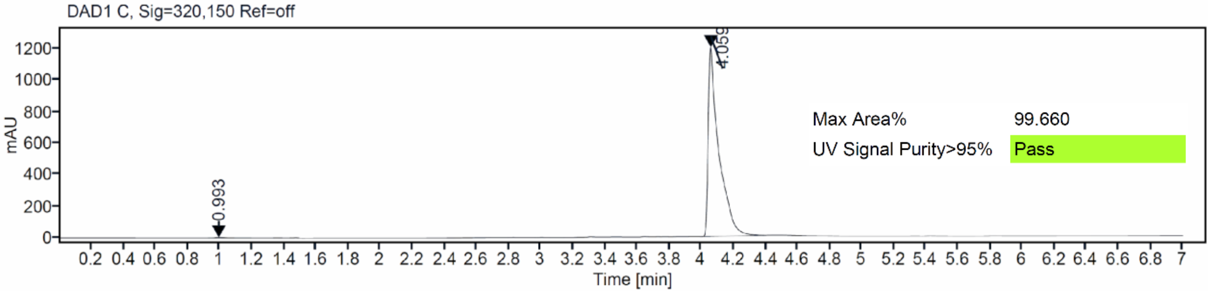
DAD1 C, Sig=320,150 Ref=off
mAU
0.993
4.059
Max Area% 99.660
UV Signal Purity>95% Pass
Time [min]

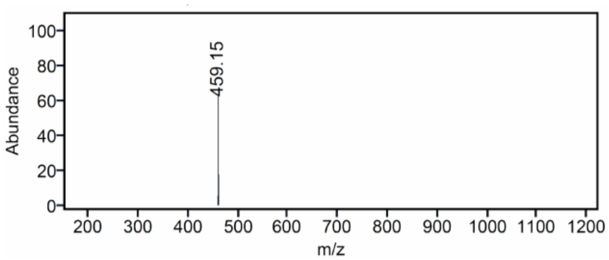
Abundance
459.15
m/z

### Mass Spectrum SmartFormula Report

**Analysis Info**

| | | | |
|---|---|---|---|
| Analysis Name | D:\Data\MS_Service\2024Q4\20241015_SKN_FG_000066-77\SKN_FG_000070_TML54_1.d | Acquisition Date | 10/15/2024 4:42:02 PM |
| Method | Positive(100-1600)_Infusion_200uL-min.m | Operator | BDAL@DE |
| Sample Name | Test1 | Instrument | micrOTOF-Q 228888.10407 |
| Comment | | | |

**Acquisition Parameter**

| | | | | | |
|---|---|---|---|---|---|
| Source Type | ESI | Ion Polarity | Positive | Set Nebulizer | 1.8 Bar |
| Focus | Active | Set Capillary | 4000 V | Set Dry Heater | 280 °C |
| Scan Begin | 100 m/z | Set End Plate Offset | -500 V | Set Dry Gas | 8.0 l/min |
| Scan End | 1600 m/z | Set Collision Cell RF | 150.0 Vpp | Set Divert Valve | Waste |

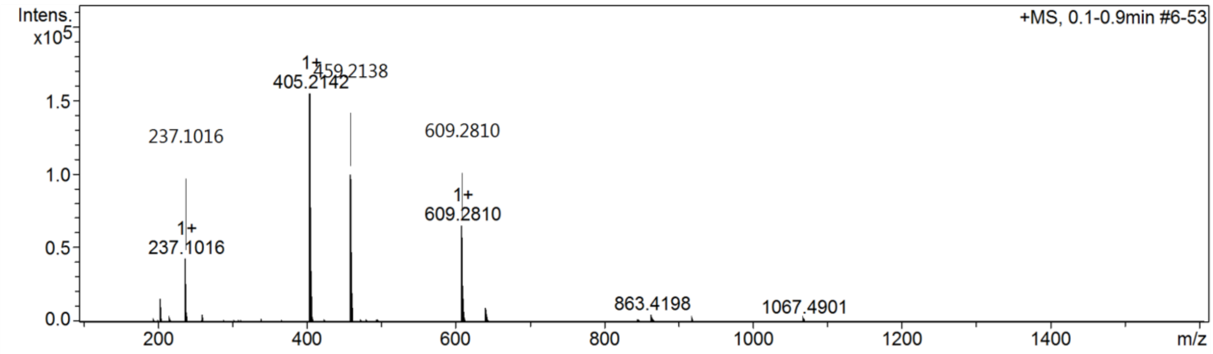

[1]H and [13]C {H} NMR, HPLC/MS (ESI) and HRMS data of compound **35**

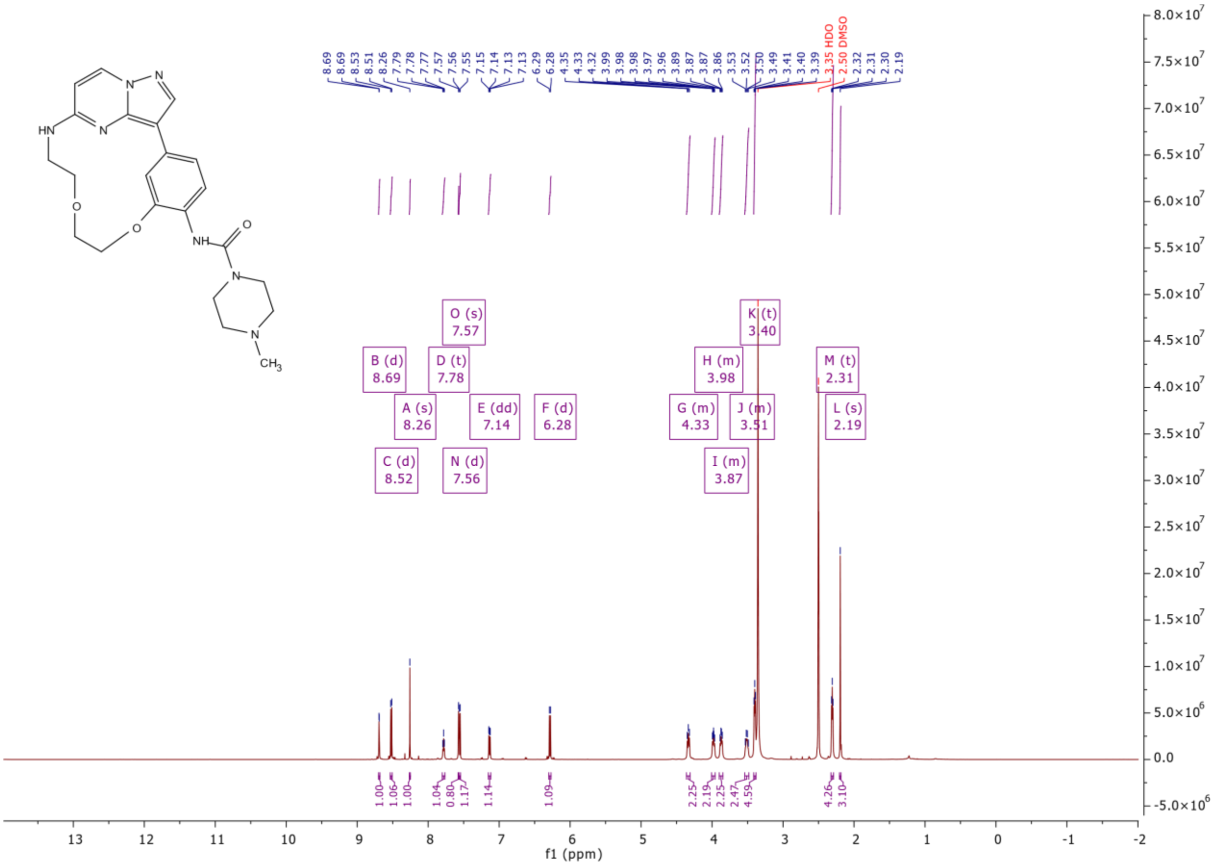

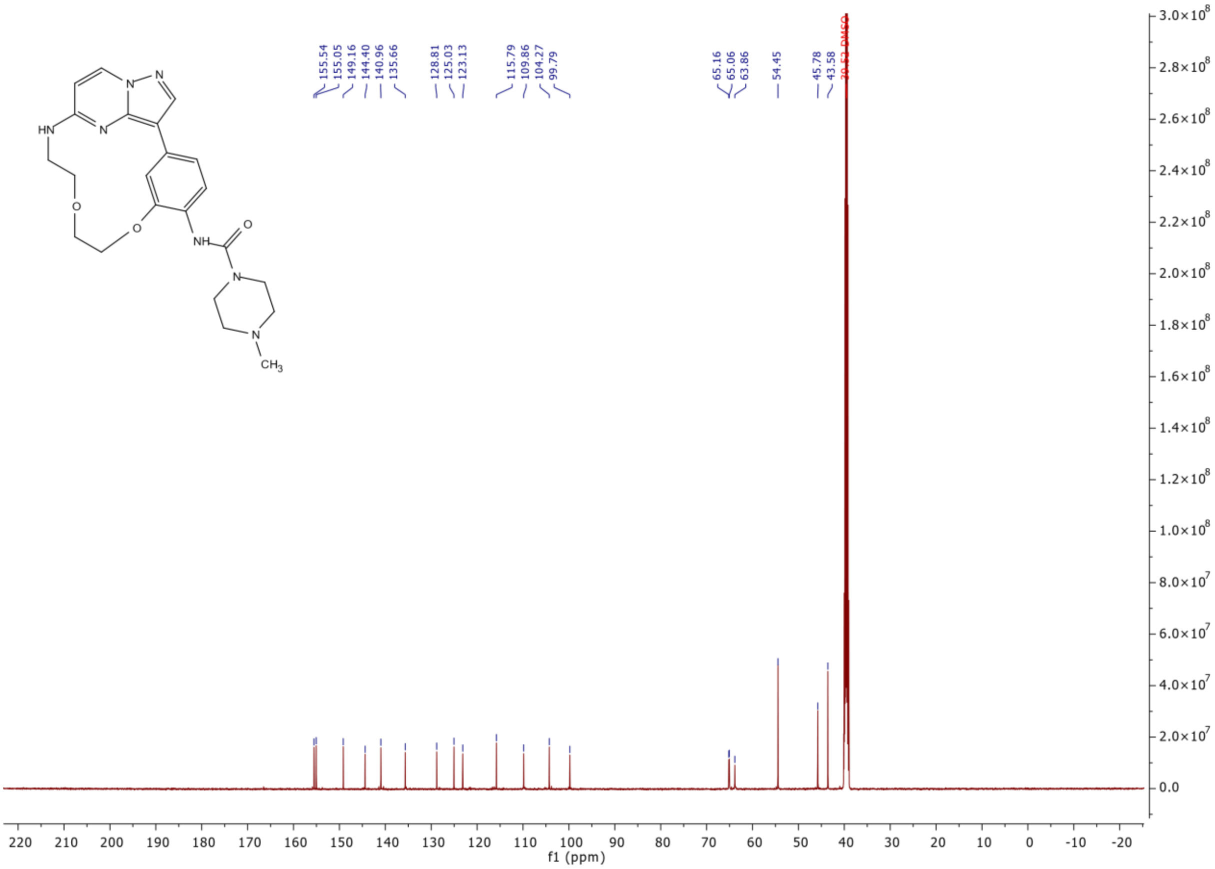
155.54
155.05
149.16
144.40
140.96
135.66
128.81
125.03
123.13
115.79
109.86
104.27
99.79
65.16
65.06
63.86
54.45
45.78
43.58
f1 (ppm)

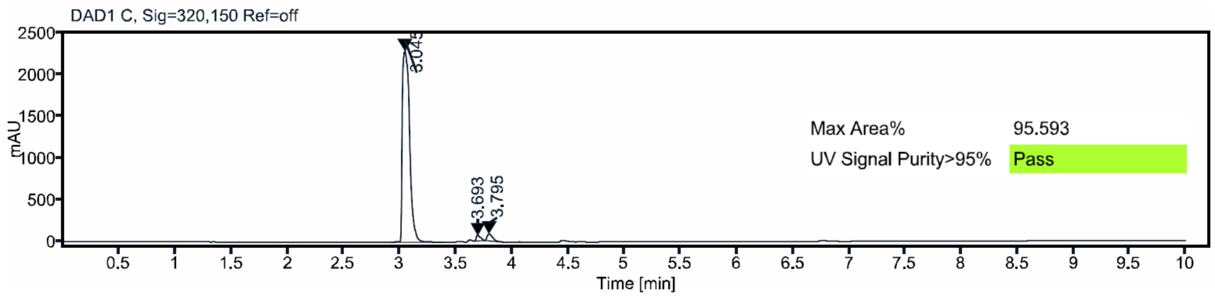
DAD1 C, Sig=320,150 Ref=off
mAU
3.693
3.795
Max Area% 95.593
UV Signal Purity>95% Pass
Time [min]

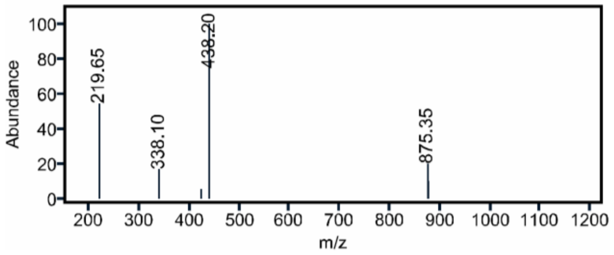
Abundance
219.65
338.10
438.20
875.35
m/z

C:\User\...\Knapp\2018\180723\JB72_E11    7/23/2018 10:28:12 AM    JB72 mit HCCA gemessen.

JB72_E11 #1-7 RT: 0.01-0.48 AV: 7 NL: 2.30E6
T: FTMS + p MALDI Full ms [400.00-700.00]

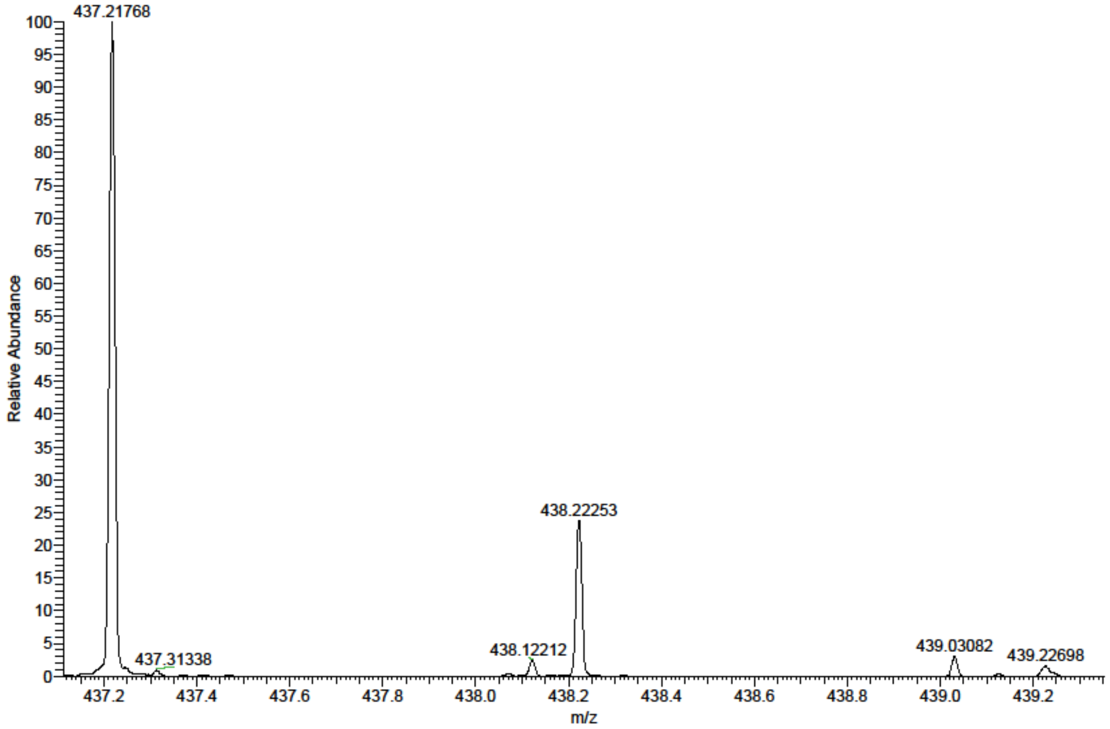

#### List of references
